# New insights from the combined discrimination of modern/ancient genome-wide shared alleles and haplotypes: Differentiated demographic history reconstruction of Tai-Kadai and Sinitic people in South China

**DOI:** 10.1101/2021.06.19.449013

**Authors:** Mengge Wang, Guanglin He, Xing Zou, Pengyu Chen, Zheng Wang, Renkuan Tang, Xiaomin Yang, Jing Chen, Meiqing Yang, Yingxiang Li, Jing Liu, Fei Wang, Jing Zhao, Jianxin Guo, Rong Hu, Lan-Hai Wei, Gang Chen, Hui-Yuan Yeh, Chuan-Chao Wang

## Abstract

Southern China was a region with mixed rice-millet farming during the Middle Neolithic period and also suggested to be the homeland of Tai-Kadai-speaking (TK) people. The archaeological evidence of animal and plant domestication has demonstrated that southern Chinese rice agriculturalists dispersed from the Yangtze River basin with the dissemination of TK, Austroasiatic (AA), Austronesian (AN) and Hmong-Mien (HM) languages. However, the formations of the inland TK-speaking people, central/southern Han Chinese and their relationships with Neolithic farmers from the Yangtze and Yellow Rivers (YR) basins are far from clear due to the limited sampling of South China. Here, we revealed the spatiotemporal demographic history of southern China by analyzing newly generated genome-wide data of 70 southeastern mainland TK speakers including Dong, Gelao and Bouyei and 45 southwestern Han Chinese together with comprehensive modern/ancient reference datasets. Southwest Han Chinese and Gelao demonstrated a closer genomic affinity to Neolithic YR farmers, while inland TKs (Dong and Bouyei) demonstrated a closer genomic affinity to coastal TK/AN-speaking islanders and Neolithic Yangtze farmers and their descendants. The shared genetic drift between inland TK/AN speaker highlighted a common origin of AN/TK groups, which may be descended from Tanshishan people or their predecessors (Xitoucun). Additionally, we found that inland TK/Sinitic could be modelled as an admixture of ancestral northern East Asian (ANEA) and ancestral southern East Asian (ASEA) sources with different proportions, in which the ANEA was phylogenetically closer to Neolithic millet farmers deriving from the YR Basin and the ASEA was phylogenetically closer to Coastal Neolithic-to-modern southern East Asians. Finally, we discovered genetic differentiation among TK people from southern China and Southeast Asia and obvious substructures between northern and southern inland Chinese TK people. The observed patterns of the spatiotemporal distribution of the northern and southern East Asian lineages in Central/southern China were also compatible with the scenario of bi-directional gene flow events from ANEA and ASEA. Conclusively, multiple lines of genomic evidence indicated millet farmers deriving from the YR basin and rice farmers deriving from the Yangtze River basin substantially contributed to the present-day mainland TK speakers and Central/southern Han Chinese, and formed the modern dual genetic admixture profile.

## Introduction

East Asia harbors more than a fifth of the global population and always has major genetic and paleoanthropological potentials since the landmass is much larger, its extensive longitudinal, latitudinal, and altitudinal breadth covers complex ecological and geographical environments, and it also shows a range of ethnic, linguistic and cultural diversity. The genetic findings and archaeological records of East Asia documented a complex demographic history of human movements and occupations, with the first archaic hominins arriving at least 1.6 million years ago and anatomically modern humans expanding into this area at least 50 thousand years ago (kya)[1–3]. The story from China largely influenced the origin, expansion and diversification of ancient populations as well as the innovation and development of material culture in East Asia, as it was principally the leading civilization in the world exerting its enormous prestige and influence on its neighbors. Geographic barriers such as mountains (e.g., the Himalayas, Qinling Mountains and the Altai Mountains), large river systems (e.g., Yellow and Yangtze Rivers), plateaus (e.g., Tibetan Plateau, Loess Plateau, Yunnan-Guizhou Plateau and Inner Mongolia Plateau), and deserts (e.g., Taklimakan Desert, Qaidam Desert and Junggar Desert) connect different biomes and may have also influenced the geographic spread of hominins, modern humans, animals and vegetative communities[4–6]. Southern China, the original birthplace of rice agriculture, occupied a key location in the southward spread of rice farmers to Southeast Asia, and was originally inhabited by the southern natives, including Tai-Kadai (TK), Hmong-Mien (HM), Austroasiatic (AA) and Austronesian (AN) speakers[7, 8].

Recently, multiple lines of archaeobotanical evidence from phytolith remains or starch grains have demonstrated that both dryland foxtail millet (also referred to as *Setaria italica L. Beauv*) and broomcorn millet (*Panicum miliaceum L*.) originated from Nanzhuangtou, Donghulin and Cishan in YR Basin and rainfed rice (*Oryza Sativa L*.) originated from Shangshan in Yangtze River Basin co-existed in Central/southern China from Neolithic periods[9, 10]. The patterns of mixed millet and rice farming in China revealed the northward spread of rice agriculture and the southward spread of millet agriculture mainly via three important north-south communication corridors[11]. However, there are still some debates focusing on the extent to which population movement propelled the spread of agriculture: whether human migrations/interactions or only cultural idea adaptation and language dispersal were accompanied by the spread of early farming lifestyle and changes in subsistence strategies? Paleoanthropological findings revealed multiple waves of rice farmers southward migrations associated with the dissemination of TK, AA, AN and HM languages and participated in the formation of modern Southeast Asians. A major focus of a genetic survey aimed to bridge the corresponding gaps in southern China based on population-scale data is on the way.

The TK or Daic languages, also referred to as Zhuang-Dong and Kra-Dai, cover a substantial area of East and Southeast Asia[12]. The high diversity of TK languages in southern China points to the origin of the TK language family in southern China. Over the past hundreds of years, the TK people gradually migrated southward, driven by various cultural, political, economic, environmental and other relevant factors, and became the majority inhabitants of Southeast Asia[8, 13, 14]. In light of the complex population history of TK-speaking groups and the important role played in the peopling of East and Southeast Asia, the analyses of genetic variations in TK-speaking populations have been conducted in several studies using different sets of autosomal[15–17], X-chromosomal[18, 19], Y-chromosomal[20, 21], mitochondrial[13, 14, 21–23] and genome-wide[7, 24] markers. Previous studies showed that the Proto-languages of TK and AN might be related[25], and TK people from Thailand are related to AN speakers based on mitogenome sequences[14]. The modeling of different demographic scenarios for Thai and Lao populations supports the spread of the ancestors of TK groups from southern China by demic diffusion[13]. Recently, He et al. have proved that Hainan TK-speaking Li survived as an isolated group for a long time and could be regarded as an unmixed proxy to model the admixture chronologies of mainland TK people and southern Han Chinese[7, 20, 26]. These genetic findings were generally based on insufficient sets of markers, sampling places, or only focused on TK speakers in Southeast Asia, and the genetic landscapes of mainland TK-speaking groups in China were roughly investigated mainly based on forensically available markers[16, 19, 27, 28]. Hitherto, much uncertainty remains regarding the origin, migration, admixture, and diversification of mainland TK populations and hence calls for detailed, systematic genome-wide analyses of mainland TK people.

Thus, to further shed light on the demographic history of mainland TK people and southern Sinitic speakers, as well as to elucidate the association between the formation of dual-mixed agricultures and genetic pools, we sampled and analyzed one Dong, one Bouyei, three Gelao populations, and three Han populations from Guizhou Province (**Figure 1A**, southwestern China) as well as collected a comprehensive modern and ancient reference datasets (including 149 Neolithic-to-Historic people from China, which was first merged to elucidate the spatiotemporal genetic history of East Asian). Guizhou is a mountainous province, lying at the eastern end of the Yungui Plateau, and its higher altitude area in the west and center, which is an ideal window for inspection of the genetic composition of millet-rice mixed farmers. Gelao and Dong languages belong to the Kadai language family, while the Bouyei language belongs to the Tai language family, and their ancestors can be traced back to ancient BaiYue people who once resided in southern China according to historical documents[29].

**Figure 1.**
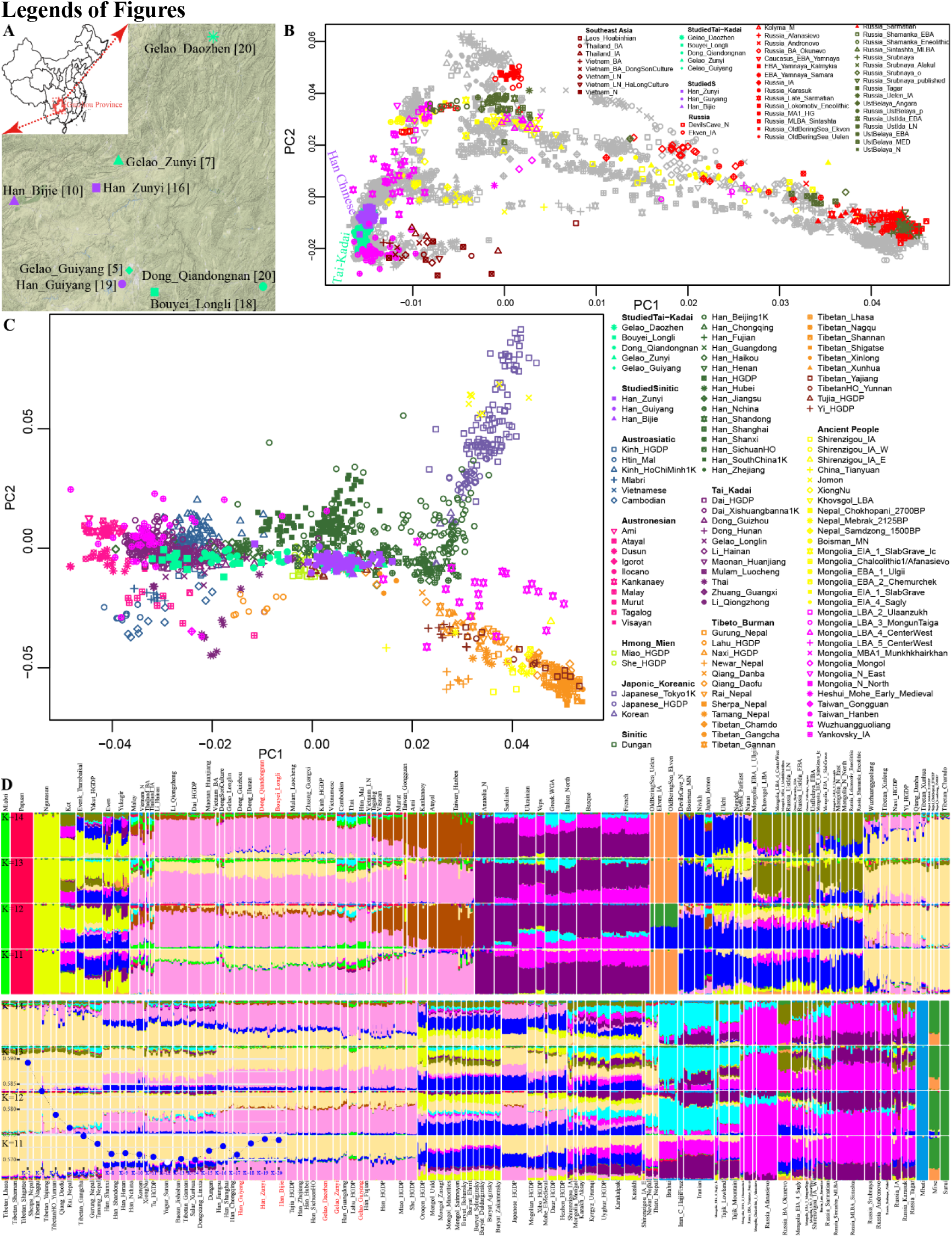
Patterns of genetic affinity and population structure of the included East Asian groups. (**A**) Geographical positions of all studied populations, Map was madeusing R package of ggmap; (**B**) Eurasian-based principal component analysis (PCA) was conducted based on the genetic variations of modern populations and Eurasian ancient people were projected onto it; (**C**) East Asian-based PCA was conducted based on the genetic variations of modern East Asians and East Asian ancient people were projected onto it; (**D**) ADMIXTURE results showed the ancestral composition of studied and reference individuals.

## Results

### Overview of genetic affinity and population structure

Patterns of genetic relationship inferred from the PCAs among Eurasian-wide populations revealed two obvious genetic clines: East-West cline and South-North cline (**Figure 1B and Supplementary Fig. 1**). The East-West cline was comprised of Tungusic-speaking Even and Evenk, Turkic, Uralic, Chukotko-Kamchatkan, Eskimo-Aleut, Indo-European, Caucasian-speaking populations and language-isolate Ket people. The South-North cline consists of AA, AN, TK, Sinitic, HM, Tibeto-Burman (TB), Koreanic, Japonic, Mongolic, Tungusic-speaking populations and language-isolate Nivkh group, starting with the AA and AN speaker at the southernmost tip and ended with the northernmost tip of Uralic-speaking Nganasan speaker and northeastern Siberians. Our newly reported Guizhou Han Chinese congregated at the Sinitic-related cluster and fall closer with Han populations from Hunan and Sichuan provinces and TK-speaking populations were scattered and generally fall within Sinitic-related and TK-related clusters. When projecting ancient samples onto the modern Eurasian-based PCAs, newly-genotyped Han Chinese showed a closer genetic affinity with Neolithic Wuzhuangguoliang individuals (~3,000 BCE) from the YR Basin[30] and newly genotyped TK-speaking people showed closer genetic relationships with Bronze Age Vietnam individuals[31],[32], Iron Age Gongguan and Hanben individuals (~1,400 BCE - 600CE) from Taiwan[30].

In the East Asian PCA plot focusing on the AA, AN, TK, Sinitic, HM, TB, Japonic, Korean-speaking populations and ancient East Asians (**Figure 1C**), we identified three meta-population clusters contributing to the genomic diversity of plotted populations: one represented by Japonic, Korean-speaking populations and Jomon samples from Japan[30, 31]; one by TB speakers (except Lahu and Tujia from HGDP), ancient northwestern and northern East Asians[30, 33–36]; and another one by AA, AN, TK, Sinitic, HM-speaking populations together with Iron Age Gongguan and Hanben people from Taiwan[30]. Studied Han populations overlapped with Chongqing Han, Sichuan Han and Han_HGDP, which also had a relatively close genetic affinity with Han Chinese from central and southern China, such as Guangdong, Fujian, Hubei, Jiangsu, Zhejiang and Shanghai. Our newly reported Bouyei and Dong samples clustered together and were surrounded by ancient Gongguan and Hanben individuals, displaying general signatures of greater affinity to southern Han Chinese, TK speakers, AA-speaking Kinh and Vietnamese. Compared with Bouyei and Dong, studied Gelao groups appeared to be shifted unexpectedly towards the direction of studied Han Chinese. Furthermore, we found that HM speakers clustered between our newly reported Bouyei, Dong, Han Chinese, and Gelao people.

The qualitative observations of model-based ADMIXTURE clustering identified the optimal K value of 12 with the lowest cross-validation error **(Supplementary Fig. 2**). The clustering pattern of involved populations was well corresponding to the linguistic affiliation (**Figure 1D and Supplementary Fig. 3**). At K=11, we observed a pink component that maximized in AN-speaking populations, ancient Gongguan and Hanben samples, followed by TK and AA speakers, and a pale-yellow component that was enriched in modern TB speakers, ancient Tibetan-related and Han-related individuals. New-generated populations were the mixed product of these above two sources with different proportions. At K = 12, a brown component maximized in AN speaker, ancient Gongguan and Hanben samples appeared, which separated AN-related groups from TK and AA speakers, suggesting the differentiated recent demographical history of those southern populations. Our newly-genotyped Han Chinese and Gelao people were grouped with other related Han populations, which were suggested to be a mixture of Tibetan-like and TK/AA-like components. The TK/AA-like component was also present at a high proportion of more than 50% in Dong and Bouyei people, while Tibetan-like and AN-like components accounted for relatively low proportions for Dong and Bouyei.

We further explored the patterns of genetic relationships of ancient and modern East Asians using the higher-density genetic variations from the 1240K dataset by including all Neolithic/Bronze-to-historic ancient populations from the Yangtze River and YR. Modern populations clustered into three groups as Han Chinese and Altaic/TB speakers, inland TKs, and island TKs respectively (**Figure 2A**). Eight newly-genotyped populations were located between northern Han and Li individuals and formed a genetic cline starting from Han and Gelao and ending with Bouyei. When we projected ancient populations from China, Mongolia, Russia, Japan, and Nepal onto the PCA plot, we observed the studied Han and Gelao groups were shifted along PC1 toward the YR farmers, while Bouyei and Dong were positioned closer to Neolithic/Bronze-Age southern East Asians (Xitoucun, Tanshishan and Hanben from Fujian and Taiwan). Model-based ADMIXTURE results among all Chinese ancients, newly-studied populations and corresponding neighbors showed that three predefined ancestral sources can best account for this gene pool of involved populations (**Figure 2B**). YR-like ancestry represented by northeast East Asian lineage also maximized in Neolithic (Mongolia_N_North) and Bronze Age Deer-Stone people (Khovsgol_LBA) from Mongolia Plateau. Blue ancestry was enriched in Middle Neolithic YR Wanggou and Xiaowu farmers, which also widely existed in both coastal (Shandong) and inland (Henan, Qinghai, Shaanxi and Inner Mongolia) Neolithic northern East Asians and modern East Asians and appeared with a relatively smaller proportion in ancients from West Liao River (WLR), Amur River (AR) and Mongolia Plateau. The ancestry of jacinth color maximized in Iron Age Hanben and Gongguan people was widely distributed among coastal Neolithic southern East Asians from Fujian province, Jomon people and modern East Asians. Modern central and southern East Asians including our newly-genotyped populations harbored two main types of ancestries (blue and jacinth), showing a dual structure in the genetic composition in line with the historic mixed farming system.

**Figure 2.**
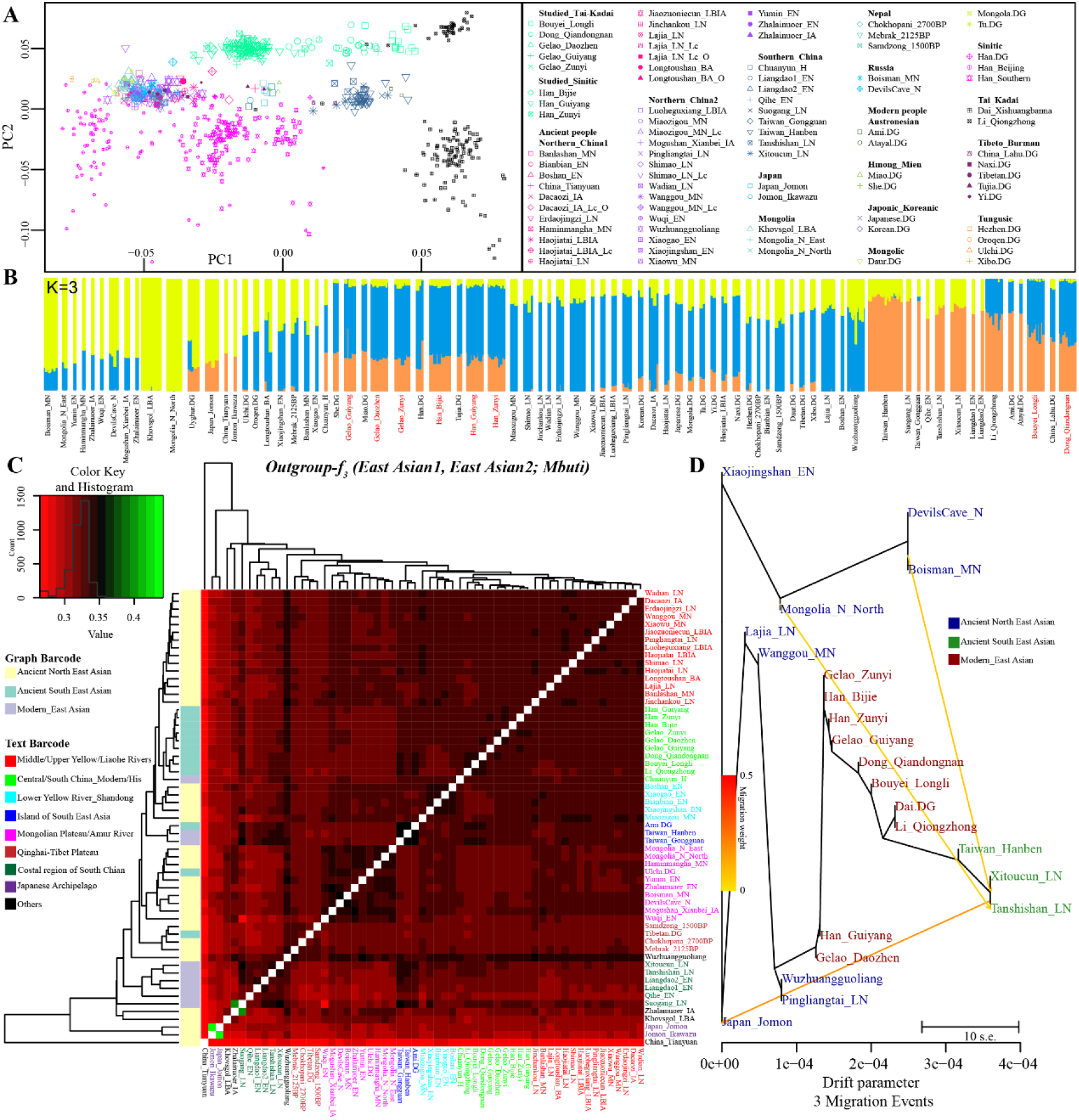
Genomic affinity between modern and ancient Asians. (**A**) PCA among modern East Asians. Ancient populations were projected onto the two-dimensional plots of modern genetic background; (**B**) Population clustering patterns of both modern and ancient populations revealed via model-based ADMIXTURE when three predefined ancestral sources were employed (K=3); (**C**) Pairwise outgroup-f_3_-statistics (Source1, Source2; Mbuti) for modern studied and ancient East Asians; (**D**) Maximum likelihood phylogeny among 10 modern and 12 ancient East Asian groups constructed using TreeMix with three migration events.

### Shared genetic drift and admixture signatures

To evaluate which modern or ancient samples had the most shared genetic drift with newly-studied populations, we first calculated outgroup-*f_3_*-statistics in the form *f_3_*(*X*, *Y*; *Yoruba*) using Human Origin Dataset (HO). In general, studied Han Chinese and Gelao populations shared more alleles with Neolithic Yellow River farmers and other Han Chinese, while TK speakers shared more alleles with TK-speaking groups and ancient Gongguan samples (**Supplementary Figs. 4~16**). We then performed high-resolution outgroup-*f_3_*-statistics based on the overlapped variants between our newly-genotyped data and 1240K dataset to further measure the allele sharing (**Supplementary Table 1**). When we focused on the shared genetic drift between eight newly-studied populations and Chinese Neolithic to historic ancient populations (**Figure 2C**), Han and Gelao showed the most shared genetic drift with inland northern East Asians related to YR farmers including Late Neolithic Pingliangtai and Haojiatai, while other inland TK speakers owned the closest relationship with Iron Age Hanben samples from Taiwan and modern island/coastal TKs, especially for Li in Qiongzhong, Hainan Island. Inland Bouyei and Dong also showed a close genetic relationship with YR farmers. We next performed admixture-*f_3_*-statistics of the form *f_3_*(*X*, *Y*; *Studied populations*) to explore potential source populations. Pairs of southern TK/AN groups with northern-TB-related groups can give significant negative *f_3_*-statistics in modeling our newly-genotyped Han, Gelao and Dong groups, but not in Bouyei (**Supplementary Figs. 13~16, and Tables 2~25**).

The allele sharing profiles revealed by *f_3_-statistics* have distinguished two ancestral populations highly associated with millet farming dispersal from North China and rice farming dispersal from Southern China for modeling our newly genotyped populations. Given that, we performed a series of *f_4_-statistics* to formally test the excess of alleles shared with any of the representative sources. Significantly negative Z-scores of *f_4_*(*Reference population1*, *Reference population2*; *Targeted populations*, *Mbuti*) in **Figures 3A~B, Supplementary Tables 26~29 and Figs. 17~24** showed that the targeted TK populations shared more alleles with modern southern East Asians (TK, AN, AA, HM and southern Han) and Coastal Neolithic/Bronze-Age southern East Asians[30, 37] than with northern East Asians, while Han and Gelao people harbored more northern East Asian-related derived alleles. Furthermore, to validate whether the eight studied populations were the direct descendants of YR or Yangtze farmers without additional admixture events, we conducted *f_4_*-statistics of the form of *f_4_*(*Yangtze*/*YR farmers*, *Studied TK*/*Sinitic*; *Reference populations*, *Mbuti*) to test if there was unbalanced allele sharing between reference populations with targeted populations when compared to proximate sources. We have not found any significantly negative signals in *f_4_*(*Ami*, *Dong*/*Bouyei*; *Reference populations*, *Mbuti*) (**Figure 3A**), which illustrated the genomic affinity between island AN and inland TKs. However, additional gene flow events among Gelao/Han Chinese populations were identified related to their predefined unique northern and southern sources. We also identified the northern East Asian-related genetic influence into Dong and Bouyei by the *f_4_*(*Neolithic southern East Asians*, *Dong*/*Bouyei*; *Reference populations*, *Mbuti*).

**Figure 3.**
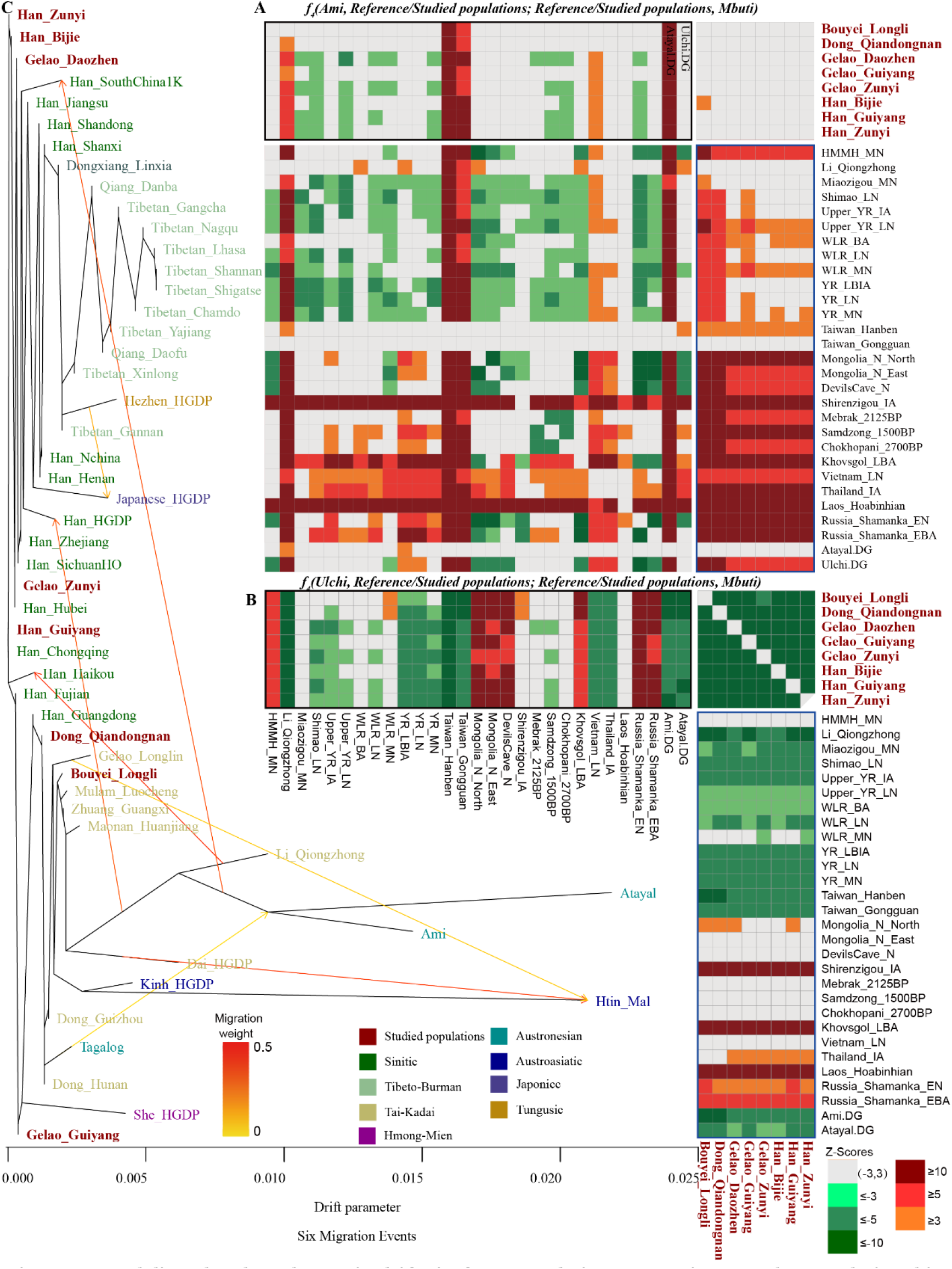
Modeling the shared genetic drift via four-population comparisons and tree relationships among modern populations. (**A**) The unbalanced allele sharing between reference populations with studied populations when compared to ASEA-related Ami revealed by f_4_(Ami, Reference/Studied populations; Reference/Studied populations, Mbuti); (**B**) The unbalanced allele sharing between reference populations with studied populations when compared to ANEA-related Ulchi revealed by f_4_(Ulchi, Reference/Studied populations; Reference/Studied populations, Mbuti); (**C**) Phylogenetic tree with maximum likelihood constructed with six migration events.

### Di-directional gene flow events between YR and Yangtze farmers since the early Neolithic period

To further reveal the detailed patterns of temporal and spatial differences of shared alleles between YR and Yangtze River populations of different prehistoric periods, we made a comprehensive population comparison among all available modern/ancient East Asians. We first conducted *f_4_*(*YR/Yangtze River 1*, *YR/Yangtze River 2*; *East Asians*, *Mbuti*) to inspect the temporal genetic differences and then conducted *f_4_*(*Contemporaneous Yangtze Farmers*, *Contemporaneous YRs*; *East Asians*, *Mbuti*) to explore the spatial differences of shared ancestry between Yangtze River and YR people. Concentrating on the temporal gene pool of the YR populations, we identified the increased modern AN/southern Han-related ancestry in the transitions from Early Neolithic to Middle Neolithic and Middle to Late Neolithic periods (**Supplementary Figs. 29~38**). We could also identify the increased shared ancestry from Coastal Neolithic southern East Asians in the aforementioned population transitions. Compared with the Late Neolithic Wadian, Haojiatai and Pingliangtai (**Supplementary Figs. 39~41**), local Bronze Age and Iron Age populations shared equal extent of alleles with studied populations and other southern East Asians (**Supplementary Figs. 39~41**). If we used the samples from Late Neolithic Shimao, Lajia and Jinchankou sites as the comparative base, we could also identify more shared southern modern and ancient ancestry in Late Bronze and Iron Age populations from Luoheguxiang and Jiaozuoniecun (**Supplementary Figs. 42~48**). Overall, the northern gene pool gradually absorbed the gene materials from the southern migrants.

For the temporal genetic changes in Southern China, we employed the Coastal southern East Asians from Early Neolithic (Qihe and Liangdao) to Iron Age (Hanben and Gongguan) and Historic period (Chuanyun) to conduct *f_4_*(*ancient southern East Asians*, *ancient northern East Asians/Tibet Plateau/Siberia/Japan*; *Studied inland TK/Sinitic*, *Mbuti*). As shown in **Supplementary Figs. 49~51**, we identified significant increased YR and inland Han/TK-related ancestry in the late Neolithic/Iron Age/Historic populations related to the Early Neolithic southern East Asians (Qihe and Liangdao). The most significant signal was found in sample from Iron Age Hanben and Gongguan and Late Neolithic mainland Tanshishan and Xitoucun sites. Compared to the late Neolithic gene pool (Suogang, Xitoucun and Tanshishan, **Supplementary Figs. 52~54**), Iron Age Gongguan and Hanben from Taiwan and Historic Chuanyun also harbored more inland AN/Han and YR-related ancestry. The same increased genetic signal was also identified during the transition from the Iron Age (Hanben) to historic time (Chuanyun, **Supplementary Figs. 55~56**).

Finally, we also explored the spatial distribution of shared genetic drift via *f_4_*(*contemporary Neolithic northern East Asians*, *Neolithic southern East Asians*; *Studied inland TK/Sinitic and East Asian ancients*, *Mbuti*). As shown in **Supplementary Figs. 57~67**, the north-south genetic structure among East Asians could be identified in all three Neolithic periods (early, middle and late stages). Here, we also pointed out that the modern Han/inland TK-speaking Gelao genetically shifted toward with Neolithic northern East Asians, while other TK and AN-speaking populations shared more alleles with Neolithic southern East Asians.

### Population substructure among TK-speaking populations

To present a fine-scale pattern of the genomic substructure within TK speakers, we performed admixture-*f_3_*-statistics and *f_4_-statistics* for all obtainable TK-speaking groups from Southeast Asia and southern China. The non-significant Z-scores of *f_3_*(*Reference population1*, *Reference population2*; *TK speakers*) suggested that Hainan Li, Yunnan Dai, BoY, CoLao, Gelao (Longlin), LaChi, Maonan, Mulam, Nung and Tay (**Supplementary Tables 34~48**) had not received substantial gene flow from current reference populations and supported that these TK people had undergone a unique genetic isolation or potential genetic drift. However, the four TK groups Dong_Guizhou, Dong_Hunan, Thai, and Zhuang_Guangxi have significant negative admixture-*f_3_*-statistics when using a northern and a southern population as a potential pair of sources (**Supplementary Tables 49~52**), suggesting recent north-south admixtures in those groups.

We used a series of *f_4_-statistics* to further explore the shared genetic drift between TK and their neighbors. The TK-speaking populations showed a close genetic affinity with each other and were genetically like HM, AA, AN speakers, or ancient southern East Asians (**Supplementary Table 53**). The significantly negative values of *f_4_*(*Reference populations*, *TK populations*; *Yangtze River related populations*, *Mbuti*) further supported that TK-speaking groups derived their main ancestry from Yangtze River related farmers, a strong genetic affinity observed in **Supplementary Table 54**.

We subsequently carried out symmetrical *f_4_-statistics* of the form *f_4_*(*TK population 1*, *TK population 2*; *Reference populations*, *Mbuti*) to explore the genomic heterogeneity among all studied TK-speaking populations (**Supplementary Table 55**). Significant-*f_4_* values observed here suggested genetic differentiation among different TK-speaking populations. The ANEA groups showed more allele sharing with Gelao people than they did with other populations and they also showed more allele sharing with Bouyei, CoLao, Dong and Zhuang than they did with Li, Dai or Nung groups. We observed that Huanjiang Maonan harbored more northern East Asian and Siberian ancestry related to northern farmers and Hunter-gatherers (Boisman_MN and DevilsCave_N) when compared with Dai. Mulam harbored more YR farmer related ancestry compared with Dai. The TK groups from Southeast Asia bored relatively more AN/AA-related ancestry compared to their northern counterparts. The significant negative Z-scores of *f_4_*(*TK populations*, *Dai*; *Reference populations*, *Mbuti*) suggested that AN-speaking Ede, Giarai, AA-speaking Mlabri, Htin_Mal or Kinh showed more allele sharing with Dai than they did with Mulam, Dong, or Tay. The ANEA-related groups showed the most shared alleles with Gelaos (from Daozhen, Guiyang, Longlin or Zunyi) and Dong (from Hunan, Guizhou, Qiandongnan), Longli Bouyei and Guangxi Zhuang, while ASEA-related groups possessed the most shared derived alleles with Dai, Li and TK-speaking populations in Southeast Asia. The studied TK-speaking groups could not be modeled as descending directly from the ASEA or ANEA because of the additional gene flow events from AA-related sources into southern TK people in Southeast Asia and from millet farmers into northern TK people in China.

### Inferring the phylogenetic relationship and estimating the admixture proportions

The patterns revealed by the Fst matrix (**Supplementary Table 56**) suggested that newly-studied Han Chinese and Gelao groups showed a close genetic affinity with each other and with neighboring northern East Asians, while Dong/Bouyei showed a close genetic affinity with AA-speaking Kinh and other southern TK speakers. Fst matrix among ancient East Asians (**Supplementary Table 57**) showed that all studied populations had the smallest genetic distance with Hanben and then followed by Neolithic YR farmers, which was consistent with the observed patterns of genetic relationship based on the outgroup-*f_3_*-statistics.

Considering the potential gene flow events between diverged human populations, we performed two graph-based models for building phylogenetic trees in treemix software. As shown in **Figure 2D**, phylogenetic relationships with three admixture events among 12 Neolithic-to-Bronze Age East Asians and 10 modern populations showed three lineages: northeast lineage consisted of Neolithic Boisman, DevilsCave and Mongolian, which also showed a close relationship with Early Neolithic Coastal northern East Asian (Xiaojingshan_EN), YR lineage comprised of Middle/Late Neolithic farmers (Lajia, Wanggou, Pingliangtai and Wuzhuangguoliang), and Yangtze River lineage comprised of Tanshishan, Xitoucun and Hanben. Jomon population was situated between northeastern lineage and YR lineage and acquired an additional gene flow from the Yangtze River lineage, which was consistent with the connection of coastal population migrations and peopling of the Japanese archipelago. Besides, we also identified two gene flows from the northeast lineage to Yangtze lineage, which showed historic population movements between northern and southern East Asians. Modern populations clustered together, and TK people showed close phylogenetic relationships with Yangtze River lineage, while others showed closer relationships with YR farmers. Furthermore, we constructed the phylogenetic topology in treemix software with the maximum likelihood when setting the value of migration edge weight to 6 (**Figure 3C**) among modern populations. We found that studied Han Chinese, Zunyi Gelao and Daozhen Gelao were phylogenetically closer to southwestern and east-central Han Chinese, and Guiyang Gelao showed a closer genetic affinity with HM-speaking She, Guangdong Han and Fujian Han, while Qiandongnan Dong and Longli Bouyei were phylogenetically closer to reference TK speakers. We observed the gene flow from AN-speaking Tagalog into AN-speaking Ami and Atayal, from Ami/Atayal into Han Chinese in southern China, from TK-speaking Dai and Longlin Gelao into AA-speaking Htin_Mal, from Qiongzhong Li into Haikou Han and Han_HGDP, and from Hezhen_HGDP into Japanese. However, no specific gene flow into studied Han Chinese or TK speakers was observed.

Based on the sharing profiles disclosed by the *f_3_* and *f_4_* analyses, as well as the tree-based phylogenetic relationship, we next built admixture graphs for each studied group to depict a demographic history of population divergence and gene flow (**Figure 4**). Here, we found all studied populations could be fitted via the admixture of the Yangtze and YR-related ancestral populations. We estimated that studied Han Chinese had approximately 57~62% ancestry from the ancestor of the Wanggou_Yangshao (~3,550-3,050 BCE, which was the main component of the aforementioned YR_MN), and approximately 38~43% ancestry from the ancestor related to the Iron Age Hanben samples (hypothesized decedents of Yangtze farmers). Furthermore, studied Han Chinese also had approximately 47~54% ancestry from the ancestor of the Upper YR Lajia_Qijia (~2,050-1,850 BCE, that was aforementioned Upper_YR_LN), and approximately 46~53% ancestry from the Hanben related ancestor. Studied Gelao groups derived about 50~60% of their ancestry from the Wanggou_Yangshao-related lineage and about 40~50% from Hanben-related lineage, which also had about 40~51% ancestry from the ancestor of the Upper YR Lajia_Qijia and about 49~60% ancestry from the ancestor of the Hanben. Qiandongnan Dong had ~64% or ~73% ancestry from this same Hanben ancestor and ~36% ancestry from a source connected with the Wanggou_Yangshao or ~27% ancestry from a source related to the Upper YR Lajia_Qijia. For Longli Bouyei, the major component was closely associated with the Hanben (~78% or ~85%), with additional Wanggou_Yangshao-related ancestry (~22%) or Upper YR Lajia_Qijia-related ancestry (~15%). This suggested a genetic cline of increasing Hanben-related ancestry stretching from TK speakers in the north of the sampling region to TK speakers in the south.

**Figure 4.**
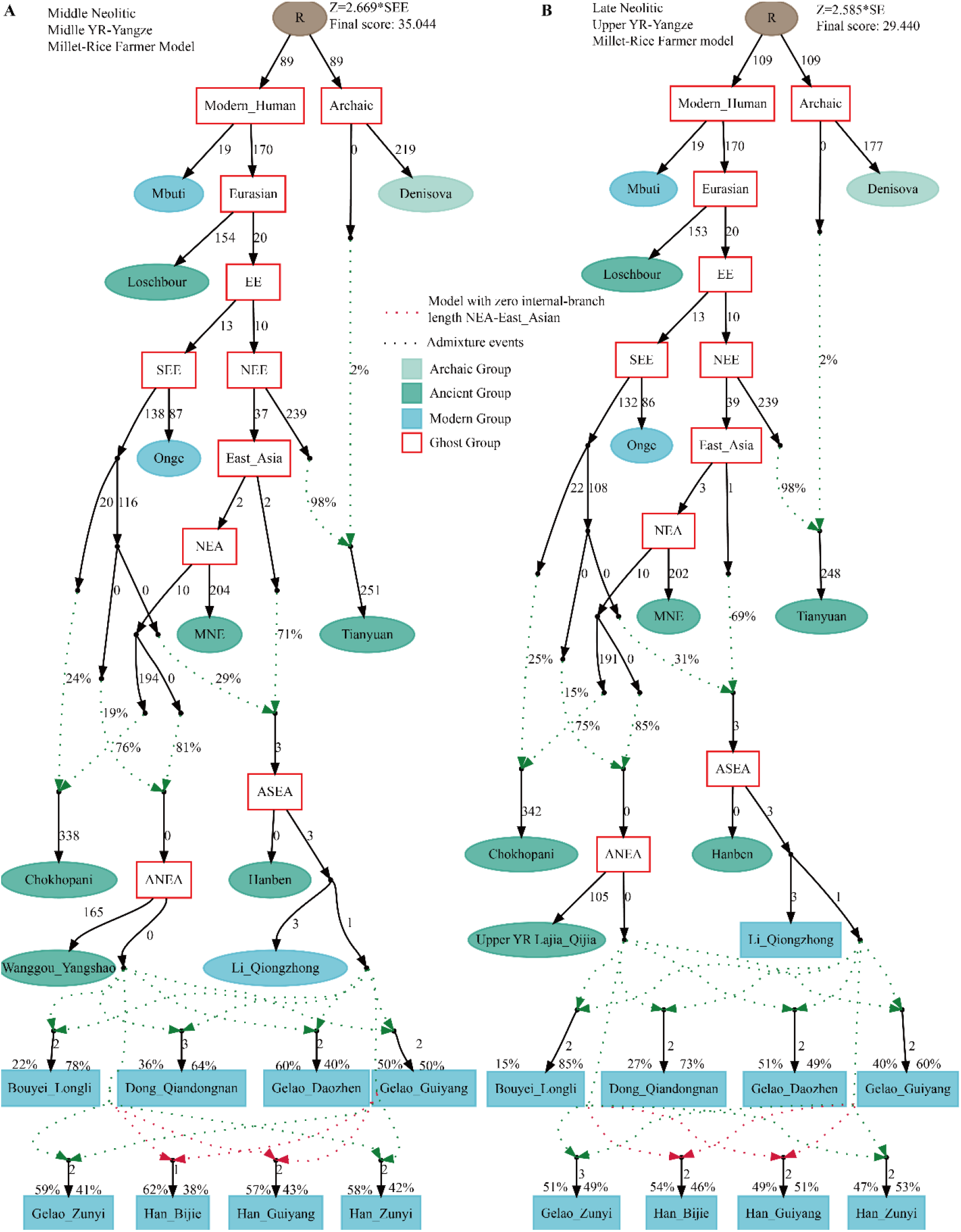
qpGraph modeling of all studied populations. (**A**) Middle Neolithic Middle Yellow River framer (Wanggou_Yangshao) was used as the Ancestral northern East Asian; (**B**) Late Neolithic Upper Yellow River framer (Upper YR Lajia_Qijia) was used as the Ancestral northern East Asian. We started with a skeleton tree that fits the data with Denisova, Mbuti, Loschbour, Onge and Tianyuan. We retained graph solutions that provided no differences of |Z| > 3 between fitted and estimated statistics. The genetic drift value is the actual test value multiplied by 1000.

We also systematically performed various *qpAdm-based* admixture models without an exact phylogeny by applying different northern source populations. Two-way admixture models with one ASEA related source as TK-speaking Li and the other as YR-related ANEA sources worked out well to explain the studied populations (p_rank1 > 0.05) (**Figure 5**). In general, Bouyei and Dong harbored a higher proportion of ASEA-related ancestry (0.653~0.908), and these two-way admixture models revealed a North-South cline of decreasing ANEA-related ancestry in northern Gelao/Han and increasing ASEA-related ancestry in southern Dong/Bouyei.

**Figure 5.**
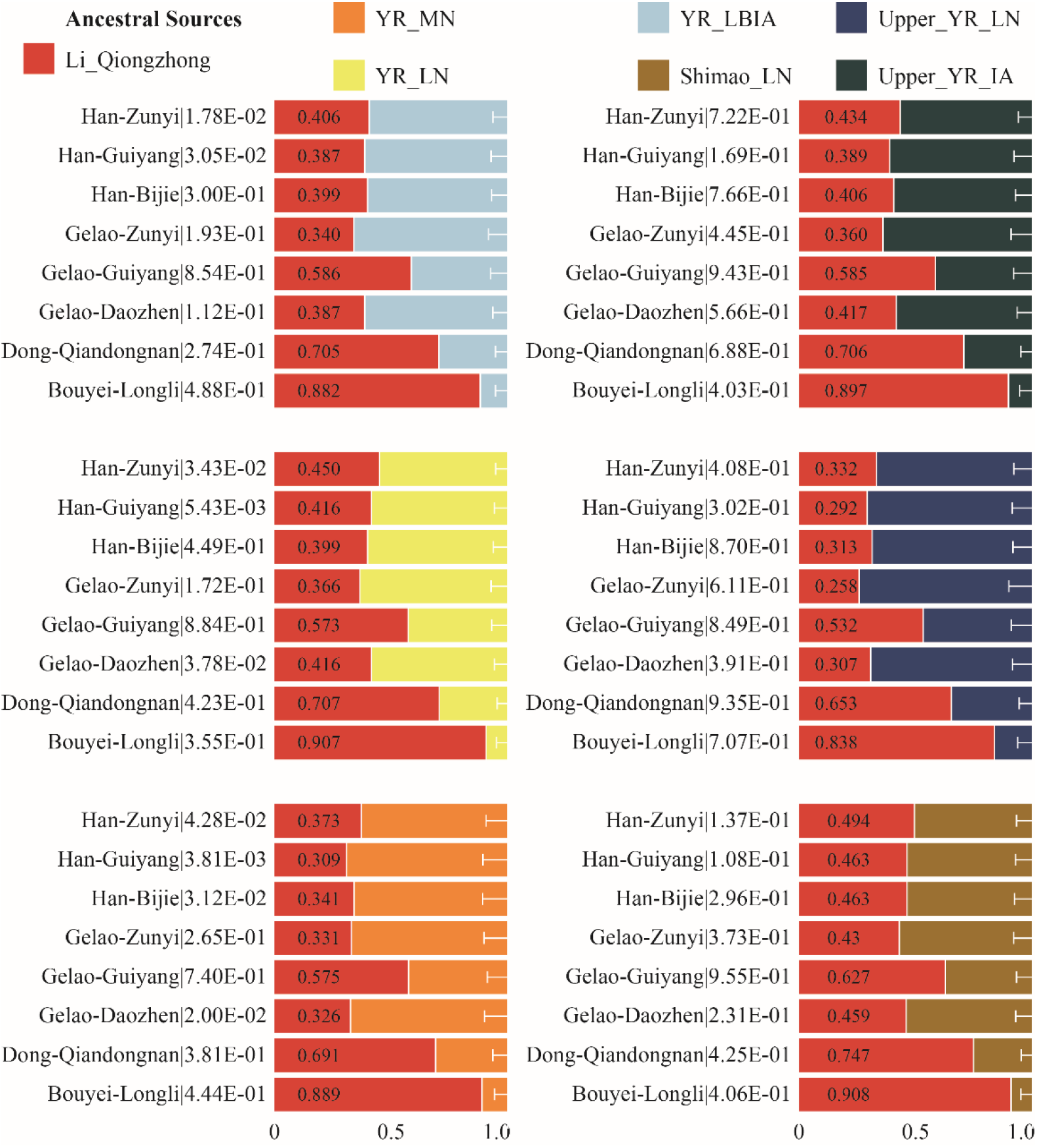
Admixture modeling of all newly studied populations using the qpWave and qpAdm approaches with different represented proximate ancestral populations. The bar denotes the standard error. The ancestry proportion of southern source were marked as the number. The p-values of qpWave when using the two-way admixture model were noted at the behind of the population name.

### Finer-scale population substructure inferred from more complex admixture models and sharing haplotypes

To further explore the variation of ancestry proportions among studied Guizhou populations and other Sinitic- and Tai-Kadai-speaking populations, we conducted pairwise *qpWave* analysis. We used eleven worldwide populations as the right reference populations, including Mbuti, Ust_Ishim, Kostenki14, Papuan, Australian, Mixe, MA1_HG, Onge, Atayal, Mongolia_N_East and Yamnaya_Samara (**Figure 6A**). This type of allele-based analysis was performed to test whether the two studied populations (Right_i_, Right_j_;) were descended from one recent common ancestor which separated from the common ancestors of a set of included outgroups (Outgroupk, Outgroupl). Here, we observed genetic substructure among populations, which are consistent with being pairwise clades within geographically/ethnically close populations compared with this relatively distant set of Outgroups. To explore the subtle population substructure among studied populations and all Tai-Kadai-speaking populations, we used a more powerful set of outgroups to re-run pairwise *qpWave* analysis (**Figure 6B**). Additional seven East Asians were added in the right populations (Bianbian_EN, Wuqi_EN, Qihe_EN, Liangdao2_EN, Yumin_EN, Chokhopani_2700BP and Xiaowu_MN). Guizhou Gelao showed a differentiated genetic history with Longlin Gelao formed a clade with Guizhou Han. While studied Dong formed a clade with geographically close Guizhou Dong, and Hunan Dong and Longli Bouyei shared a close relationship with southern Tai-Kadai speakers from Guangxi and Southeastern East Asian. Population substructure was further confirmed in the three-way admixture *qpAdm*-based ancestry proportion estimation (**Figure 6C~E**). Here, we used spatiotemporally Yellow River basin farmers, inland Southern East Asians and Coastal Southern East Asians as three sources. We obtained 901 well-fitted three-population admixture proportions and found Han Chinese and Gelao harbored more northern East Asian related ancestry. Focused on the ancestry amount of southern East Asians, we observed that southern Tai-Kadai-speaking populations obtained approximate gene flow from two used southern sources.

**Figure 6.**
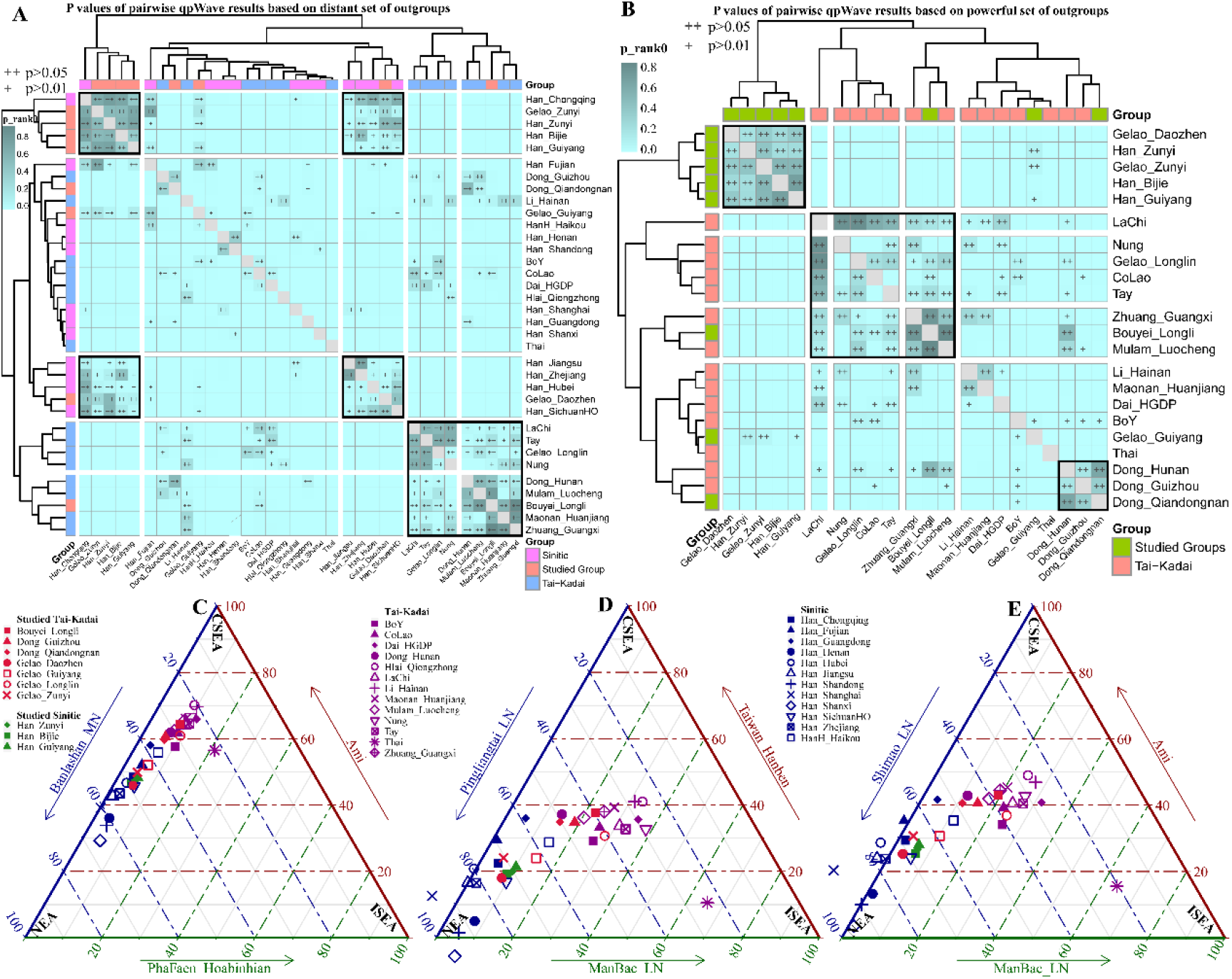
The genetic difference between Tai-Kadai speakers and other East Asians. (**A**) Pairwise qpWave results among the studied populations and referenced Sinitic/Tai-Kadai-speaking populations based on the relatively distant set of outgroups. (**B**) Pairwise qpWave results among the studied populations and referenced Tai-Kadai-speaking populations based on the powerful set of outgroups. (**C~E**) Ancestry composition of three-way admixture qpAdm-based results among Tai-Kadai and Sinitic speakers. Spatiotemporally diverse populations from Yellow River basin, coastal region of southern East Asian and inland region of southern East Asian. NEA: Northern East Asian; ISEA: inland Southern East Asian; CSEA: Coastal Southern East Asian.

We also explored the finer-scale population structure based on the genome-wide patterns of shared haplotype chunks based on a dataset with higher SNP density (600K). We phased the haplotype fragments of 328 people from Tai-Kadai, Sinitic and Tibeto-Burman language families. The genetic relationship based on the independent SNP frequency showed five clusters: Hainan Tai-Kadai Halai, Guizhou Tai-Kadai Bouyei and Dong, Tibeto-Burmans, Hainan Sinitic Hans, and Guizhou mixed cluster included Gelaos and Hans (**Figure 7A**). Patterns of shared ancestry inferred from haplotype data not only confirmed the aforementioned clustered, but also revealed population subclusters among populations from one allele-based cluster, such as genetic differentiation between Bouyei and Dong (**Figure 7B~D**). We reconstructed the population relationships based on the individual pairwise coincidence and found that two main branches and four subbranches. Guizhou Bouyeis shared the most ancestry chunks with Dongs, followingly harbored a close genetic relationship with Hans and inland Gelaos. Compared with Tibeto-Burman-speaking populations, all included Hainan and Guizhou Tai-Kadai-speaking populations shared more common ancestry or more gene flow events (**Figure E~F**). Our observed patterns of shared haplotypes were consistent with the shared ancestry inferred from the model-based ADMIXTURE result (**Figure 7G**), TreeMix-based phylogenetic relationship (**Figure 7H**) and pairwise Fst distance (**Figure 7I**). IBD fragments also confirmed the status that genetic heterogeneity between Tai-Kadai and Tibeto-Burman people and even reveal subtle genetic differences between Guizhou Tai-Kadai people and Guizhou Hans, as well as Guizhou Tai-Kadai people and Hainan HaLai (**Figure 7J**). We finally explored the natural selection signals (**Figure 7K**) based on the shared haplotype heterogeneity and length. Results from the iHS revealed natural signals associated with the susceptibility of clinical diseases or metabolism pathway-regulated genes.

**Figure 7.**
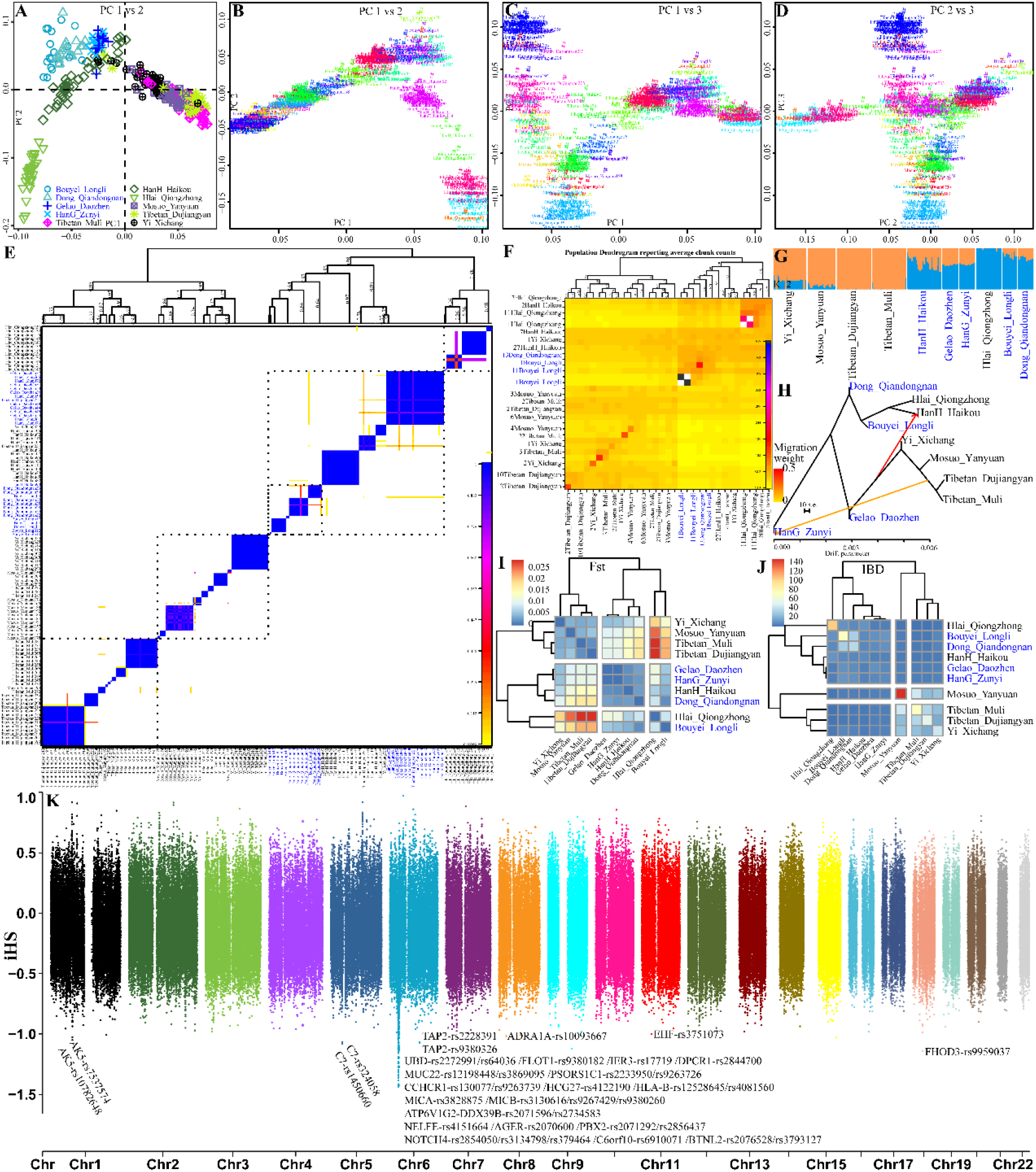
Shared ancestry between Tai-Kadai-speakers and their adjoin populations from southern China inferred from haplotype chunks. (**A**) PCA result based on the shared alleles among 328 southern Chinese people from 10 Sino-Tibetan or Tai-Kadai-speaking populations. (**B~D**) Two-dimensional plots of analyzed individuals based on their shared ancestry fragments. (**E**) Pairwise coincidence among 328 people based on the co-ancestry coefficient based on the ChromoPainter-based shared haplotype patterns. (**F**) Population dendrogram based on the average chunk counts of genetically homogeneous populations. Model-based ADMIXTURE (**G**) and TreeMix (**H**) results of haplotype-analyzed datasets. (**I**) Pairwise genetic distance among 10 populations shared three genetic branches. (**J**) Average shared population-level IBD fragments showed two branches among 10 included populations.

### Investigating the genetic continuity and admixture based on uniparental markers

Among the maternal haplogroups in our studied Han Chinese, subclades of southern prevalent haplogroups (B4, B5, B6, F1, F2, M7, M33, M74 and R9) accounted for the highest frequency (60.00%), and some subclades of northern haplogroups, for example, A14, A5, C7, D4, D5, G2, G3, M8, M9, M10 and Z, also showed high frequencies (40.00%). As for paternally inherited haplogroups, we identified high frequencies (88.46%) of Han Chinese characteristic sub-haplogroups of O1a1a*, O1b1a1*, O2a1c1a1a1a*, O2a2a1a2a1a2 and O2a2b1a* in studied Han Chinese. Besides, subclades C2b1b1a, N1a3~ and Q1a1a1 were sporadically distributed in targeted Han populations.

All 32 Gelao individuals could be assigned into 30 specific matrilineal subclades, haplogroup D4 was the most prevailing haplogroup (21.88%), followed by F1 (18.75%), M7 (12.50%), F2 (9.38%) and B5 (6.25%), other haplogroups (A5, A15, B4, B5, D2, G1, G2, M8, M10, M12 and N9) were sporadically distributed in Gelao groups with relatively lower frequencies. Generally, the prevailing haplogroups in southern China and Southeast Asia (B, F, M7 and M12) accounted for 53.13% of the Gelao maternal gene pool, and the predominant haplogroups in North China (A, D, G, M8, M10 and N9) accounted for 46.87%. Furthermore, 15 Gelao male individuals were assigned into 15 terminal sub-haplogroups, haplogroup O2a was the most prevailing haplogroup with a frequency of approximately 53.33%, haplogroups O1 and C2 tied for second with the frequency of 20.00%, respectively, and only one Daozhen Gelao was assigned to haplogroup Q1a1a1.

We noticed that subclades of southern prevailing haplogroups (B4, B5, F1, F3, F4, M7 and R9) accounted for 85.00% of the Qiandongnan Dong maternal gene pool, and subclades of southern prevailing haplogroups (B4, B5, F1, M7, M74 and R9) accounted for 66.67% of the Longli Bouyei maternal gene pool. The dominant paternal haplogroup in Qiandongnan Dong was O1 (37.50%) and O2 (37.50%), and in Longli Bouyei was O2 (42.86%). Moreover, we identified characteristic lineage C2 and characteristic lineage N1 in our Qiandongnan Dong samples, and characteristic lineage C1 and Tibetan characteristic lineage D1a1a were found in Longli Bouyei individuals.

## Discussion

The demographic history of the TK speakers and spatiotemporal genetic history of ancient East Asians are ones marked by repeated mixing of diverse southern and northern East Asian gene pools. However, rather than simple waves of migration, divergent events and genetic admixtures among the TK people and ancient East Asians have been tangled and variable, which has provoked debates over the past decades[7, 21, 38–40]. We have submitted the first genome-wide SNP data of TK-speaking Li islanders in the previous study[7] and provided genomic evidence that Li islander could be adopted as an unmixed proxy to model the demographic events of mainland TK speakers and southern Han Chinese. However, more genetic surveys for inland TK people needed to be conducted, which can provide additional insights into the formation process of southern Chinese populations. Thus, in this study, we have generated and analyzed genome-wide SNP data from 70 mainland TK individuals and 45 surrounding Han Chinese and comprehensively co-analyzed with modern and ancient East Asians, and we found an explicit genomic admixture pattern regarding TK speakers and southern Han Chinese.

### Genomic affinity among Guizhou Sinitic/TK speakers and northern/southern Neolithic farmers

We found extensive genetic diversity, geographic affinity and genetic differentiation among studied TK groups in the PCA, that is, Bouyei and Dong were genetically most like present-day AA, AN, TK speakers from the southmost tip of China and Southeast Asia. We also found their closer genetic affinity with Coastal Neolithic Southern East Asians based on *f_3_* and *f_4_*-statistics, represented by Neolithic Tianshishan and Xitoucun people, as well as the ancient Taiwan Hanben and Gongguan individuals. While TK-speaking Gelao groups showed a closer genetic affinity with the southern Han Chinese and a closer relationship with the farmers originated from dryland millet agriculture from YR, which showed an additional genetic flow event from millet farmers for Guizhou Gelao. Studied Han Chinese were genetically most like southwestern Han and Neolithic Pingliangtai and Haojiatai individuals from the YR Basin, followed by the Coastal Neolithic northern East Asian (Xiaojingshan) and aforementioned Coastal Neolithic southern East Asians. The shared drift revealed by HO-based and 1240K-based outgroup-*f_3_*-statistics also shed light on the findings observed in PCAs and ADMIXTURE. Additionally, the genetic affinity between TK and AN speakers and significantly negative Z-scores of *f_4_*(*East Asians*, *Studied TK speakers*; *Taiwan ancients/AN speakers*, *Mbuti*) were in keeping with the linguistic affinity between the TK and AN language families[41] and supported the hypothesis that TK and AN groups were descended from a common ancestral population related to Fujian Neolithic rice people. Further supporting genomic evidence obtained from the largest shared genetic drift in *f_3_*(*mainland/island TK*, *ancient southern coastal/island East Asian/AN*, *Mbuti*) and close phylogenetic relationship from *Treemix/qpGrapch* frameworks. Our genome-wide evidence of the genomic affinity between mainland/island TK and AN and Coastal Neolithic southern East Asian populations supported that the common ancestor of the Proto-AN and Proto-TK speakers originated from mainland southern China and then it disseminated from Taiwan, the island of Southeast Asia, Southwest Pacific and African island of Madagascar via the marine route, and it also spread southwestward via the inland route from southern China to Southeast Asia with the rainfed rice agriculture dissemination. Here, we provided one genomic-based case for the co-dispersal of language and agriculture.

### Genetic continuity and proximate ancestral populations of southern East Asians

The shared profiles uncovered via ADMIXTURE, *f_3_*-statistics and *f_4_*-statistics revealed that the primary ancestry of the targeted TK and Han populations was characterized by two proxy sources – ANEA and ASEA. The ANEA was mainly represented by millet farmers deriving from the YR Basin, also strongly associated with Tibetan-like individuals and Neolithic hunter-gatherers from Mongolia Plateau and Russia Far East. The reconstruction of the deep history of East Asia[30, 42] documented ancient millet farmers, termed as YR Ghost Population, that was a plausible vector for the spread of Sino-Tibetan languages and mixed with southern agriculturalists on the central plain to form the ancestors of Han Chinese. The ASEA was mainly represented by the modern proxy of Qiongzhong Li, AN-speaking Ami, Atayal or ancient rice farmers (islanders of Hanben and Gongguan and Coastal Neolithic Tanshishan and Xitoucun people) deriving from the Yangtze River Basin. Wang et al. has shown that ancient Hanben and Gongguan individuals were consistent with being nearly direct descendants of Yangtze Valley’s first farmers who were termed as Yangtze River Ghost Population and also obtained additional gene flow from northern East Asian farmers[30]. The population-scale genomic composition of studied TK speakers highlighted that the unmixed Qiongzhong Li could be used as the proxy of Proto-TK speakers[7] and provided supporting evidence that ancient rice farmers originating from the Yangtze River Basin had contributed their ancestry to present-day TK-speaking populations. Furthermore, the strongly negative signals in *f_3_*(*Longli Bouyei/Qiandongnan Dong*, *Tibetan/YR-farmer-related populations*; *Studied Han Chinese/Gelao people*) revealed that Longli Bouyei or Qiandongnan Dong also could be used as proxy source populations, suggesting that the ancestor of coastal TKs has participated in the formation of inland TK and southern Han Chinese. Our findings of genetic continuity/admixture between studied populations and ANEA/ASEA resonated with the ancestral profiles of two-way admixture revealed by *qpAdm/qpGraph*-based models. Furthermore, the gene flow from ANEA into studied Han Chinese was evident in the northern dominant maternal lineages A5, A14, C7, D4, D5, G2, G3, M8, M9, M10 and Z and paternal Y-chromosomal lineages O2a2b1a*, C2b1b1a, N 1a3~ and Q1a1a1. The formation of studied Han Chinese seemed to receive great influences from southern Chinese minorities with the presence of maternal lineages B4, M7, M33, M74 and R9 and paternal lineages O1a1a*, O1b1a1* and O2a2a1a2*. Similarly, we identified a series of motif matrilineal lineages A5, A15, C7, D2, D4, D5, G1, G2, M8 and N9, and Y-chromosomal lineages C2*, O2a2b1a*, N1 and Q1a1a1 reflecting the gene flow between targeted TK speakers and ANEA. This observed uniparental genetic landscape was consistent with the previously reported demic diffusion model of the formation of modern East Asians[43].

### Genetic heterogeneity among TK-speaking groups

We observed that newly-studied Gelao groups tended to be shifted unexpectedly in the direction of studied Han Chinese and reference northern Han Chinese and Inland Neolithic northern East Asians from YR (both lower and middle YR Yangshao and Longshan ancients and Upper YR Neolithic Qijia farmers), while Longli Bouyei and Qiandongnan Dong tended to be clustered with southern references: including TK speakers, AA-speaking Kinh, Vietnamese together with ancient Gongguan and Hanben individuals and Coastal Neolithic southern East Asians. Results of ADMIXTURE analysis showed that newly-genotyped Gelao people could be modeled as a mix of a similar proportion of Tibetan-like and TK/AA-like components, and TK/AA-like component was present in Qiandongnan Dong or Longli Bouyei with a significantly higher proportion (> 50%). Furthermore, values of *outgroup-f_3_-statistics* revealed that studied Gelao populations shared most derived alleles with Neolithic northern East Asian, while Qiandongnan Dong and Longli Bouyei showed more allele sharing with southern Chinese ancients, and modern AN. Specifically, the results of symmetry *f_4_*(*Studied population1*, *studied population2*; *Source populations*, *Mbuti*) demonstrated that ANEA-related populations showed more derived alleles with Gelao people, and ASEA-related populations showed more ancestry with Qiandongnan Dong and Longli Bouyei relative to their counterparts, which was consistent with the genetic patterns revealed by PCAs. Observations of *qpAdm/qpGraph-based* admixture models highlighted the differentiated shared profiles revealed by PCAs, ADMIXTURE analyses and outgroup-*f_3_-statistics*, which showed a genetic cline of increasing ASEA-related ancestry stretching from northern TK people to southern TK people. Furthermore, the genetic findings of admixture-*f_3_-statistics* and *f_4_-statistics* exclusively for all involved TK-speaking populations further demonstrated that Gelao groups (except Longlin Gelao) possessed the most ANEA-related ancestry, and Dong populations, Longli Bouyei, Longlin Gelao and Guangxi Zhuang also possessed more ANEA-related ancestry, while Dai, Li groups and TK-speaking populations in Southeast Asia possessed the most ASEA-related ancestry, and Huanjiang Maonan and Luocheng Mulam also possessed more ASEA-related ancestry. Fine-scale genetic structure among TK people showed the TK people dispersed from southern China to Southeast Asia and mixed with different local or resident indigenous populations and formed the observed obsessed genetic landscape.

## Conclusion

Here, we generated new genome-wide SNP genotypes in 70 inland TK-speaking and 45 Sinitic-speaking individuals and comprehensively explored the spatiotemporal differences of shared genetic ancestry of ancient and modern East Asians. Our findings showed extensive genetic diversity of the mainland TK people that were associated with heterogeneous ancestry sharing profiles and uniparental lineages. We found that the genomic makeup of studied Han Chinese received great influences from surrounding TK-speaking groups but maintained their main ancestry descended from Coastal/Inland Neolithic YR East Asians (Pingliangtai, Haojiatai, Xiaojingshan and Lajia). The *f*-statistics-based admixture models revealed two primary proxy sources, which were referred to as the ASEA represented by Neolithic/Iron Age southern East Asians (Hanben, Xitoucun, Tanshishan and Qihe) and modern southern indigenous AA (Ami and Atayal) and TK speakers (Li), and the ANEA represented by Neolithic northern millet farmers deriving from the YR Basin, Tibetan-related populations, Neolithic hunter-gatherers from Mongolia or DevilsCave_N. Our *qpWave/qpAdm-based* and *qpGraph-based* findings further supported those two-way admixture models with different proportions fitted the demographic events of southwestern Han Chinese and mainland TK people, in which ancient YR millet farmers as the northern source, and Qiongzhong Li, island/coastal AN and their ancestors as the southern source. We also found evidence for substantial genetic divergences between TK people residing in the south of the sampling area and TK people residing in the north of the sampling area, which was consistent with the barrier effect of karst landform and its higher altitudes in the west and center in the process of population migration, contacts and subsequent genetic admixture. Finally, our modern and ancient genomic evidence supported the common origin of modern AA and TK-speaking populations: the most possible original birthplace is coastal southern China with the development and expansion of Yangtze rice farmers and their Proto-languages, consistent with the famous language-agriculture co-dispersal model.

## Materials and methods

### Sample collection and genotyping

We sampled 70 TK-speaking (20 Daozhen Gelao, 18 Longli Bouyei, 20 Qiandongnan Dong, 7 Zunyi Gelao, and 5 Guiyang Gelao) and 45 Sinitic-speaking (16 Zunyi Han, 19 Guiyang Han, and 10 Bijie Han) individuals from Guizhou, southwest China. The geographic coordinates of sampling locations per population are displayed in **Figure 1A**. All samples were collected randomly, and each participant had signed the written informed consent. This project was approved by the Medical Ethics Committee of Xiamen University (XDYX201909) and all procedures were performed following the recommendations stated in the Helsinki Declaration of 2000[44]. The extraction and purification of human genomic DNA were performed using the PureLink Genomic DNA Mini Kit (Thermo Fisher Scientific) and then the quantification was conducted using the Quantifiler Human DNA Quantification Kit (Thermo Fisher Scientific) on an Applied Biosystems 7500 Real-time PCR System (Thermo Fisher Scientific) following the manufacturer’s guidelines. All qualified samples were genotyped via the Affymetrix WeGene V1 Arrays and SNP calling yielding a confidence value > 0.1 was regarded as missing data.

### Reference dataset assembly

The newly-genotyped 115 individuals together with 191 ancient and 383 modern East Asians reported in Wang et al. 2020[30] and 24 ancient individuals from northern and southern East Asia[37], 55 ancient genomes from YR, West Liao River (WLR) and Amur River (AR) basins[45] were merged separately with Human Origin (HO) or 1240K datasets from David Reich Lab (https://reich.hms.harvard.edu/datasets). The merged HO-based dataset covered 120,447 overlapped autosomal variants in 11,662 individuals and the merged 1240K-based dataset covered 393,358 overlapped autosomal variants in 7,411 individuals.

### Analysis of genetic relationship and population structure

We employed principal component analysis (PCA) and model-based clustering analysis to visualize how the populations cluster. We performed PCA at the individual level using the smartpca program of EIGENSOFT v.6.1.4[46] with the following parameters: numoutlieriter: 0 and lsqproject: YES. We applied PLINK v.1.9[47] with parameters --indep-pairwise 200 25 0.4 to remove SNPs in strong linkage disequilibrium. We ran ADMIXTURE (v.1.3.0)[48] with the 10-fold cross-validation (--cv = 10) in 100 bootstraps with different random seeds to investigate the genetic-ancestry coefficients of targeted populations, predefining the number of ancestral populations in the range of 2 to 20.

### Estimation of allele sharing and gene flow

We conducted a series of *f_3_*- and *f_4_-statistics* using the *qp3Pop* and *qpDstat* programs of ADMIXTOOLS[49, 50] with default parameters. We calculated the *outgroup-f_3_-statistics* of the form of *f_3_(X, Y; Yoruba*) to measure the shared drift between populations X and Y since their divergence from the outgroup (here we used Mbuti as the baseline of the outgroup), and then we performed admixture-*f_3_-statistics* of the form of *f_3_*(*X*, *Y*; *Studied populations*) to evaluate the potential admixture signals from different source populations in the targeted populations. Based on the results of *outgroup-f_3_-statistics* and the admixture-*f_3_statistics*, we computed the *f_4_*(*W*, *X*; *Y*, *Outgroup*) to formally test whether W or X harbored more population Y related ancestry.

### Modeling the admixture graph

We ran TreeMix v.1.13[51] with migration events varying from 0 to 17 to generate the admixture graph with the maximum likelihood and simulate the genetic drift between studied populations and potential ancestral modern and ancient source populations. To further model population histories that fit the genomic data, we used the *qpGraph* program implemented in the ADMIXTOOLS package[52] to create the admixture topologies and calculate the best-fitting admixture proportions and branch lengths based on the results of the *f-statistics* (results from *f_2_*, *f_3_ and f_4_* values based on the population combinations of the pairs, triples, and quadruples). We started with a topology that fits the data with Mbuti, Denisova, Loschbour, Onge and Tianyuan. We accepted the admixture graph as a good fit when there were no internal zero branch lengths and the absolute value of Z-score of the worst *f_4_*-statistic was below three.

### Inference of mixture proportions

We applied the *qpWave* and *qpAdm[53]* in the ADMIXTOOLS package to test whether a group of targeted populations is coincident with being related via N streams of source populations from a basic set of outgroup populations and then estimate the corresponding ancestry proportions. We used the Mbuti, Romania_Oase1, Ust_Ishim, China_Tianyuan, Kostenki14, Yana_UP, Israel_Natufian, Anatolia_N, Motala_HG, Onge, Papuan, BedouinB, Eskimo_Sireniki, Kalash, Australian, Greenland_Saqqaq and Karitiana as outgroups.

### FineStructure and ChromoPainter

Genome-wide SNP data was pbased using SHAPEIT [46]. Recipient chromosome of tagated individual was painted using all avalbel donor chromosomes as the potential sources, and chromosome painting was conducted using ChromoPainter and ChromoCombine [47]. Further genetic structure based on the coancestry matrix was explored using FineStructure v.4 [47].

### Uniparental haplogroup assignment

We assigned each consensus sequence into a Y-chromosomal or mtDNA haplogroup using an in-house script following the recommendations of the International Society of Genetic Genealogy (ISOGG, http://www.isogg.org/) and mtDNA tree Build 17[54] (http://www.phylotree.org/).

## Acknowledgements

This study was supported by the National Natural Science Foundation of China (31801040), Nanqiang Outstanding Young Talents Program of Xiamen University (X2123302), Major Project of National Social Science Foundation of China (20&ZD248), European Research Council (ERC) grant to Dan Xu (ERC-2019-ADG-883700-TRAM) and Fundamental Research Funds for the Central Universities (ZK1144). GLH was supported by Project funded by China Postdoctoral Science Foundation (2021).

## Data Availability

The Genome-wide variation data have been deposited in the Genome Variation Map (GVM) in Big Data Center, Beijing Institute of Genomics (BIG), Chinese Academy of Science, under accession numbers xxxxxx that are publicly accessible at http://bigd.big.ac.cn/gvm/getProjectDetail?project= xxxxxx (will be available upon publication).

## Reference

1. Bae CJ, Douka K, Petraglia MD. Human Colonization of Asia in the Late Pleistocene: An Introduction to Supplement 17. Current Anthropology. 2017;58(S17):S373–S82. doi: 10.1086/694420.

2. Higham C. Hunter-gatherers in southeast Asia: from prehistory to the present. Hum Biol. 2013;85(1-3):21–43. Epub 2013/12/04. doi: 10.3378/027.085.0302. PubMed PMID: 24297219.

3. Fu Q, Meyer M, Gao X, Stenzel U, Burbano HA, Kelso J, et al. DNA analysis of an early modern human from Tianyuan Cave, China. Proc Natl Acad Sci U S A. 2013;110(6):2223–7. Epub 2013/01/24. doi: 10.1073/pnas.1221359110. PubMed PMID: 23341637; PubMed Central PMCID: PMCPMC3568306.

4. Li YC, Ye WJ, Jiang CG, Zeng Z, Tian JY, Yang LQ, et al. River Valleys Shaped the Maternal Genetic Landscape of Han Chinese. Mol Biol Evol. 2019;36(8):1643–52. Epub 2019/05/22. doi: 10.1093/molbev/msz072. PubMed PMID: 31112995.

5. Ren LL, Dong GH, Liu FW, Guedes JD, Flad RK, Ma MM, et al. Foraging and farming: archaeobotanical and zooarchaeological evidence for Neolithic exchange on the Tibetan Plateau. Antiquity. 2020;94(375):637–52. doi: 10.15184/aqy.2020.35. PubMed PMID: WOS:000538867100011.

6. Jeong C, Ozga AT, Witonsky DB, Malmstrom H, Edlund H, Hofman CA, et al. Long-term genetic stability and a high-altitude East Asian origin for the peoples of the high valleys of the Himalayan arc. Proc Natl Acad Sci U S A. 2016;113(27):7485–90. Epub 2016/06/22. doi: 10.1073/pnas.1520844113. PubMed PMID: 27325755; PubMed Central PMCID: PMCPMC4941446.

7. He G, Wang Z, Guo J, Wang M, Zou X, Tang R, et al. Inferring the population history of Tai-Kadai-speaking people and southernmost Han Chinese on Hainan Island by genome-wide array genotyping. Eur J Hum Genet. 2020. Epub 2020/03/04. doi: 10.1038/s41431-020-0599-7. PubMed PMID: 32123326.

8. Kampuansai J, Kutanan W, Tassi F, Kaewgahya M, Ghirotto S, Kangwanpong D. Effect of migration patterns on maternal genetic structure: a case of Tai-Kadai migration from China to Thailand. J Hum Genet. 2017;62(2):223–8. Epub 2016/09/09. doi: 10.1038/jhg.2016.112. PubMed PMID: 27604557.

9. Yang XY, Wan ZW, Perry L, Lu HY, Wang Q, Zhao CH, et al. Early millet use in northern China. Proc Natl Acad Sci U S A. 2012;109(10):3726–30. doi: 10.1073/pnas.1115430109. PubMed PMID: WOS:000301117700031.

10. Zuo XX, Lu HY, Jiang LP, Zhang JP, Yang XY, Huan XJ, et al. Dating rice remains through phytolith carbon-14 study reveals domestication at the beginning of the Holocene. Proc Natl Acad Sci U S A. 2017;114(25):6486–91. doi: 10.1073/pnas.1704304114. PubMed PMID: WOS:000403687300029.

11. He KY, Lu HY, Zhang JP, Wang C, Huan XJ. Prehistoric evolution of the dualistic structure mixed rice and millet farming in China. Holocene. 2017;27(12):1885–98. doi: 10.1177/0959683617708455. PubMed PMID: WOS:000417713200007.

12. The prehistory of the Daic (Tai-Kadai) speaking peoples and the hypothesis of an Austronesian connection. 2013.

13. Kutanan W, Kampuansai J, Srikummool M, Kangwanpong D, Ghirotto S, Brunelli A, et al. Complete mitochondrial genomes of Thai and Lao populations indicate an ancient origin of Austroasiatic groups and demic diffusion in the spread of Tai-Kadai languages. Hum Genet. 2017;136(1):85–98. Epub 2016/11/12. doi: 10.1007/s00439-016-1742-y. PubMed PMID: 27837350; PubMed Central PMCID: PMCPMC5214972 procedures performed in studies involving human participants were approved by Chiang Mai University, Khon Kaen University, Naruesuan University, and the Ethics Commission of the University of Leipzig Medical Faculty. Informed consent Informed consent was obtained from all individual participants included in the study.

14. Kutanan W, Kampuansai J, Brunelli A, Ghirotto S, Pittayaporn P, Ruangchai S, et al. New insights from Thailand into the maternal genetic history of Mainland Southeast Asia. Eur J Hum Genet. 2018;26(6):898–911. Epub 2018/02/28. doi: 10.1038/s41431-018-0113-7. PubMed PMID: 29483671; PubMed Central PMCID: PMCPMC5974021.

15. Liu J, Du W, Wang M, Liu C, Wang S, He G, et al. Forensic features, genetic diversity and structure analysis of three Chinese populations using 47 autosomal InDels. Forensic Sci Int Genet. 2020;45:102227. Epub 2019/12/23. doi: 10.1016/j.fsigen.2019.102227. PubMed PMID: 31865224.

16. He G, Wang Z, Zou X, Wang M, Liu J, Wang S, et al. Tai-Kadai-speaking Gelao population: Forensic features, genetic diversity and population structure. Forensic Sci Int Genet. 2019;40:e231–e9. Epub 2019/03/27. doi: 10.1016/j.fsigen.2019.03.013. PubMed PMID: 30910535.

17. Liu J, Ye Z, Wang Z, Zou X, He G, Wang M, et al. Genetic diversity and phylogenetic analysis of Chinese Han and Li ethnic populations from Hainan Island by 30 autosomal insertion/deletion polymorphisms. Forensic Sciences Research. 2019:1–7. doi: 10.1080/20961790.2019.1672933.

18. He G, Zou X, Wang M, Liu J, Wang F, Hou Y, et al. Population genetics, diversity, forensic characteristics of four Chinese populations inferred from X-chromosomal short tandem repeats. Leg Med (Tokyo). 2020;43:101677. Epub 2020/01/27. doi: 10.1016/j.legalmed.2020.101677. PubMed PMID: 31982839.

19. Chen P, He G, Zou X, Wang M, Luo H, Yu L, et al. Genetic structure and polymorphisms of Gelao ethnicity residing in southwest china revealed by X-chromosomal genetic markers. Sci Rep. 2018;8(1):14585. Epub 2018/10/03. doi: 10.1038/s41598-018-32945-7. PubMed PMID: 30275508; PubMed Central PMCID: PMCPMC6167355.

20. Song M, Wang Z, Zhang Y, Zhao C, Lang M, Xie M, et al. Forensic characteristics and phylogenetic analysis of both Y-STR and Y-SNP in the Li and Han ethnic groups from Hainan Island of China. Forensic Sci Int Genet. 2019;39:e14–e20. Epub 2018/12/14. doi: 10.1016/j.fsigen.2018.11.016. PubMed PMID: 30522950.

21. Kutanan W, Kampuansai J, Srikummool M, Brunelli A, Ghirotto S, Arias L, et al. Contrasting Paternal and Maternal Genetic Histories of Thai and Lao Populations. Mol Biol Evol. 2019;36(7):1490–506. Epub 2019/04/14. doi: 10.1093/molbev/msz083. PubMed PMID: 30980085; PubMed Central PMCID: PMCPMC6573475.

22. Liu C, Wang SY, Zhao M, Xu ZY, Hu YH, Chen F, et al. Mitochondrial DNA polymorphisms in Gelao ethnic group residing in Southwest China. Forensic Sci Int Genet. 2011;5(1):e4–10. Epub 2010/05/25. doi: 10.1016/j.fsigen.2010.04.007. PubMed PMID: 20494640.

23. Duong NT, Macholdt E, Ton ND, Arias L, Schroder R, Van Phong N, et al. Complete human mtDNA genome sequences from Vietnam and the phylogeography of Mainland Southeast Asia. Sci Rep. 2018;8(1):11651. Epub 2018/08/05. doi: 10.1038/s41598-018-29989-0. PubMed PMID: 30076323; PubMed Central PMCID: PMCPMC6076260.

24. Liu D, Duong NT, Ton ND, Van Phong N, Pakendorf B, Hai NV, et al. Extensive ethnolinguistic diversity in Vietnam reflects multiple sources of genetic diversity. 2019. doi: 10.1101/857367.

25. Habu J, Lape PV, Olsen JW. Handbook of East and Southeast Asian Archaeology: Springer; 2017.

26. Wang M, He G, Su Y, Wang S, Zou X, Liu J, et al. Massively parallel sequencing of mitogenome sequences reveals the forensic features and maternal diversity of Tai-Kadai-speaking Hlai Islanders. Forensic Science International: Genetics. 2020.

27. Wang L, Lv M, Zaumsegel D, Zhang L, Liu F, Xiang J, et al. A comparative study of insertion/deletion polymorphisms applied among Southwest, South and Northwest Chinese populations using Investigator((R)) DIPplex. Forensic Sci Int Genet. 2016;21:10–4. Epub 2015/12/15. doi: 10.1016/j.fsigen.2015.08.005. PubMed PMID: 26656953.

28. Feng R, Zhao Y, Chen S, Li Q, Fu Y, Zhao L, et al. Genetic analysis of 50 Y-STR loci in Dong, Miao, Tujia, and Yao populations from Hunan. Int J Legal Med. 2020;134(3):981–3. Epub 2019/07/03. doi: 10.1007/s00414-019-02115-z. PubMed PMID: 31263947.

29. Weinstein JL. Empire and Identity in Guizhou: Local Resistance to Qing Expansion: University of Washington Press; 2013.

30. Wang C-C, Yeh H-Y, Popov AN, Zhang H-Q, Matsumura H, Sirak K, et al. The Genomic Formation of Human Populations in East Asia. Nature. 2020. doi: 10.1101/2020.03.25.004606.

31. McColl H, Racimo F, Vinner L, Demeter F, Gakuhari T, Moreno-Mayar JV, et al. The prehistoric peopling of Southeast Asia. Science. 2018;361(6397):88–92. Epub 2018/07/07. doi: 10.1126/science.aat3628. PubMed PMID: 29976827.

32. Lipson M, Cheronet O, Mallick S, Rohland N, Oxenham M, Pietrusewsky M, et al. Ancient genomes document multiple waves of migration in Southeast Asian prehistory. Science. 2018;361(6397):92–5. Epub 2018/05/19. doi: 10.1126/science.aat3188. PubMed PMID: 29773666; PubMed Central PMCID: PMCPMC6476732.

33. Ning C, Wang CC, Gao S, Yang Y, Zhang X, Wu X, et al. Ancient Genomes Reveal Yamnaya-Related Ancestry and a Potential Source of Indo-European Speakers in Iron Age Tianshan. Curr Biol. 2019;29(15):2526–32 e4. Epub 2019/07/30. doi: 10.1016/j.cub.2019.06.044. PubMed PMID: 31353181.

34. Yang MA, Gao X, Theunert C, Tong H, Aximu-Petri A, Nickel B, et al. 40,000-Year-Old Individual from Asia Provides Insight into Early Population Structure in Eurasia. Curr Biol. 2017;27(20):3202–8 e9. Epub 2017/10/17. doi: 10.1016/j.cub.2017.09.030. PubMed PMID: 29033327; PubMed Central PMCID: PMCPMC6592271.

35. Jeong C, Wilkin S, Amgalantugs T, Bouwman AS, Taylor WTT, Hagan RW, et al. Bronze Age population dynamics and the rise of dairy pastoralism on the eastern Eurasian steppe. Proc Natl Acad Sci U S A. 2018;115(48):E11248–E55. Epub 2018/11/07. doi: 10.1073/pnas.1813608115. PubMed PMID: 30397125; PubMed Central PMCID: PMCPMC6275519.

36. Damgaard PB, Marchi N, Rasmussen S, Peyrot M, Renaud G, Korneliussen T, et al. 137 ancient human genomes from across the Eurasian steppes. Nature. 2018;557(7705):369–74. Epub 2018/05/11. doi: 10.1038/s41586-018-0094-2. PubMed PMID: 29743675.

37. Yang MA, Fan X, Sun B, Chen C, Lang J, Ko YC, et al. Ancient DNA indicates human population shifts and admixture in northern and southern China. Science. 2020;369(6501):282–8. Epub 2020/05/16. doi: 10.1126/science.aba0909. PubMed PMID: 32409524.

38. Peng MS, He JD, Liu HX, Zhang YP. Tracing the legacy of the early Hainan Islanders--a perspective from mitochondrial DNA. BMC Evol Biol. 2011;11(1):46. Epub 2011/02/18. doi: 10.1186/1471-2148-11-46. PubMed PMID: 21324107; PubMed Central PMCID: PMCPMC3048540.

39. Li D-N, Wang C-C, Lu Y, Qin Z-D, Yang K, Lin X-J, et al. Three phases for the early peopling of Hainan Island viewed from mitochondrial DNA. Journal of Systematics and Evolution. 2013;51(6):671–80. doi: 10.1111/jse.12024. PubMed PMID: WOS:000326574200004.

40. Mengge W, Guanglin H, Yongdong S, Shouyu W, Xing Z, Jing L, et al. Massively parallel sequencing of mitogenome sequences reveals the forensic features and maternal diversity of tai-kadai-speaking hlai islanders. Forensic Sci Int Genet. 2020;47:102303. Epub 2020/05/04. doi: 10.1016/j.fsigen.2020.102303. PubMed PMID: 32361554.

41. Sagart L. The higher phylogeny of Austronesian and the position of Tai-Kadai. Oceanic Linguistics. 2004;43(2):411–44.

42. Reich D. Who we are and how we got here: Ancient DNA and the new science of the human past: Oxford University Press; 2018.

43. Wen B, Li H, Lu D, Song X, Zhang F, He Y, et al. Genetic evidence supports demic diffusion of Han culture. Nature. 2004;431(7006):302–5. Epub 2004/09/17. doi: 10.1038/nature02878. PubMed PMID: 15372031.

44. Association WM. World Medical Association Declaration of Helsinki. Ethical principles for medical research involving human subjects. Bulletin of the World Health Oiganization. 2001;79(4):373.

45. Ning C, Li T, Wang K, Zhang F, Li T, Wu X, et al. Ancient genomes from northern China suggest links between subsistence changes and human migration. Nat Commun. 2020;11(1):2700. Epub 2020/06/03. doi: 10.1038/s41467-020-16557-2. PubMed PMID: 32483115.

46. Patterson N, Price AL, Reich D. Population structure and eigenanalysis. PLoS Genet. 2006;2(12):e190. Epub 2006/12/30. doi: 10.1371/journal.pgen.0020190. PubMed PMID: 17194218; PubMed Central PMCID: PMCPMC1713260.

47. Chang CC, Chow CC, Tellier LC, Vattikuti S, Purcell SM, Lee JJ. Second-generation PLINK: rising to the challenge of larger and richer datasets. Gigascience. 2015;4:7. Epub 2015/02/28. doi: 10.1186/s13742-015-0047-8. PubMed PMID: 25722852; PubMed Central PMCID: PMCPMC4342193.

48. Alexander DH, Novembre J, Lange K. Fast model-based estimation of ancestry in unrelated individuals. Genome Res. 2009;19(9):1655–64. Epub 2009/08/04. doi: 10.1101/gr.094052.109. PubMed PMID: 19648217; PubMed Central PMCID: PMCPMC2752134.

49. Reich D, Thangaraj K, Patterson N, Price AL, Singh L. Reconstructing Indian population history. Nature. 2009;461(7263):489–94. Epub 2009/09/26. doi: 10.1038/nature08365. PubMed PMID: 19779445; PubMed Central PMCID: PMCPMC2842210.

50. Patterson N, Moorjani P, Luo Y, Mallick S, Rohland N, Zhan Y, et al. Ancient admixture in human history. Genetics. 2012;192(3):1065–93. Epub 2012/09/11. doi: 10.1534/genetics.112.145037. PubMed PMID: 22960212; PubMed Central PMCID: PMCPMC3522152.

51. Pickrell JK, Pritchard JK. Inference of population splits and mixtures from genome-wide allele frequency data. PLoS Genet. 2012;8(11):e1002967. Epub 2012/11/21. doi: 10.1371/journal.pgen.1002967. PubMed PMID: 23166502; PubMed Central PMCID: PMCPMC3499260.

52. Fu Q, Hajdinjak M, Moldovan OT, Constantin S, Mallick S, Skoglund P, et al. An early modern human from Romania with a recent Neanderthal ancestor. Nature. 2015;524(7564):216–9. Epub 2015/06/23. doi: 10.1038/nature14558. PubMed PMID: 26098372; PubMed Central PMCID: PMCPMC4537386.

53. Haak W, Lazaridis I, Patterson N, Rohland N, Mallick S, Llamas B, et al. Massive migration from the steppe was a source for Indo-European languages in Europe. Nature. 2015;522(7555):207–11. Epub 2015/03/04. doi: 10.1038/nature14317. PubMed PMID: 25731166; PubMed Central PMCID: PMCPMC5048219.

54. van Oven M, Kayser M. Updated comprehensive phylogenetic tree of global human mitochondrial DNA variation. Hum Mutat. 2009;30(2):E386–94. Epub 2008/10/15. doi: 10.1002/humu.20921. PubMed PMID: 18853457.

