## Supplementary Figures S41-67 for "New insights from the combined discrimination of modern/ancient genome-wide shared alleles and haplotypes: Differentiated demographic history reconstruction of Tai-Kadai and Sinitic people in South China"

[1]State Key Laboratory of Cellular Stress Biology, School of Life Sciences, Department of Anthropology and Ethnology, Institute of Anthropology, State Key Laboratory of Marine Environmental Science, Xiamen University, Xiamen 361005, PR China
[2]Institute of Forensic Medicine, West China School of Basic Science and Forensic Medicine, Sichuan University, Chengdu 610000, PR China
[3]School of Forensic Medicine, Zunyi Medical University, Zunyi 563000, Guizhou, PR China
[4]Department of Forensic Medicine, College of Basic Medicine, Chongqing Medical University, Chongqin 400016, PR China
[5]Department of Forensic Medicine, Guizhou Medical University, Guiyang 550000, PR China
[6]Hunan Key Lab of Bioinformatics, School of Computer Science and Engineering, Central South University, Changsha, 410075, PR China
[7]School of Humanities, Nanyang Technological University, Nanyang, 639798, Singapore
[8]School of Basic Medical Sciences, Zhejiang University School of Medicine, Hangzhou 310000, PR China
[9]Institute of Asian Civilizations, Zhejiang University, Hangzhou 310000, PR China
*These authors contributed equally to this work and should be considered co-first authors.

**[#]Corresponding author**
**Guanglin He**
**Affiliation**: State Key Laboratory of Cellular Stress Biology, School of Life Sciences, Department of Anthropology and Ethnology, Institute of Anthropology, State Key Laboratory of Marine Environmental Science, Xiamen University, Xiamen 361005, PR China; School of Humanities, Nanyang Technological University
**Hui-Yuan Yeh**
**Affiliation**: School of Humanities, Nanyang Technological University, Nanyang, 639798, Singapore
**Chuan-Chao Wang**
**Affiliation**: State Key Laboratory of Cellular Stress Biology, School of Life Sciences, Department of Anthropology and Ethnology, Institute of Anthropology, State Key Laboratory of Marine Environmental Science, Xiamen University, Xiamen 361005, PR China

### Contents of Supplementary Figures

Supplementary Fig. 41A. Temporal changes of shared genetic drift of ancient populations from Yellow River Basin and other region of northern East Asia assessed via *$f_4$(Pingliangtai_LN, Inland/Coastal Neolithic/Bronze Age northern East Asian/Tibet Plateau/Siberia/Japan; Studied inland TK/Sinitic, Mbuti)*. 5
Supplementary Fig. 41B. Temporal changes of shared genetic drift of ancient populations from Yellow River Basin and other region of northern East Asia assessed via *$f_4$(Pingliangtai_LN, Inland/Coastal Neolithic/Bronze Age northern East Asian/Tibet Plateau/Siberia/Japan; Inland/Coastal Neolithic/Bronze Age southern East Asian, Mbuti)*. 6
Supplementary Fig. 42A. Temporal changes of shared genetic drift of ancient populations from Yellow River Basin and other region of northern East Asia assessed via *$f_4$(Shimao_LN, Inland/Coastal Neolithic/Bronze Age northern East Asian/Tibet Plateau/Siberia/Japan; Studied inland TK/Sinitic, Mbuti)*. 7
Supplementary Fig. 42B. Temporal changes of shared genetic drift of ancient populations from Yellow

River Basin and other region of northern East Asia assessed via *$f_4$(Shimao_LN, Inland/Coastal Neolithic/Bronze Age northern East Asian/Tibet Plateau/Siberia/Japan; Inland/Coastal Neolithic/Bronze Age southern East Asian, Mbuti)*. 8
Supplementary Fig. 43A. Temporal changes of shared genetic drift of ancient populations from Yellow River Basin and other region of northern East Asia assessed via *$f_4$(Jinchankou_LN, Inland/Coastal Neolithic/Bronze Age northern East Asian/Tibet Plateau/Siberia/Japan; Studied inland TK/Sinitic, Mbuti)*. 9
Supplementary Fig. 43B. Temporal changes of shared genetic drift of ancient populations from Yellow River Basin and other region of northern East Asia assessed via *$f_4$(Jinchankou_LN, Inland/Coastal Neolithic/Bronze Age northern East Asian/Tibet Plateau/Siberia/Japan; Inland/Coastal Neolithic/Bronze Age southern East Asian, Mbuti)*. 10
Supplementary Fig. 44A. Temporal changes of shared genetic drift of ancient populations from Yellow River Basin and other region of northern East Asia assessed via *$f_4$(Lajia_LN, Inland/Coastal Neolithic/Bronze Age northern East Asian/Tibet Plateau/Siberia/Japan; Studied inland TK/Sinitic, Mbuti)*. 11
Supplementary Fig. 44B. Temporal changes of shared genetic drift of ancient populations from Yellow River Basin and other region of northern East Asia assessed via *$f_4$(Lajia_LN, Inland/Coastal Neolithic/Bronze Age northern East Asian/Tibet Plateau/Siberia/Japan; Inland/Coastal Neolithic/Bronze Age southern East Asian, Mbuti)*. 12
Supplementary Fig. 45. Temporal changes of shared genetic drift of ancient populations from Yellow River Basin and other region of northern East Asia assessed via *$f_4$(Luoheguxiang_LBIA, Inland/Coastal Neolithic/Bronze Age northern East Asian/Tibet Plateau/Siberia/Japan; Studied inland TK/Sinitic, Mbuti)*. 13
Supplementary Fig. 46. Temporal changes of shared genetic drift of ancient populations from Yellow River Basin and other region of northern East Asia assessed via *$f_4$(Jiaozuoniecun_LBIA, Inland/Coastal Neolithic/Bronze Age northern East Asian/Tibet Plateau/Siberia/Japan; Studied inland TK/Sinitic, Mbuti)*. 15
Supplementary Fig. 47. Temporal changes of shared genetic drift of ancient populations from Yellow River Basin and other region of northern East Asia assessed via *$f_4$(Haojiatai_LBIA, Inland/Coastal Neolithic/Bronze Age northern East Asian/Tibet Plateau/Siberia/Japan; Studied inland TK/Sinitic, Mbuti)*. 16
Supplementary Fig. 48. Temporal changes of shared genetic drift of ancient populations from Yellow River Basin and other region of northern East Asia assessed via *$f_4$(Dacaozi_IA, Inland/Coastal Neolithic/Bronze Age northern East Asian/Tibet Plateau/Siberia/Japan; Studied inland TK/Sinitic, Mbuti)*. 17
Supplementary Fig. 49A. Temporal changes of shared genetic drift of ancient populations from Yangtze River surrounding region in southern East Asia assessed via *$f_4$(Qihe_EN, Inland/Coastal Neolithic/Bronze Age southern East Asian; Studied inland TK/Sinitic, Mbuti)*. 18
Supplementary Fig. 49B. Temporal changes of shared genetic drift of ancient populations from Yangtze River surrounding region in southern East Asia assessed via *$f_4$(Qihe_EN, Inland/Coastal Neolithic/Bronze Age southern East Asian; Inland/Coastal Neolithic/Bronze Age northern East Asian/Tibet Plateau/Siberia/Japan, Mbuti)*. 19
Supplementary Fig. 50A. Temporal changes of shared genetic drift of ancient populations from Yangtze River surrounding region in southern East Asia assessed via *$f_4$(Liangdao1_EN, Inland/Coastal*

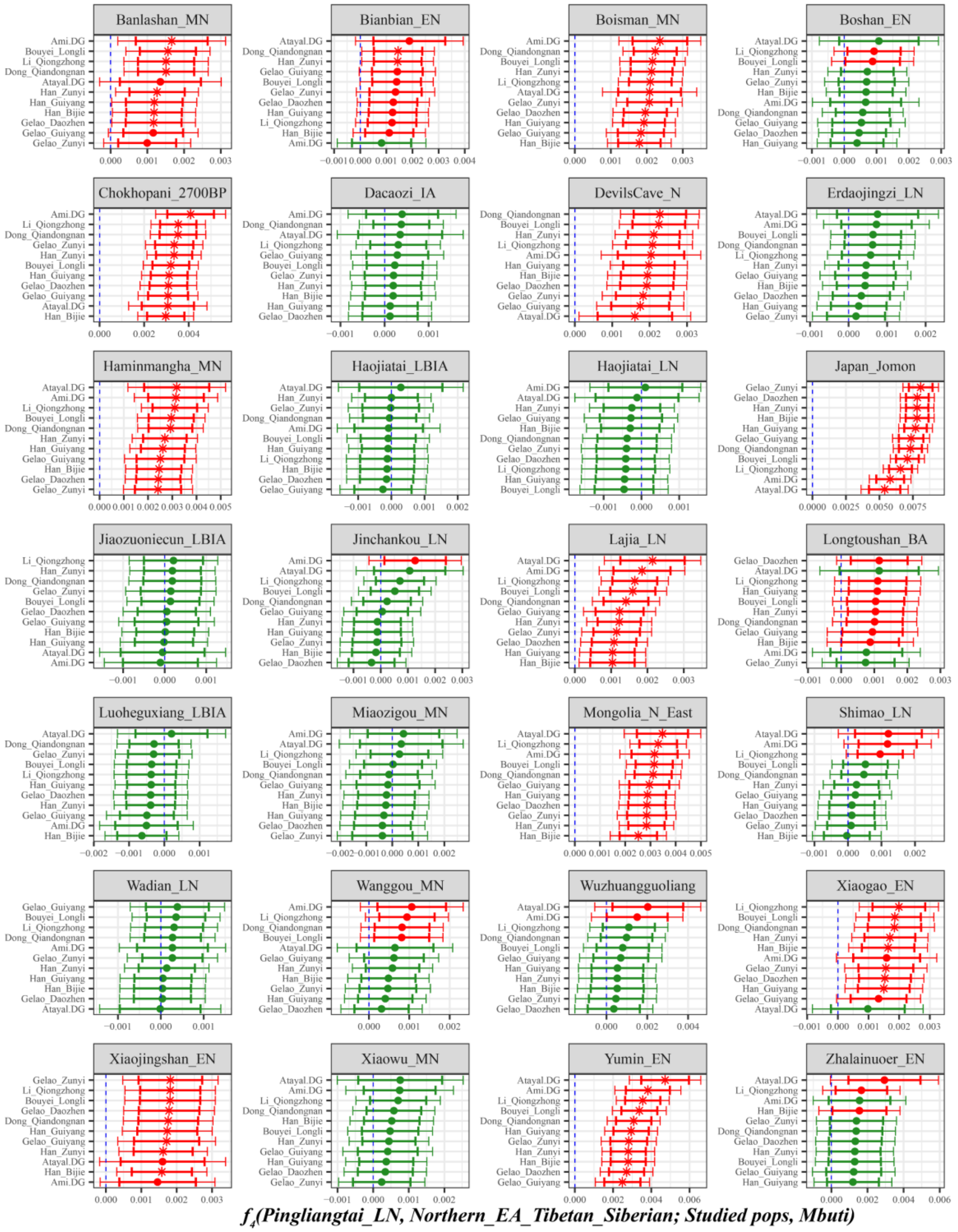

**Supplementary Fig. 41A. Temporal changes of shared genetic drift of ancient populations from Yellow River Basin and other region of northern East Asia assessed via *$f_4$(Pingliangtai_LN, Inland/Coastal Neolithic/Bronze Age northern East Asian/Tibet Plateau/Siberia/Japan; Studied inland TK/Sinitic, Mbuti)*.**

The dashed blue line indicates the zero $f_4$ value. The red asterisk indicates the absolute Z-score value larger than 3, the red point for absolute Z-score value ranging from two to three, and the green point for absolute Z-score value ranging from zero to two. The thick bar denotes two standard errors, and the thin bar for three standard errors. Figures were grouped via the second population in the $f_4$-statistics of Inland/Coastal Neolithic/Bronze Age northern East Asian/Tibet Plateau/Siberia/Japan, which was labeled as the red colour with some significant negative values. Significant negative values denoted the included second populations harboured more central/southern Sinitic or TK related ancestry. Significant positive $f_4$ values indicated the first population had more central/southern Sinitic or TK related ancestry.

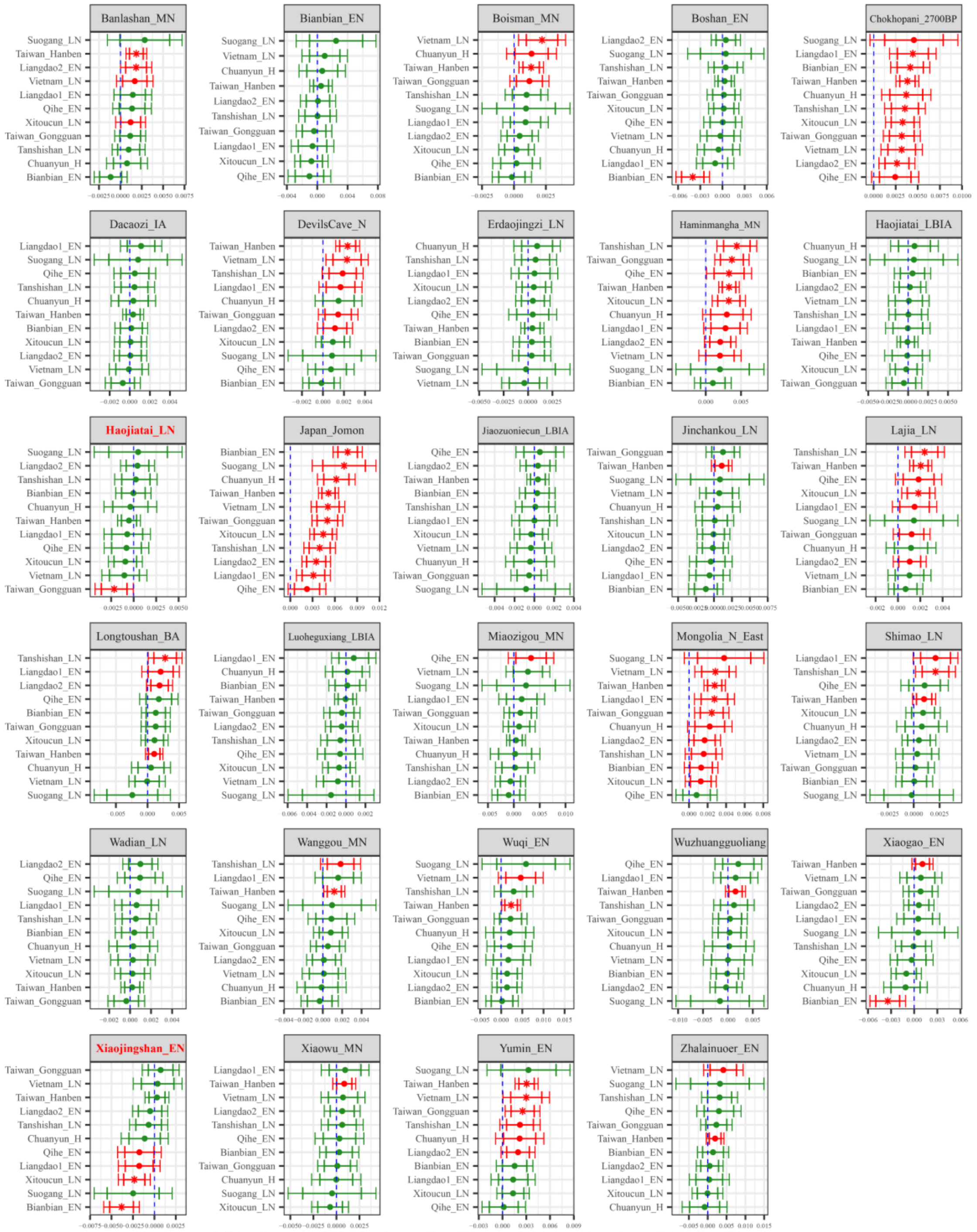

**Supplementary Fig. 41B. Temporal changes of shared genetic drift of ancient populations from Yellow River Basin and other region of northern East Asia assessed via $f_4$*(Pingliangtai_LN, Inland/Coastal Neolithic/Bronze Age northern East Asian/Tibet Plateau/Siberia/Japan; Inland/Coastal Neolithic/Bronze Age southern East Asian, Mbuti)*.**

The dashed blue line indicates the zero $f_4$ value. The red asterisk indicates the absolute Z-score value larger than 3, the red point for absolute Z-score value ranging from two to three, and the green point for absolute Z-score value ranging from zero to two. The thick bar denotes two standard errors, and the thin bar for three standard errors. Figures were grouped via the second population in the $f_4$-statistics of Inland/Coastal Neolithic/Bronze Age northern East Asian/Tibet Plateau/Siberia/Japan, which was labeled as the red colour with some significant negative values. Significant negative values denoted the included second populations harboured more ancient southern East Asian related ancestry. Significant positive $f_4$ values indicated the first population had more ancient southern East Asian related ancestry.

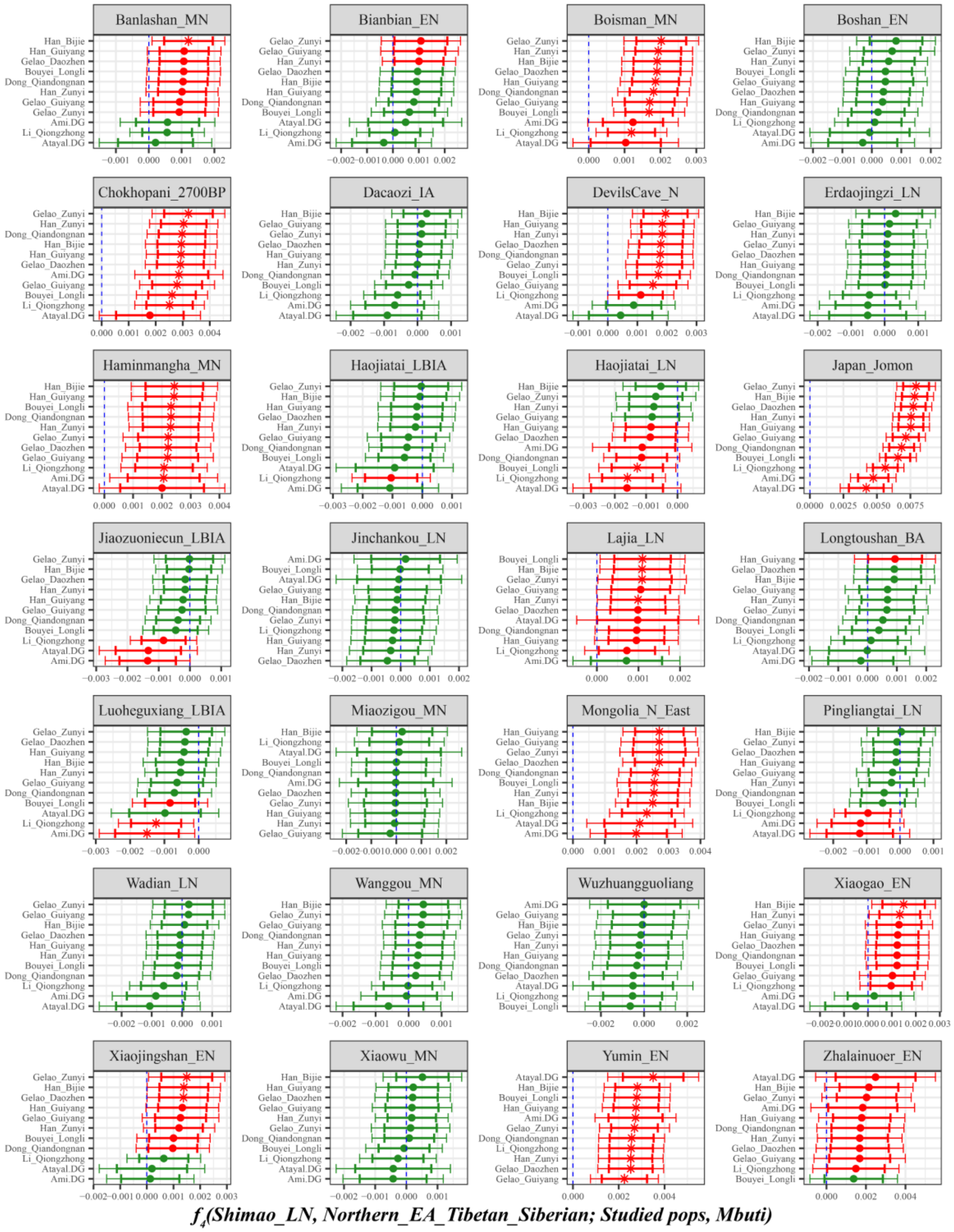

**Supplementary Fig. 42A. Temporal changes of shared genetic drift of ancient populations from Yellow River Basin and other region of northern East Asia assessed via *$f_4$(Shimao_LN, Inland/Coastal Neolithic/Bronze Age northern East Asian/Tibet Plateau/Siberia/Japan; Studied inland TK/Sinitic, Mbuti)*.**

The dashed blue line indicates the zero $f_4$ value. The red asterisk indicates the absolute Z-score value larger than 3, the red point for absolute Z-score value ranging from two to three, and the green point for absolute Z-score value ranging from zero to two. The thick bar denotes two standard errors, and the thin bar for three standard errors. Figures were grouped via the second population in the $f_4$-statistics of Inland/Coastal Neolithic/Bronze Age northern East Asian/Tibet Plateau/Siberia/Japan, which was labeled as the red colour with some significant negative values. Significant negative values denoted the included second populations harboured more central/southern Sinitic or TK related ancestry. Significant positive

$f_4$ values indicated the first population had more central/southern Sinitic or TK related ancestry.

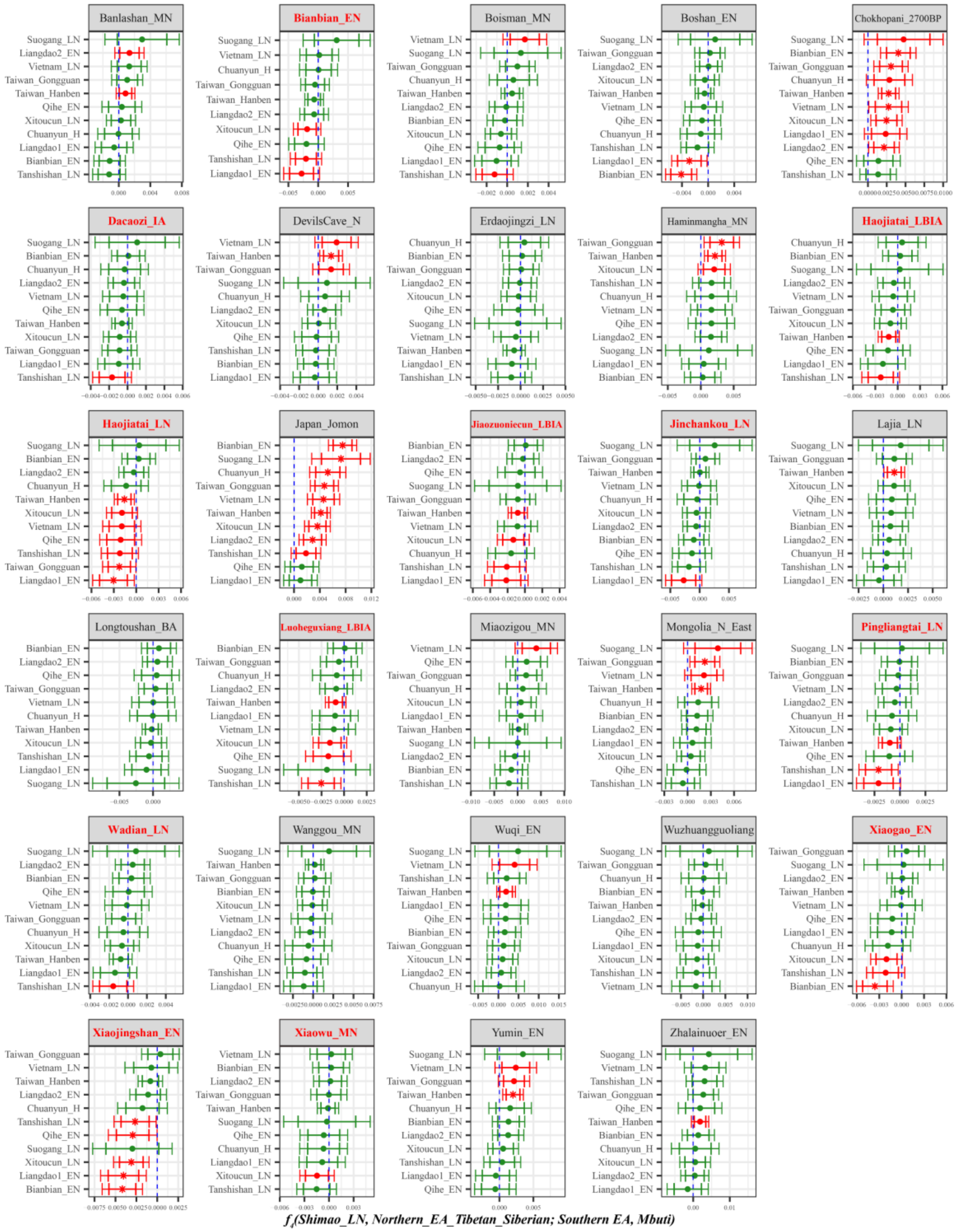

**Supplementary Fig. 42B. Temporal changes of shared genetic drift of ancient populations from Yellow River Basin and other region of northern East Asia assessed via *f4(Shimao_LN, Inland/Coastal Neolithic/Bronze Age northern East Asian/Tibet Plateau/Siberia/Japan; Inland/Coastal Neolithic/Bronze Age southern East Asian, Mbuti)*.**

The dashed blue line indicates the zero $f_4$ value. The red asterisk indicates the absolute Z-score value larger than 3, the red point for absolute Z-score value ranging from two to three, and the green point for absolute Z-score value ranging from zero to two. The thick bar denotes two standard errors, and the thin bar for three standard errors. Figures were grouped via the second population in the $f_4$-statistics of Inland/Coastal Neolithic/Bronze Age northern East Asian/Tibet Plateau/Siberia/Japan, which was labeled as the red colour with some significant negative values. Significant negative values denoted the included second populations harboured more ancient southern East Asian related ancestry. Significant positive $f_4$ values indicated the first population had more ancient southern East Asian related ancestry.

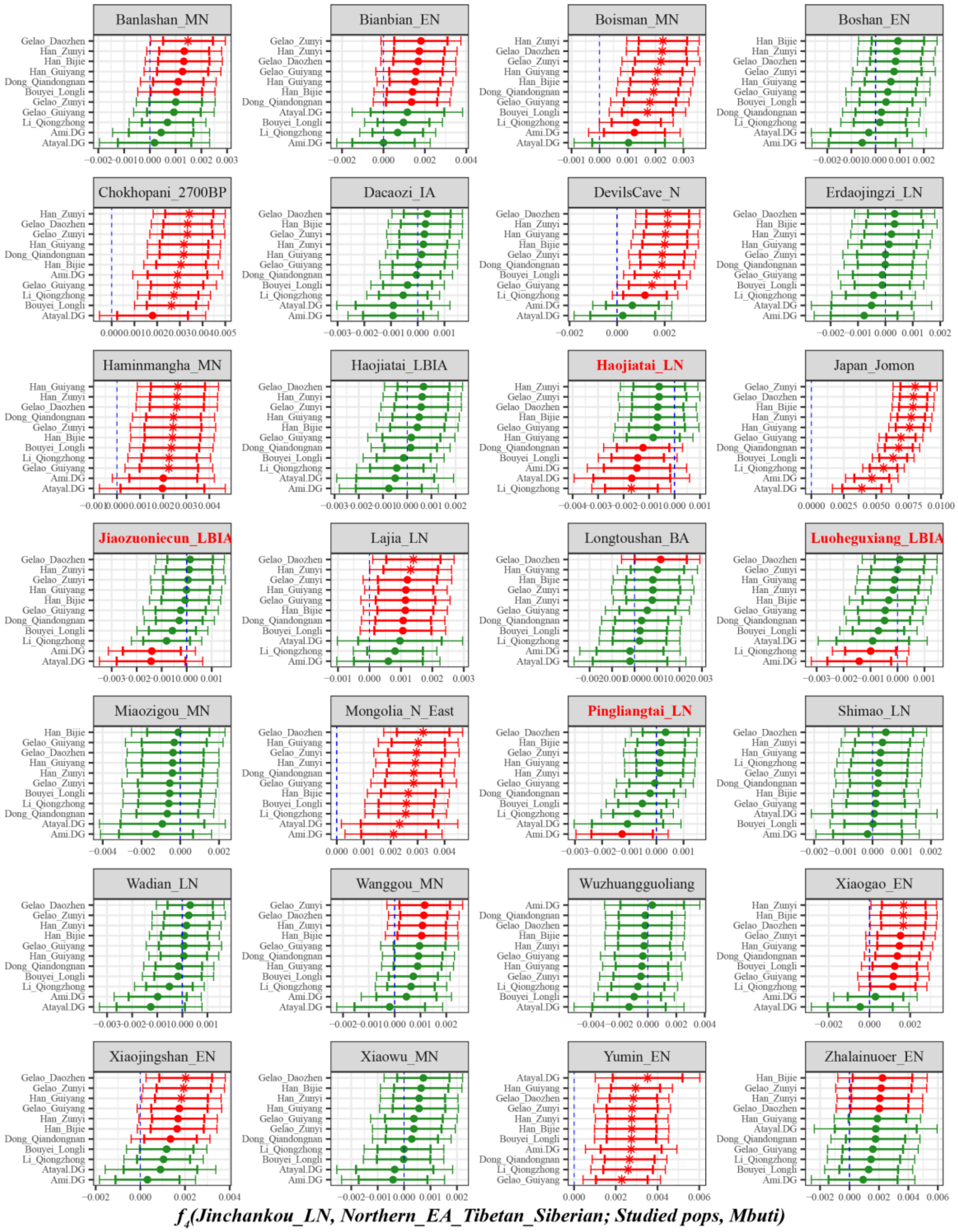

**Supplementary Fig. 43A. Temporal changes of shared genetic drift of ancient populations from Yellow River Basin and other region of northern East Asia assessed via *$f_4$(Jinchankou_LN, Inland/Coastal Neolithic/Bronze Age northern East Asian/Tibet Plateau/Siberia/Japan; Studied inland TK/Sinitic, Mbuti)*.**

The dashed blue line indicates the zero $f_4$ value. The red asterisk indicates the absolute Z-score value larger than 3, the red point for absolute Z-score value ranging from two to three, and the green point for absolute Z-score value ranging from zero to two. The thick bar denotes two standard errors, and the thin bar for three standard errors. Figures were grouped via the second population in the $f_4$-statistics of Inland/Coastal Neolithic/Bronze Age northern East Asian/Tibet Plateau/Siberia/Japan, which was labeled as the red colour with some significant negative values. Significant negative values denoted the included second populations harboured more central/southern Sinitic or TK related ancestry. Significant positive $f_4$ values indicated the first population had more central/southern Sinitic or TK related ancestry.

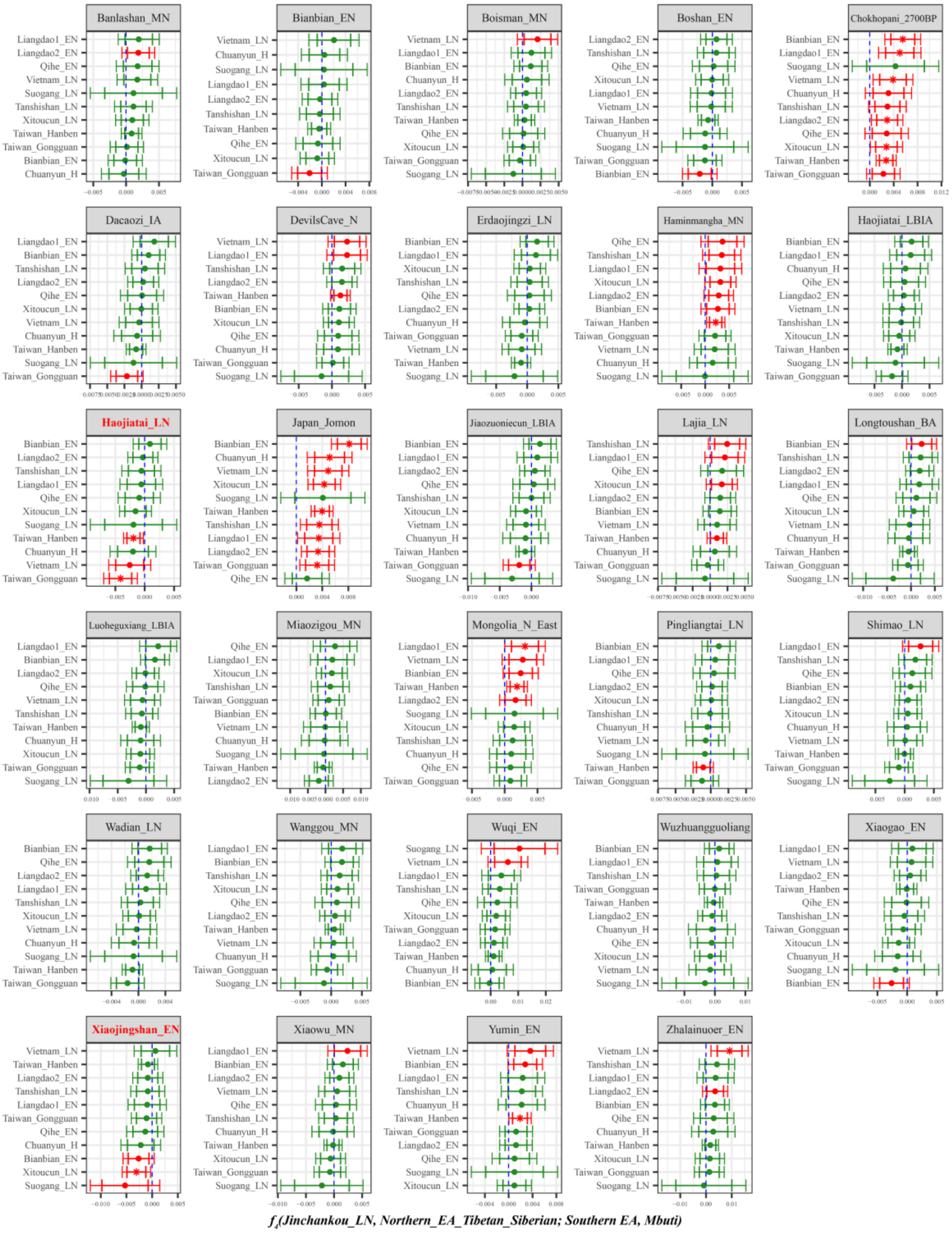

**Supplementary Fig. 43B. Temporal changes of shared genetic drift of ancient populations from Yellow River Basin and other region of northern East Asia assessed via *f*₄*(Jinchankou_LN, Inland/Coastal Neolithic/Bronze Age northern East Asian/Tibet Plateau/Siberia/Japan; Inland/Coastal Neolithic/Bronze Age southern East Asian, Mbuti)*.**

The dashed blue line indicates the zero $f_4$ value. The red asterisk indicates the absolute Z-score value larger than 3, the red point for absolute Z-score value ranging from two to three, and the green point for absolute Z-score value ranging from zero to two. The thick bar denotes two standard errors, and the thin bar for three standard errors. Figures were grouped via the second population in the $f_4$-statistics of

Inland/Coastal Neolithic/Bronze Age northern East Asian/Tibet Plateau/Siberia/Japan, which was labeled as the red colour with some significant negative values. Significant negative values denoted the included second populations harboured more ancient southern East Asian related ancestry. Significant positive $f_4$ values indicated the first population had more ancient southern East Asian related ancestry.

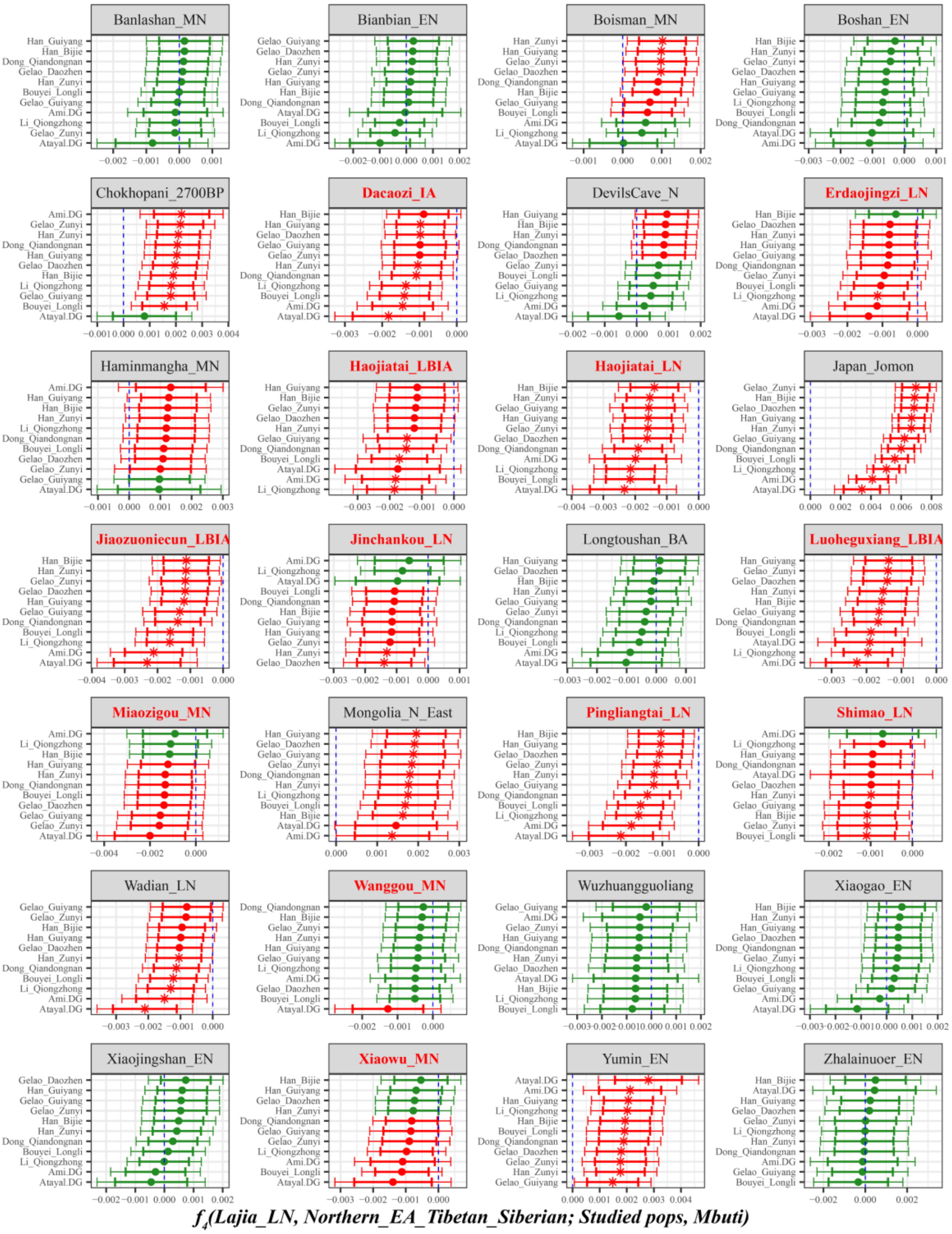

**Supplementary Fig. 44A. Temporal changes of shared genetic drift of ancient populations from Yellow River Basin and other region of northern East Asia assessed via *$f_4$(Lajia_LN, Inland/Coastal Neolithic/Bronze Age northern East Asian/Tibet Plateau/Siberia/Japan; Studied inland TK/Sinitic, Mbuti)*.**

The dashed blue line indicates the zero $f_4$ value. The red asterisk indicates the absolute Z-score value larger than 3, the red point for absolute Z-score value ranging from two to three, and the green point for absolute Z-score value ranging from zero to two. The thick bar denotes two standard errors, and the thin

bar for three standard errors. Figures were grouped via the second population in the $f_4$-statistics of Inland/Coastal Neolithic/Bronze Age northern East Asian/Tibet Plateau/Siberia/Japan, which was labeled as the red colour with some significant negative values. Significant negative values denoted the included second populations harboured more central/southern Sinitic or TK related ancestry. Significant positive $f_4$ values indicated the first population had more central/southern Sinitic or TK related ancestry.

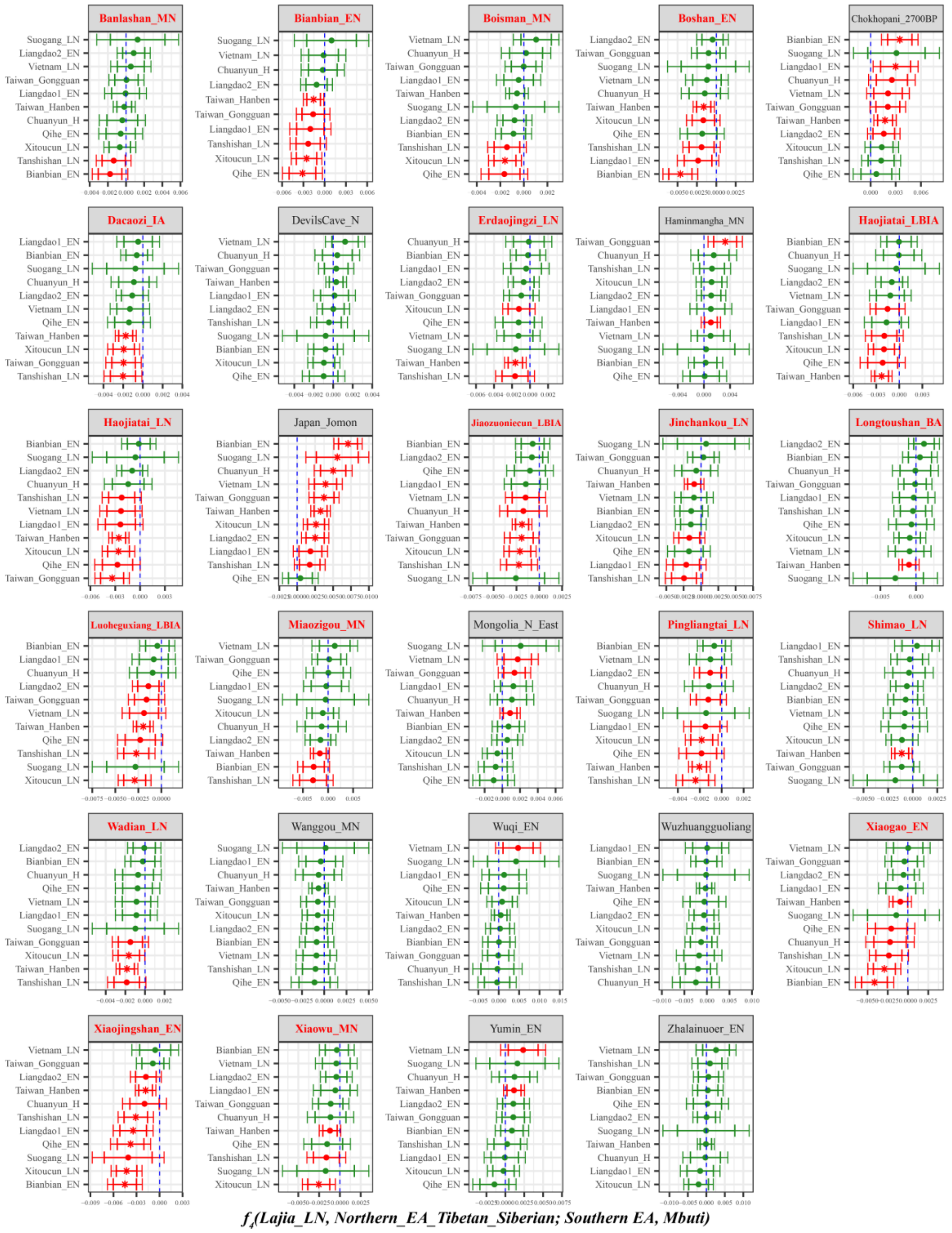

**Supplementary Fig. 44B. Temporal changes of shared genetic drift of ancient populations from Yellow River Basin and other region of northern East Asia assessed via *f₄*(*Lajia_LN, Inland/Coastal Neolithic/Bronze Age northern East Asian/Tibet Plateau/Siberia/Japan; Inland/Coastal Neolithic/Bronze Age southern East Asian, Mbuti*).**

The dashed blue line indicates the zero $f_4$ value. The red asterisk indicates the absolute Z-score value larger than 3, the red point for absolute Z-score value ranging from two to three, and the green point for absolute Z-score value ranging from zero to two. The thick bar denotes two standard errors, and the thin bar for three standard errors. Figures were grouped via the second population in the $f_4$-statistics of Inland/Coastal Neolithic/Bronze Age northern East Asian/Tibet Plateau/Siberia/Japan, which was labeled as the red colour with some significant negative values. Significant negative values denoted the included second populations harboured more ancient southern East Asian related ancestry. Significant positive $f_4$ values indicated the first population had more ancient southern East Asian related ancestry.

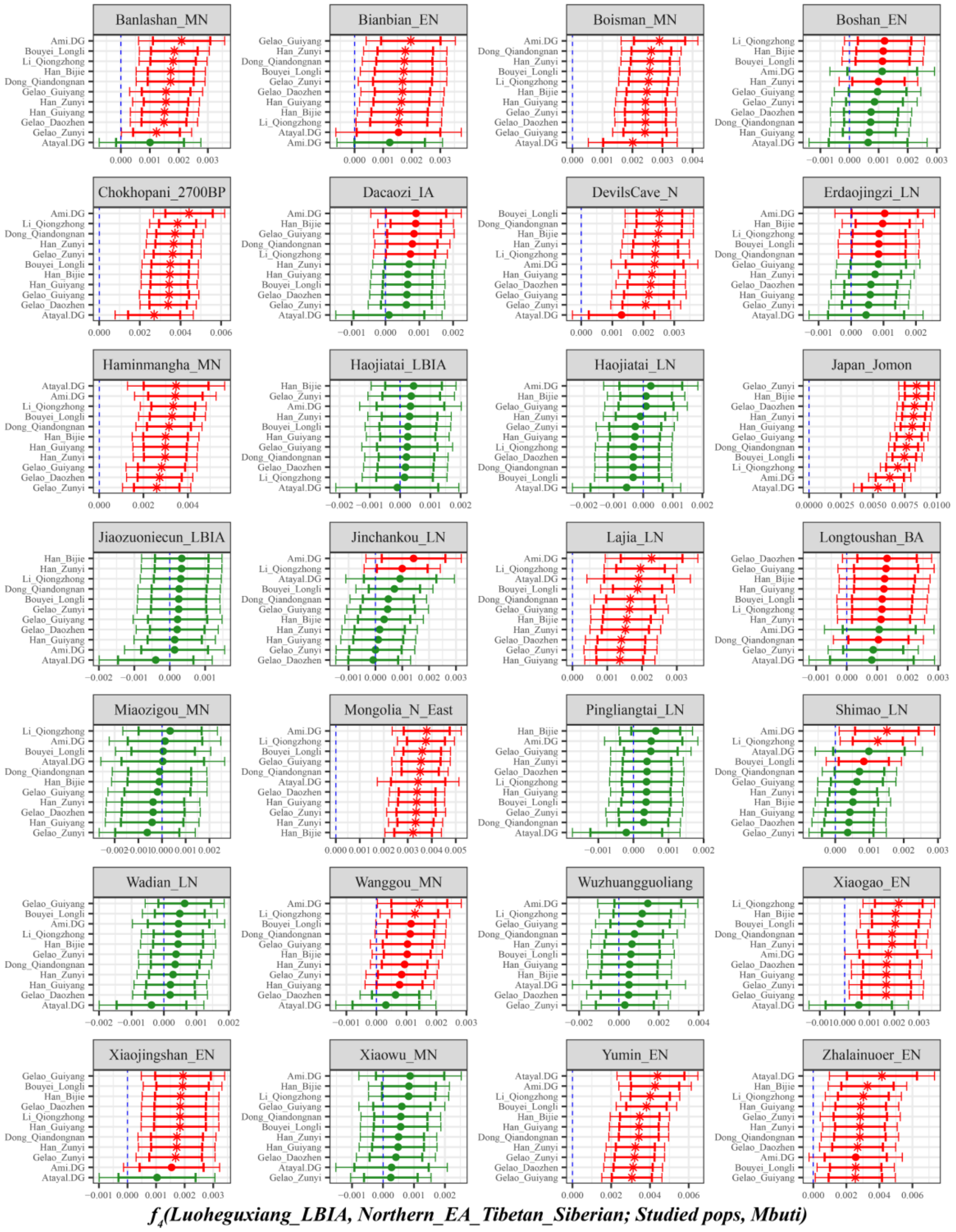

**Supplementary Fig. 45. Temporal changes of shared genetic drift of ancient populations from Yellow River Basin and other region of northern East Asia assessed via *$f_4$(Luoheguxiang_LBIA, Inland/Coastal Neolithic/Bronze Age northern East Asian/Tibet Plateau/Siberia/Japan; Studied inland TK/Sinitic, Mbuti)*.**

The dashed blue line indicates the zero $f_4$ value. The red asterisk indicates the absolute Z-score value larger than 3, the red point for absolute Z-score value ranging from two to three, and the green point for absolute Z-score value ranging from zero to two. The thick bar denotes two standard errors, and the thin bar for three standard errors. Figures were grouped via the second population in the $f_4$-statistics of Inland/Coastal Neolithic/Bronze Age northern East Asian/Tibet Plateau/Siberia/Japan, which was labeled as the red colour with some significant negative values. Significant negative values denoted the included second populations harboured more central/southern Sinitic or TK related ancestry. Significant positive $f_4$ values indicated the first population had more central/southern Sinitic or TK related ancestry.

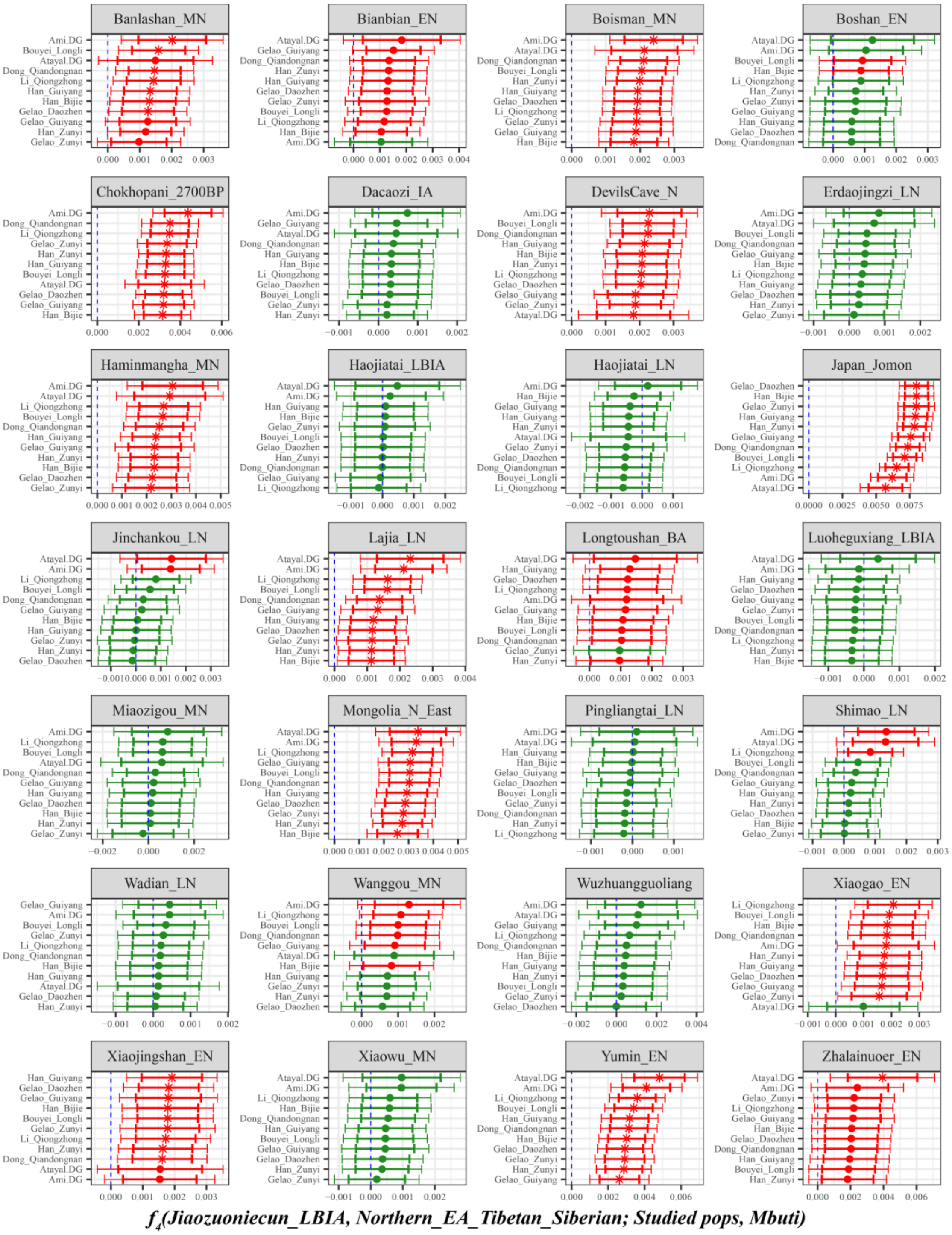

**Supplementary Fig. 46. Temporal changes of shared genetic drift of ancient populations from Yellow River Basin and other region of northern East Asia assessed via $f_4$*(Jiaozuoniecun_LBIA, Inland/Coastal Neolithic/Bronze Age northern East Asian/Tibet Plateau/Siberia/Japan; Studied inland TK/Sinitic, Mbuti)*.**

The dashed blue line indicates the zero $f_4$ value. The red asterisk indicates the absolute Z-score value larger than 3, the red point for absolute Z-score value ranging from two to three, and the green point for absolute Z-score value ranging from zero to two. The thick bar denotes two standard errors, and the thin bar for three standard errors. Figures were grouped via the second population in the $f_4$-statistics of Inland/Coastal Neolithic/Bronze Age northern East Asian/Tibet Plateau/Siberia/Japan, which was labeled as the red colour with some significant negative values. Significant negative values denoted the included second populations harboured more central/southern Sinitic or TK related ancestry. Significant positive $f_4$ values indicated the first population had more central/southern Sinitic or TK related ancestry.

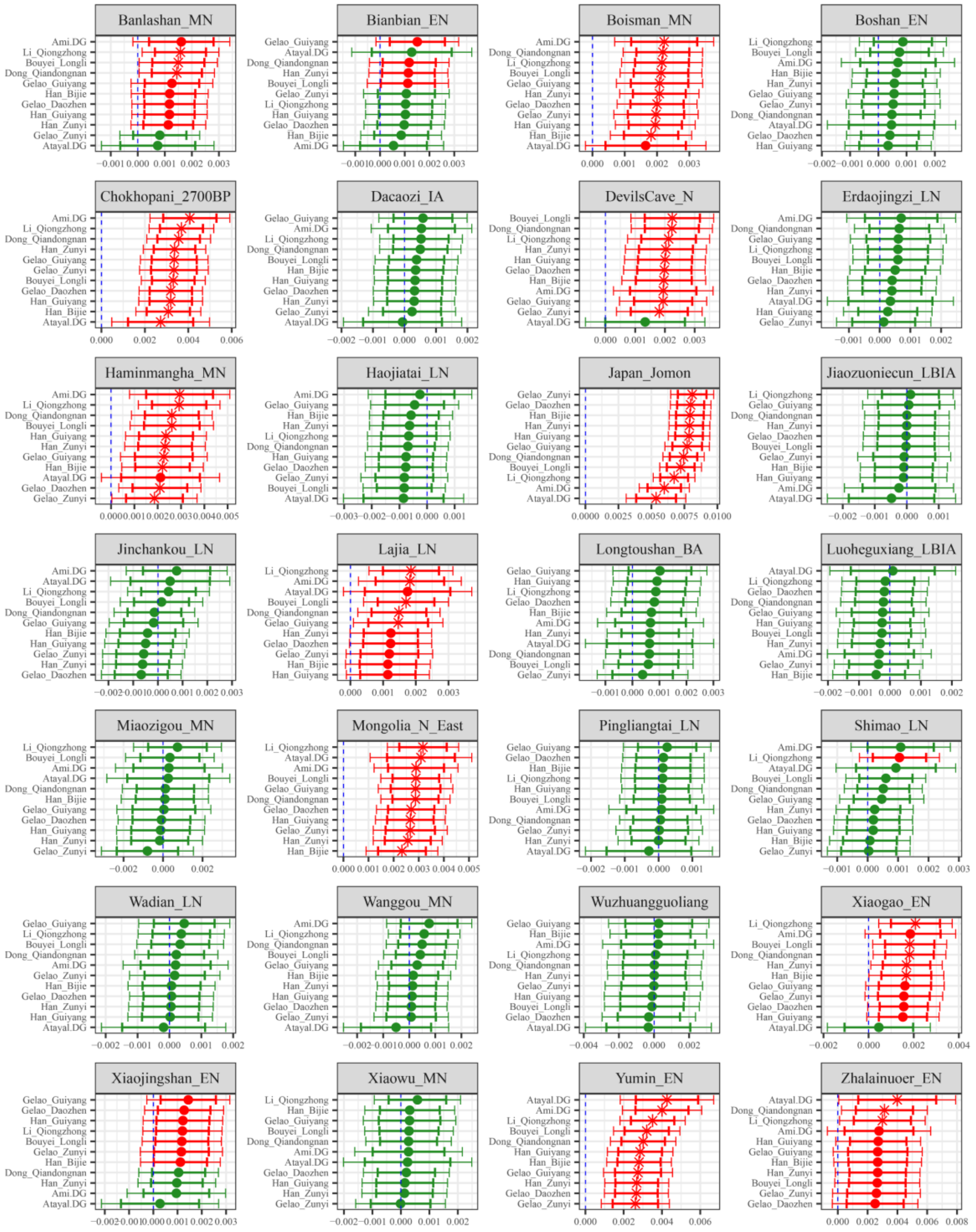

*$f_4$(Haojiatai_LBIA, Northern_EA_Tibetan_Siberian; Studied pops, Mbuti)*

**Supplementary Fig. 47. Temporal changes of shared genetic drift of ancient populations from Yellow River Basin and other region of northern East Asia assessed via *$f_4$(Haojiatai_LBIA, Inland/Coastal Neolithic/Bronze Age northern East Asian/Tibet Plateau/Siberia/Japan; Studied inland TK/Sinitic, Mbuti)*.**

The dashed blue line indicates the zero $f_4$ value. The red asterisk indicates the absolute Z-score value larger than 3, the red point for absolute Z-score value ranging from two to three, and the green point for absolute Z-score value ranging from zero to two. The thick bar denotes two standard errors, and the thin bar for three standard errors. Figures were grouped via the second population in the $f_4$-statistics of Inland/Coastal Neolithic/Bronze Age northern East Asian/Tibet Plateau/Siberia/Japan, which was labeled as the red colour with some significant negative values. Significant negative values denoted the included second populations harboured more central/southern Sinitic or TK related ancestry. Significant positive $f_4$ values indicated the first population had more central/southern Sinitic or TK related ancestry.

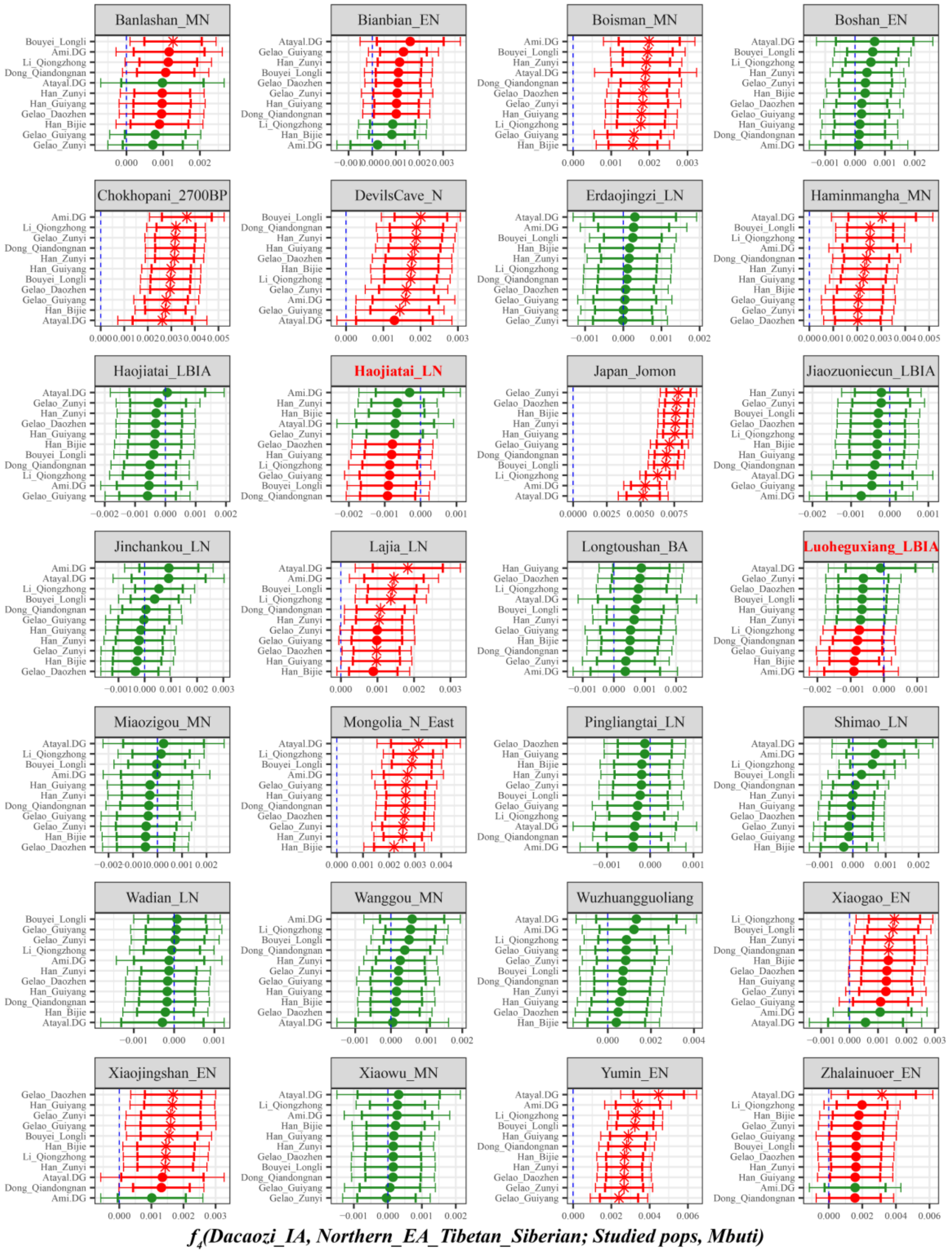

**Supplementary Fig. 48. Temporal changes of shared genetic drift of ancient populations from Yellow River Basin and other region of northern East Asia assessed via *$f_4$(Dacaozi_IA, Inland/Coastal Neolithic/Bronze Age northern East Asian/Tibet Plateau/Siberia/Japan; Studied inland TK/Sinitic, Mbuti)*.**

The dashed blue line indicates the zero $f_4$ value. The red asterisk indicates the absolute Z-score value larger than 3, the red point for absolute Z-score value ranging from two to three, and the green point for absolute Z-score value ranging from zero to two. The thick bar denotes two standard errors, and the thin bar for three standard errors. Figures were grouped via the second population in the $f_4$-statistics of Inland/Coastal Neolithic/Bronze Age northern East Asian/Tibet Plateau/Siberia/Japan, which was labeled as the red colour with some significant negative values. Significant negative values denoted the included second populations harboured more central/southern Sinitic or TK related ancestry. Significant positive

$f_4$ values indicated the first population had more central/southern Sinitic or TK related ancestry.

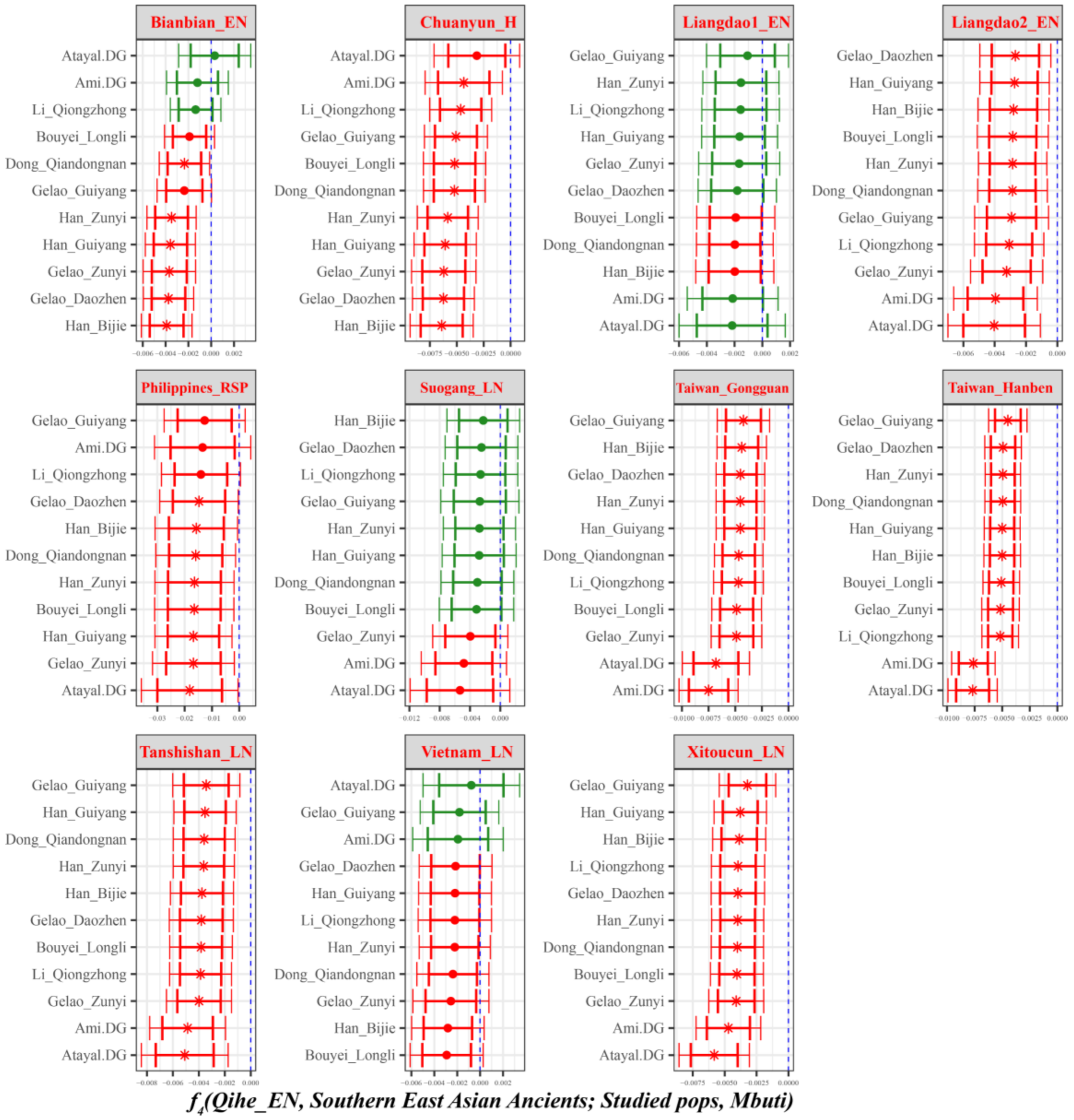

**Supplementary Fig. 49A. Temporal changes of shared genetic drift of ancient populations from Yangtze River surrounding region in southern East Asia assessed via *$f_4$(Qihe_EN, Inland/Coastal Neolithic/Bronze Age southern East Asian; Studied inland TK/Sinitic, Mbuti)*.**

The dashed blue line indicates the zero $f_4$ value. The red asterisk indicates the absolute Z-score value larger than 3, the red point for absolute Z-score value ranging from two to three, and the green point for absolute Z-score value ranging from zero to two. The thick bar denotes two standard errors, and the thin bar for three standard errors. Figures were grouped via the second population in the $f_4$-statistics of Inland/Coastal Neolithic/Bronze Age southern East Asian, which was labeled as the red colour with some significant negative values. Significant negative values denoted the included second populations harboured more central/southern Sinitic or TK related ancestry. Significant positive $f_4$ values indicated the first population had more central/southern Sinitic or TK related ancestry.

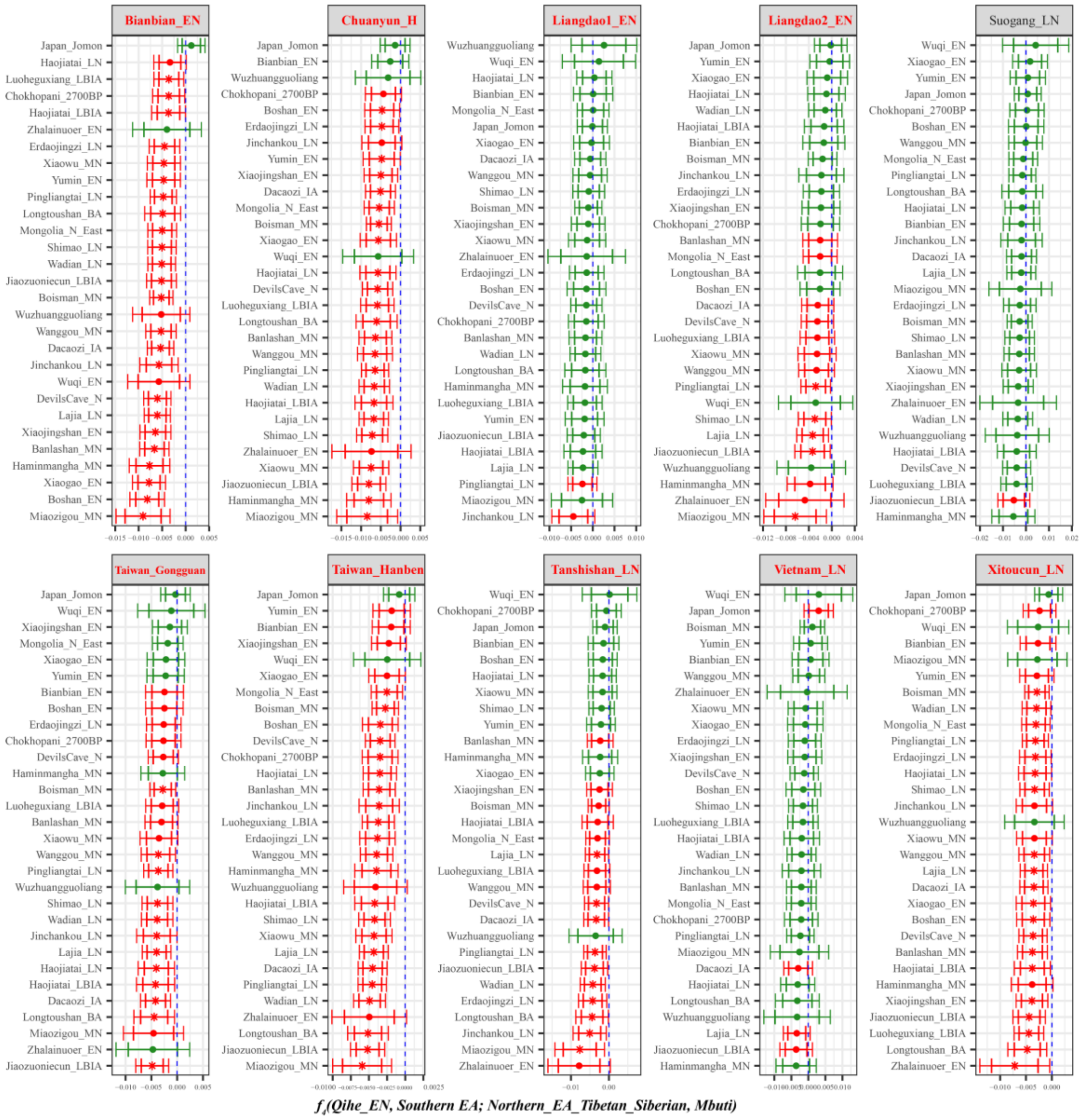

**Supplementary Fig. 49B. Temporal changes of shared genetic drift of ancient populations from Yangtze River surrounding region in southern East Asia assessed via *$f_4$(Qihe_EN, Inland/Coastal Neolithic/Bronze Age southern East Asian; Inland/Coastal Neolithic/Bronze Age northern East Asian/Tibet Plateau/Siberia/Japan, Mbuti)*.**

The dashed blue line indicates the zero $f_4$ value. The red asterisk indicates the absolute Z-score value larger than 3, the red point for absolute Z-score value ranging from two to three, and the green point for absolute Z-score value ranging from zero to two. The thick bar denotes two standard errors, and the thin bar for three standard errors. Figures were grouped via the second population in the $f_4$-statistics of Inland/Coastal Neolithic/Bronze Age southern East Asian, which was labeled as the red colour with some significant negative values. Significant negative values denoted the included second populations harboured more ancient northern East Asian related ancestry. Significant positive $f_4$ values indicated the first population had more ancient northern East Asian related ancestry.

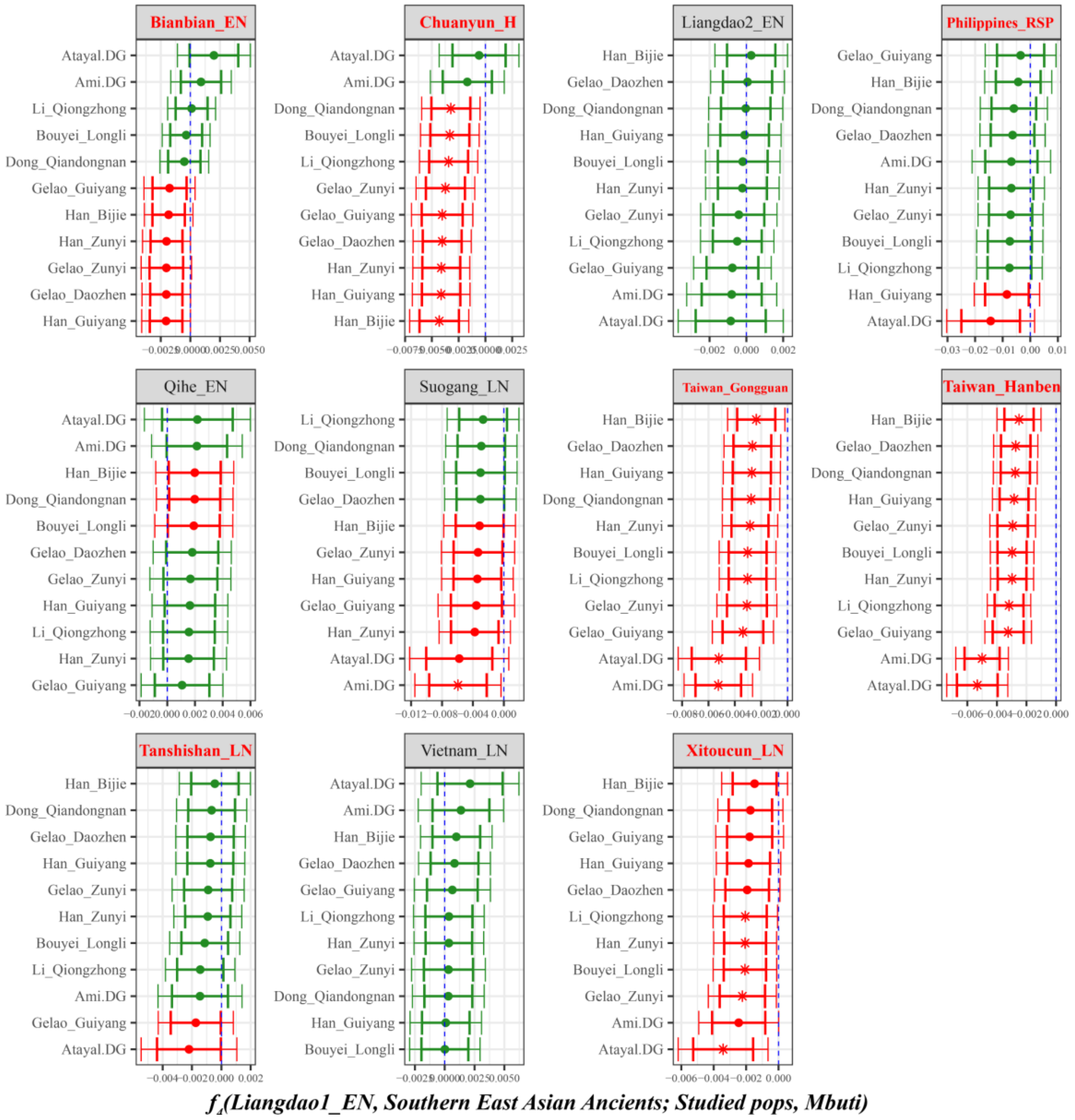

**Supplementary Fig. 50A. Temporal changes of shared genetic drift of ancient populations from Yangtze River surrounding region in southern East Asia assessed via *$f_4$(Liangdao1_EN, Inland/Coastal Neolithic/Bronze Age southern East Asian; Studied inland TK/Sinitic, Mbuti)*.**
The dashed blue line indicates the zero $f_4$ value. The red asterisk indicates the absolute Z-score value larger than 3, the red point for absolute Z-score value ranging from two to three, and the green point for absolute Z-score value ranging from zero to two. The thick bar denotes two standard errors, and the thin bar for three standard errors. Figures were grouped via the second population in the $f_4$-statistics of Inland/Coastal Neolithic/Bronze Age southern East Asian, which was labeled as the red colour with some significant negative values. Significant negative values denoted the included second populations harboured more central/southern Sinitic or TK related ancestry. Significant positive $f_4$ values indicated the first population had more central/southern Sinitic or TK related ancestry.

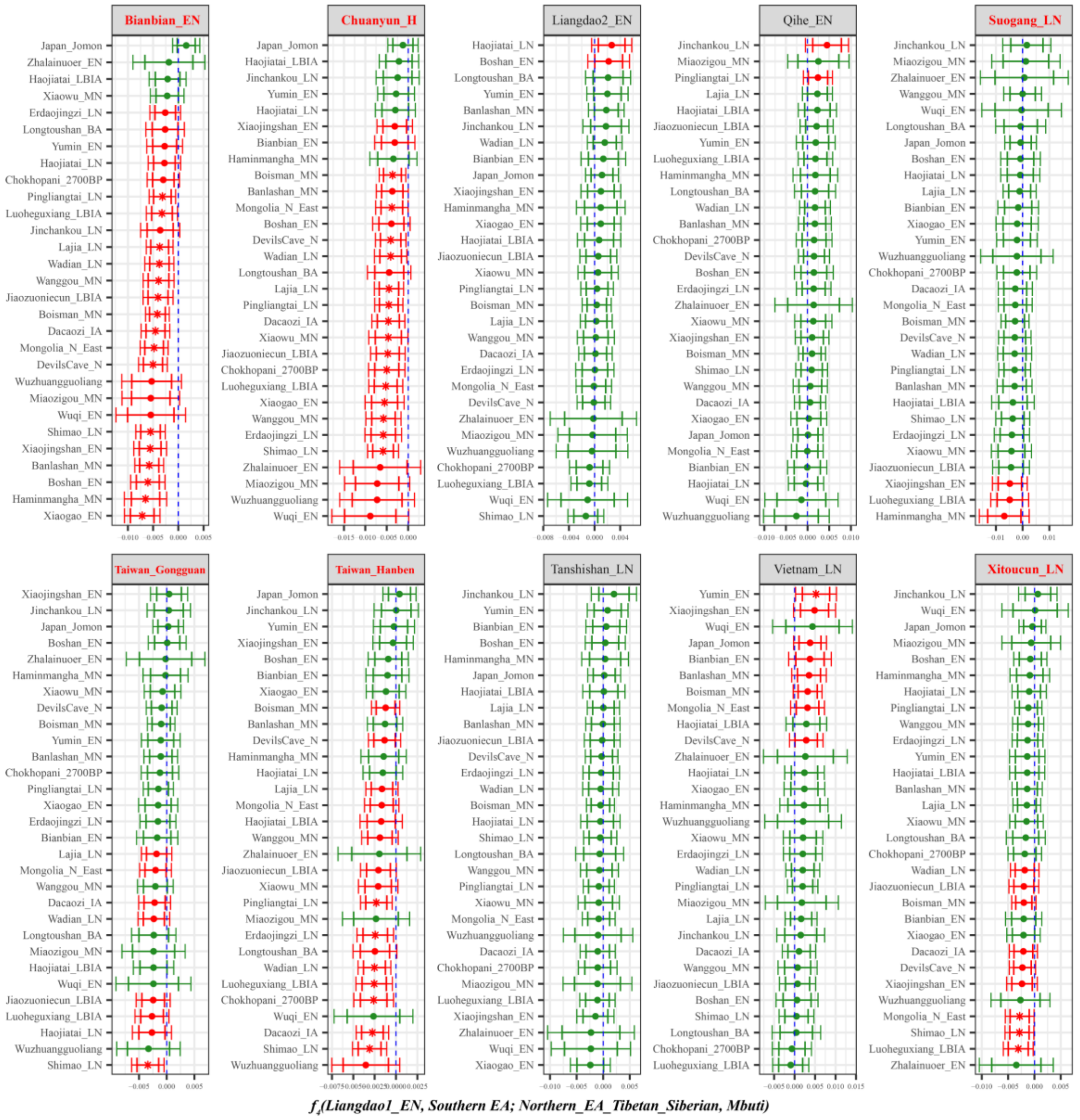

**Supplementary Fig. 50B. Temporal changes of shared genetic drift of ancient populations from Yangtze River surrounding region in southern East Asia assessed via *$f_4$(Liangdao1_EN, Inland/Coastal Neolithic/Bronze Age southern East Asian; Inland/Coastal Neolithic/Bronze Age northern East Asian/Tibet Plateau/Siberia/Japan, Mbuti)*.**

The dashed blue line indicates the zero $f_4$ value. The red asterisk indicates the absolute Z-score value larger than 3, the red point for absolute Z-score value ranging from two to three, and the green point for absolute Z-score value ranging from zero to two. The thick bar denotes two standard errors, and the thin bar for three standard errors. Figures were grouped via the second population in the $f_4$-statistics of Inland/Coastal Neolithic/Bronze Age southern East Asian, which was labeled as the red colour with some significant negative values. Significant negative values denoted the included second populations harboured more ancient northern East Asian related ancestry. Significant positive $f_4$ values indicated the first population had more ancient northern East Asian related ancestry.

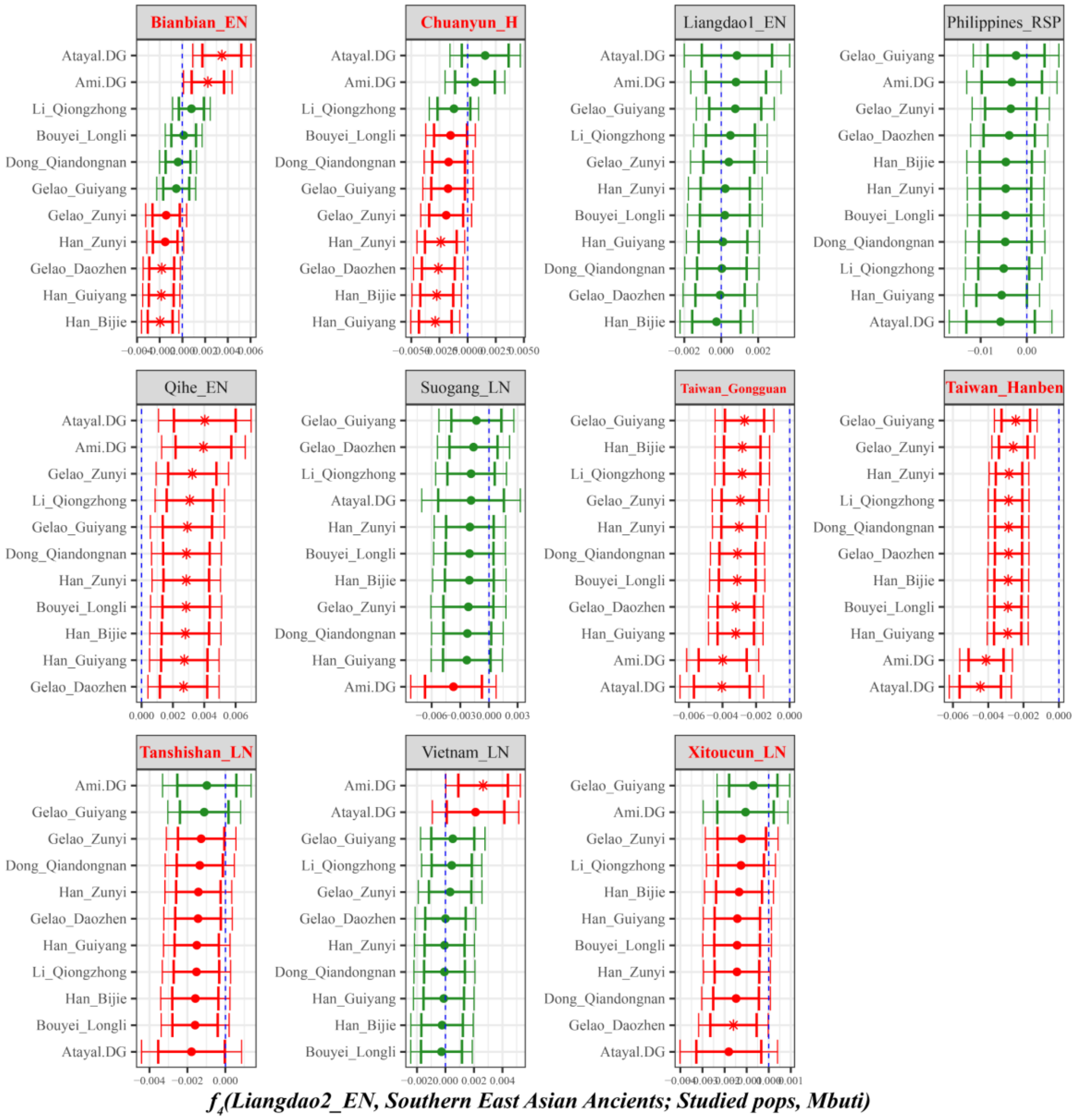

**Supplementary Fig. 51A. Temporal changes of shared genetic drift of ancient populations from Yangtze River surrounding region in southern East Asia assessed via *$f_4$(Liangdao2_EN, Inland/Coastal Neolithic/Bronze Age southern East Asian; Studied inland TK/Sinitic, Mbuti)*.**
The dashed blue line indicates the zero $f_4$ value. The red asterisk indicates the absolute Z-score value larger than 3, the red point for absolute Z-score value ranging from two to three, and the green point for absolute Z-score value ranging from zero to two. The thick bar denotes two standard errors, and the thin bar for three standard errors. Figures were grouped via the second population in the $f_4$-statistics of Inland/Coastal Neolithic/Bronze Age southern East Asian, which was labeled as the red colour with some significant negative values. Significant negative values denoted the included second populations harboured more central/southern Sinitic or TK related ancestry. Significant positive $f_4$ values indicated the first population had more central/southern Sinitic or TK related ancestry.

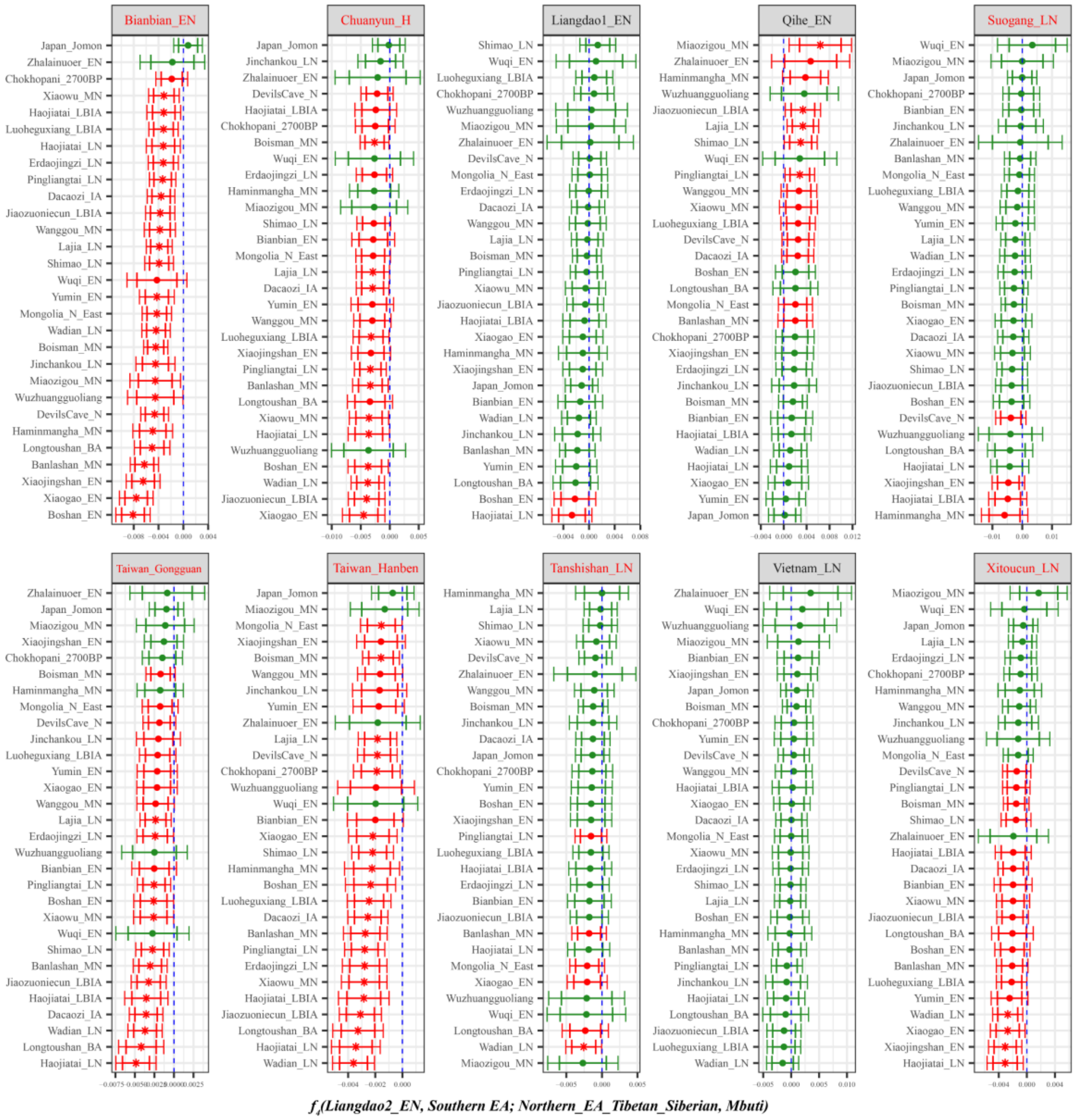

**Supplementary Fig. 51B. Temporal changes of shared genetic drift of ancient populations from Yangtze River surrounding region in southern East Asia assessed via *f₄*(Liangdao2_EN, Inland/Coastal Neolithic/Bronze Age southern East Asian; Inland/Coastal Neolithic/Bronze Age northern East Asian/Tibet Plateau/Siberia/Japan, Mbuti).**

The dashed blue line indicates the zero $f_4$ value. The red asterisk indicates the absolute Z-score value larger than 3, the red point for absolute Z-score value ranging from two to three, and the green point for absolute Z-score value ranging from zero to two. The thick bar denotes two standard errors, and the thin bar for three standard errors. Figures were grouped via the second population in the $f_4$-statistics of Inland/Coastal Neolithic/Bronze Age southern East Asian, which was labeled as the red colour with some significant negative values. Significant negative values denoted the included second populations harboured more ancient northern East Asian related ancestry. Significant positive $f_4$ values indicated the first population had more ancient northern East Asian related ancestry.

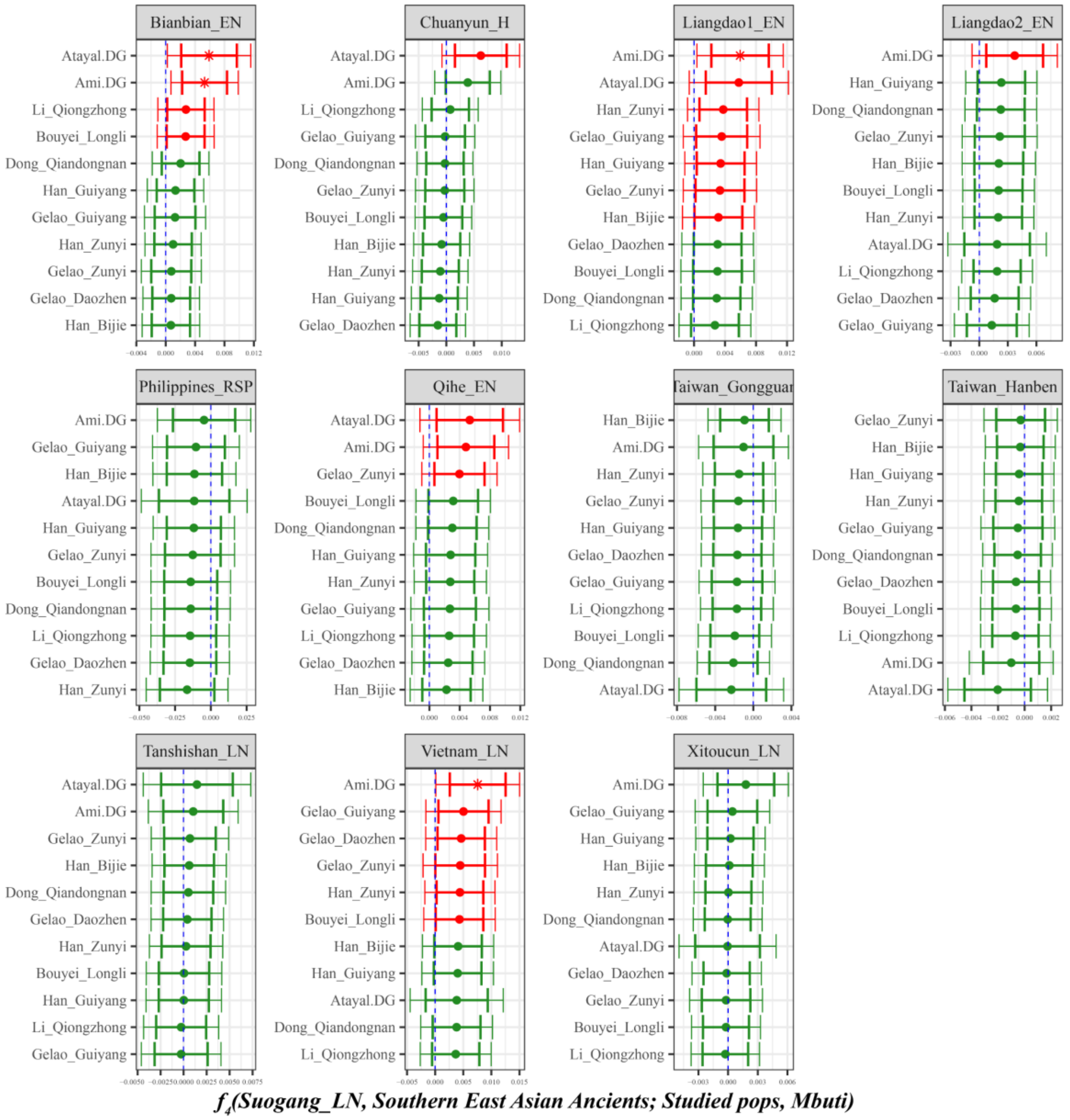

**Supplementary Fig. 52A. Temporal changes of shared genetic drift of ancient populations from Yangtze River surrounding region in southern East Asia assessed via *$f_4$(Suogang_LN, Inland/Coastal Neolithic/Bronze Age southern East Asian; Studied inland TK/Sinitic, Mbuti)*.**

The dashed blue line indicates the zero $f_4$ value. The red asterisk indicates the absolute Z-score value larger than 3, the red point for absolute Z-score value ranging from two to three, and the green point for absolute Z-score value ranging from zero to two. The thick bar denotes two standard errors, and the thin bar for three standard errors. Figures were grouped via the second population in the $f_4$-statistics of Inland/Coastal Neolithic/Bronze Age southern East Asian, which was labeled as the red colour with some significant negative values. Significant negative values denoted the included second populations harboured more central/southern Sinitic or TK related ancestry. Significant positive $f_4$ values indicated the first population had more central/southern Sinitic or TK related ancestry.

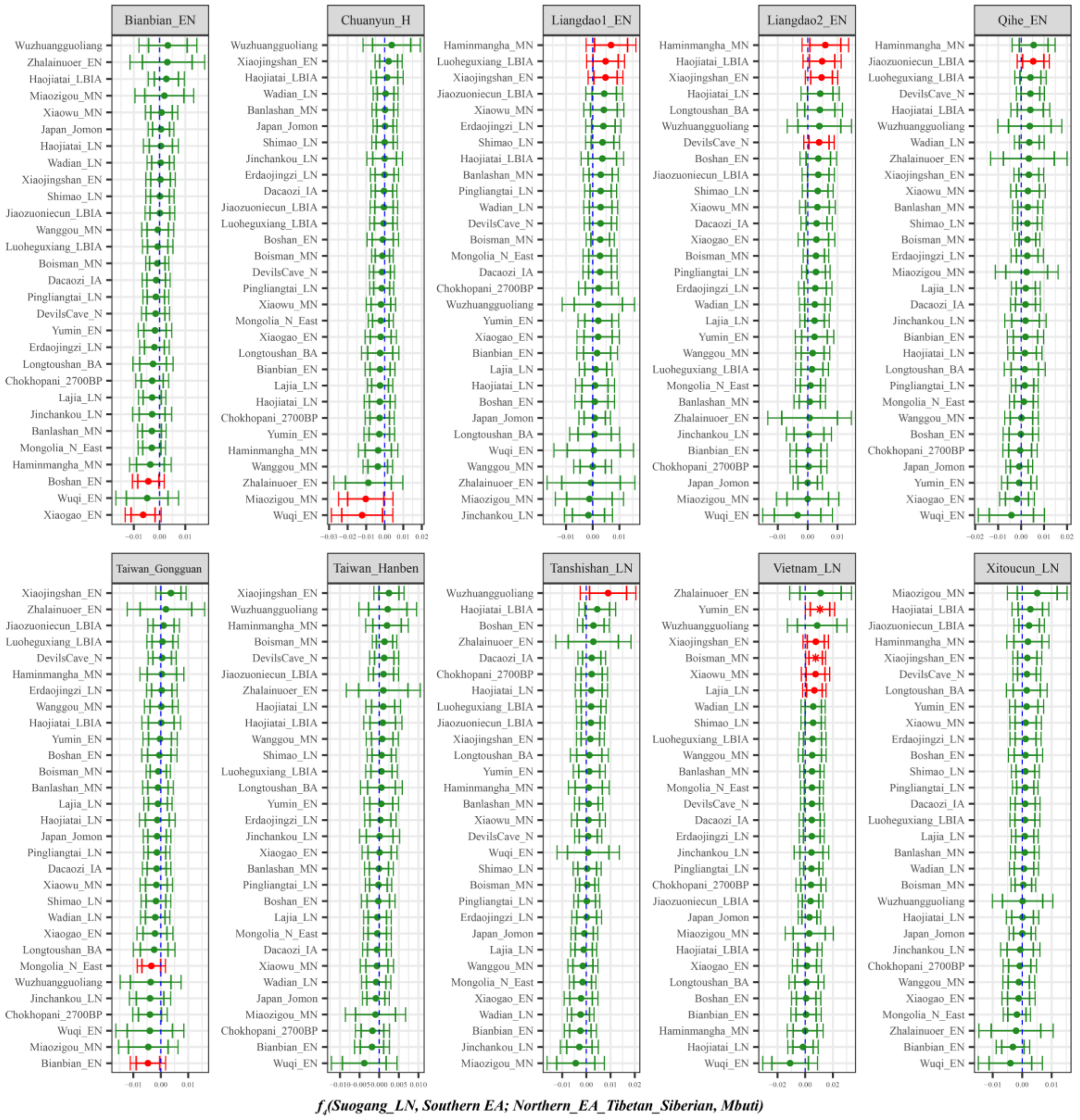

**Supplementary Fig. 52B. Temporal changes of shared genetic drift of ancient populations from Yangtze River surrounding region in southern East Asia assessed via *$f_4$(Suogang_LN, Inland/Coastal Neolithic/Bronze Age southern East Asian; Inland/Coastal Neolithic/Bronze Age northern East Asian/Tibet Plateau/Siberia/Japan, Mbuti)*.**

The dashed blue line indicates the zero $f_4$ value. The red asterisk indicates the absolute Z-score value larger than 3, the red point for absolute Z-score value ranging from two to three, and the green point for absolute Z-score value ranging from zero to two. The thick bar denotes two standard errors, and the thin bar for three standard errors. Figures were grouped via the second population in the $f_4$-statistics of Inland/Coastal Neolithic/Bronze Age southern East Asian, which was labeled as the red colour with some significant negative values. Significant negative values denoted the included second populations harboured more ancient northern East Asian related ancestry. Significant positive $f_4$ values indicated the first population had more ancient northern East Asian related ancestry.

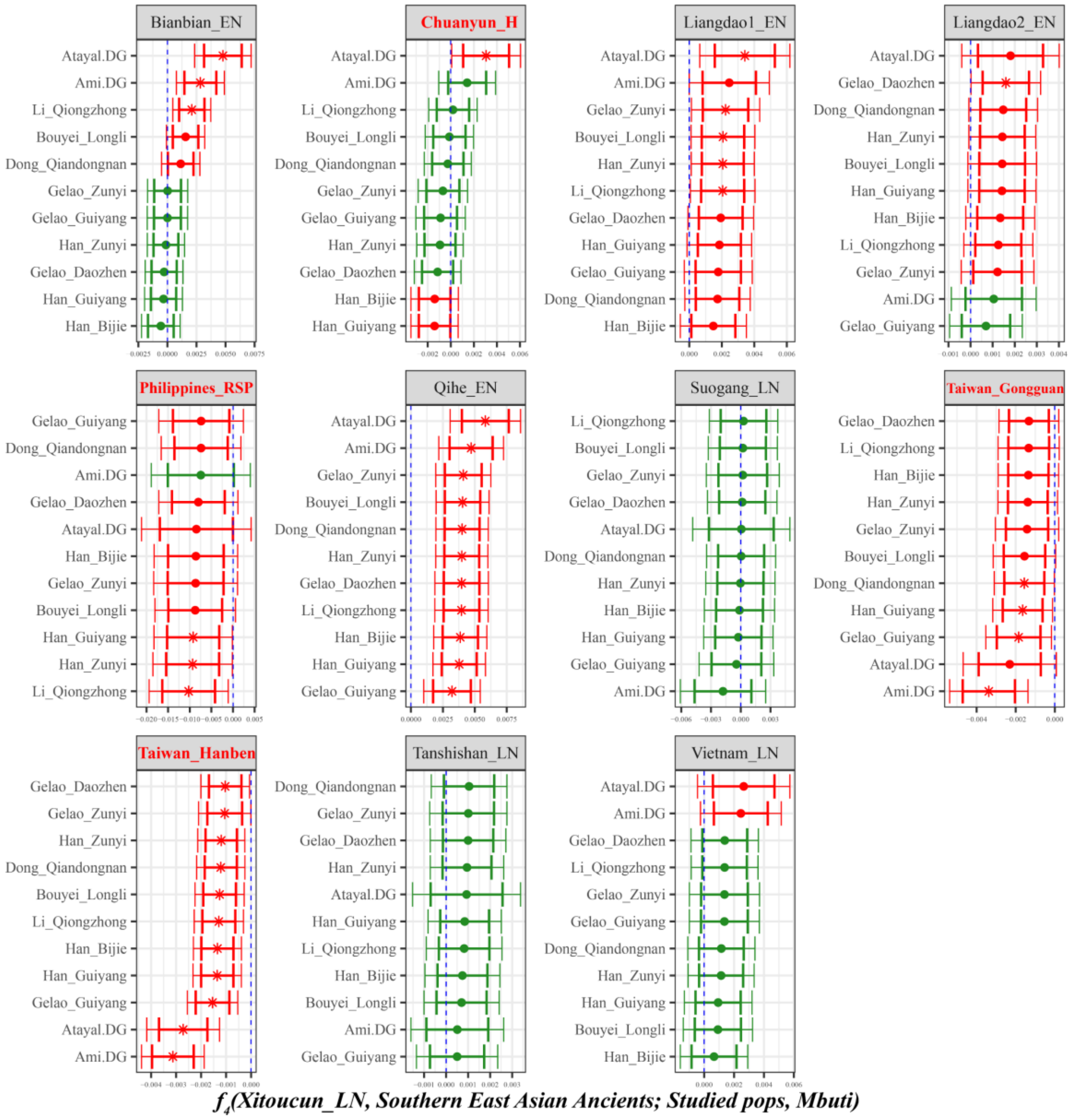

**Supplementary Fig. 53A. Temporal changes of shared genetic drift of ancient populations from Yangtze River surrounding region in southern East Asia assessed via *f*4*(Xitoucun_LN, Inland/Coastal Neolithic/Bronze Age southern East Asian; Studied inland TK/Sinitic, Mbuti)*.**
The dashed blue line indicates the zero $f_4$ value. The red asterisk indicates the absolute Z-score value larger than 3, the red point for absolute Z-score value ranging from two to three, and the green point for absolute Z-score value ranging from zero to two. The thick bar denotes two standard errors, and the thin bar for three standard errors. Figures were grouped via the second population in the $f_4$-statistics of Inland/Coastal Neolithic/Bronze Age southern East Asian, which was labeled as the red colour with some significant negative values. Significant negative values denoted the included second populations harboured more central/southern Sinitic or TK related ancestry. Significant positive $f_4$ values indicated the first population had more central/southern Sinitic or TK related ancestry.

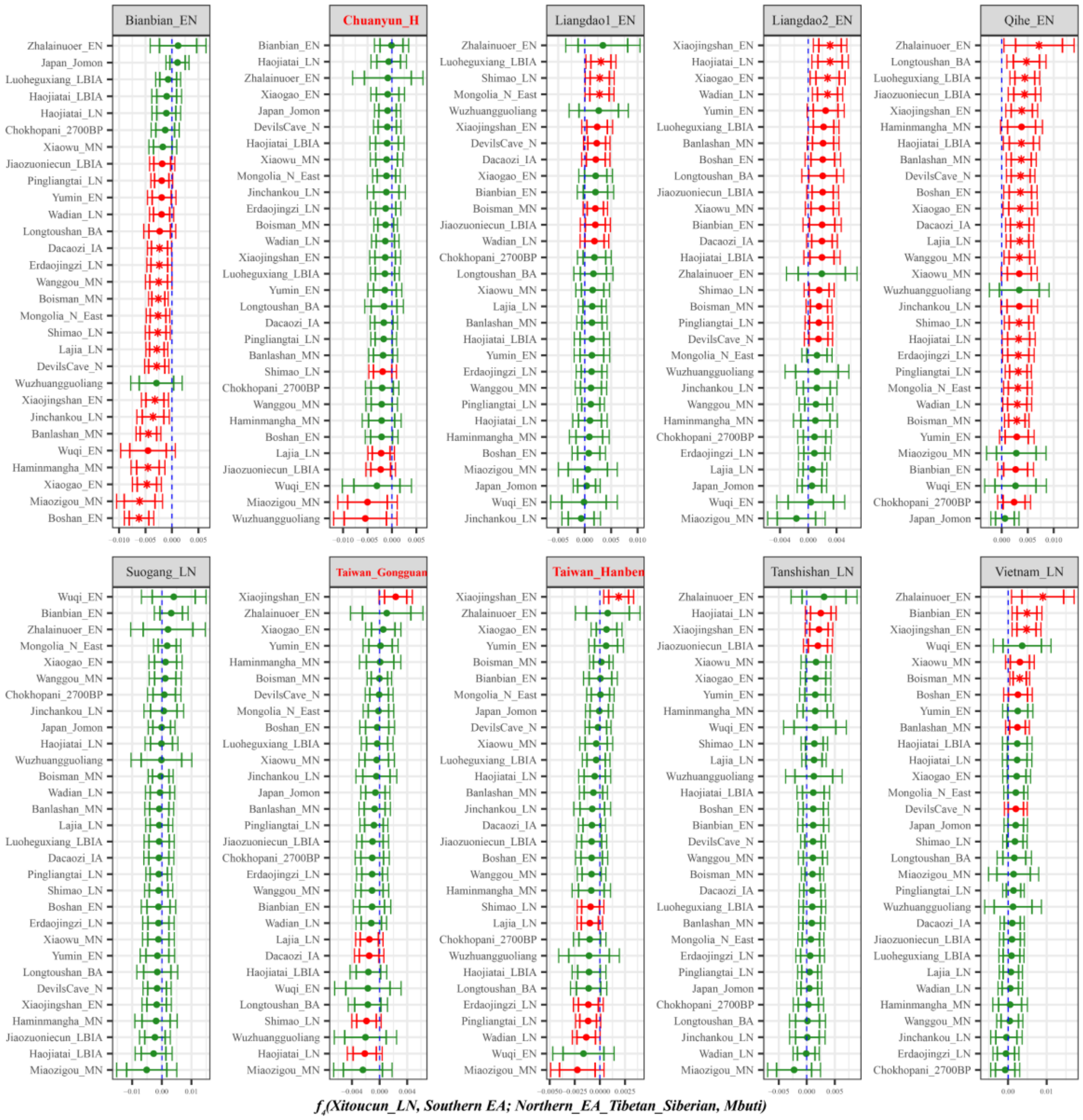

**Supplementary Fig. 53B. Temporal changes of shared genetic drift of ancient populations from Yangtze River surrounding region in southern East Asia assessed via *$f_4$(Xitoucun_LN, Inland/Coastal Neolithic/Bronze Age southern East Asian; Inland/Coastal Neolithic/Bronze Age northern East Asian/Tibet Plateau/Siberia/Japan, Mbuti)*.**

The dashed blue line indicates the zero $f_4$ value. The red asterisk indicates the absolute Z-score value larger than 3, the red point for absolute Z-score value ranging from two to three, and the green point for absolute Z-score value ranging from zero to two. The thick bar denotes two standard errors, and the thin bar for three standard errors. Figures were grouped via the second population in the $f_4$-statistics of Inland/Coastal Neolithic/Bronze Age southern East Asian, which was labeled as the red colour with some significant negative values. Significant negative values denoted the included second populations harboured more ancient northern East Asian related ancestry. Significant positive $f_4$ values indicated the first population had more ancient northern East Asian related ancestry.

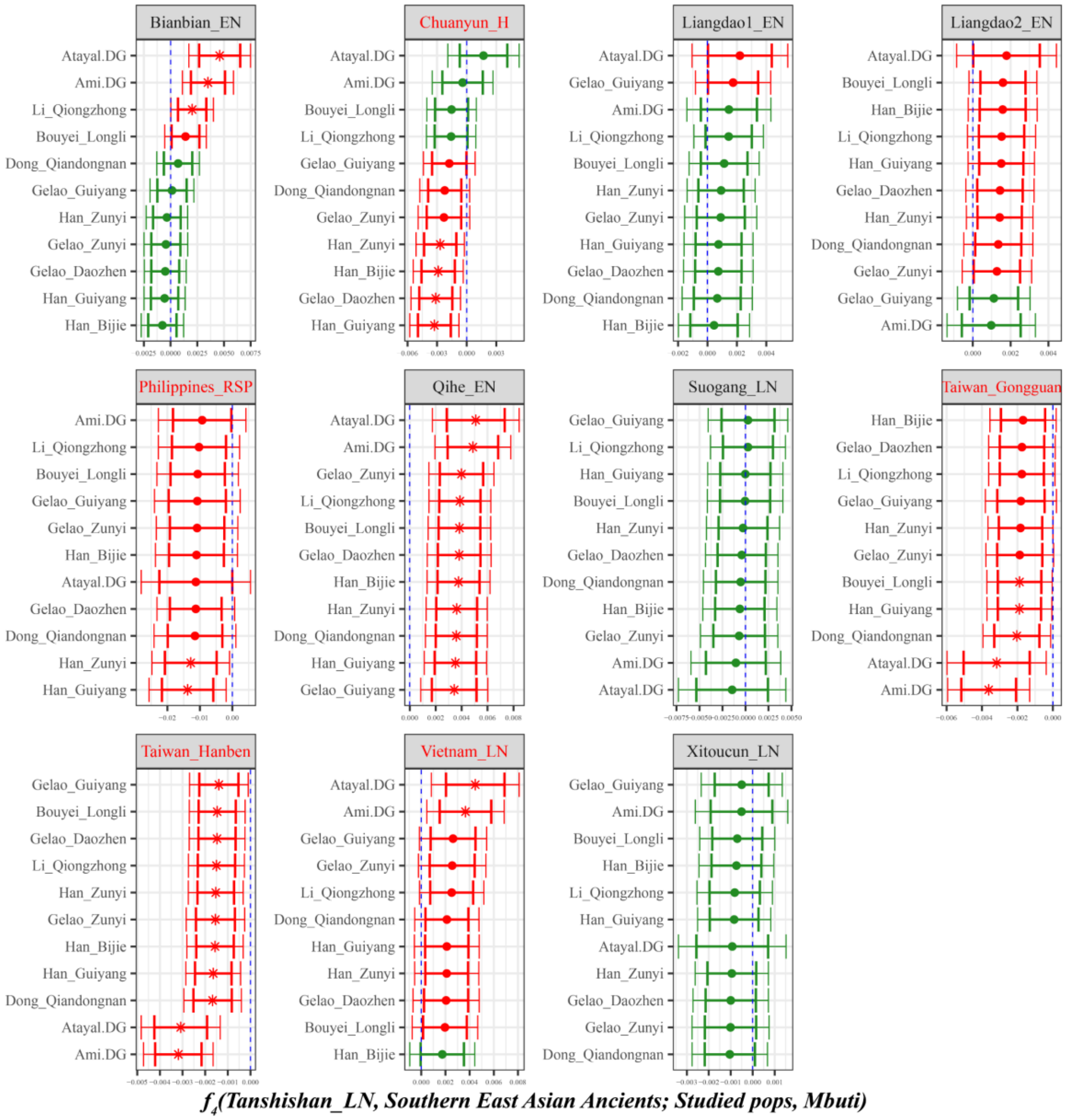

**Supplementary Fig. 54A. Temporal changes of shared genetic drift of ancient populations from Yangtze River surrounding region in southern East Asia assessed via *$f_4$(Tanshishan_LN, Inland/Coastal Neolithic/Bronze Age southern East Asian; Studied inland TK/Sinitic, Mbuti)*.**
The dashed blue line indicates the zero $f_4$ value. The red asterisk indicates the absolute Z-score value larger than 3, the red point for absolute Z-score value ranging from two to three, and the green point for absolute Z-score value ranging from zero to two. The thick bar denotes two standard errors, and the thin bar for three standard errors. Figures were grouped via the second population in the $f_4$-statistics of Inland/Coastal Neolithic/Bronze Age southern East Asian, which was labeled as the red colour with some significant negative values. Significant negative values denoted the included second populations harboured more central/southern Sinitic or TK related ancestry. Significant positive $f_4$ values indicated the first population had more central/southern Sinitic or TK related ancestry.

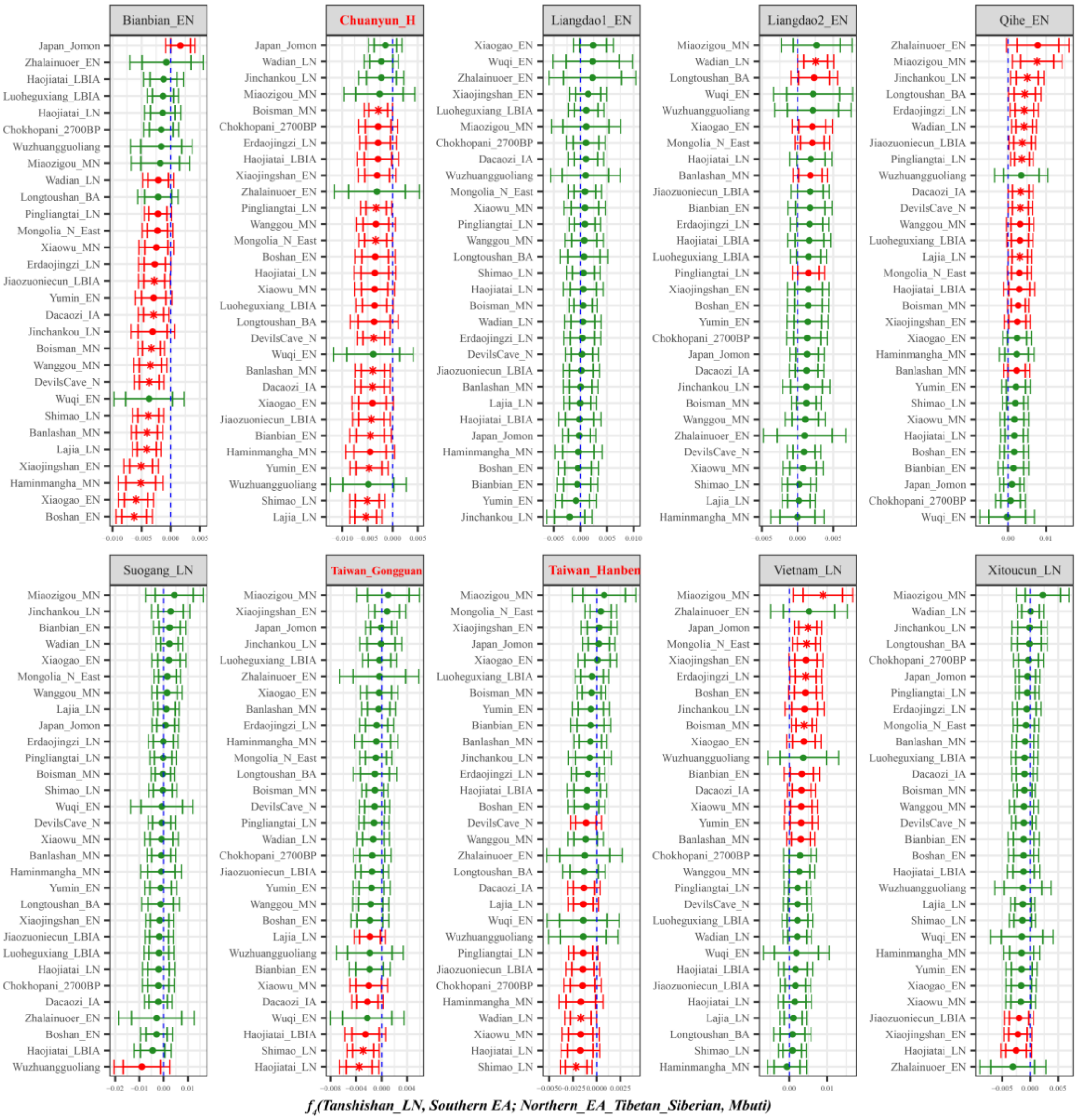

**Supplementary Fig. 54B. Temporal changes of shared genetic drift of ancient populations from Yangtze River surrounding region in southern East Asia assessed via *$f_4$(Tanshishan_LN, Inland/Coastal Neolithic/Bronze Age southern East Asian; Inland/Coastal Neolithic/Bronze Age northern East Asian/Tibet Plateau/Siberia/Japan, Mbuti)*.**

The dashed blue line indicates the zero $f_4$ value. The red asterisk indicates the absolute Z-score value larger than 3, the red point for absolute Z-score value ranging from two to three, and the green point for absolute Z-score value ranging from zero to two. The thick bar denotes two standard errors, and the thin bar for three standard errors. Figures were grouped via the second population in the $f_4$-statistics of Inland/Coastal Neolithic/Bronze Age southern East Asian, which was labeled as the red colour with some significant negative values. Significant negative values denoted the included second populations harboured more ancient northern East Asian related ancestry. Significant positive $f_4$ values indicated the first population had more ancient northern East Asian related ancestry.

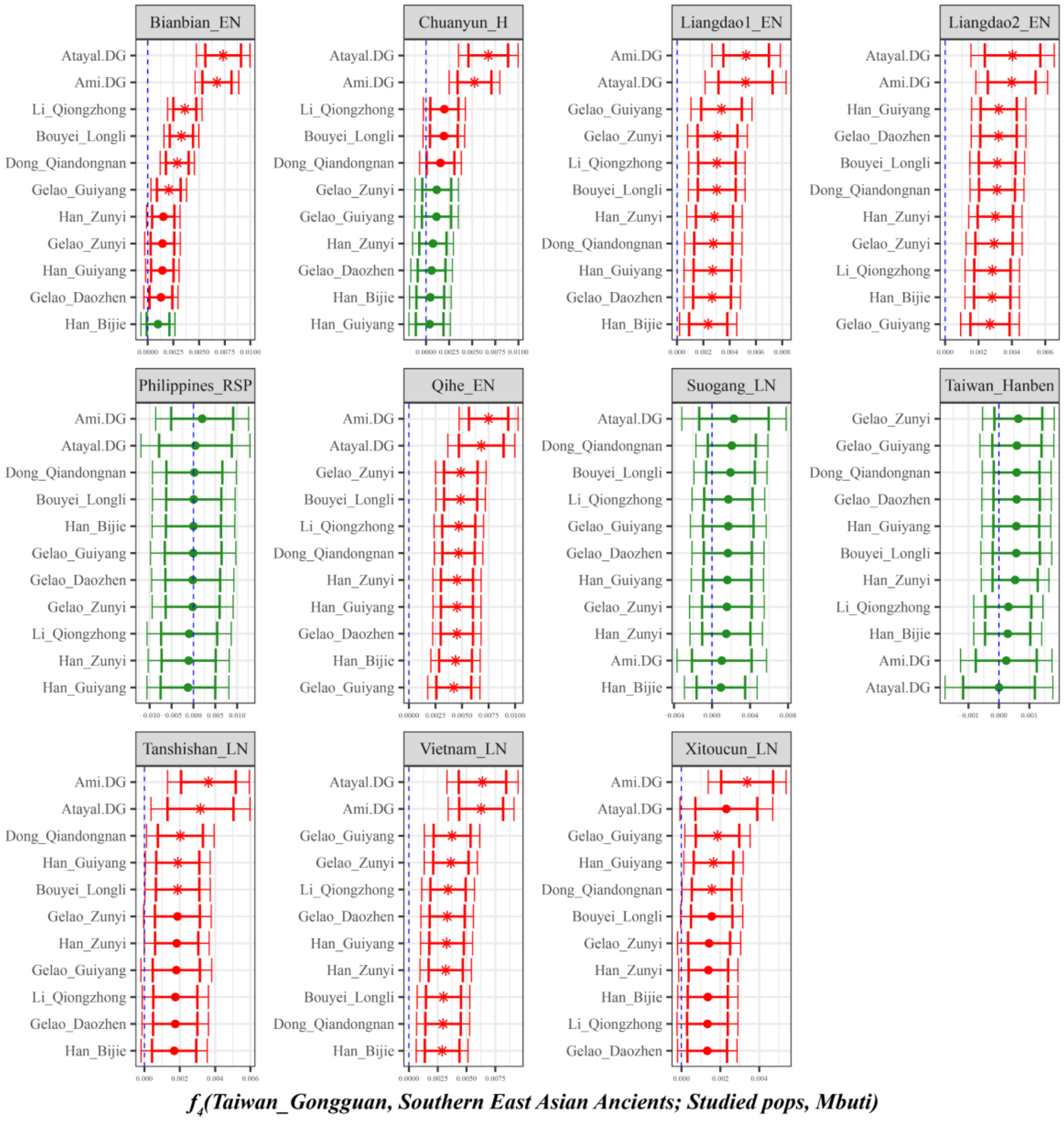

**Supplementary Fig. 55A. Temporal changes of shared genetic drift of ancient populations from Yangtze River surrounding region in southern East Asia assessed via *$f_4$(Taiwan_Gongguan, Inland/Coastal Neolithic/Bronze Age southern East Asian; Studied inland TK/Sinitic, Mbuti)*.**

The dashed blue line indicates the zero $f_4$ value. The red asterisk indicates the absolute Z-score value larger than 3, the red point for absolute Z-score value ranging from two to three, and the green point for absolute Z-score value ranging from zero to two. The thick bar denotes two standard errors, and the thin bar for three standard errors. Figures were grouped via the second population in the $f_4$-statistics of Inland/Coastal Neolithic/Bronze Age southern East Asian, which was labeled as the red colour with some significant negative values. Significant negative values denoted the included second populations harboured more central/southern Sinitic or TK related ancestry. Significant positive $f_4$ values indicated the first population had more central/southern Sinitic or TK related ancestry.

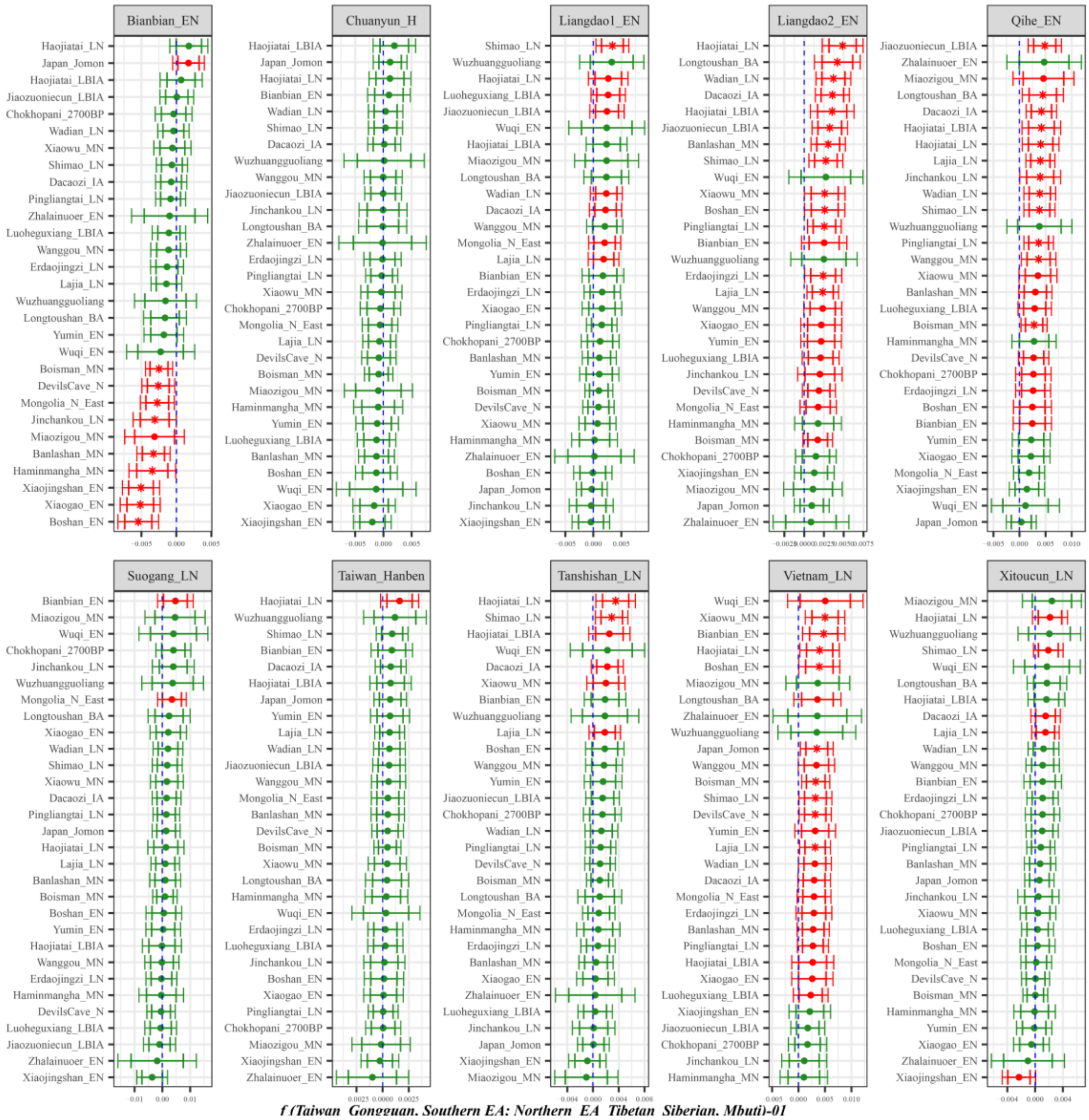

**Supplementary Fig. 55B. Temporal changes of shared genetic drift of ancient populations from Yangtze River surrounding region in southern East Asia assessed via $f_4$*(Taiwan_Gongguan, Inland/Coastal Neolithic/Bronze Age southern East Asian; Inland/Coastal Neolithic/Bronze Age northern East Asian/Tibet Plateau/Siberia/Japan, Mbuti)*.**

The dashed blue line indicates the zero $f_4$ value. The red asterisk indicates the absolute Z-score value larger than 3, the red point for absolute Z-score value ranging from two to three, and the green point for absolute Z-score value ranging from zero to two. The thick bar denotes two standard errors, and the thin bar for three standard errors. Figures were grouped via the second population in the $f_4$-statistics of Inland/Coastal Neolithic/Bronze Age southern East Asian, which was labeled as the red colour with some significant negative values. Significant negative values denoted the included second populations harboured more ancient northern East Asian related ancestry. Significant positive $f_4$ values indicated the first population had more ancient northern East Asian related ancestry.

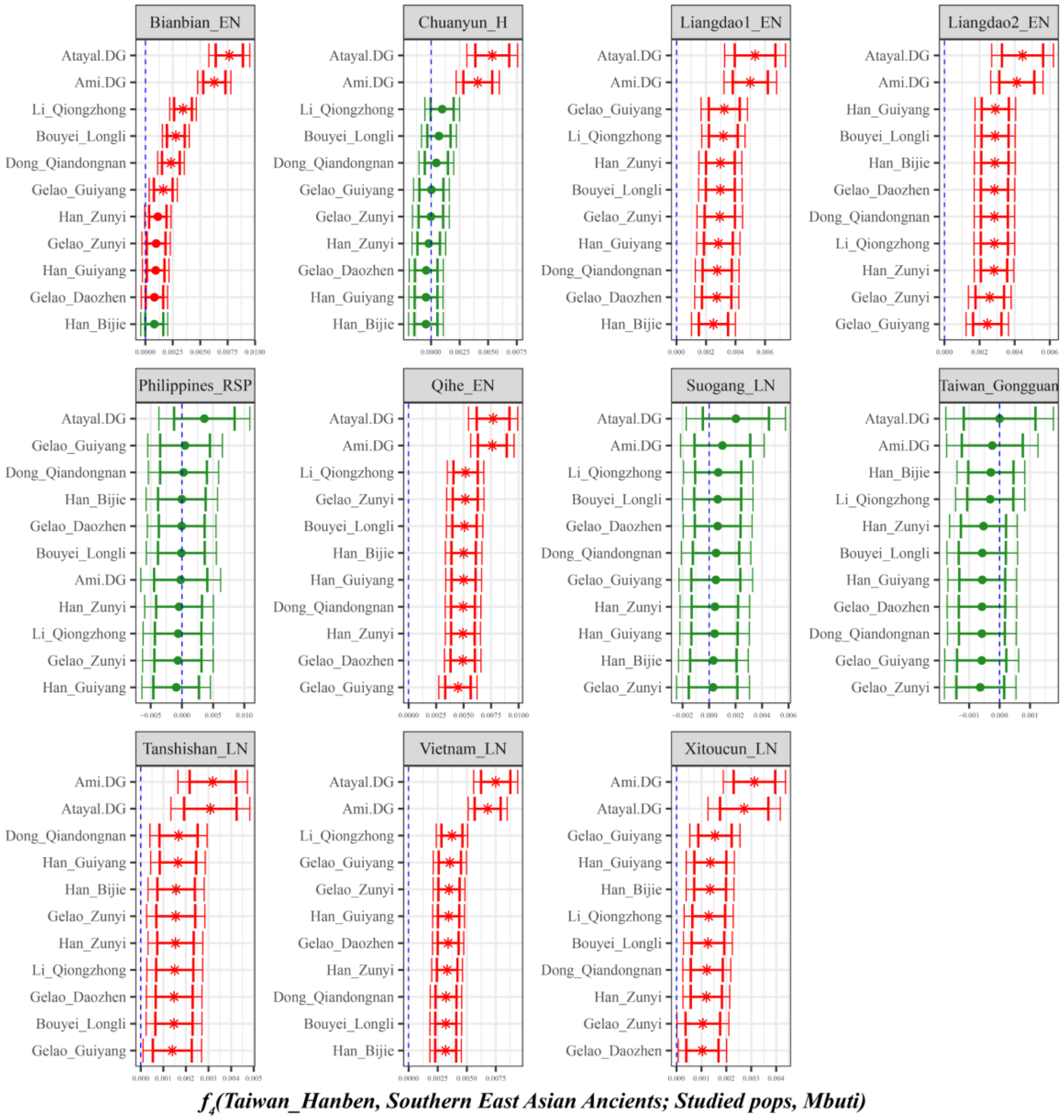

**Supplementary Fig. 56A. Temporal changes of shared genetic drift of ancient populations from Yangtze River surrounding region in southern East Asia assessed via *$f_4$(Taiwan_Hanben, Inland/Coastal Neolithic/Bronze Age southern East Asian; Studied inland TK/Sinitic, Mbuti)*.**

The dashed blue line indicates the zero $f_4$ value. The red asterisk indicates the absolute Z-score value larger than 3, the red point for absolute Z-score value ranging from two to three, and the green point for absolute Z-score value ranging from zero to two. The thick bar denotes two standard errors, and the thin bar for three standard errors. Figures were grouped via the second population in the $f_4$-statistics of Inland/Coastal Neolithic/Bronze Age southern East Asian, which was labeled as the red colour with some significant negative values. Significant negative values denoted the included second populations harboured more central/southern Sinitic or TK related ancestry. Significant positive $f_4$ values indicated the first population had more central/southern Sinitic or TK related ancestry.

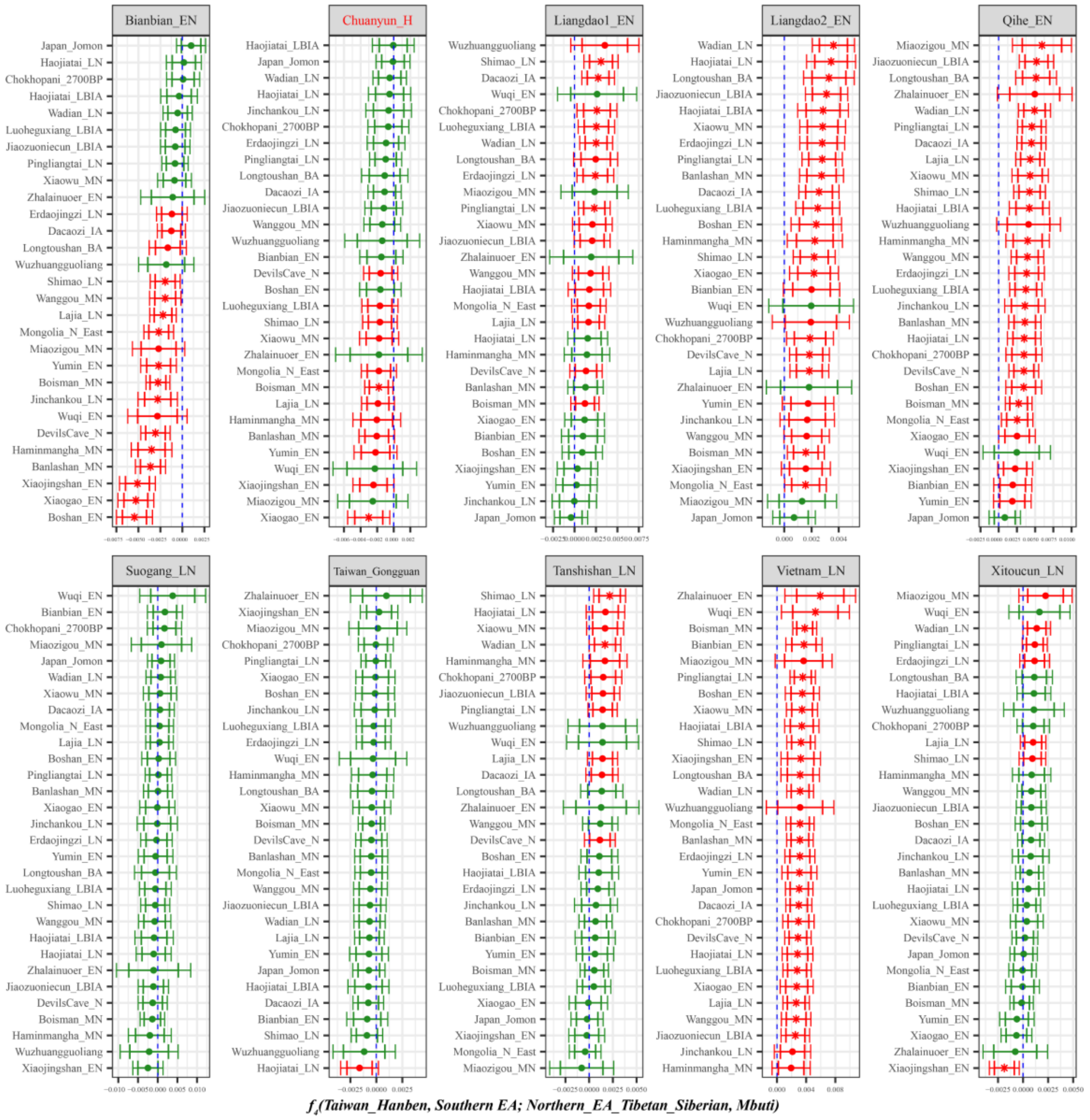

**Supplementary Fig. 56B. Temporal changes of shared genetic drift of ancient populations from Yangtze River surrounding region in southern East Asia assessed via $f_4$*(Taiwan_Hanben, Inland/Coastal Neolithic/Bronze Age southern East Asian; Inland/Coastal Neolithic/Bronze Age northern East Asian/Tibet Plateau/Siberia/Japan, Mbuti)*.**

The dashed blue line indicates the zero $f_4$ value. The red asterisk indicates the absolute Z-score value larger than 3, the red point for absolute Z-score value ranging from two to three, and the green point for absolute Z-score value ranging from zero to two. The thick bar denotes two standard errors, and the thin bar for three standard errors. Figures were grouped via the second population in the $f_4$-statistics of Inland/Coastal Neolithic/Bronze Age southern East Asian, which was labeled as the red colour with some significant negative values. Significant negative values denoted the included second populations harboured more ancient northern East Asian related ancestry. Significant positive $f_4$ values indicated the first population had more ancient northern East Asian related ancestry.

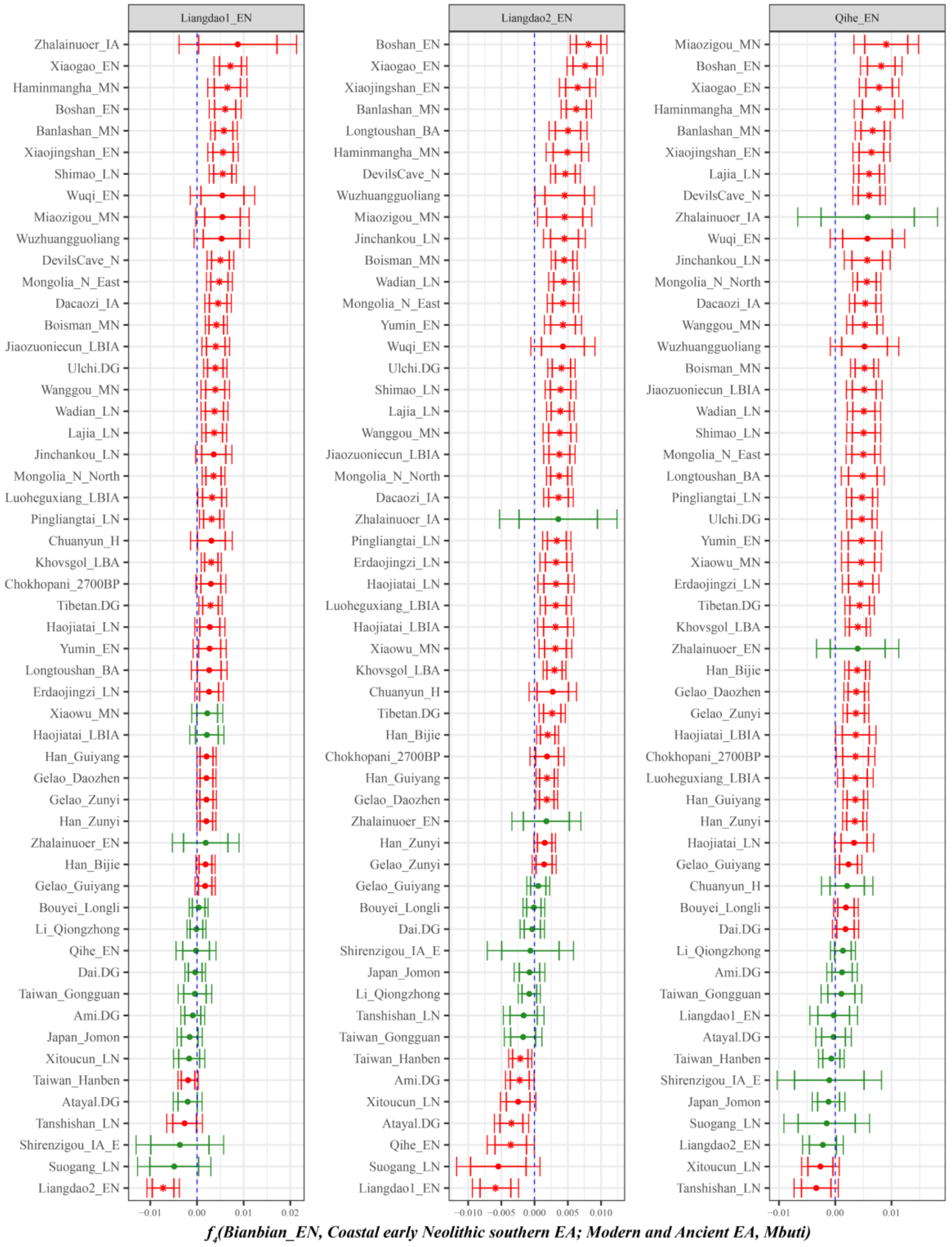

**Supplementary Fig. 57. Spatial difference of shared genetic drift of ancient populations from northern and southern East Asia assessed in the early Neolithic period via *$f_4$(Bianbian_EN, Inland/Coastal early Neolithic southern East Asian; Studied inland TK/Sinitic and East Asian ancients, Mbuti)*.**

The dashed blue line indicates the zero $f_4$ value. The red asterisk indicates the absolute Z-score value larger than 3, the red point for absolute Z-score value ranging from two to three, and the green point for absolute Z-score value ranging from zero to two. The thick bar denotes two standard errors, and the thin bar for three standard errors. Figures were grouped via the second population in the $f_4$-statistics of Inland/Coastal early Neolithic southern East Asian. Significant negative values denoted the studied inland TK/Sinitic and East Asian ancients shared more alleles with Inland/Coastal early Neolithic southern East Asian, while significant positive $f_4$ values indicated the targeted populations shared more alleles with Bianbian_EN related ancestral populations.

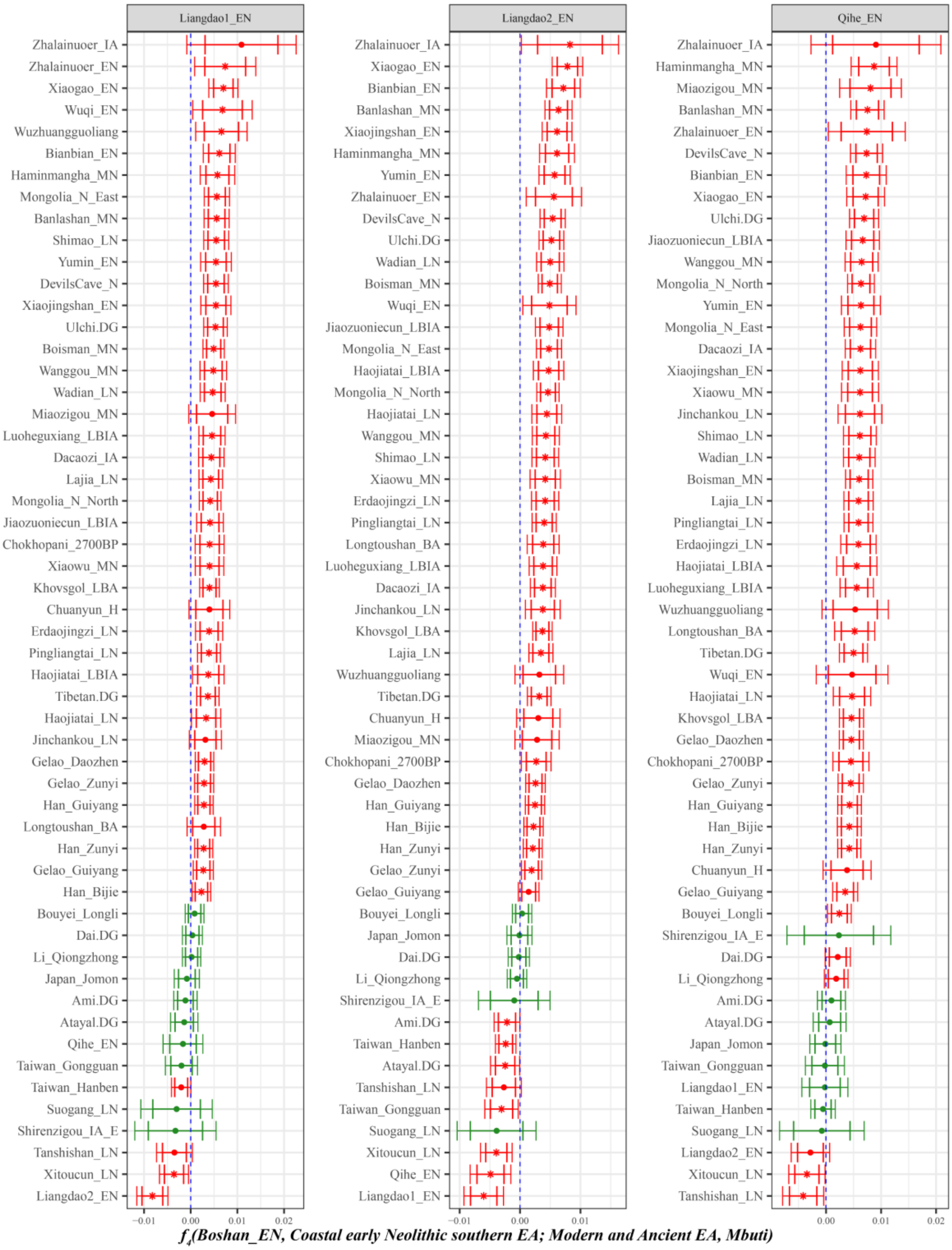

**Supplementary Fig. 58. Spatial difference of shared genetic drift of ancient populations from northern and southern East Asia assessed in the early Neolithic period via *$f_4$(Boshan_EN, Inland/Coastal early Neolithic southern East Asian; Studied inland TK/Sinitic and East Asian ancients, Mbuti)*.**

The dashed blue line indicates the zero $f_4$ value. The red asterisk indicates the absolute Z-score value larger than 3, the red point for absolute Z-score value ranging from two to three, and the green point for absolute Z-score value ranging from zero to two. The thick bar denotes two standard errors, and the thin bar for three standard errors. Figures were grouped via the second population in the $f_4$-statistics of Inland/Coastal early Neolithic southern East Asian. Significant negative values denoted the studied inland TK/Sinitic and East Asian ancients shared more alleles with Inland/Coastal early Neolithic southern East Asian, while significant positive $f_4$ values indicated the targeted populations shared more alleles with Boshan_EN related ancestral populations.

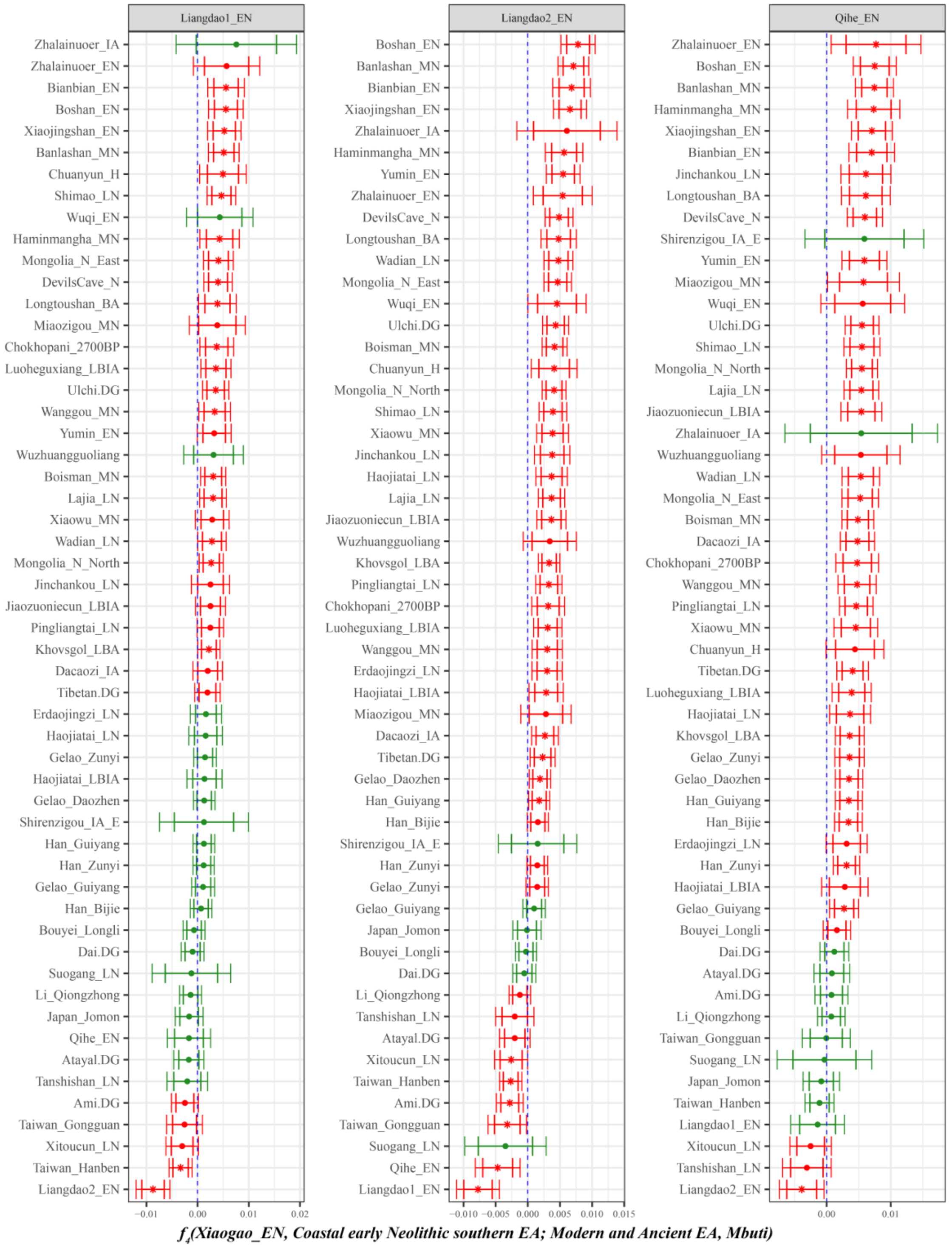

**Supplementary Fig. 59. Spatial difference of shared genetic drift of ancient populations from northern and southern East Asia assessed in the early Neolithic period via *$f_4$(Xiaogao_EN, Inland/Coastal early Neolithic southern East Asian; Studied inland TK/Sinitic and East Asian ancients, Mbuti)*.**

The dashed blue line indicates the zero $f_4$ value. The red asterisk indicates the absolute Z-score value larger than 3, the red point for absolute Z-score value ranging from two to three, and the green point for absolute Z-score value ranging from zero to two. The thick bar denotes two standard errors, and the thin bar for three standard errors. Figures were grouped via the second population in the $f_4$-statistics of Inland/Coastal early Neolithic southern East Asian. Significant negative values denoted the studied inland TK/Sinitic and East Asian ancients shared more alleles with Inland/Coastal early Neolithic southern East Asian, while significant positive $f_4$ values indicated the targeted populations shared more alleles with Xiaogao_EN related ancestral populations.

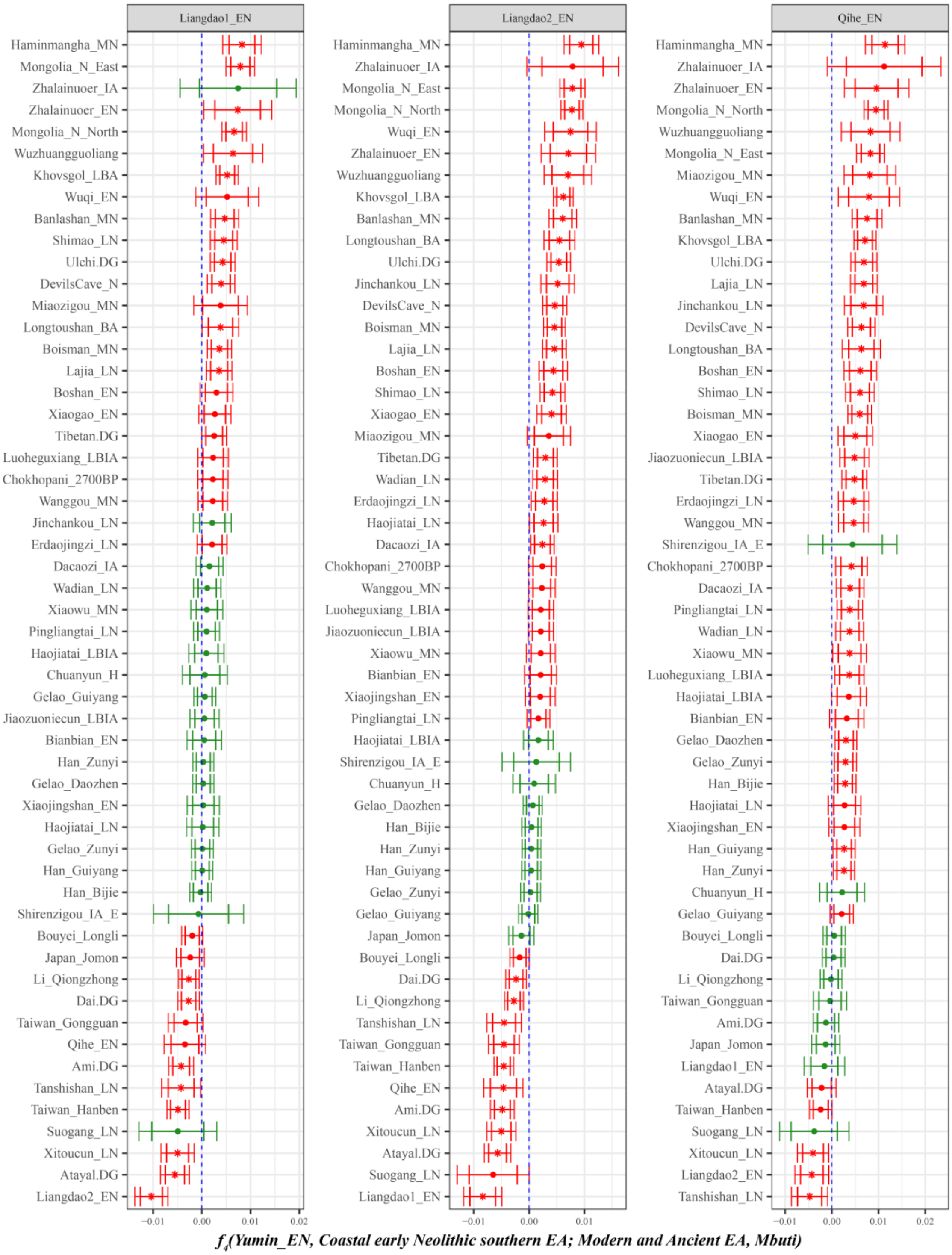

**Supplementary Fig. 60. Spatial difference of shared genetic drift of ancient populations from northern and southern East Asia assessed in the early Neolithic period via *$f_4$(Yumin_EN, Inland/Coastal early Neolithic southern East Asian; Studied inland TK/Sinitic and East Asian ancients, Mbuti)*.**

The dashed blue line indicates the zero $f_4$ value. The red asterisk indicates the absolute Z-score value larger than 3, the red point for absolute Z-score value ranging from two to three, and the green point for absolute Z-score value ranging from zero to two. The thick bar denotes two standard errors, and the thin bar for three standard errors. Figures were grouped via the second population in the $f_4$-statistics of Inland/Coastal early Neolithic southern East Asian. Significant negative values denoted the studied inland TK/Sinitic and East Asian ancients shared more alleles with Inland/Coastal early Neolithic southern East Asian, while significant positive $f_4$ values indicated the targeted populations shared more alleles with Yumin_EN related ancestral populations.

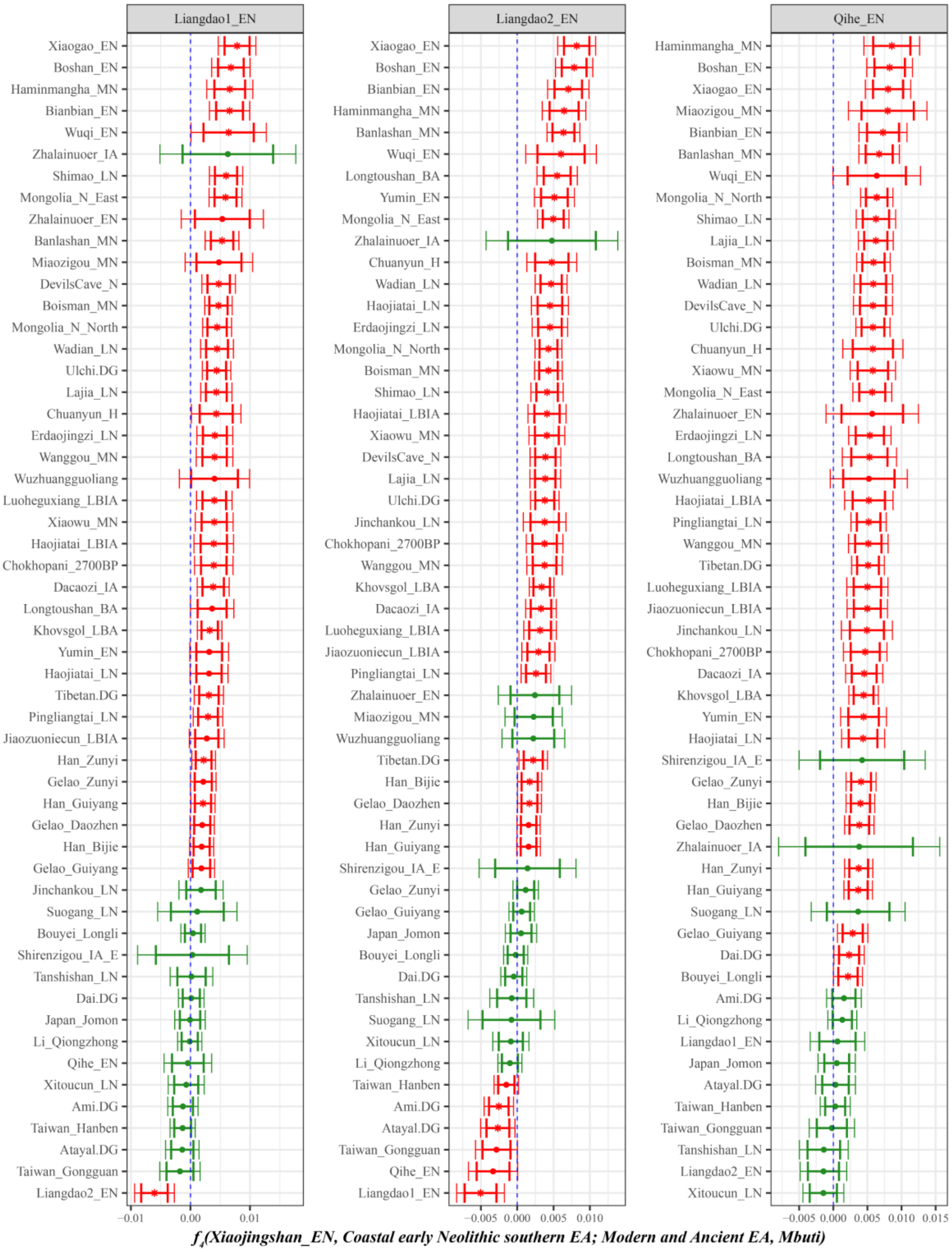

**Supplementary Fig. 61. Spatial difference of shared genetic drift of ancient populations from northern and southern East Asia assessed in the early Neolithic period via *$f_4$(Xiaojingshan_EN, Inland/Coastal early Neolithic southern East Asian; Studied inland TK/Sinitic and East Asian ancients, Mbuti)*.**

The dashed blue line indicates the zero $f_4$ value. The red asterisk indicates the absolute Z-score value larger than 3, the red point for absolute Z-score value ranging from two to three, and the green point for absolute Z-score value ranging from zero to two. The thick bar denotes two standard errors, and the thin

bar for three standard errors. Figures were grouped via the second population in the $f_4$-statistics of Inland/Coastal early Neolithic southern East Asian. Significant negative values denoted the studied inland TK/Sinitic and East Asian ancients shared more alleles with Inland/Coastal early Neolithic southern East Asian, while significant positive $f_4$ values indicated the targeted populations shared more alleles with Xiaojingshan_EN related ancestral populations.

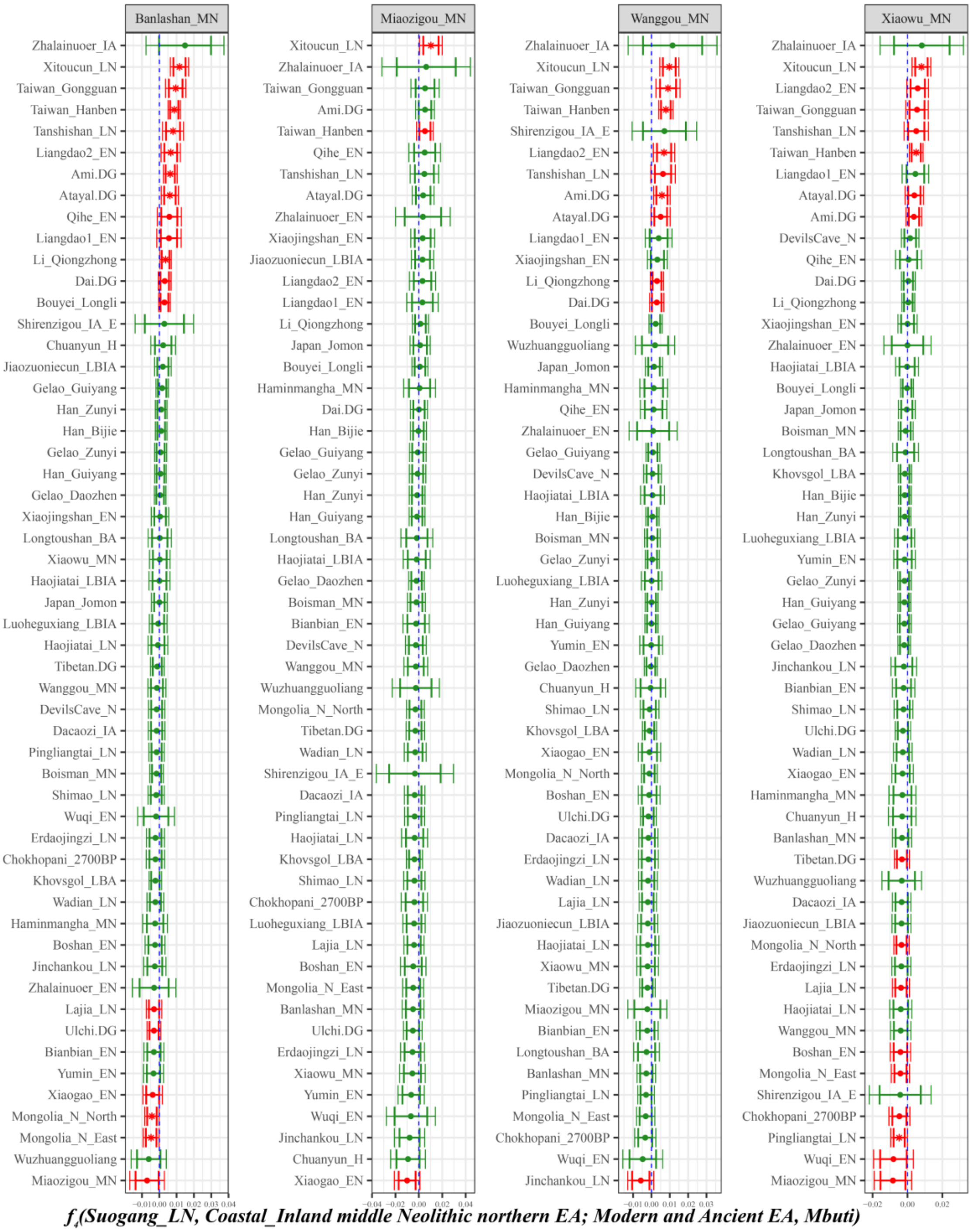

**Supplementary Fig. 62. Spatial difference of shared genetic drift of ancient populations from northern and southern East Asia assessed in the middle Neolithic period via $f_4$*(Suogang_LN, Inland/Coastal middle Neolithic northern East Asian; Studied inland TK/Sinitic and East Asian ancients, Mbuti)*.**

The dashed blue line indicates the zero $f_4$ value. The red asterisk indicates the absolute Z-score value larger than 3, the red point for absolute Z-score value ranging from two to three, and the green point for absolute Z-score value ranging from zero to two. The thick bar denotes two standard errors, and the thin

bar for three standard errors. Figures were grouped via the second population in the $f_4$-statistics of Inland/Coastal middle Neolithic northern East Asian. Significant negative values denoted the studied inland TK/Sinitic and East Asian ancients shared more alleles with Inland/Coastal middle Neolithic northern East Asian, while significant positive $f_4$ values indicated the targeted populations shared more alleles with Suogang_LN related ancestral populations.

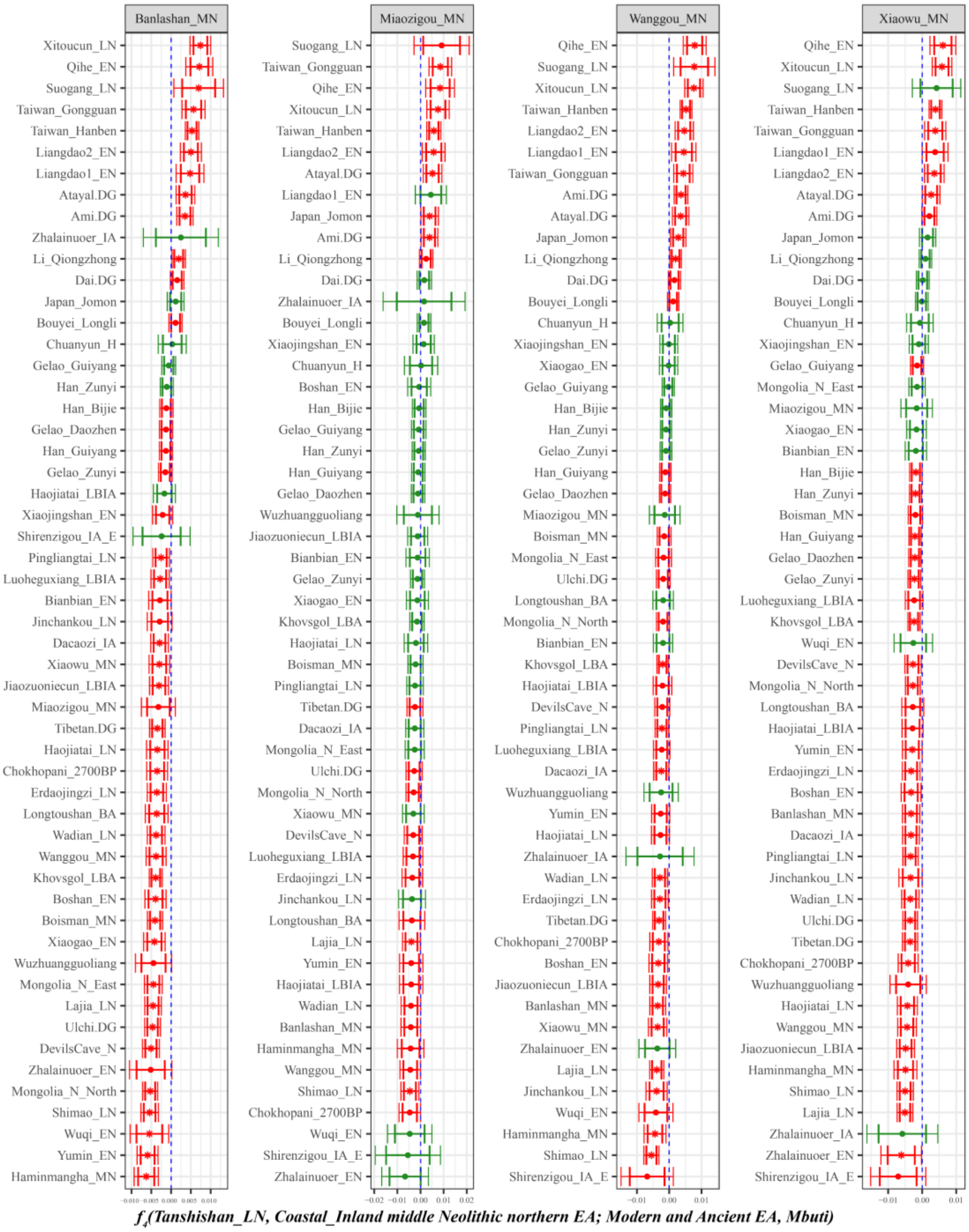

**Supplementary Fig. 63. Spatial difference of shared genetic drift of ancient populations from northern and southern East Asia assessed in the middle Neolithic period via $f_4$*(Tanshishan_LN, Inland/Coastal middle Neolithic northern East Asian; Studied inland TK/Sinitic and East Asian ancients, Mbuti)*.**

The dashed blue line indicates the zero $f_4$ value. The red asterisk indicates the absolute Z-score value larger than 3, the red point for absolute Z-score value ranging from two to three, and the green point for absolute Z-score value ranging from zero to two. The thick bar denotes two standard errors, and the thin bar for three standard errors. Figures were grouped via the second population in the $f_4$-statistics of Inland/Coastal middle Neolithic northern East Asian. Significant negative values denoted the studied inland TK/Sinitic and East Asian ancients shared more alleles with Inland/Coastal middle Neolithic northern East Asian, while significant positive $f_4$ values indicated the targeted populations shared more alleles with Tanshishan_LN related ancestral populations.

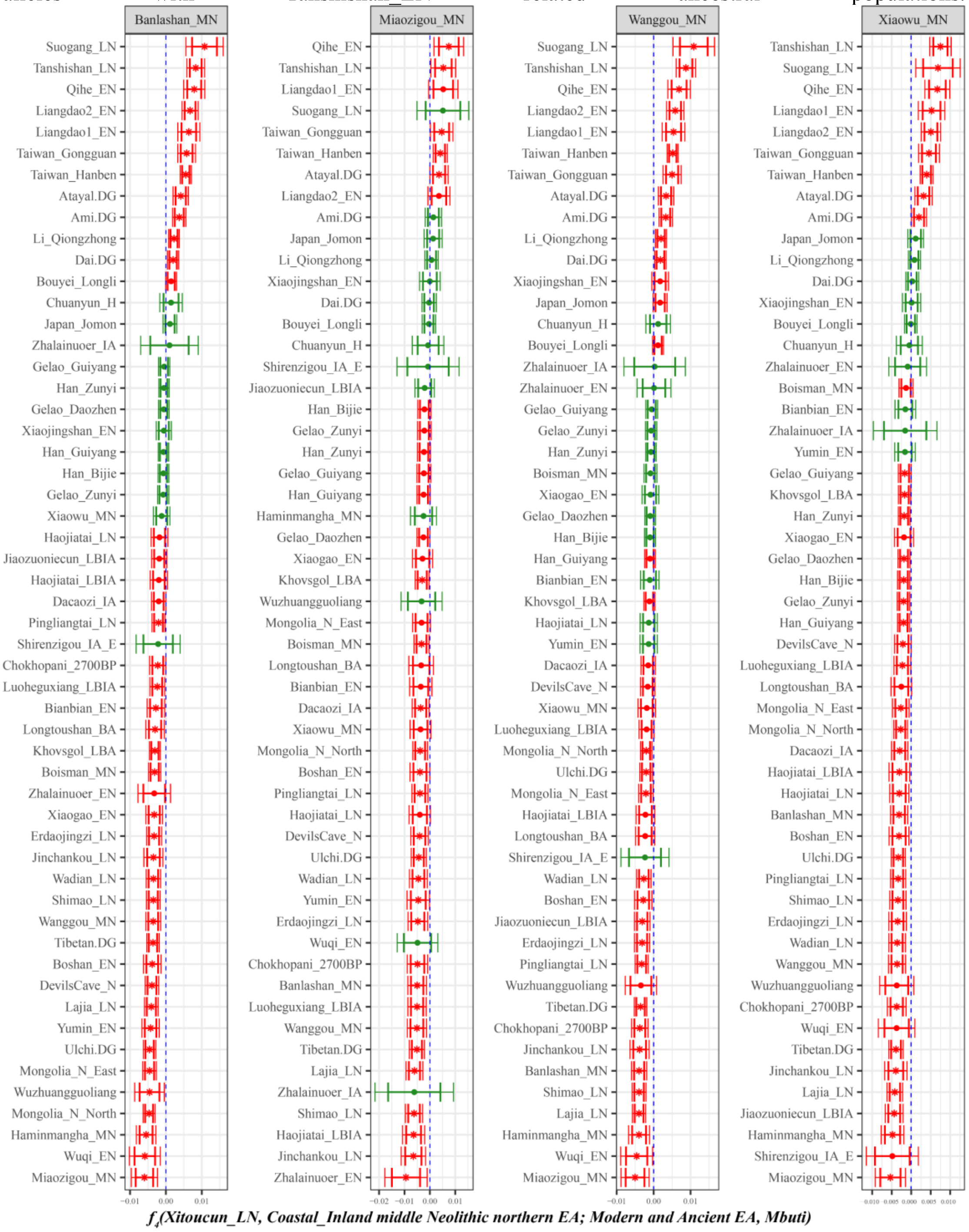

**Supplementary Fig. 64. Spatial difference of shared genetic drift of ancient populations from northern and southern East Asia assessed in the middle Neolithic period via $f_4$*(Xitoucun_LN, Inland/Coastal middle Neolithic northern East Asian; Studied inland TK/Sinitic and East Asian ancients, Mbuti)*.**

The dashed blue line indicates the zero $f_4$ value. The red asterisk indicates the absolute Z-score value larger than 3, the red point for absolute Z-score value ranging from two to three, and the green point for absolute Z-score value ranging from zero to two. The thick bar denotes two standard errors, and the thin bar for three standard errors. Figures were grouped via the second population in the $f_4$-statistics of Inland/Coastal middle Neolithic northern East Asian. Significant negative values denoted the studied inland TK/Sinitic and East Asian ancients shared more alleles with Inland/Coastal middle Neolithic northern East Asian, while significant positive $f_4$ values indicated the targeted populations shared more alleles with Xitoucun_LN related ancestral populations.

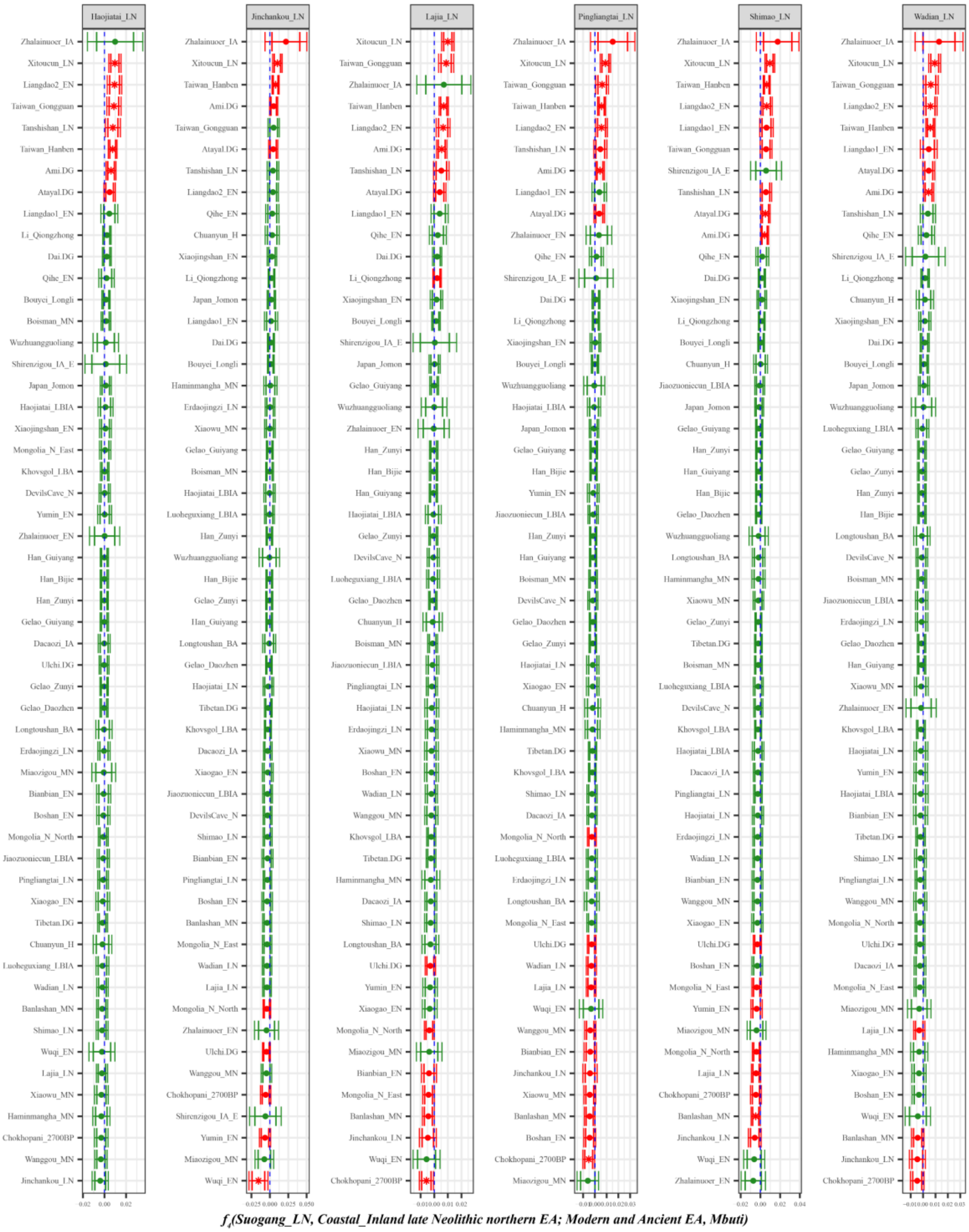

**Supplementary Fig. 65. Spatial difference of shared genetic drift of ancient populations from northern and southern East Asia assessed in the late Neolithic period via *$f_4$(Suogang_LN, Inland/Coastal late Neolithic northern East Asian; Studied inland TK/Sinitic and East Asian ancients, Mbuti)*.**

The dashed blue line indicates the zero $f_4$ value. The red asterisk indicates the absolute Z-score value larger than 3, the red point for absolute Z-score value ranging from two to three, and the green point for absolute Z-score value ranging from zero to two. The thick bar denotes two standard errors, and the thin bar for three standard errors. Figures were grouped via the second population in the $f_4$-statistics of Inland/Coastal late Neolithic northern East Asian. Significant negative values denoted the studied inland TK/Sinitic and East Asian ancients shared more alleles with Inland/Coastal late Neolithic northern East Asian, while significant positive $f_4$ values indicated the targeted populations shared more alleles with

Suogang_LN related ancestral populations.

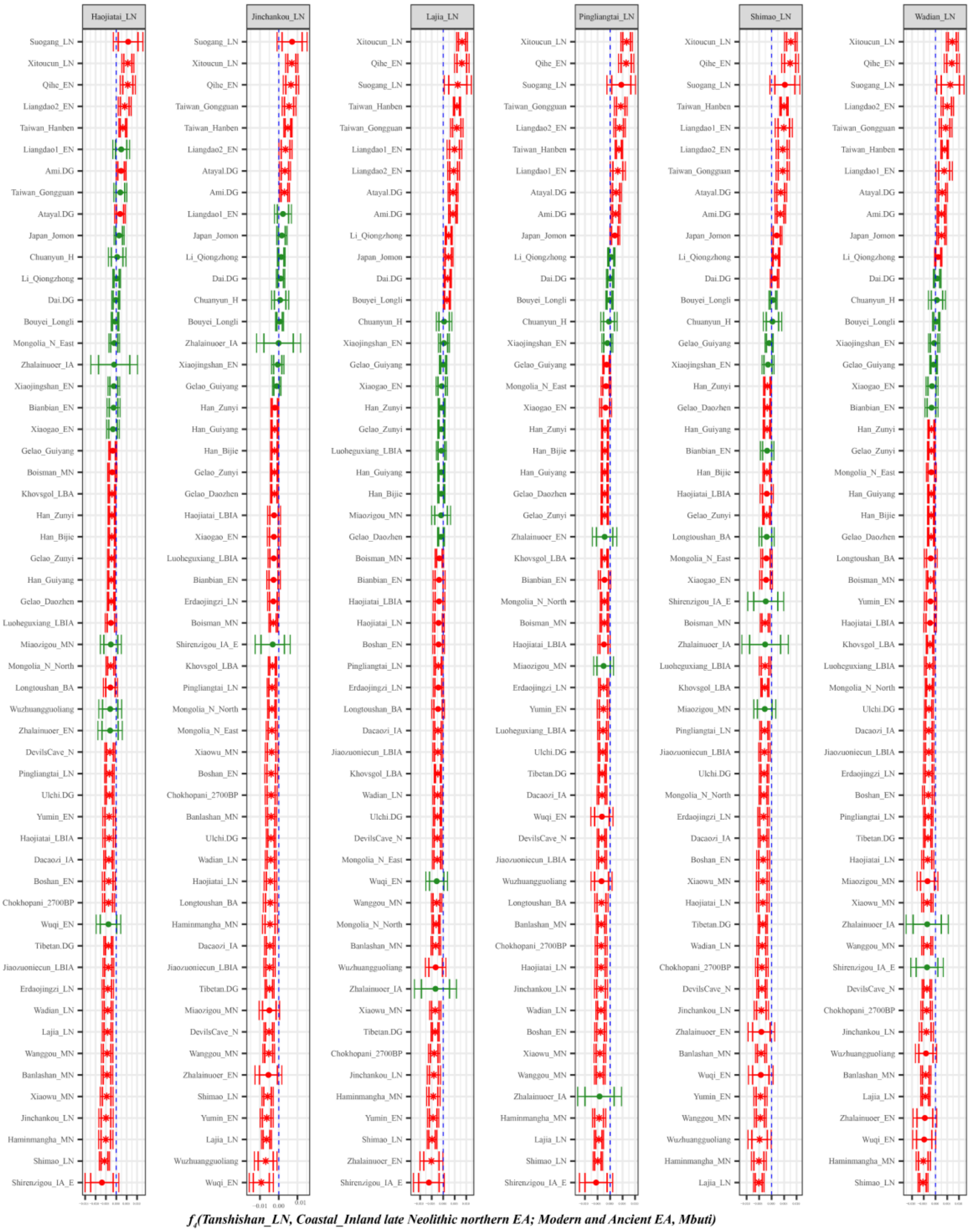

**Supplementary Fig. 66. Spatial difference of shared genetic drift of ancient populations from northern and southern East Asia assessed in the late Neolithic period via $f_4$*(Tanshishan_LN, Inland/Coastal late Neolithic northern East Asian; Studied inland TK/Sinitic and East Asian ancients, Mbuti)*.**

The dashed blue line indicates the zero $f_4$ value. The red asterisk indicates the absolute Z-score value larger than 3, the red point for absolute Z-score value ranging from two to three, and the green point for absolute Z-score value ranging from zero to two. The thick bar denotes two standard errors, and the thin bar for three standard errors. Figures were grouped via the second population in the $f_4$-statistics of Inland/Coastal late Neolithic northern East Asian. Significant negative values denoted the studied inland TK/Sinitic and East Asian ancients shared more alleles with Inland/Coastal late Neolithic northern East Asian, while significant positive $f_4$ values indicated the targeted populations shared more alleles with

Tanshishan_LN related ancestral populations.

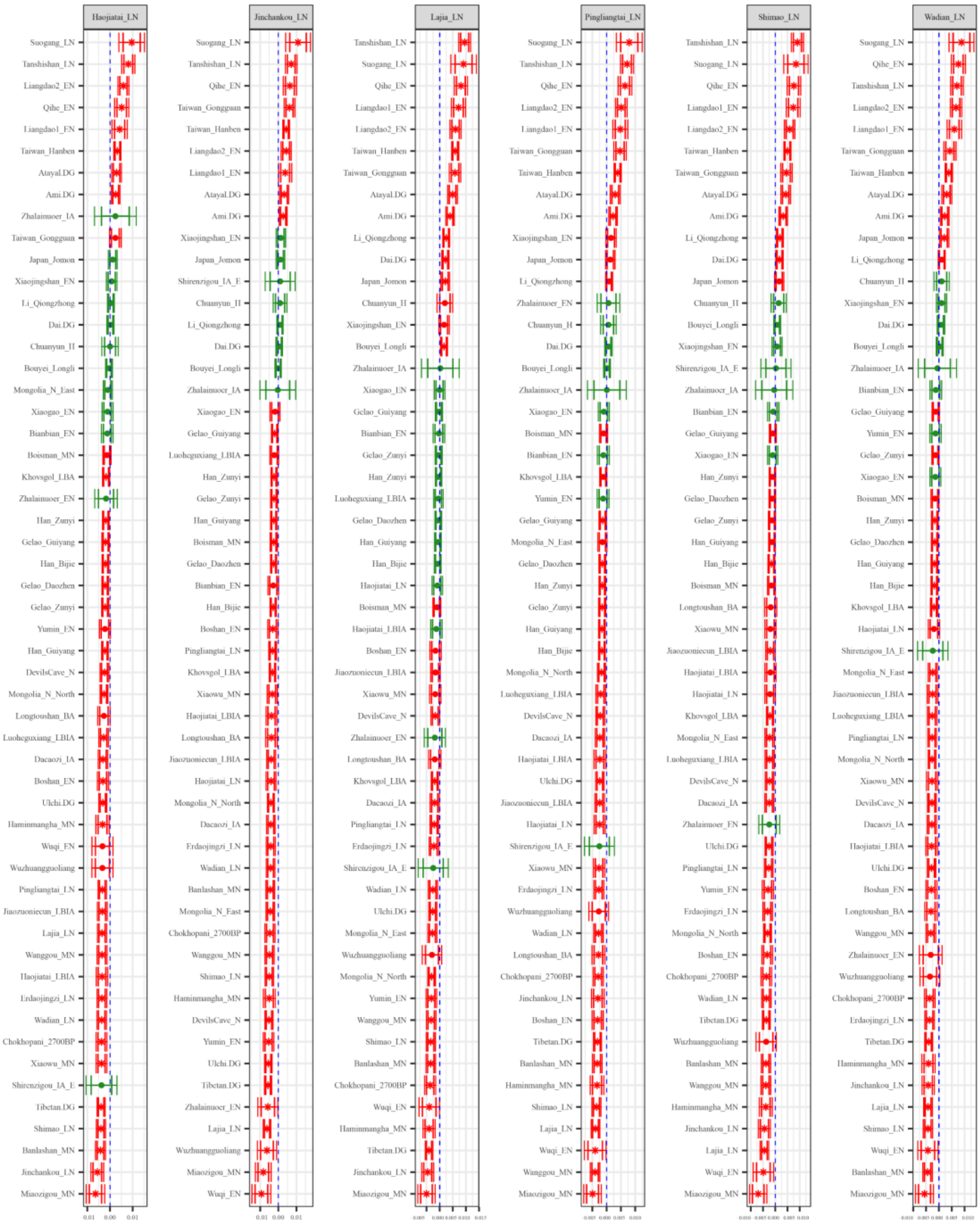

**Supplementary Fig. 67. Spatial difference of shared genetic drift of ancient populations from northern and southern East Asia assessed in the late Neolithic period via *$f_4$(Xitoucun_LN, Inland/Coastal late Neolithic northern East Asian; Studied inland TK/Sinitic and East Asian ancients, Mbuti)*.**

The dashed blue line indicates the zero $f_4$ value. The red asterisk indicates the absolute Z-score value larger than 3, the red point for absolute Z-score value ranging from two to three, and the green point for absolute Z-score value ranging from zero to two. The thick bar denotes two standard errors, and the thin bar for three standard errors. Figures were grouped via the second population in the $f_4$-statistics of Inland/Coastal late Neolithic northern East Asian. Significant negative values denoted the studied inland TK/Sinitic and East Asian ancients shared more alleles with Inland/Coastal late Neolithic northern East Asian, while significant positive $f_4$ values indicated the targeted populations shared more alleles with Xitoucun_LN related ancestral populations.
