## Supplementary Figures S21-40 for "New insights from the combined discrimination of modern/ancient genome-wide shared alleles and haplotypes: Differentiated demographic history reconstruction of Tai-Kadai and Sinitic people in South China"

### Contents of Supplementary Figures

Supplementary Fig. 24. The Z-scores of $f_4$*(Reference population1, Reference population2; Bouyei_Longli, Mbuti)* revealed the allele sharing between Longli Bouyei and reference populations. ..8
Supplementary Fig. 25. Results of genetic continuity and admixture between modern studied populations and Middle Yellow River Basin ancient populations. ..............................................................9
Supplementary Fig. 26. Results of genetic continuity and admixture between modern studied populations

and Upper Yellow River Basin ancient populations. 10
Supplementary Fig. 27. Results of genetic continuity and admixture between modern studied populations and Hunter-Gatherers from Siberia and Qinghai-Tibet Plateau high-altitude adaptive ancient populations. 11
Supplementary Fig. 28. Results of genetic continuity and admixture between modern studied populations and Taiwan ancient populations. 12
Supplementary Fig. 29A. Temporal changes of shared genetic drift of ancient populations from Yellow River Basin and other region of northern East Asia assessed via *$f_4$(Bianbian_EN, Inland/Coastal Neolithic/Bronze Age northern East Asian/Tibet Plateau/Siberia/Japan; Studied inland TK/Sinitic, Mbuti)*. 13
Supplementary Fig. 29B. Temporal changes of shared genetic drift of ancient populations from Yellow River Basin and other region of northern East Asia assessed via *$f_4$(Bianbian_EN, Inland/Coastal Neolithic/Bronze Age northern East Asian/Tibet Plateau/Siberia/Japan; Inland/Coastal Neolithic/Bronze Age southern East Asian, Mbuti)*. 14
Supplementary Fig. 30A. Temporal changes of shared genetic drift of ancient populations from Yellow River Basin and other region of northern East Asia assessed via *$f_4$(Boshan_EN, Inland/Coastal Neolithic/Bronze Age northern East Asian/Tibet Plateau/Siberia/Japan; Studied inland TK/Sinitic, Mbuti)*. 15
Supplementary Fig. 30B. Temporal changes of shared genetic drift of ancient populations from Yellow River Basin and other region of northern East Asia assessed via *$f_4$(Boshan_EN, Inland/Coastal Neolithic/Bronze Age northern East Asian/Tibet Plateau/Siberia/Japan; Inland/Coastal Neolithic/Bronze Age southern East Asian, Mbuti)*. 16
Supplementary Fig. 31A. Temporal changes of shared genetic drift of ancient populations from Yellow River Basin and other region of northern East Asia assessed via *$f_4$(Xiaogao_EN, Inland/Coastal Neolithic/Bronze Age northern East Asian/Tibet Plateau/Siberia/Japan; Studied inland TK/Sinitic, Mbuti)*. 17
Supplementary Fig. 31B. Temporal changes of shared genetic drift of ancient populations from Yellow River Basin and other region of northern East Asia assessed via *$f_4$(Xiaogao_EN, Inland/Coastal Neolithic/Bronze Age northern East Asian/Tibet Plateau/Siberia/Japan; Inland/Coastal Neolithic/Bronze Age southern East Asian, Mbuti)*. 18
Supplementary Fig. 32A. Temporal changes of shared genetic drift of ancient populations from Yellow River Basin and other region of northern East Asia assessed via *$f_4$(Yumin_EN, Inland/Coastal Neolithic/Bronze Age northern East Asian/Tibet Plateau/Siberia/Japan; Studied inland TK/Sinitic, Mbuti)*. 19
Supplementary Fig. 32B. Temporal changes of shared genetic drift of ancient populations from Yellow River Basin and other region of northern East Asia assessed via *$f_4$(Yumin_EN, Inland/Coastal Neolithic/Bronze Age northern East Asian/Tibet Plateau/Siberia/Japan; Inland/Coastal Neolithic/Bronze Age southern East Asian, Mbuti)*. 21
Supplementary Fig. 33A. Temporal changes of shared genetic drift of ancient populations from Yellow River Basin and other region of northern East Asia assessed via *$f_4$(Xiaojingshan_EN, Inland/Coastal Neolithic/Bronze Age northern East Asian/Tibet Plateau/Siberia/Japan; Studied inland TK/Sinitic, Mbuti)*. 22
Supplementary Fig. 33B. Temporal changes of shared genetic drift of ancient populations from Yellow River Basin and other region of northern East Asia assessed via *$f_4$(Xiaojingshan_EN, Inland/Coastal*

Supplementary Fig. 39A. Temporal changes of shared genetic drift of ancient populations from Yellow River Basin and other region of northern East Asia assessed via *$f_4$(Wadian_LN, Inland/Coastal*

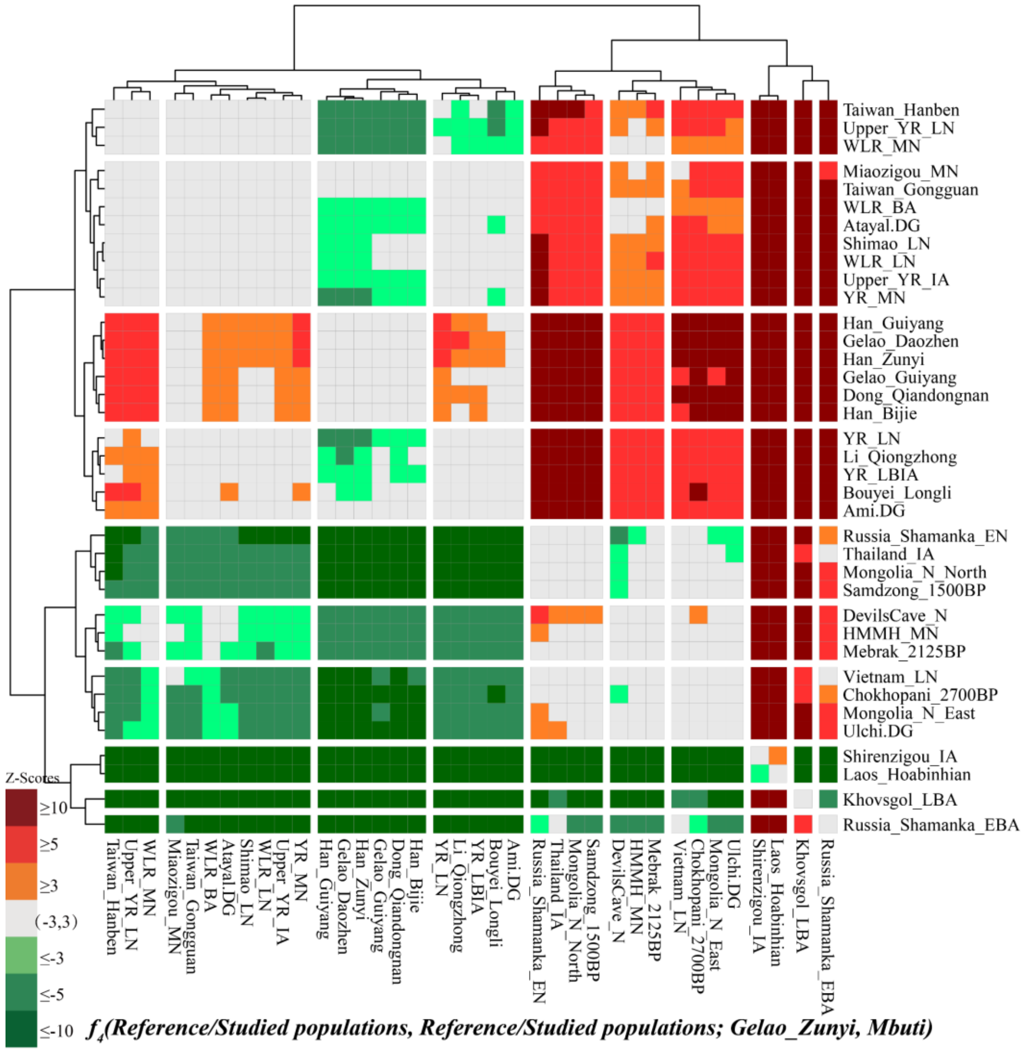

**Supplementary Fig. 21. The Z-scores of *f*₄*(Reference population1, Reference population2; Gelao_Zunyi, Mbuti)* revealed the allele sharing between Zunyi Gelao and reference populations.**

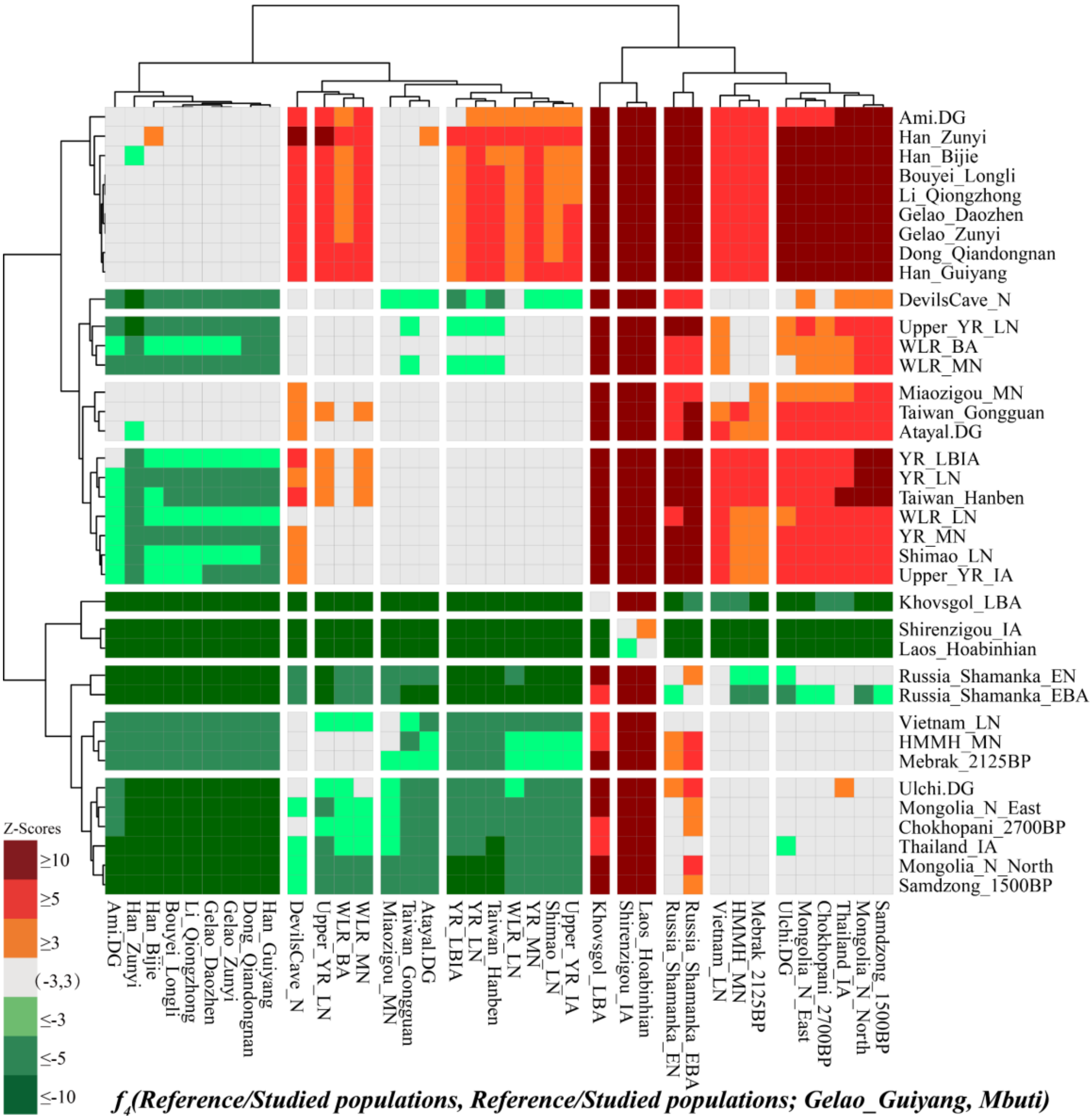

**Supplementary Fig. 22. The Z-scores of *$f_4$(Reference population1, Reference population2; Gelao_Guiyang, Mbuti)* revealed the allele sharing between Guiyang Gelao and reference populations.**

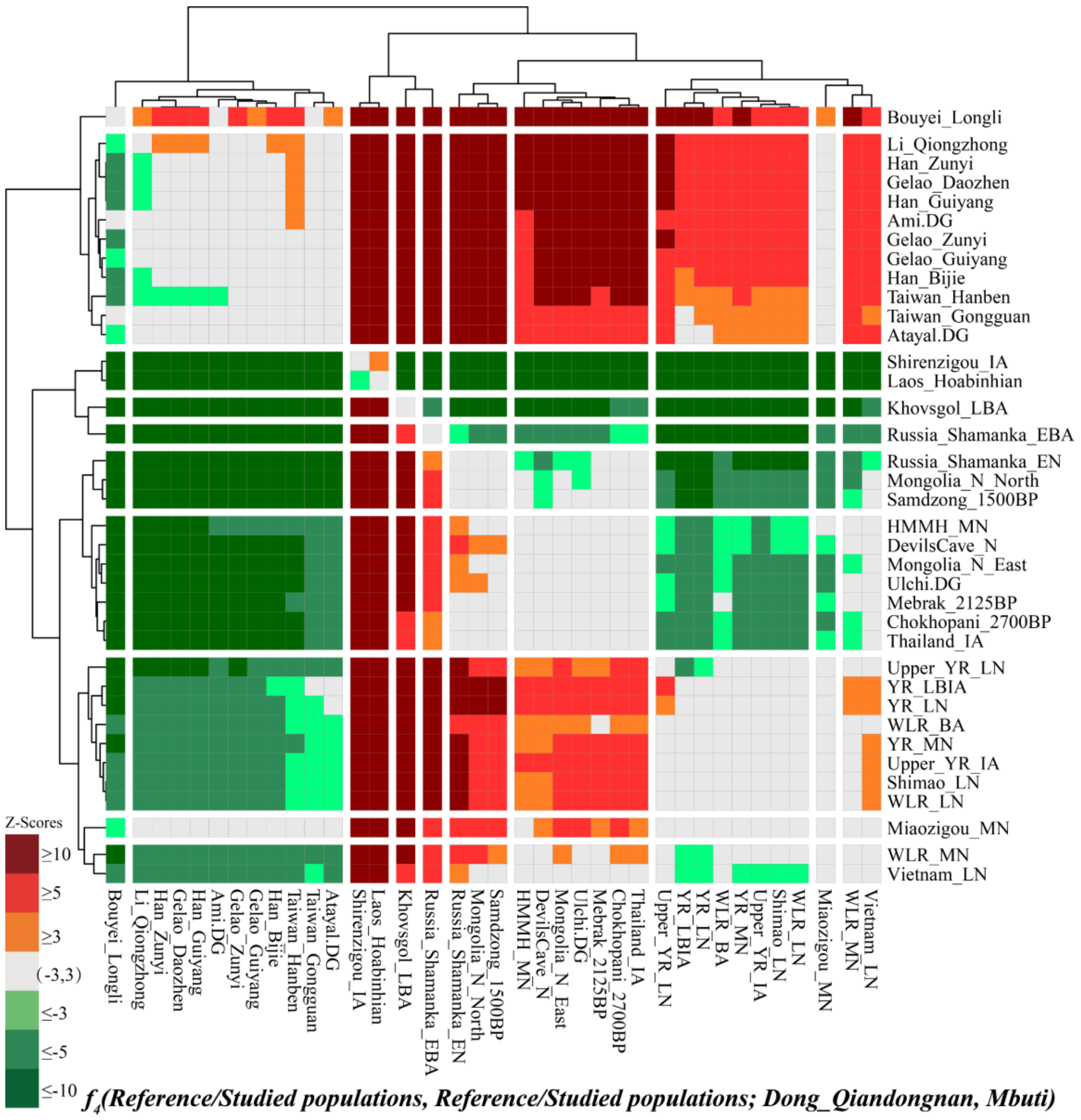

**Supplementary Fig. 23. The Z-scores of *$f_4$(Reference population1, Reference population2; Dong_Qiandongnan, Mbuti)* revealed the allele sharing between Qiandongnan Dong and reference populations.**

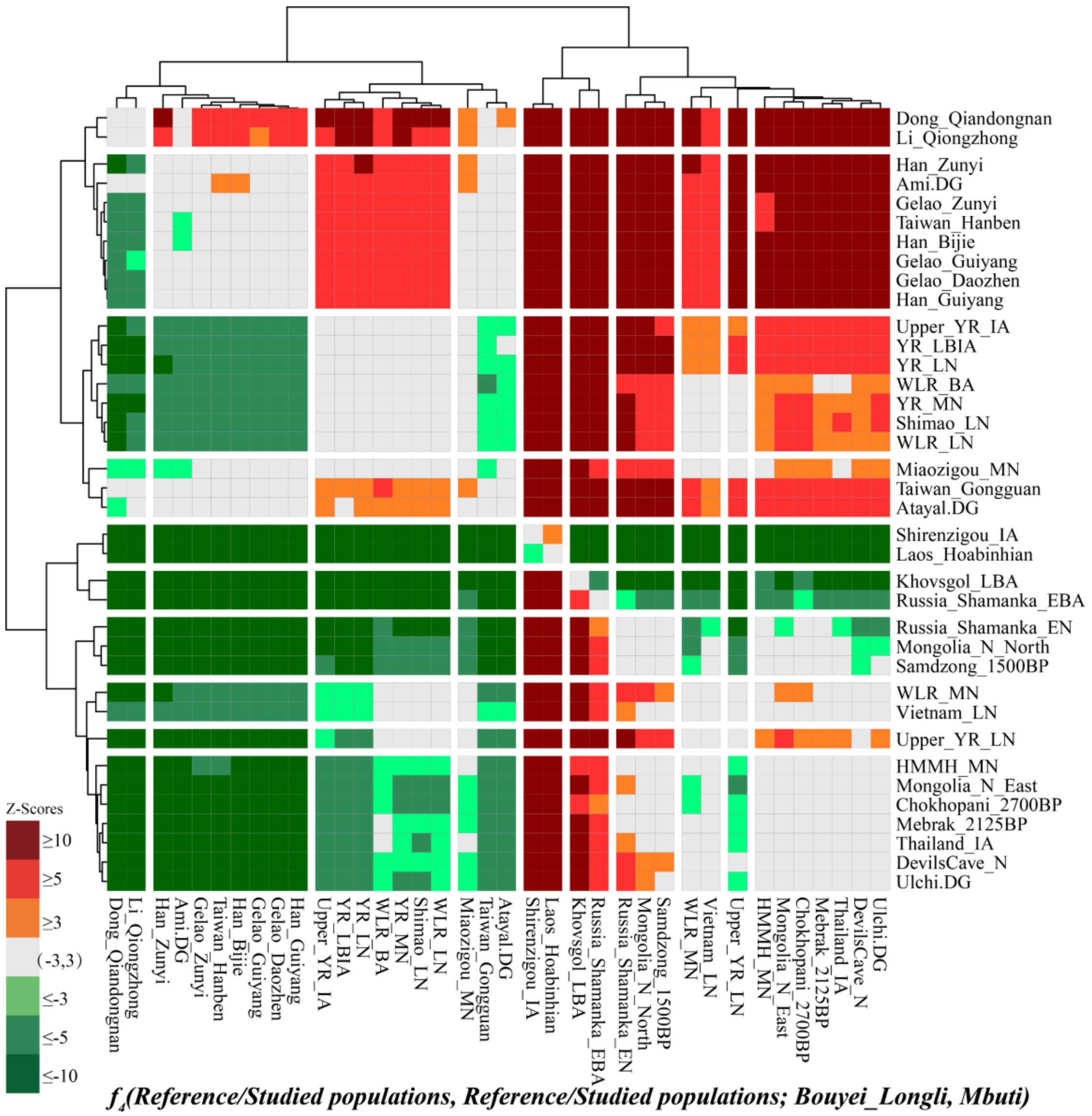

**Supplementary Fig. 24. The Z-scores of *f4(Reference population1, Reference population2; Bouyei_Longli, Mbuti)* revealed the allele sharing between Longli Bouyei and reference populations.**

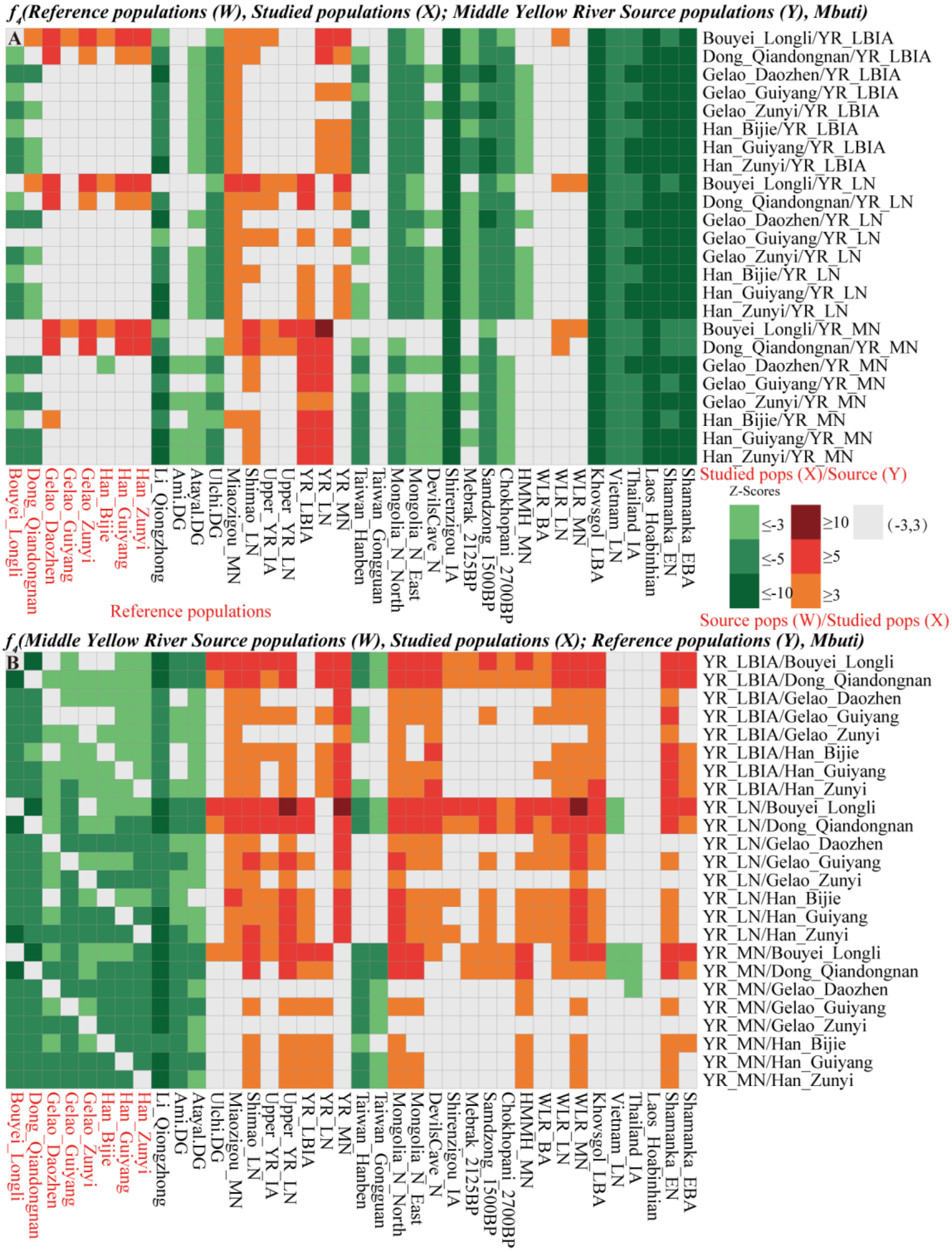

**Supplementary Fig. 25. Results of genetic continuity and admixture between modern studied populations and Middle Yellow River Basin ancient populations.**

(A) Genetic continuity between studied populations and Middle Yellow River source populations revealed by *$f_4$(Reference populations, Studied populations; Middle Yellow River Source populations, Mbuti)*; (B) The unbalanced allele sharing between reference populations with studied populations when compared to Middle Yellow River source populations revealed by *$f_4$(Middle Yellow River Source populations, Studied populations; Reference populations, Mbuti)*.

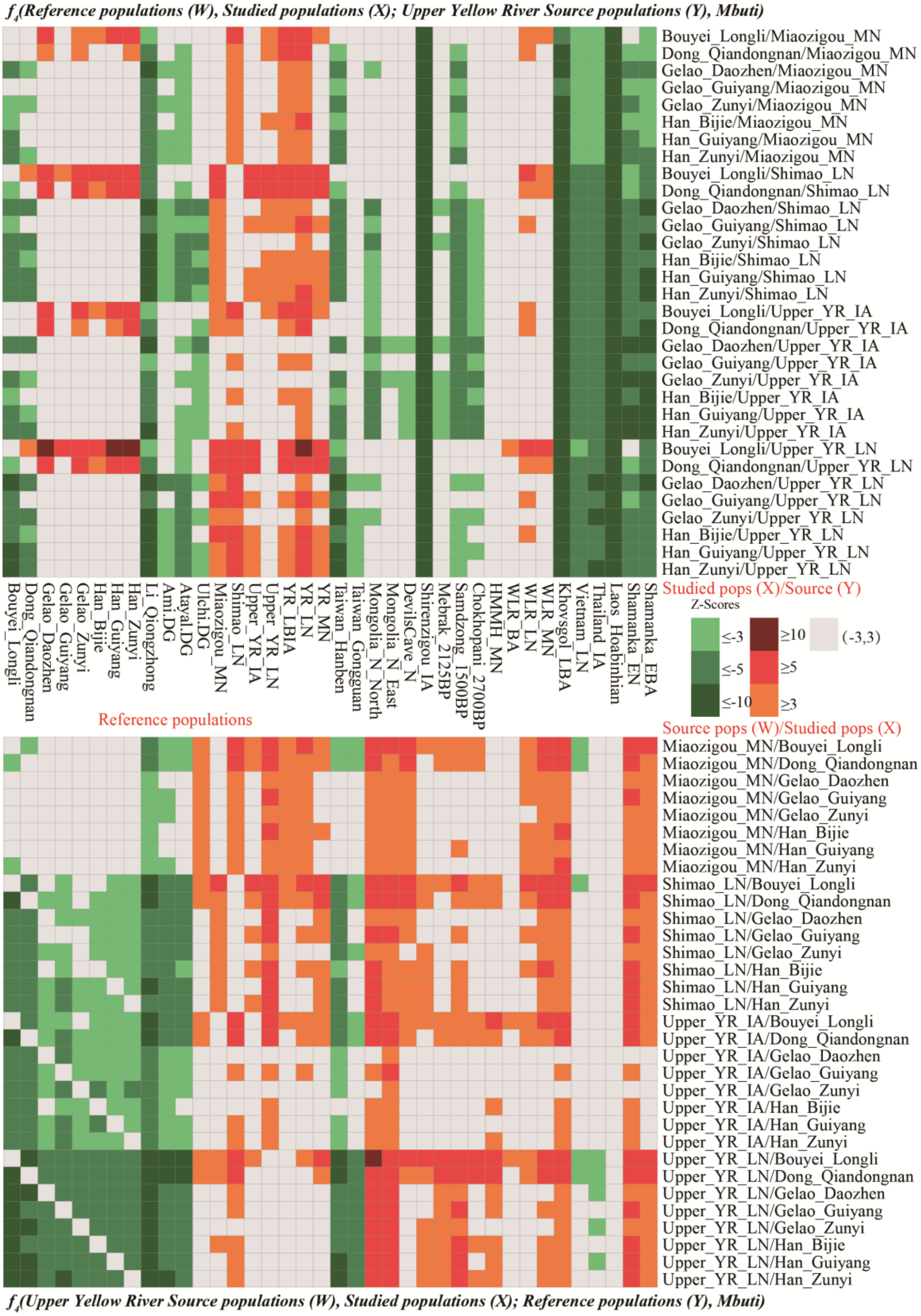

**Supplementary Fig. 26. Results of genetic continuity and admixture between modern studied populations and Upper Yellow River Basin ancient populations.**

(A) Genetic continuity between studied populations and Upper Yellow River source populations revealed by $f_4$*(Reference populations, Studied populations; Upper Yellow River Source populations, Mbuti)*; (B) The unbalanced allele sharing between reference populations with studied populations when compared to Upper Yellow River source populations revealed by $f_4$*(Upper Yellow River Source populations, Studied populations; Reference populations, Mbuti)*.

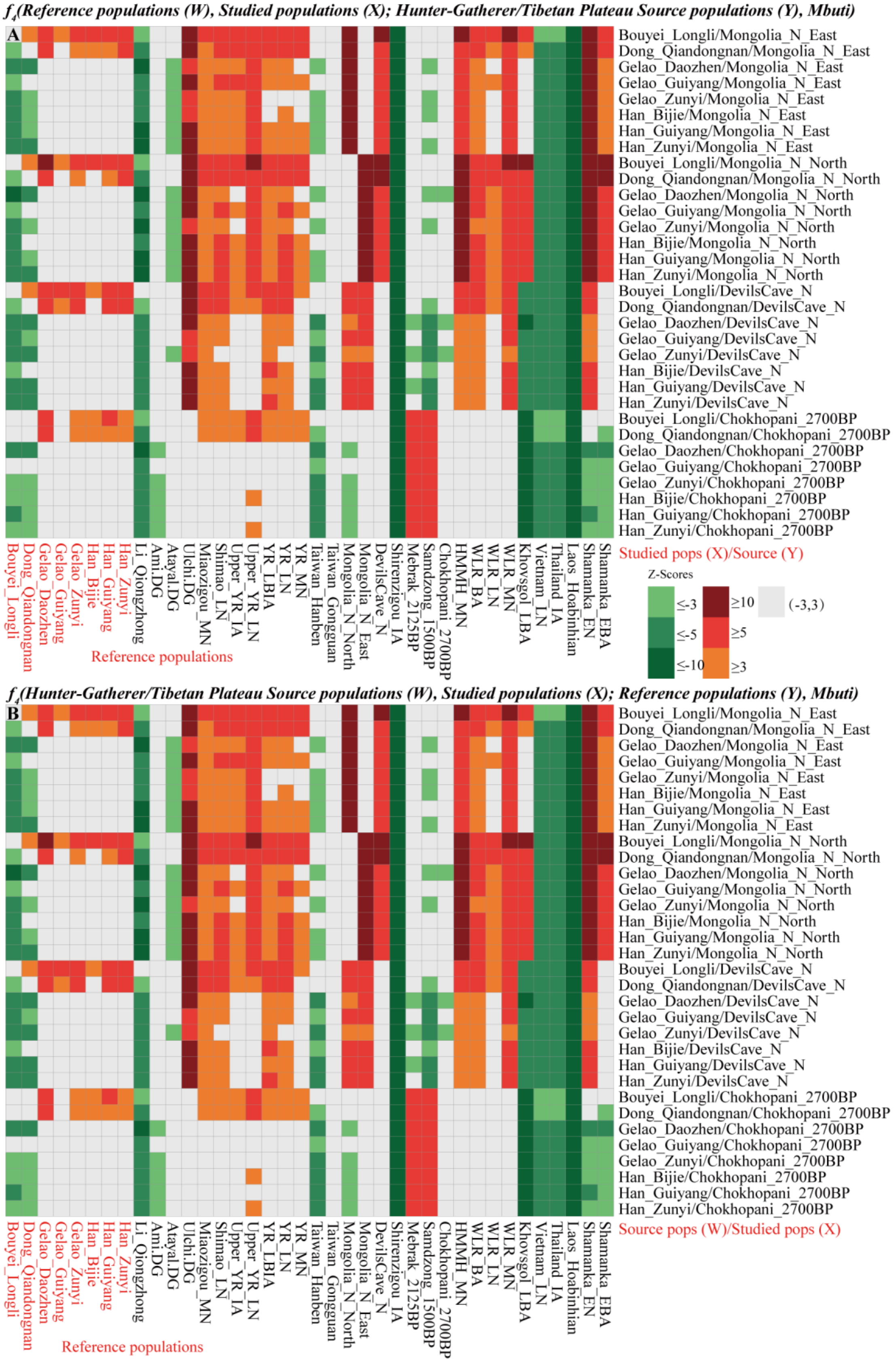

**Supplementary Fig. 27. Results of genetic continuity and admixture between modern studied populations and Hunter-Gatherers from Siberia and Qinghai-Tibet Plateau high-altitude adaptive ancient populations.**

(A) Genetic continuity between studied populations and Neolithic hunter-gatherers from Mongolia or

DevilsCave_N or Tibetan Plateau source populations revealed by $f_4$*(Reference populations, Studied populations; Hunter-Gatherer/Tibetan Plateau Source populations, Mbuti)*; (B) The unbalanced allele sharing between reference populations with studied populations when compared to Neolithic hunter-gatherers from Mongolia or DevilsCave_N or Tibetan Plateau source populations revealed by $f_4$*(Hunter-Gatherer/Tibetan Plateau Source populations, Studied populations; Reference populations, Mbuti)*.

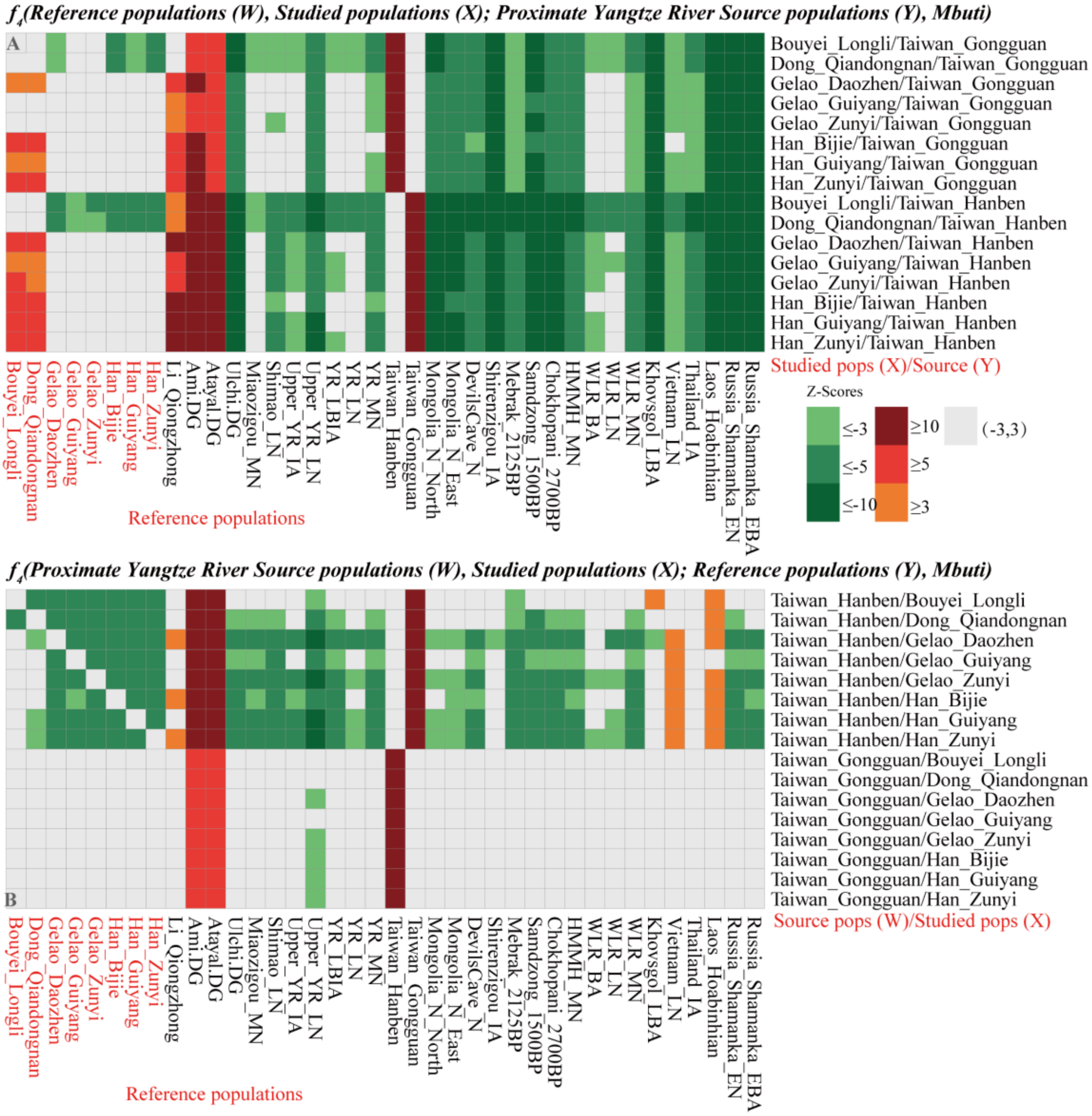

**Supplementary Fig. 28. Results of genetic continuity and admixture between modern studied populations and Taiwan ancient populations.**

(A) Genetic continuity between studied populations and proxy Yangtze River source populations revealed by $f_4$*(Reference populations, Studied populations; Proximate Yangtze River Source populations, Mbuti)*; (B) The unbalanced allele sharing between reference populations with studied populations when compared to proxy Yangtze River source populations revealed by $f_4$*(Proximate Yangtze River Source populations, Studied populations; Reference populations, Mbuti)*.

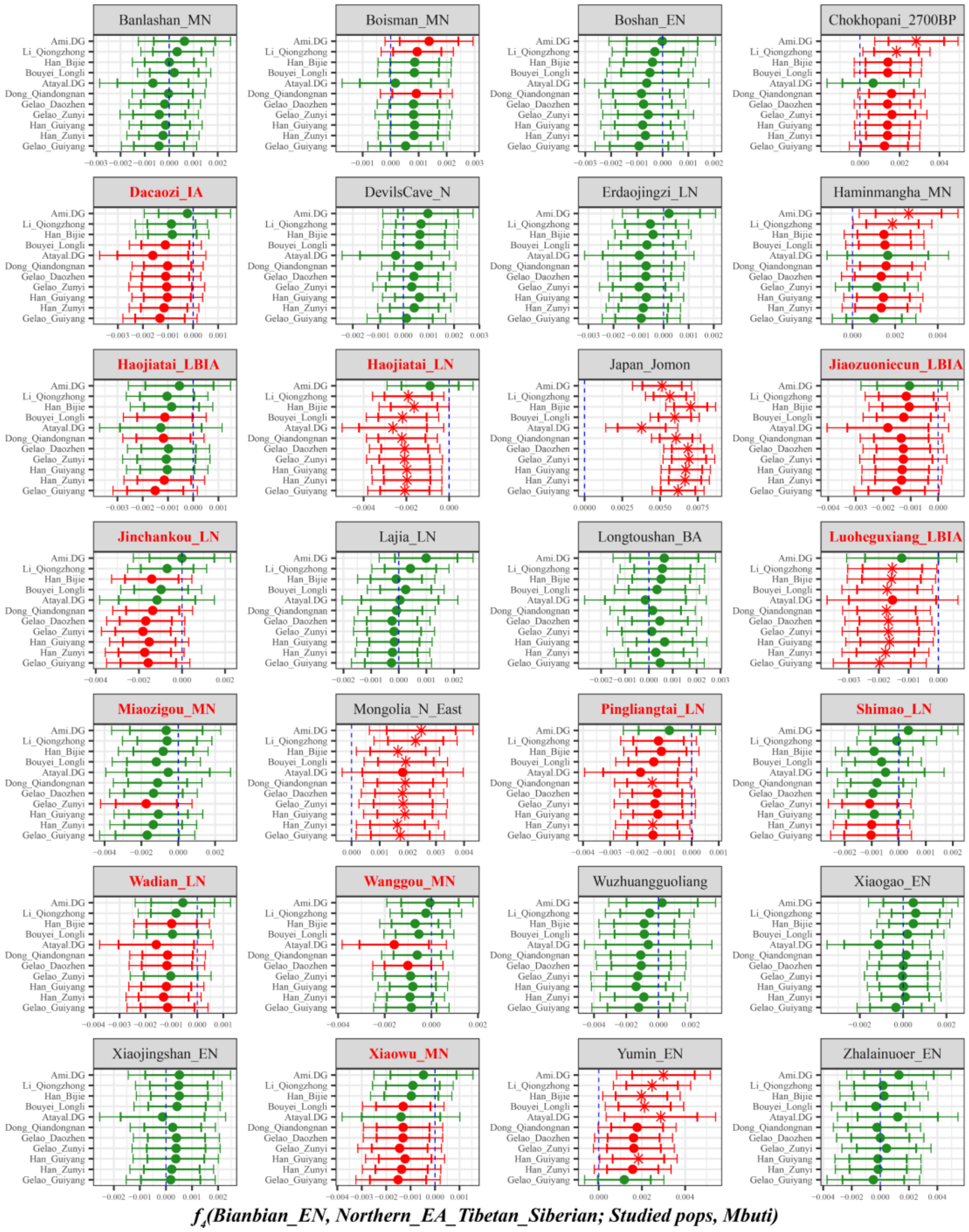

**Supplementary Fig. 29A. Temporal changes of shared genetic drift of ancient populations from Yellow River Basin and other region of northern East Asia assessed via *$f_4$(Bianbian_EN, Inland/Coastal Neolithic/Bronze Age northern East Asian/Tibet Plateau/Siberia/Japan; Studied inland TK/Sinitic, Mbuti)*.**

The dashed blue line indicates the zero $f_4$ value. The red asterisk indicates the absolute Z-score value larger than 3, the red point for absolute Z-score value ranging from two to three, and the green point for absolute Z-score value ranging from zero to two. The thick bar denotes two standard errors, and the thin bar for three standard errors. Figures were grouped via the second population in the $f_4$-statistics of Inland/Coastal Neolithic/Bronze Age northern East Asian/Tibet Plateau/Siberia/Japan, which was labeled as the red colour with some significant negative values. Significant negative values denoted the included second populations harboured more central/southern Sinitic or TK related ancestry. Significant positive $f_4$ values indicated the first population had more central/southern Sinitic or TK related ancestry. Here, compared with early Neolithic population of Bianbian, we observed middle Neolithic Xiaowu, Miaozigou and Wanggou people, late Neolithic Wadian, Shimao, Pingliangtai and Haojiatai people,

Bronze/Iron Age Dacaozi, Haojiaotai, Jiaozuoniecun and Luoheguxiang people had increased TK or southern modern Han related ancestry.

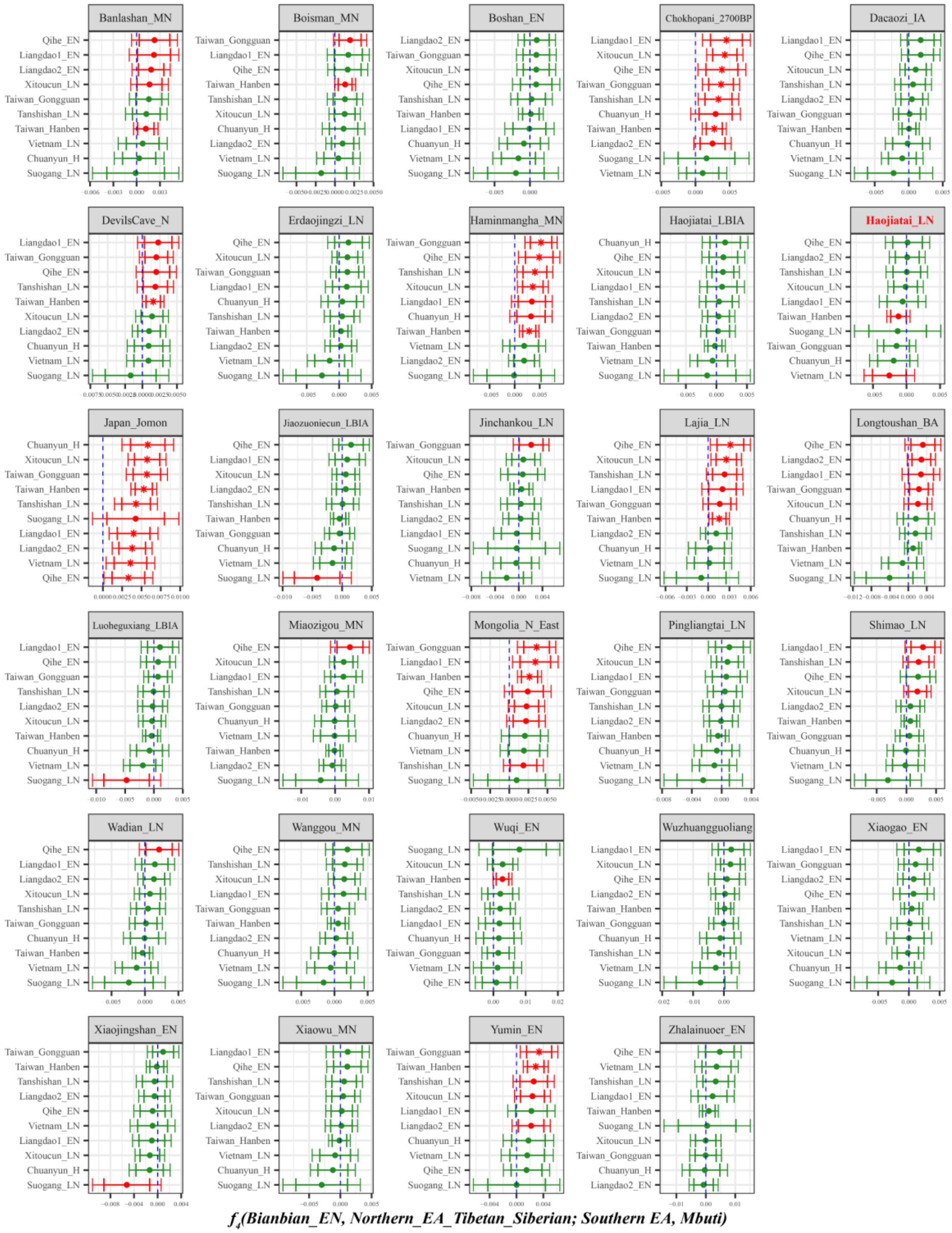

**Supplementary Fig. 29B. Temporal changes of shared genetic drift of ancient populations from Yellow River Basin and other region of northern East Asia assessed via $f_4$*(Bianbian_EN, Inland/Coastal Neolithic/Bronze Age northern East Asian/Tibet Plateau/Siberia/Japan; Inland/Coastal Neolithic/Bronze Age southern East Asian, Mbuti)*.**

The dashed blue line indicates the zero $f_4$ value. The red asterisk indicates the absolute Z-score value larger than 3, the red point for absolute Z-score value ranging from two to three, and the green point for absolute Z-score value ranging from zero to two. The thick bar denotes two standard errors, and the thin bar for three standard errors. Figures were grouped via the second population in the $f_4$-statistics of Inland/Coastal Neolithic/Bronze Age northern East Asian/Tibet Plateau/Siberia/Japan, which was labeled

as the red colour with some significant negative values. Significant negative values denoted the included second populations harboured more ancient southern East Asian related ancestry. Significant positive $f_4$ values indicated the first population had more ancient southern East Asian related ancestry.

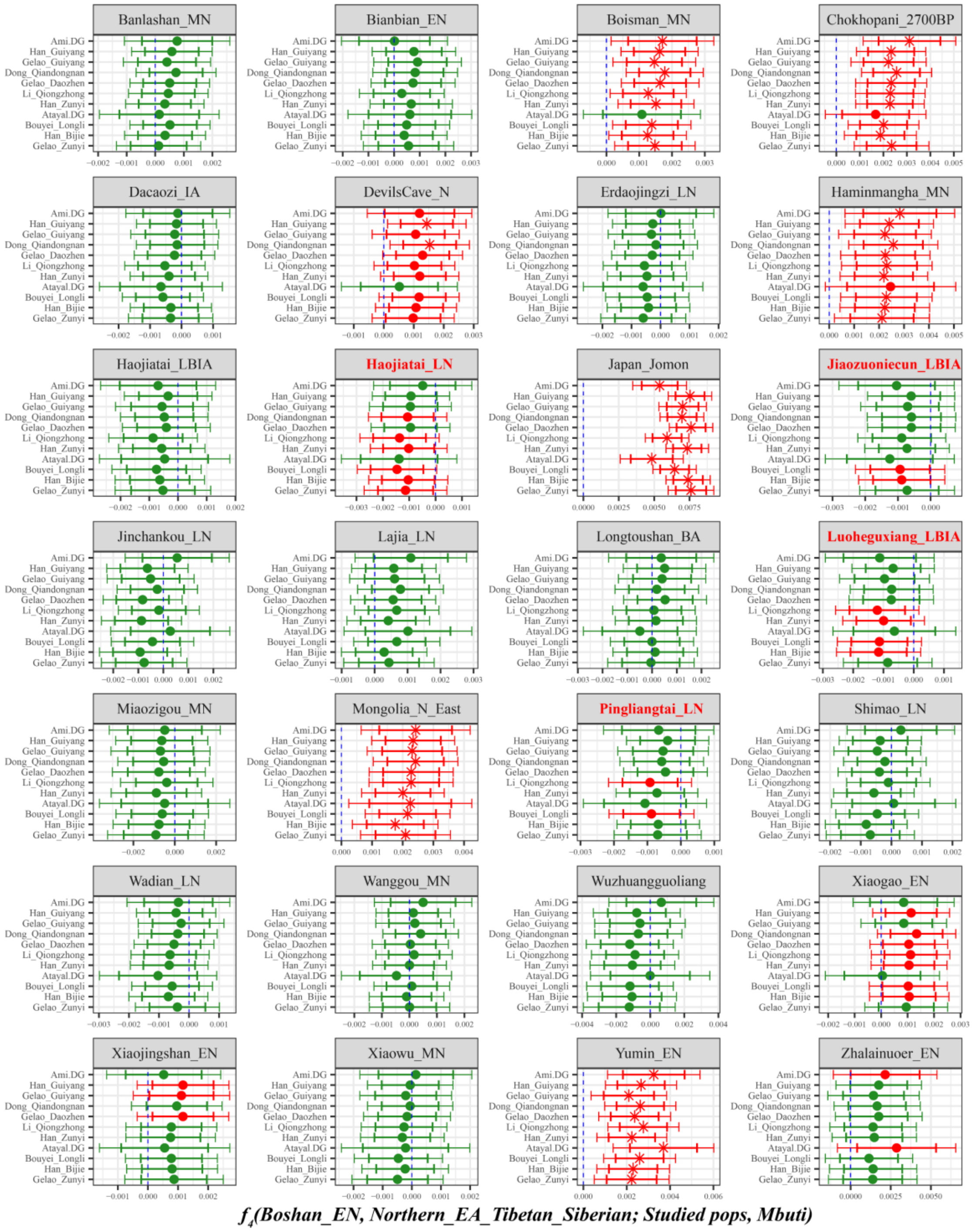

**Supplementary Fig. 30A. Temporal changes of shared genetic drift of ancient populations from Yellow River Basin and other region of northern East Asia assessed via *$f_4$(Boshan_EN, Inland/Coastal Neolithic/Bronze Age northern East Asian/Tibet Plateau/Siberia/Japan; Studied inland TK/Sinitic, Mbuti)*.**

The dashed blue line indicates the zero $f_4$ value. The red asterisk indicates the absolute Z-score value larger than 3, the red point for absolute Z-score value ranging from two to three, and the green point for absolute Z-score value ranging from zero to two. The thick bar denotes two standard errors, and the thin bar for three standard errors. Figures were grouped via the second population in the $f_4$-statistics of Inland/Coastal Neolithic/Bronze Age northern East Asian/Tibet Plateau/Siberia/Japan, which was labeled

as the red colour with some significant negative values. Significant negative values denoted the included second populations harboured more central/southern Sinitic or TK related ancestry. Significant positive $f_4$ values indicated the first population had more central/southern Sinitic or TK related ancestry.

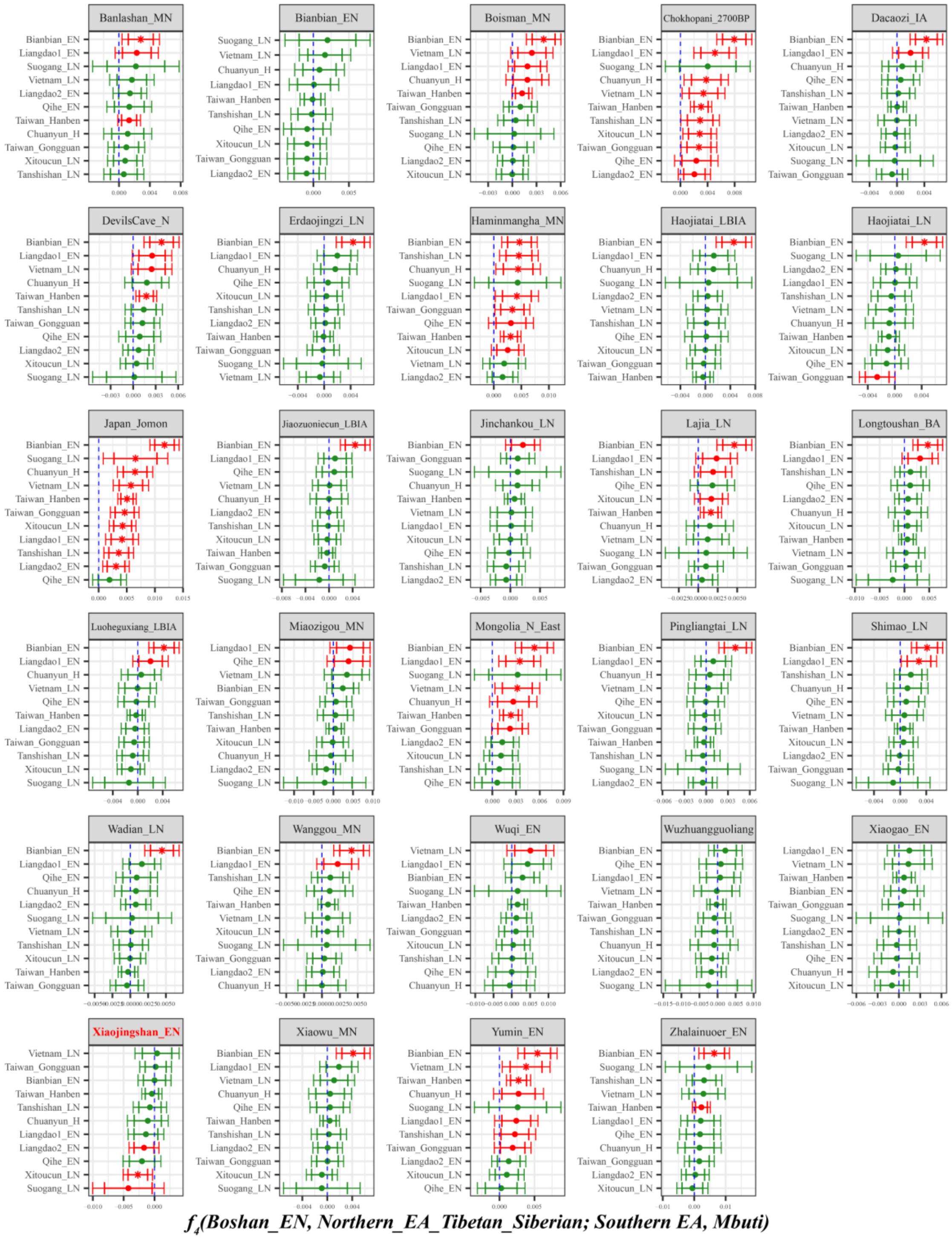

**Supplementary Fig. 30B. Temporal changes of shared genetic drift of ancient populations from Yellow River Basin and other region of northern East Asia assessed via *$f_4$(Boshan_EN, Inland/Coastal Neolithic/Bronze Age northern East Asian/Tibet Plateau/Siberia/Japan; Inland/Coastal Neolithic/Bronze Age southern East Asian, Mbuti)*.**

The dashed blue line indicates the zero $f_4$ value. The red asterisk indicates the absolute Z-score value larger than 3, the red point for absolute Z-score value ranging from two to three, and the green point for absolute Z-score value ranging from zero to two. The thick bar denotes two standard errors, and the thin bar for three standard errors. Figures were grouped via the second population in the $f_4$-statistics of

Inland/Coastal Neolithic/Bronze Age northern East Asian/Tibet Plateau/Siberia/Japan, which was labeled as the red colour with some significant negative values. Significant negative values denoted the included second populations harboured more ancient southern East Asian related ancestry. Significant positive $f_4$ values indicated the first population had more ancient southern East Asian related ancestry.

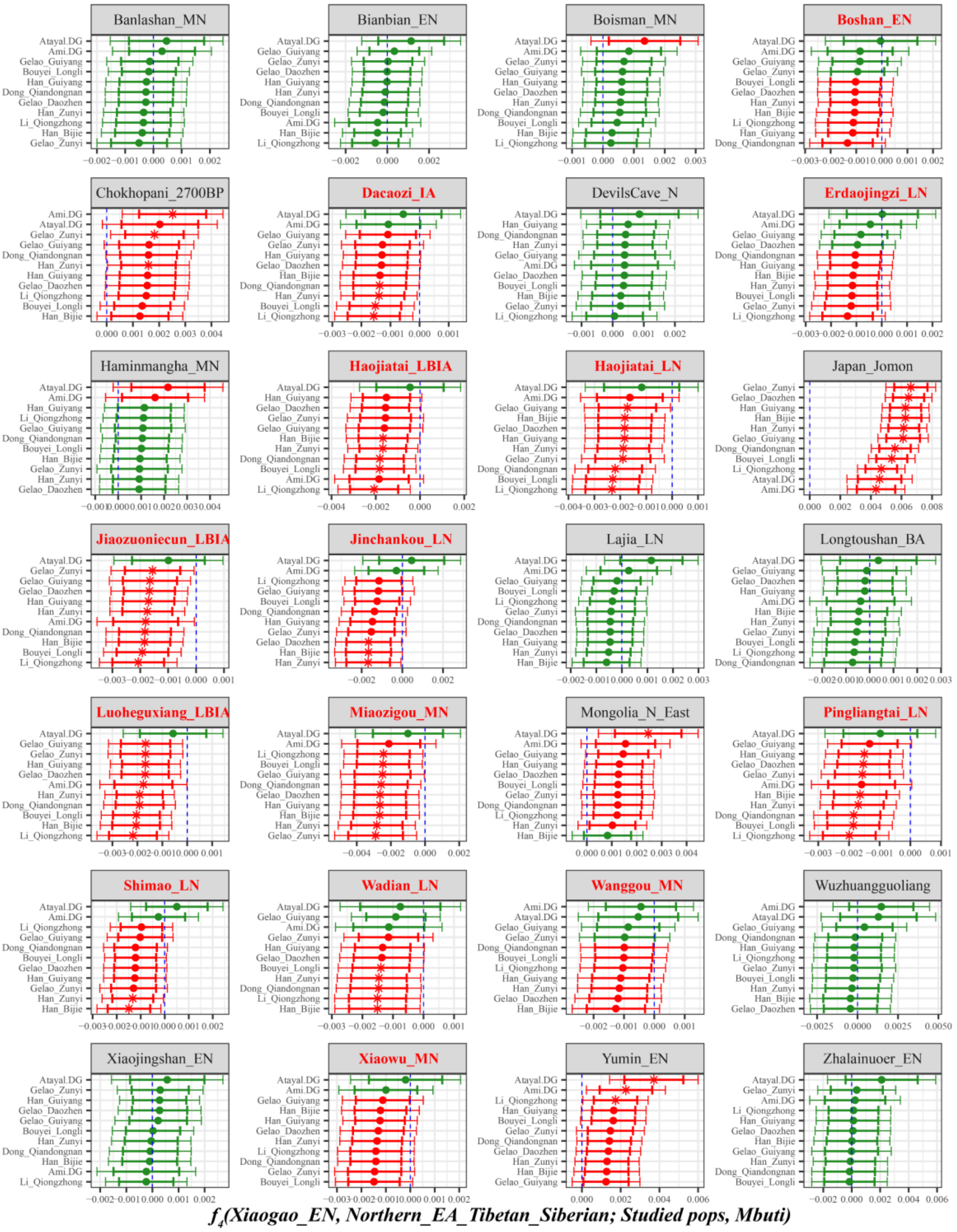

**Supplementary Fig. 31A. Temporal changes of shared genetic drift of ancient populations from Yellow River Basin and other region of northern East Asia assessed via *f4(Xiaogao_EN, Inland/Coastal Neolithic/Bronze Age northern East Asian/Tibet Plateau/Siberia/Japan; Studied inland TK/Sinitic, Mbuti)*.**

The dashed blue line indicates the zero $f_4$ value. The red asterisk indicates the absolute Z-score value larger than 3, the red point for absolute Z-score value ranging from two to three, and the green point for absolute Z-score value ranging from zero to two. The thick bar denotes two standard errors, and the thin bar for three standard errors. Figures were grouped via the second population in the $f_4$-statistics of Inland/Coastal Neolithic/Bronze Age northern East Asian/Tibet Plateau/Siberia/Japan, which was labeled as the red colour with some significant negative values. Significant negative values denoted the included second populations harboured more central/southern Sinitic or TK related ancestry. Significant positive $f_4$ values indicated the first population had more central/southern Sinitic or TK related ancestry.

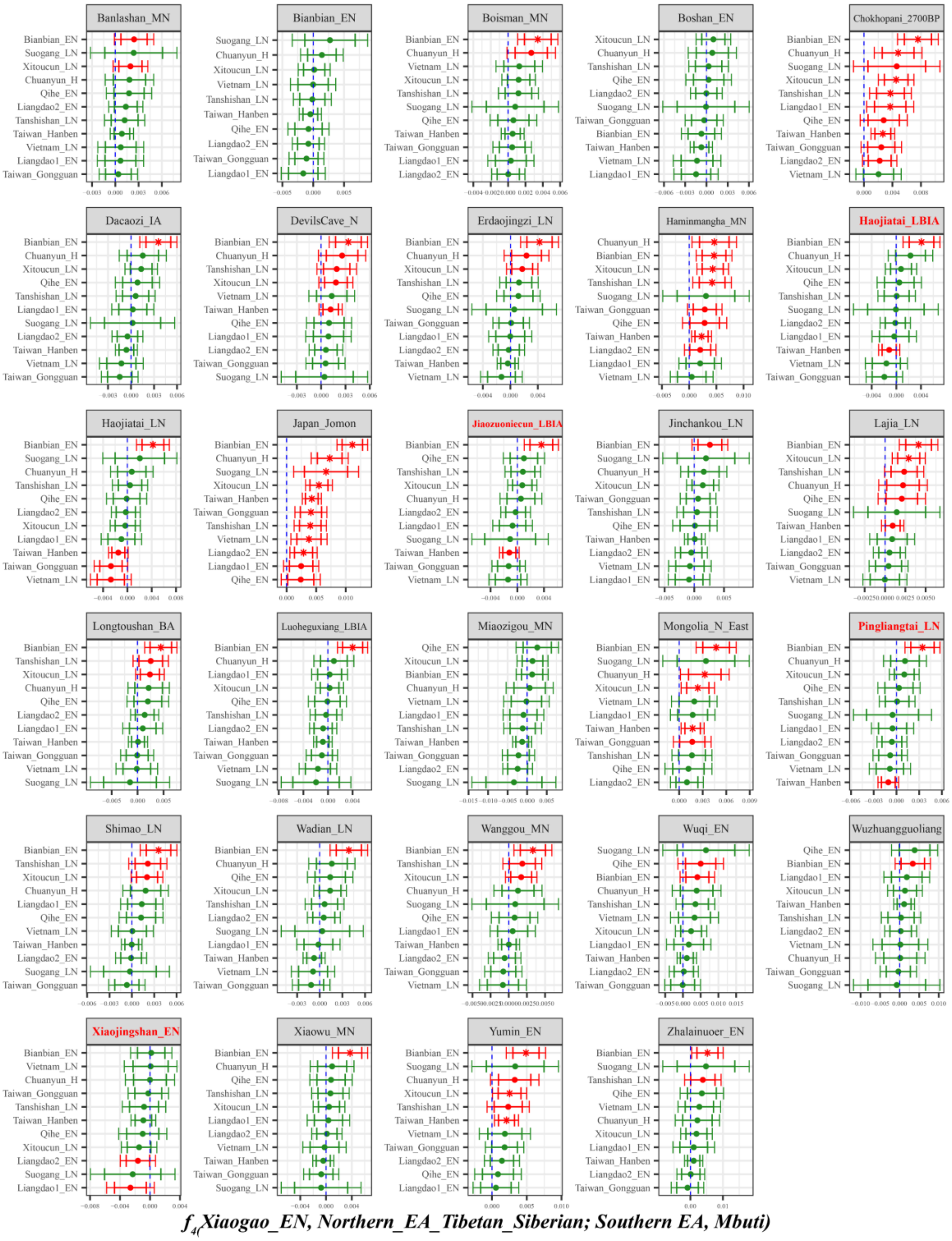

**Supplementary Fig. 31B. Temporal changes of shared genetic drift of ancient populations from Yellow River Basin and other region of northern East Asia assessed via *$f_4$(Xiaogao_EN, Inland/Coastal Neolithic/Bronze Age northern East Asian/Tibet Plateau/Siberia/Japan; Inland/Coastal Neolithic/Bronze Age southern East Asian, Mbuti)*.**

The dashed blue line indicates the zero $f_4$ value. The red asterisk indicates the absolute Z-score value larger than 3, the red point for absolute Z-score value ranging from two to three, and the green point for

absolute Z-score value ranging from zero to two. The thick bar denotes two standard errors, and the thin bar for three standard errors. Figures were grouped via the second population in the $f_4$-statistics of Inland/Coastal Neolithic/Bronze Age northern East Asian/Tibet Plateau/Siberia/Japan, which was labeled as the red colour with some significant negative values. Significant negative values denoted the included second populations harboured more ancient southern East Asian related ancestry. Significant positive $f_4$ values indicated the first population had more ancient southern East Asian related ancestry.

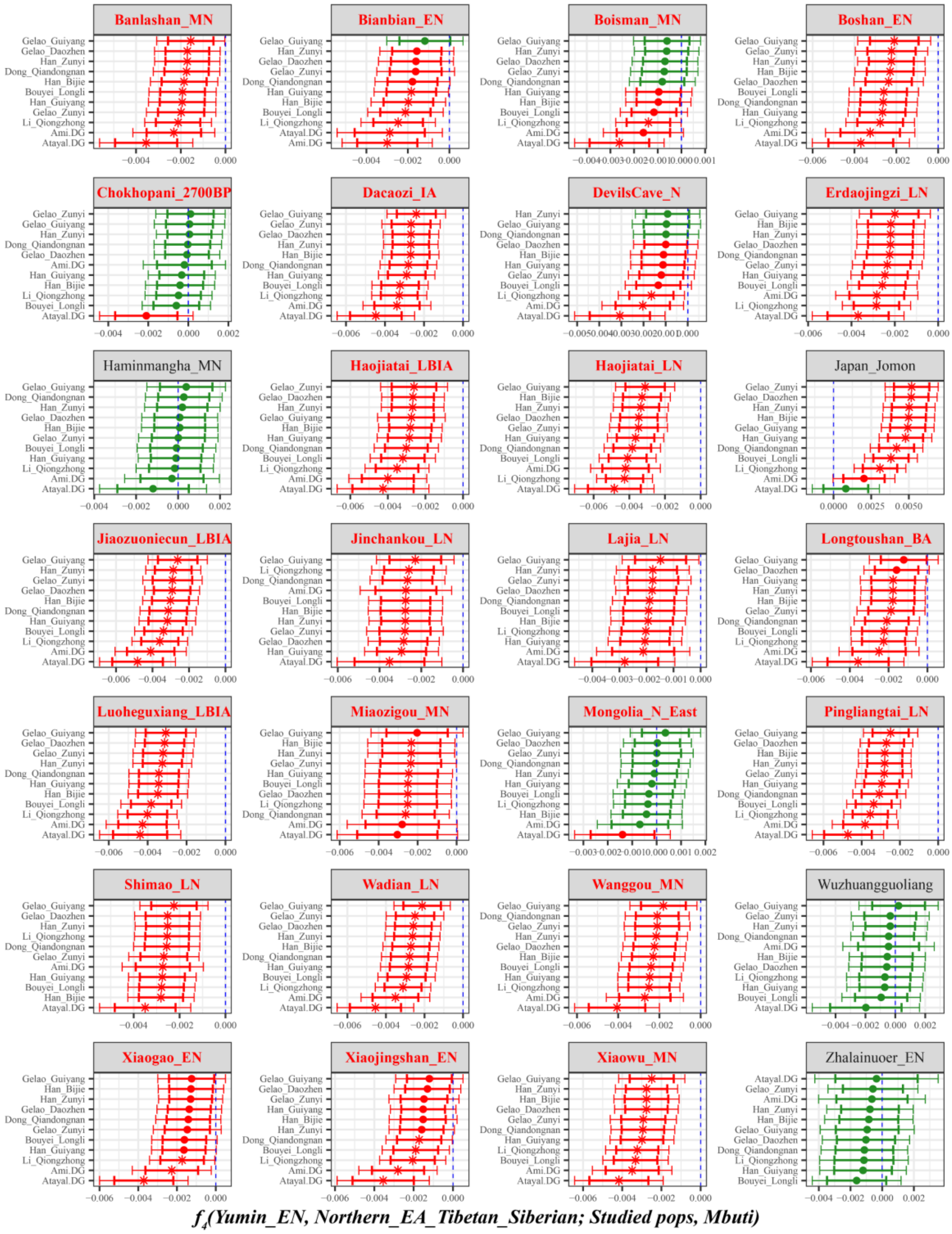

**Supplementary Fig. 32A. Temporal changes of shared genetic drift of ancient populations from Yellow River Basin and other region of northern East Asia assessed via *$f_4$(Yumin_EN, Inland/Coastal Neolithic/Bronze Age northern East Asian/Tibet Plateau/Siberia/Japan; Studied inland TK/Sinitic, Mbuti)*.**

The dashed blue line indicates the zero $f_4$ value. The red asterisk indicates the absolute Z-score value larger than 3, the red point for absolute Z-score value ranging from two to three, and the green point for absolute Z-score value ranging from zero to two. The thick bar denotes two standard errors, and the thin bar for three standard errors. Figures were grouped via the second population in the $f_4$-statistics of Inland/Coastal Neolithic/Bronze Age northern East Asian/Tibet Plateau/Siberia/Japan, which was labeled as the red colour with some significant negative values. Significant negative values denoted the included second populations harboured more central/southern Sinitic or TK related ancestry. Significant positive $f_4$ values indicated the first population had more central/southern Sinitic or TK related ancestry.

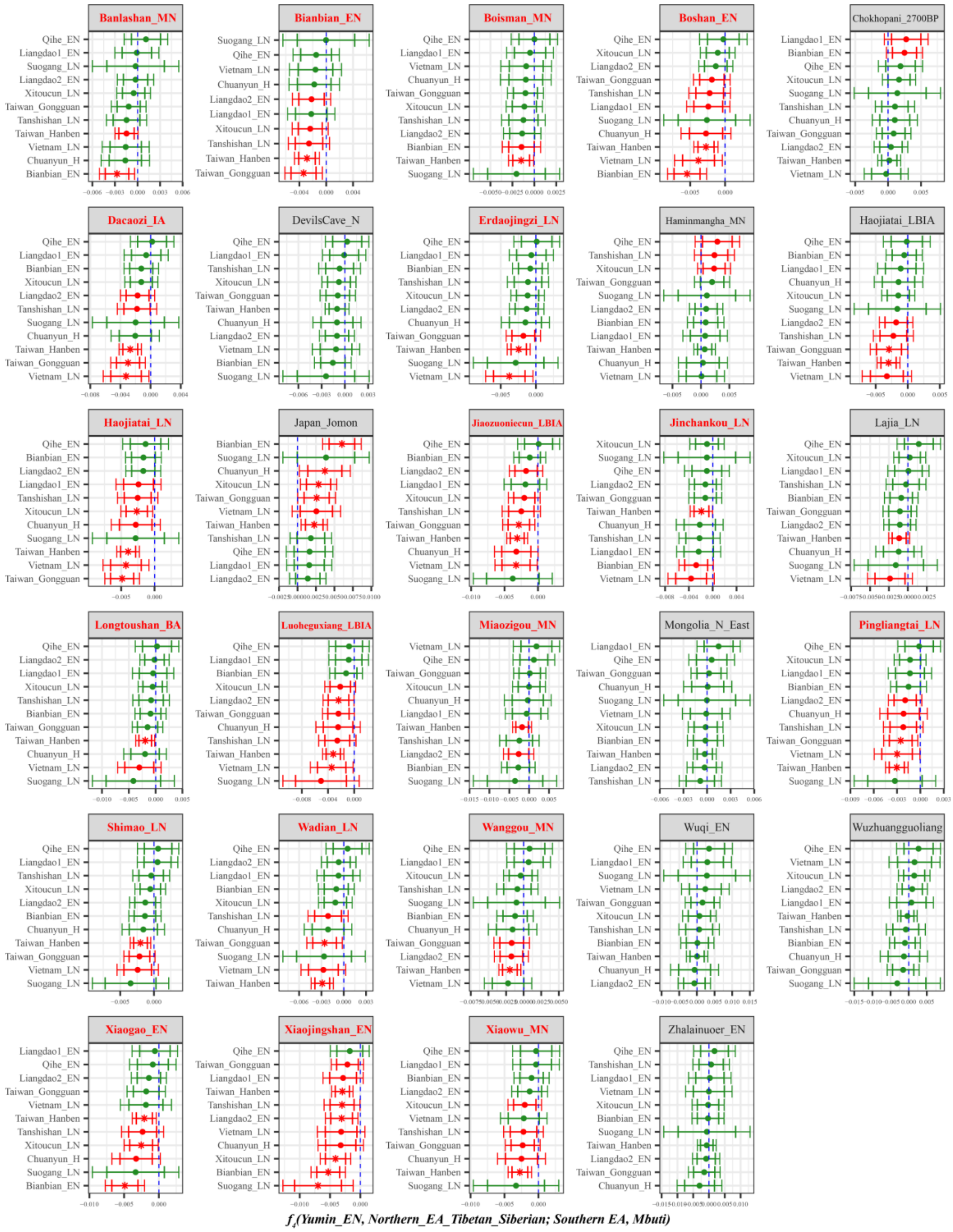

**Supplementary Fig. 32B. Temporal changes of shared genetic drift of ancient populations from Yellow River Basin and other region of northern East Asia assessed via *f4(Yumin_EN, Inland/Coastal Neolithic/Bronze Age northern East Asian/Tibet Plateau/Siberia/Japan; Inland/Coastal Neolithic/Bronze Age southern East Asian, Mbuti)*.**

The dashed blue line indicates the zero $f_4$ value. The red asterisk indicates the absolute Z-score value larger than 3, the red point for absolute Z-score value ranging from two to three, and the green point for absolute Z-score value ranging from zero to two. The thick bar denotes two standard errors, and the thin bar for three standard errors. Figures were grouped via the second population in the $f_4$-statistics of Inland/Coastal Neolithic/Bronze Age northern East Asian/Tibet Plateau/Siberia/Japan, which was labeled as the red colour with some significant negative values. Significant negative values denoted the included second populations harboured more ancient southern East Asian related ancestry. Significant positive $f_4$ values indicated the first population had more ancient southern East Asian related ancestry.

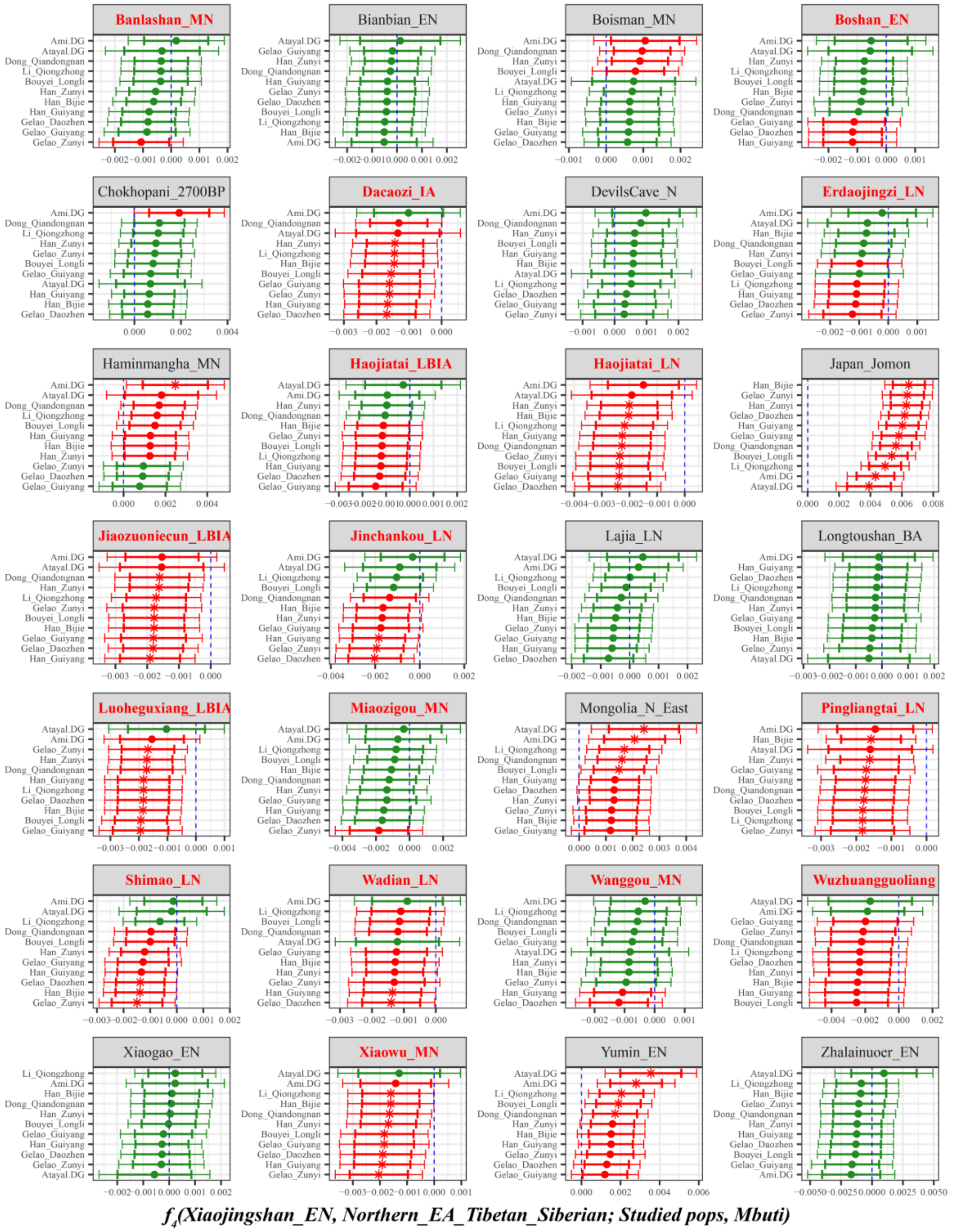

**Supplementary Fig. 33A. Temporal changes of shared genetic drift of ancient populations from Yellow River Basin and other region of northern East Asia assessed via *$f_4$(Xiaojingshan_EN, Inland/Coastal Neolithic/Bronze Age northern East Asian/Tibet Plateau/Siberia/Japan; Studied inland TK/Sinitic, Mbuti)*.**

The dashed blue line indicates the zero $f_4$ value. The red asterisk indicates the absolute Z-score value larger than 3, the red point for absolute Z-score value ranging from two to three, and the green point for absolute Z-score value ranging from zero to two. The thick bar denotes two standard errors, and the thin bar for three standard errors. Figures were grouped via the second population in the $f_4$-statistics of Inland/Coastal Neolithic/Bronze Age northern East Asian/Tibet Plateau/Siberia/Japan, which was labeled as the red colour with some significant negative values. Significant negative values denoted the included second populations harboured more central/southern Sinitic or TK related ancestry. Significant positive $f_4$ values indicated the first population had more central/southern Sinitic or TK related ancestry.

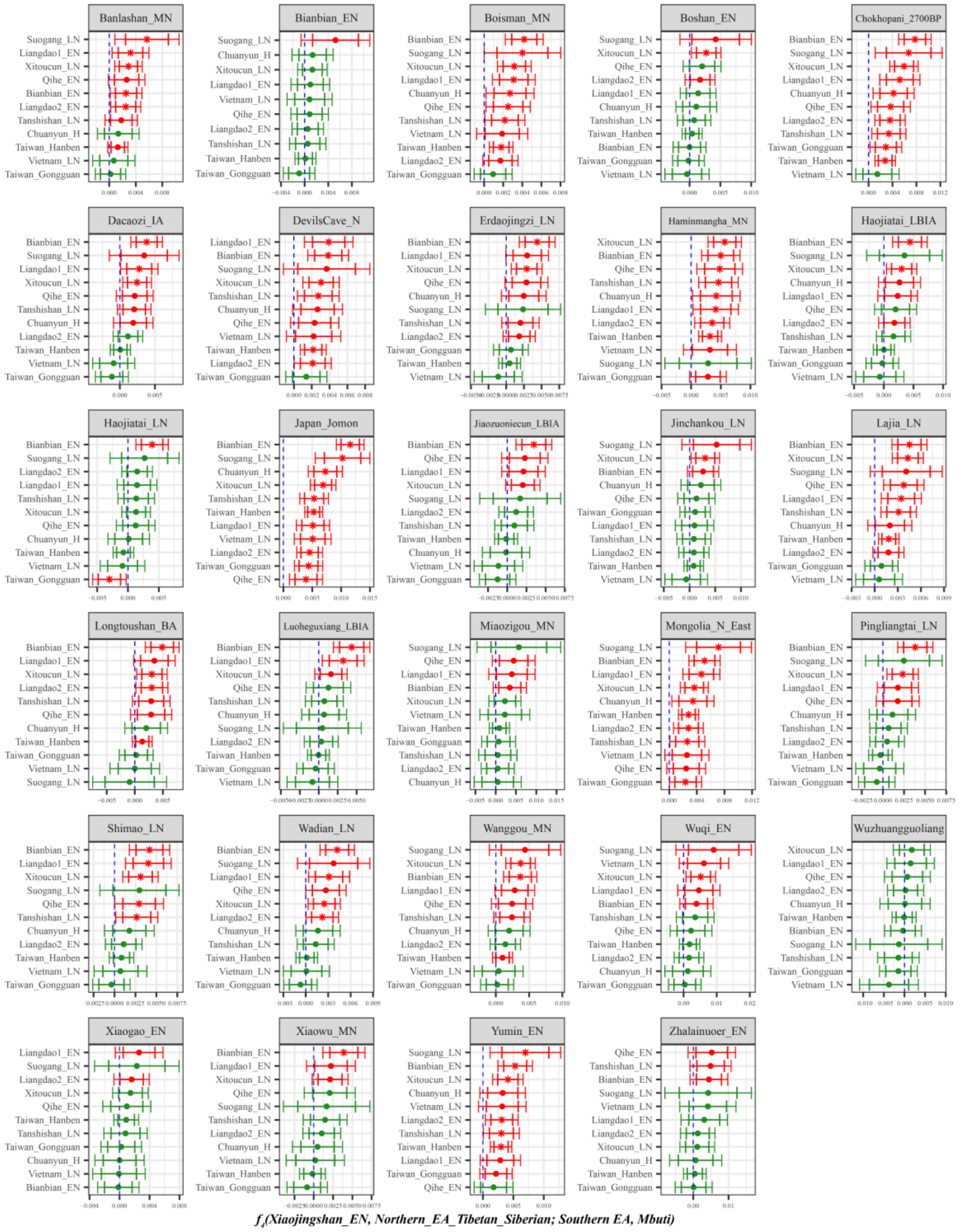

**Supplementary Fig. 33B. Temporal changes of shared genetic drift of ancient populations from Yellow River Basin and other region of northern East Asia assessed via $f_4$*(Xiaojingshan_EN, Inland/Coastal Neolithic/Bronze Age northern East Asian/Tibet Plateau/Siberia/Japan; Inland/Coastal Neolithic/Bronze Age southern East Asian, Mbuti)*.**

The dashed blue line indicates the zero $f_4$ value. The red asterisk indicates the absolute Z-score value larger than 3, the red point for absolute Z-score value ranging from two to three, and the green point for absolute Z-score value ranging from zero to two. The thick bar denotes two standard errors, and the thin bar for three standard errors. Figures were grouped via the second population in the $f_4$-statistics of Inland/Coastal Neolithic/Bronze Age northern East Asian/Tibet Plateau/Siberia/Japan, which was labeled as the red colour with some significant negative values. Significant negative values denoted the included second populations harboured more ancient southern East Asian related ancestry. Significant positive $f_4$ values indicated the first population had more ancient southern East Asian related ancestry.

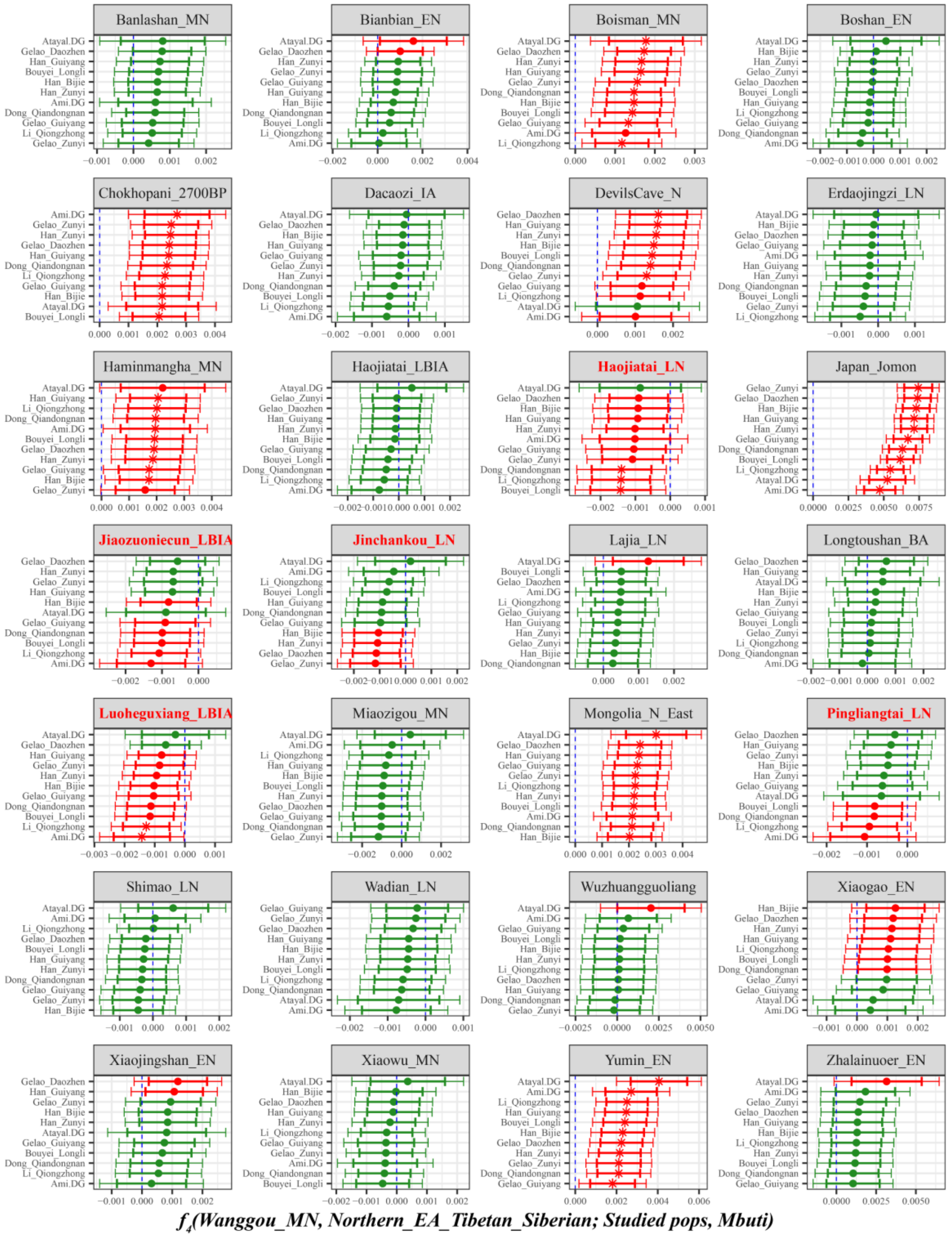

**Supplementary Fig. 34A. Temporal changes of shared genetic drift of ancient populations from Yellow River Basin and other region of northern East Asia assessed via *$f_4$(Wanggou_MN, Inland/Coastal Neolithic/Bronze Age northern East Asian/Tibet Plateau/Siberia/Japan; Studied inland TK/Sinitic, Mbuti)*.**

The dashed blue line indicates the zero $f_4$ value. The red asterisk indicates the absolute Z-score value larger than 3, the red point for absolute Z-score value ranging from two to three, and the green point for absolute Z-score value ranging from zero to two. The thick bar denotes two standard errors, and the thin bar for three standard errors. Figures were grouped via the second population in the $f_4$-statistics of Inland/Coastal Neolithic/Bronze Age northern East Asian/Tibet Plateau/Siberia/Japan, which was labeled as the red colour with some significant negative values. Significant negative values denoted the included second populations harboured more central/southern Sinitic or TK related ancestry. Significant positive

$f_4$ values indicated the first population had more central/southern Sinitic or TK related ancestry.

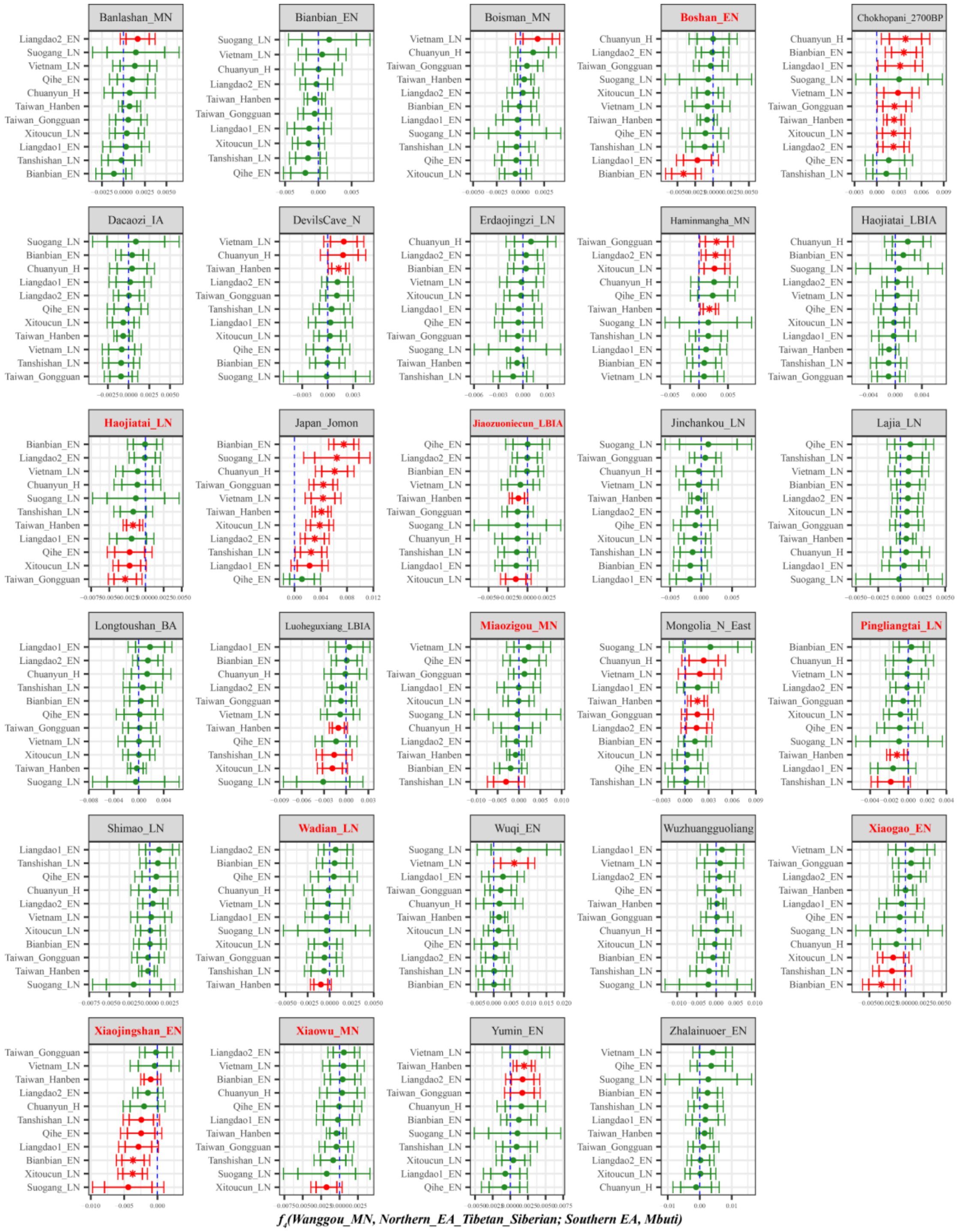

**Supplementary Fig. 34B. Temporal changes of shared genetic drift of ancient populations from Yellow River Basin and other region of northern East Asia assessed via $f_4$*(Wanggou_MN, Inland/Coastal Neolithic/Bronze Age northern East Asian/Tibet Plateau/Siberia/Japan; Inland/Coastal Neolithic/Bronze Age southern East Asian, Mbuti)*.**

The dashed blue line indicates the zero $f_4$ value. The red asterisk indicates the absolute Z-score value larger than 3, the red point for absolute Z-score value ranging from two to three, and the green point for absolute Z-score value ranging from zero to two. The thick bar denotes two standard errors, and the thin bar for three standard errors. Figures were grouped via the second population in the $f_4$-statistics of Inland/Coastal Neolithic/Bronze Age northern East Asian/Tibet Plateau/Siberia/Japan, which was labeled as the red colour with some significant negative values. Significant negative values denoted the included second populations harboured more ancient southern East Asian related ancestry. Significant positive $f_4$ values indicated the first population had more ancient southern East Asian related ancestry.

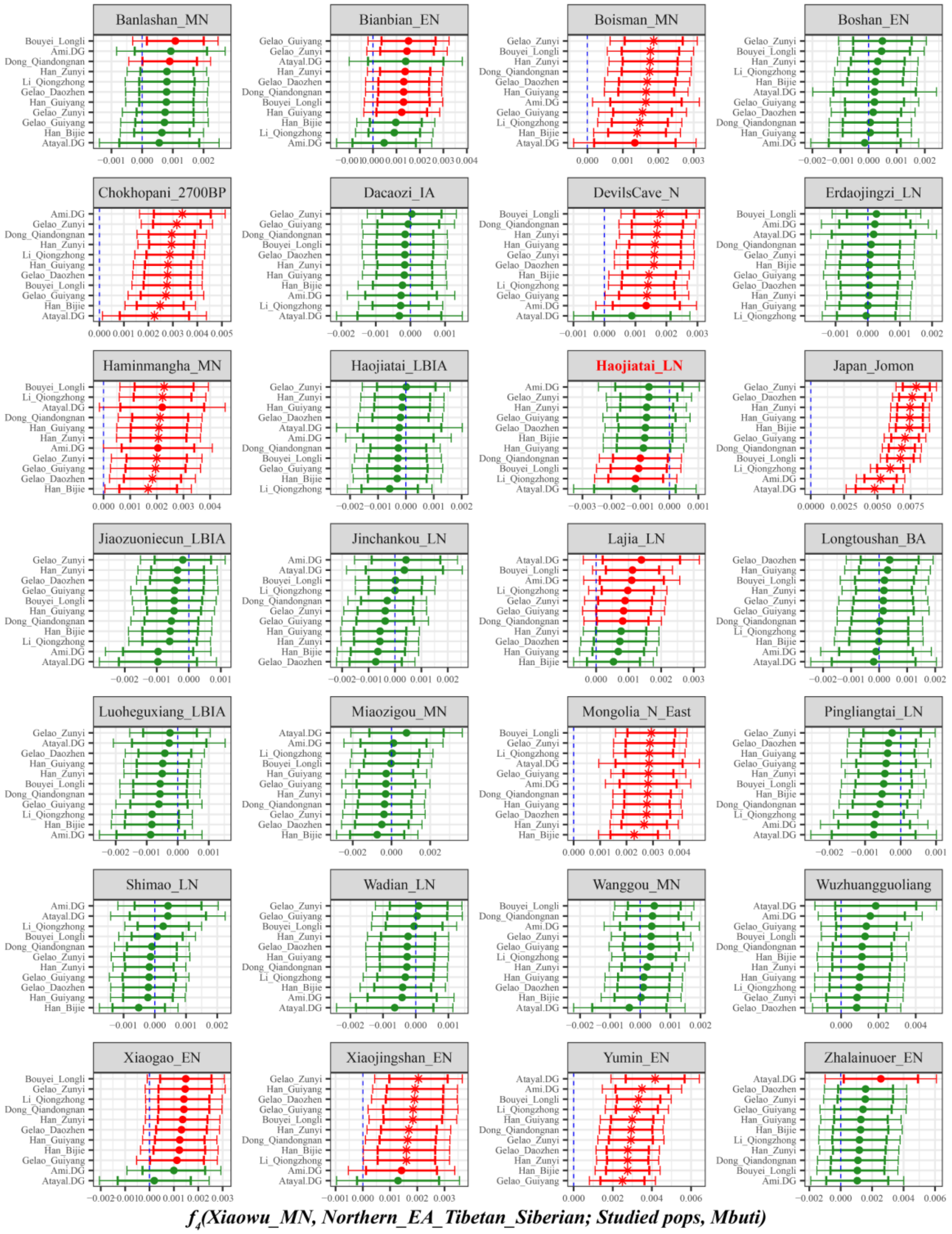

**Supplementary Fig. 35A. Temporal changes of shared genetic drift of ancient populations from Yellow River Basin and other region of northern East Asia assessed via *$f_4$(Xiaowu_MN, Inland/Coastal Neolithic/Bronze Age northern East Asian/Tibet Plateau/Siberia/Japan; Studied inland TK/Sinitic, Mbuti)*.**

The dashed blue line indicates the zero $f_4$ value. The red asterisk indicates the absolute Z-score value larger than 3, the red point for absolute Z-score value ranging from two to three, and the green point for absolute Z-score value ranging from zero to two. The thick bar denotes two standard errors, and the thin bar for three standard errors. Figures were grouped via the second population in the $f_4$-statistics of Inland/Coastal Neolithic/Bronze Age northern East Asian/Tibet Plateau/Siberia/Japan, which was labeled as the red colour with some significant negative values. Significant negative values denoted the included second populations harboured more central/southern Sinitic or TK related ancestry. Significant positive

$f_4$ values indicated the first population had more central/southern Sinitic or TK related ancestry.

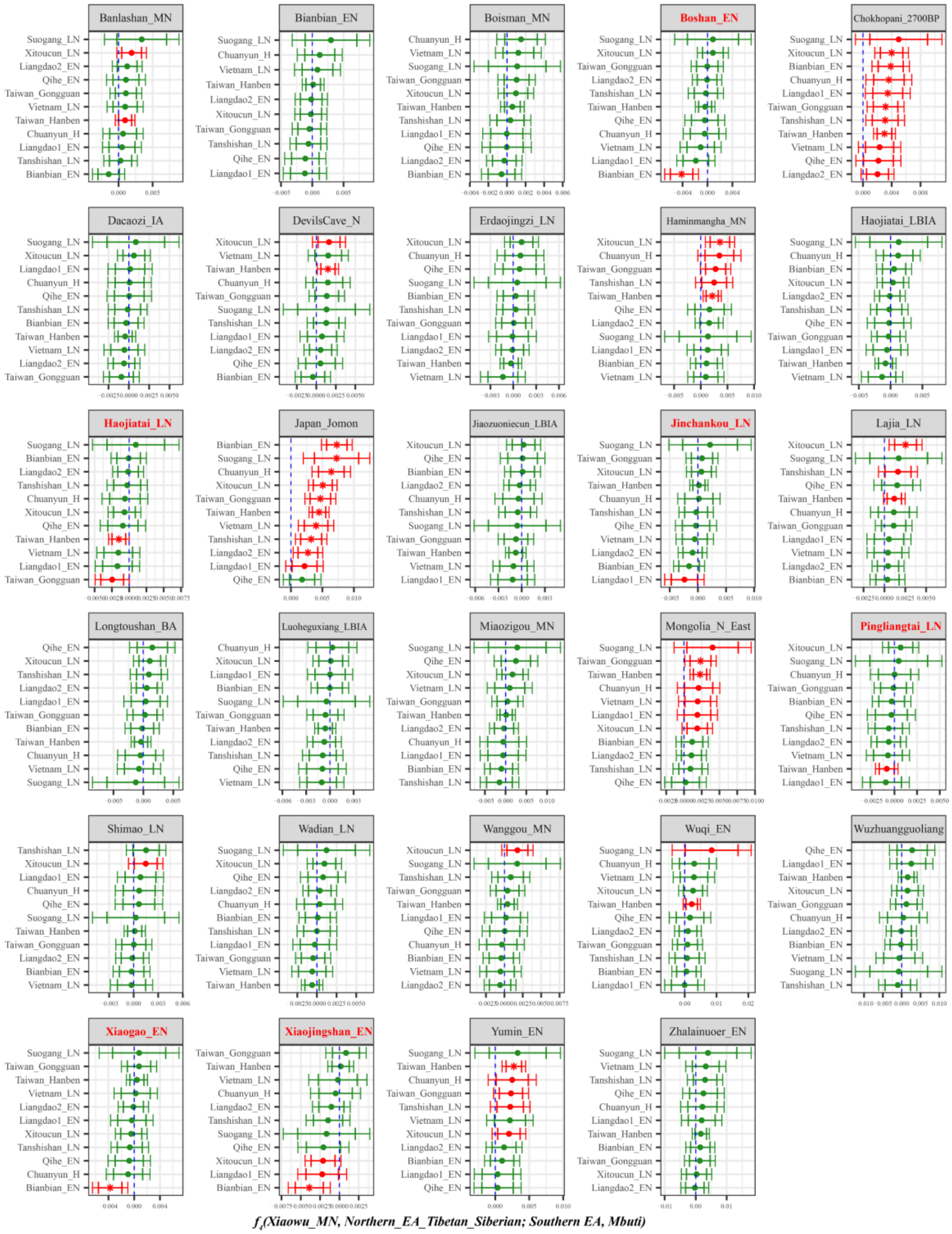

**Supplementary Fig. 35B. Temporal changes of shared genetic drift of ancient populations from Yellow River Basin and other region of northern East Asia assessed via $f_4$*(Xiaowu _MN, Inland/Coastal Neolithic/Bronze Age northern East Asian/Tibet Plateau/Siberia/Japan; Inland/Coastal Neolithic/Bronze Age southern East Asian, Mbuti)*.**

The dashed blue line indicates the zero $f_4$ value. The red asterisk indicates the absolute Z-score value larger than 3, the red point for absolute Z-score value ranging from two to three, and the green point for absolute Z-score value ranging from zero to two. The thick bar denotes two standard errors, and the thin bar for three standard errors. Figures were grouped via the second population in the $f_4$-statistics of Inland/Coastal Neolithic/Bronze Age northern East Asian/Tibet Plateau/Siberia/Japan, which was labeled as the red colour with some significant negative values. Significant negative values denoted the included second populations harboured more ancient southern East Asian related ancestry. Significant positive $f_4$ values indicated the first population had more ancient southern East Asian related ancestry.

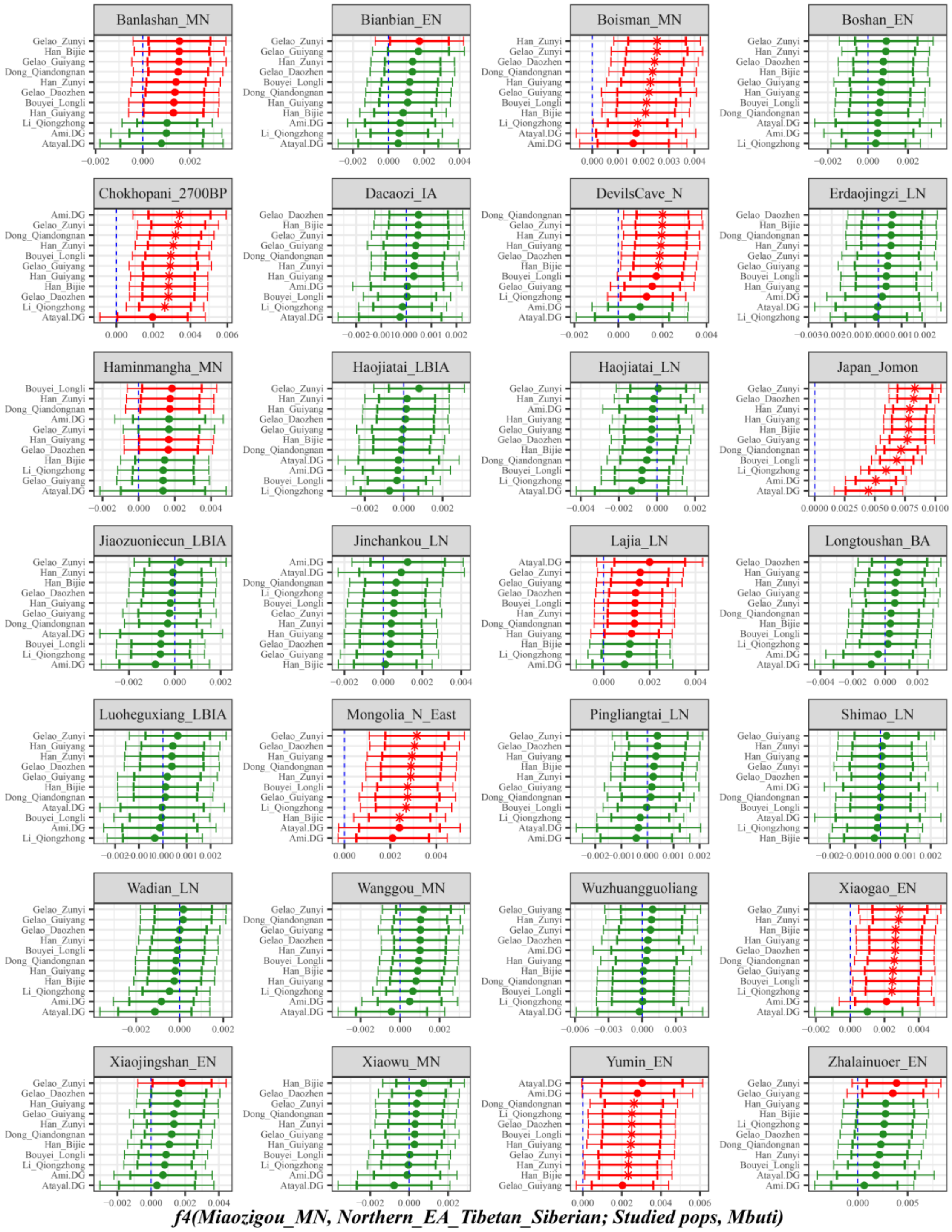

**Supplementary Fig. 36A. Temporal changes of shared genetic drift of ancient populations from Yellow River Basin and other region of northern East Asia assessed via *$f_4$(Miaozigou_MN, Inland/Coastal Neolithic/Bronze Age northern East Asian/Tibet Plateau/Siberia/Japan; Studied inland TK/Sinitic, Mbuti)*.**

The dashed blue line indicates the zero $f_4$ value. The red asterisk indicates the absolute Z-score value larger than 3, the red point for absolute Z-score value ranging from two to three, and the green point for absolute Z-score value ranging from zero to two. The thick bar denotes two standard errors, and the thin bar for three standard errors. Figures were grouped via the second population in the $f_4$-statistics of Inland/Coastal Neolithic/Bronze Age northern East Asian/Tibet Plateau/Siberia/Japan, which was labeled

as the red colour with some significant negative values. Significant negative values denoted the included second populations harboured more central/southern Sinitic or TK related ancestry. Significant positive $f_4$ values indicated the first population had more central/southern Sinitic or TK related ancestry.

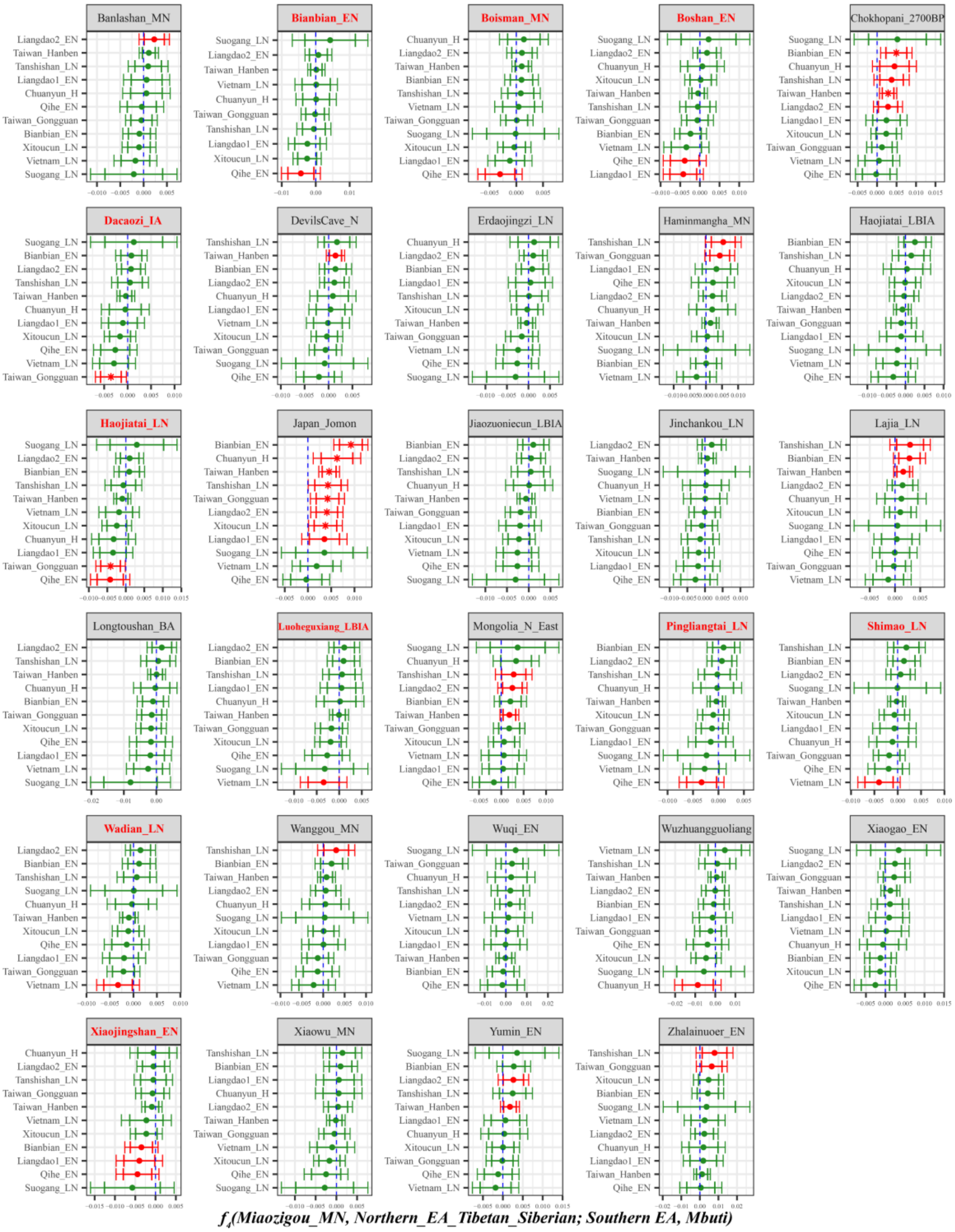

**Supplementary Fig. 36B. Temporal changes of shared genetic drift of ancient populations from Yellow River Basin and other region of northern East Asia assessed via $f_4$*(Miaozigou_MN, Inland/Coastal Neolithic/Bronze Age northern East Asian/Tibet Plateau/Siberia/Japan; Inland/Coastal Neolithic/Bronze Age southern East Asian, Mbuti)*.**

The dashed blue line indicates the zero $f_4$ value. The red asterisk indicates the absolute Z-score value larger than 3, the red point for absolute Z-score value ranging from two to three, and the green point for absolute Z-score value ranging from zero to two. The thick bar denotes two standard errors, and the thin bar for three standard errors. Figures were grouped via the second population in the $f_4$-statistics of

Inland/Coastal Neolithic/Bronze Age northern East Asian/Tibet Plateau/Siberia/Japan, which was labeled as the red colour with some significant negative values. Significant negative values denoted the included second populations harboured more ancient southern East Asian related ancestry. Significant positive $f_4$ values indicated the first population had more ancient southern East Asian related ancestry.

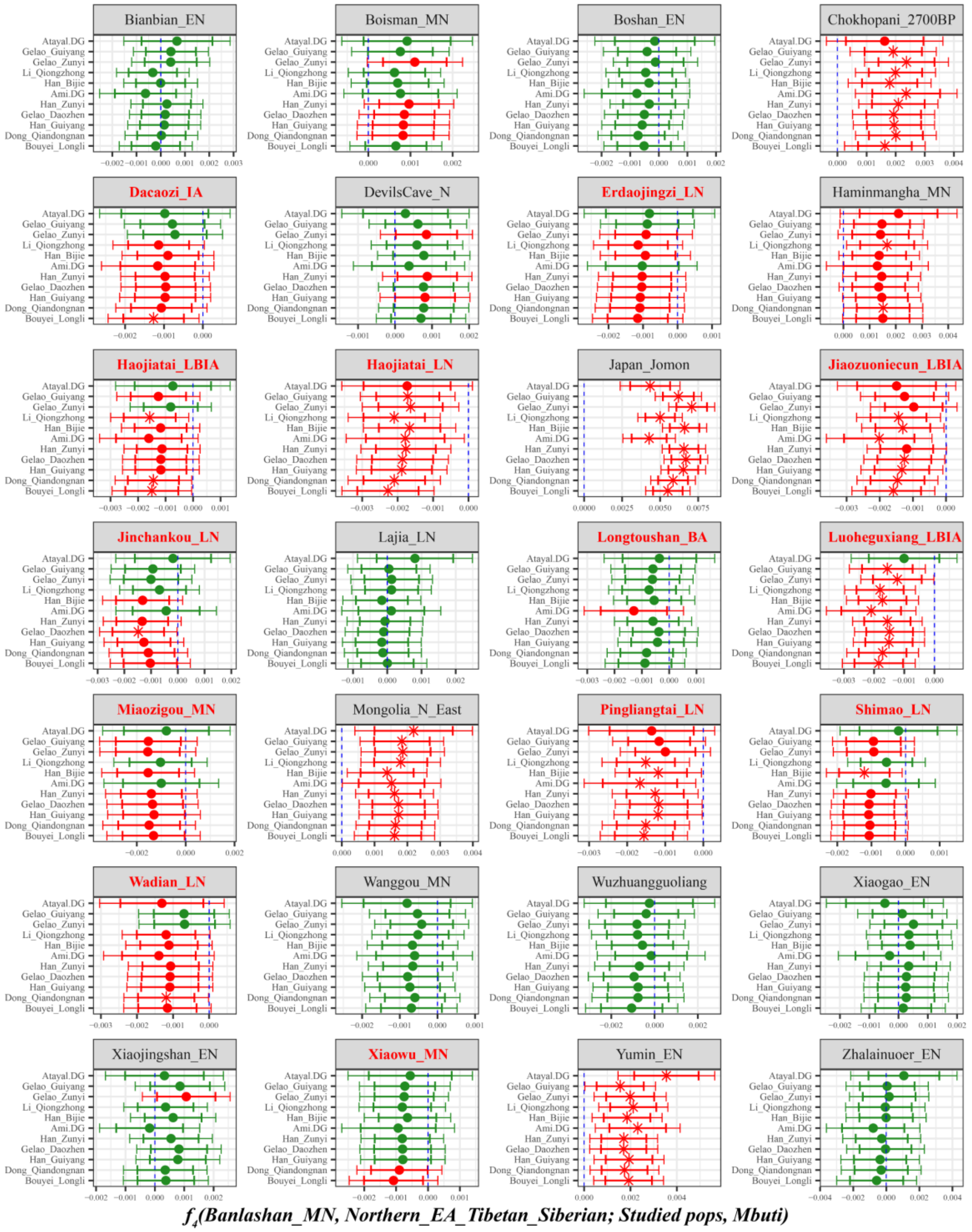

**Supplementary Fig. 37A. Temporal changes of shared genetic drift of ancient populations from Yellow River Basin and other region of northern East Asia assessed via $f_4$(Banlashan_MN, *Inland/Coastal Neolithic/Bronze Age northern East Asian/Tibet Plateau/Siberia/Japan; Studied inland TK/Sinitic, Mbuti)*.**

The dashed blue line indicates the zero $f_4$ value. The red asterisk indicates the absolute Z-score value larger than 3, the red point for absolute Z-score value ranging from two to three, and the green point for absolute Z-score value ranging from zero to two. The thick bar denotes two standard errors, and the thin bar for three standard errors. Figures were grouped via the second population in the $f_4$-statistics of

Inland/Coastal Neolithic/Bronze Age northern East Asian/Tibet Plateau/Siberia/Japan, which was labeled as the red colour with some significant negative values. Significant negative values denoted the included second populations harboured more central/southern Sinitic or TK related ancestry. Significant positive $f_4$ values indicated the first population had more central/southern Sinitic or TK related ancestry.

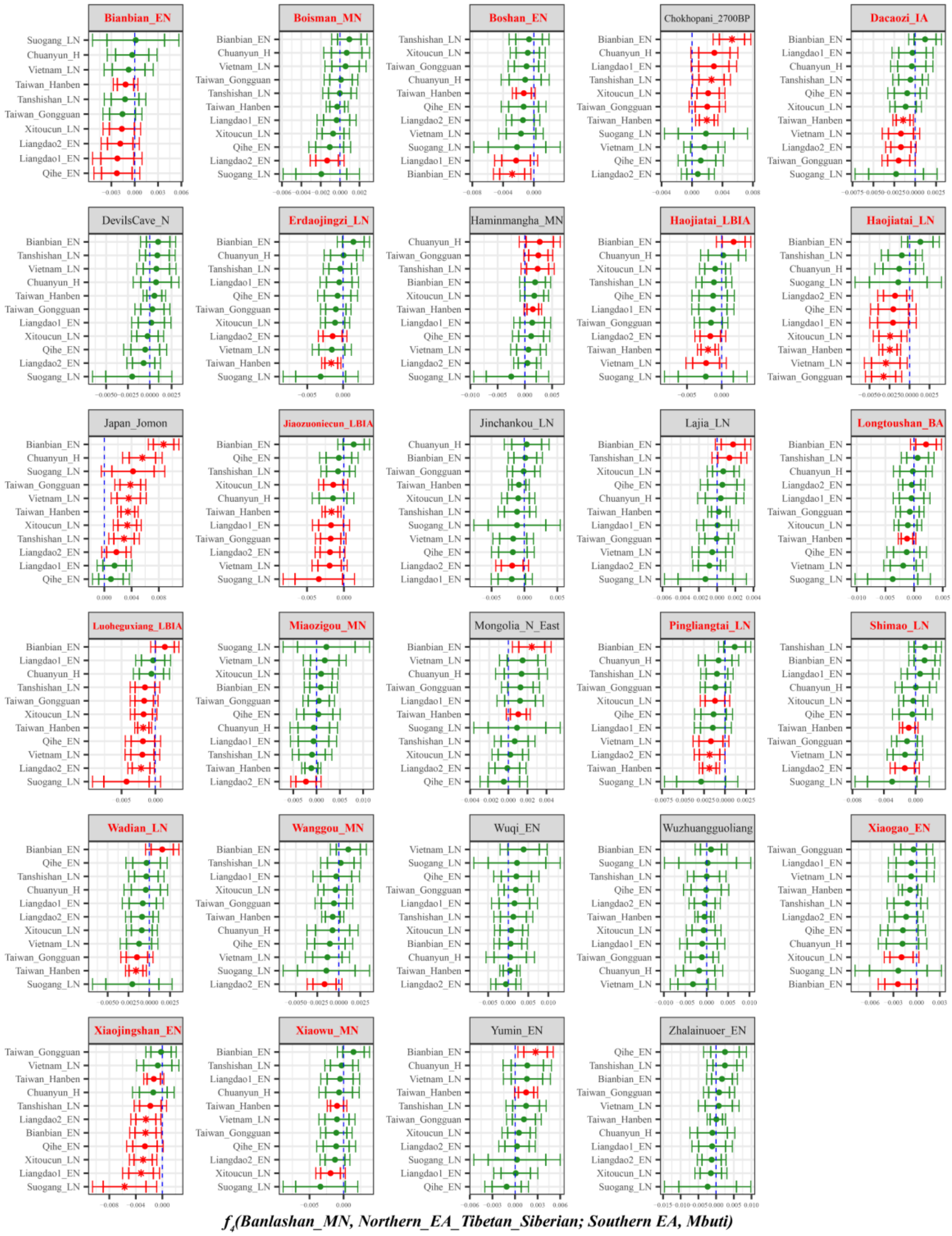

**Supplementary Fig. 37B. Temporal changes of shared genetic drift of ancient populations from Yellow River Basin and other region of northern East Asia assessed via *$f_4$(Banlashan_MN, Inland/Coastal Neolithic/Bronze Age northern East Asian/Tibet Plateau/Siberia/Japan; Inland/Coastal Neolithic/Bronze Age southern East Asian, Mbuti)*.**

The dashed blue line indicates the zero $f_4$ value. The red asterisk indicates the absolute Z-score value larger than 3, the red point for absolute Z-score value ranging from two to three, and the green point for absolute Z-score value ranging from zero to two. The thick bar denotes two standard errors, and the thin bar for three standard errors. Figures were grouped via the second population in the $f_4$-statistics of Inland/Coastal Neolithic/Bronze Age northern East Asian/Tibet Plateau/Siberia/Japan, which was labeled as the red colour with some significant negative values. Significant negative values denoted the included second populations harboured more ancient southern East Asian related ancestry. Significant positive $f_4$ values indicated the first population had more ancient southern East Asian related ancestry.

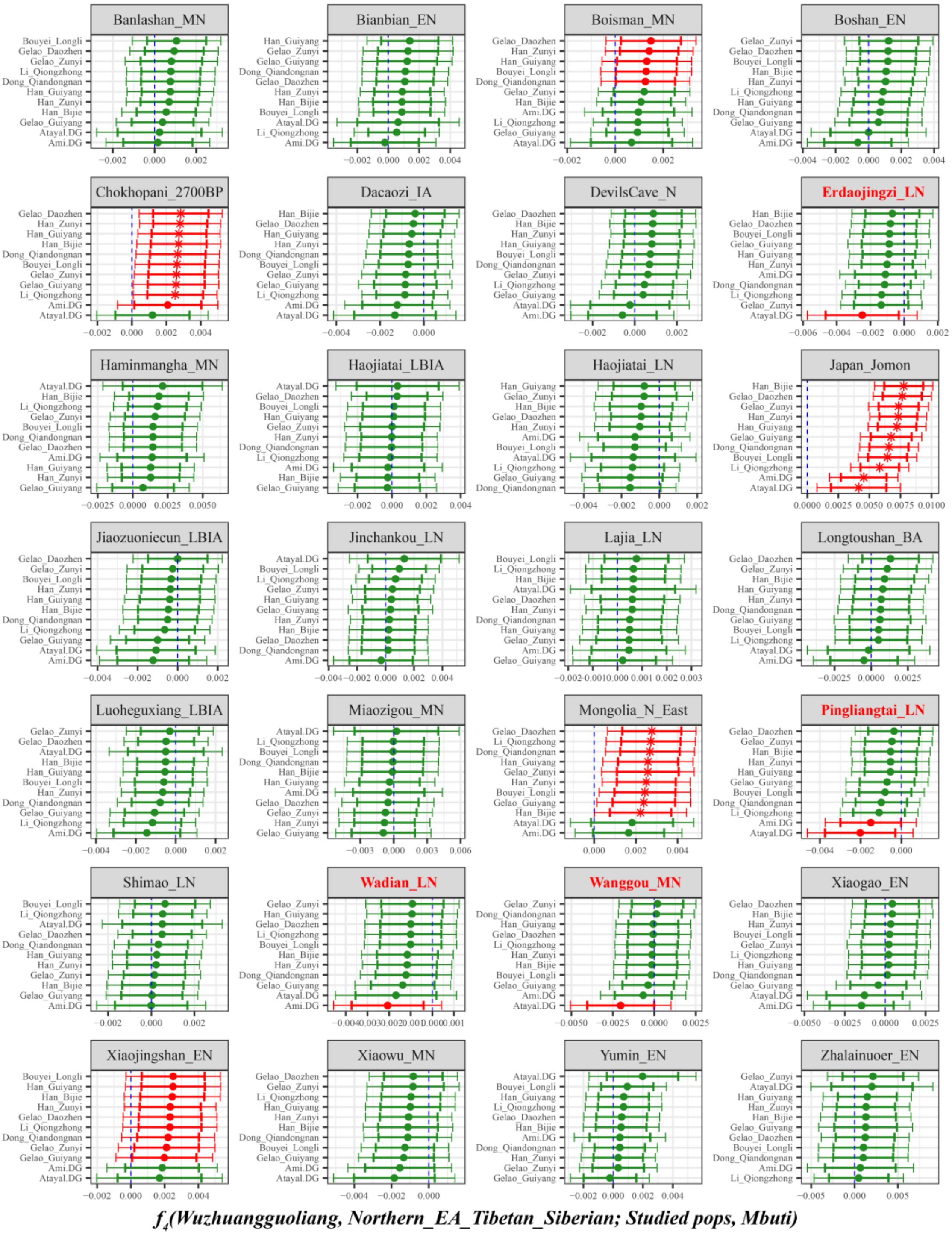

**Supplementary Fig. 38A. Temporal changes of shared genetic drift of ancient populations from Yellow River Basin and other region of northern East Asia assessed via *$f_4$(Wuzhuangguoliang, Inland/Coastal Neolithic/Bronze Age northern East Asian/Tibet Plateau/Siberia/Japan; Studied inland TK/Sinitic, Mbuti)*.**

The dashed blue line indicates the zero $f_4$ value. The red asterisk indicates the absolute Z-score value larger than 3, the red point for absolute Z-score value ranging from two to three, and the green point for

absolute Z-score value ranging from zero to two. The thick bar denotes two standard errors, and the thin bar for three standard errors. Figures were grouped via the second population in the $f_4$-statistics of Inland/Coastal Neolithic/Bronze Age northern East Asian/Tibet Plateau/Siberia/Japan, which was labeled as the red colour with some significant negative values. Significant negative values denoted the included second populations harboured more central/southern Sinitic or TK related ancestry. Significant positive $f_4$ values indicated the first population had more central/southern Sinitic or TK related ancestry.

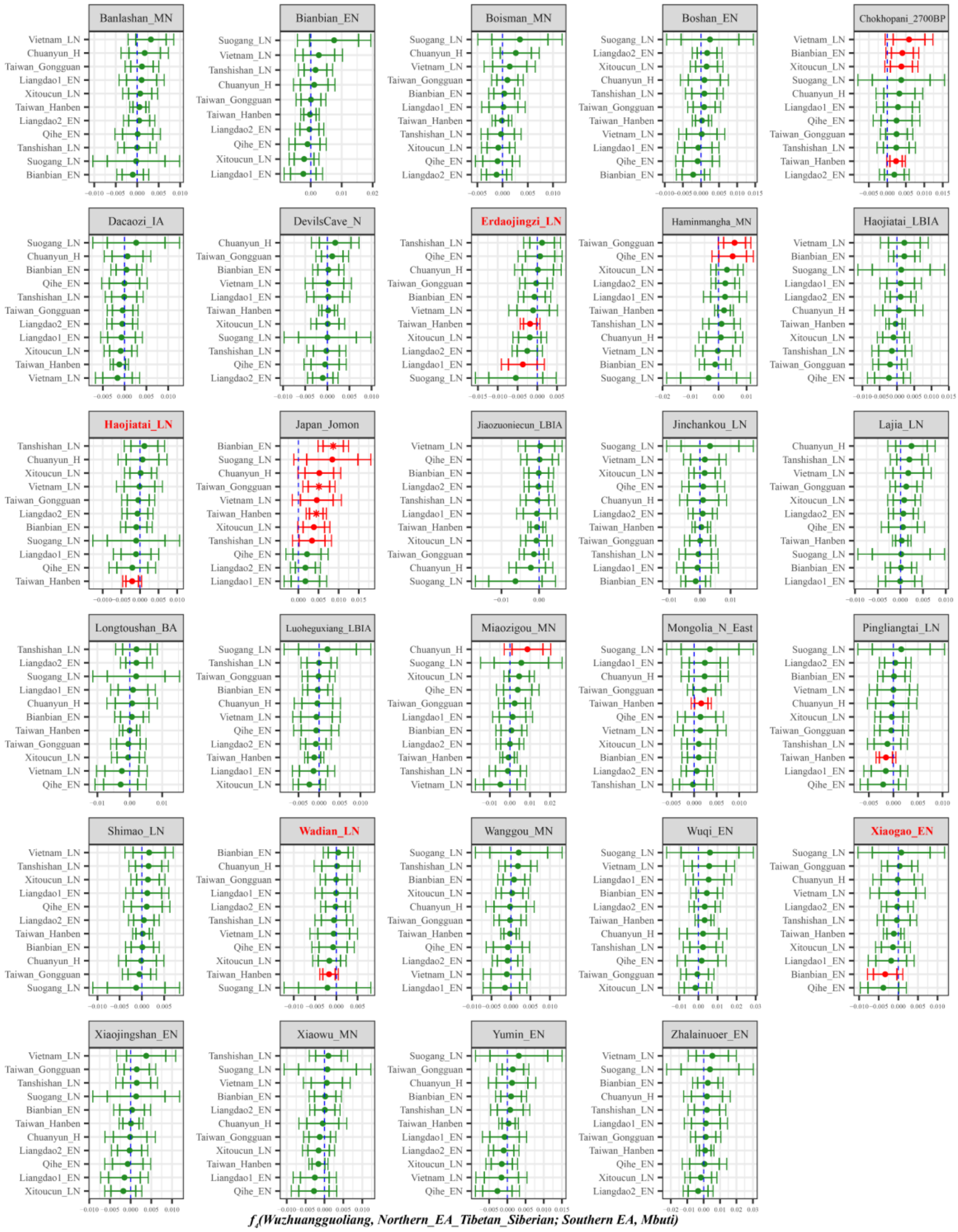

**Supplementary Fig. 38B. Temporal changes of shared genetic drift of ancient populations from Yellow River Basin and other region of northern East Asia assessed via *f₄(Wuzhuangguoliang, Inland/Coastal Neolithic/Bronze Age northern East Asian/Tibet Plateau/Siberia/Japan; Inland/Coastal Neolithic/Bronze Age southern East Asian, Mbuti)*.**

The dashed blue line indicates the zero $f_4$ value. The red asterisk indicates the absolute Z-score value larger than 3, the red point for absolute Z-score value ranging from two to three, and the green point for

absolute Z-score value ranging from zero to two. The thick bar denotes two standard errors, and the thin bar for three standard errors. Figures were grouped via the second population in the $f_4$-statistics of Inland/Coastal Neolithic/Bronze Age northern East Asian/Tibet Plateau/Siberia/Japan, which was labeled as the red colour with some significant negative values. Significant negative values denoted the included second populations harboured more ancient southern East Asian related ancestry. Significant positive $f_4$ values indicated the first population had more ancient southern East Asian related ancestry.

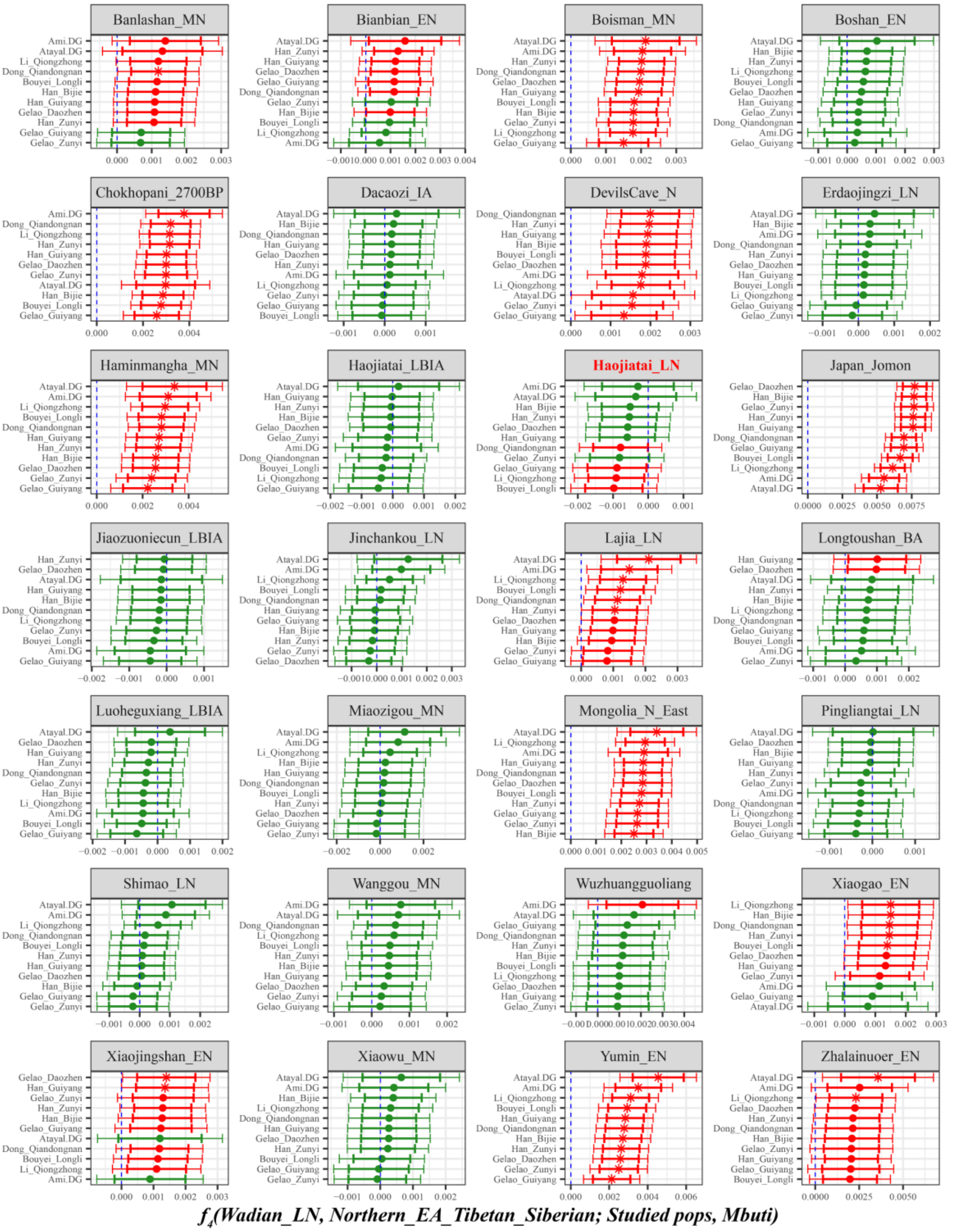

**Supplementary Fig. 39A. Temporal changes of shared genetic drift of ancient populations from Yellow River Basin and other region of northern East Asia assessed via *$f_4$(Wadian_LN, Inland/Coastal Neolithic/Bronze Age northern East Asian/Tibet Plateau/Siberia/Japan; Studied inland TK/Sinitic, Mbuti)*.**

The dashed blue line indicates the zero $f_4$ value. The red asterisk indicates the absolute Z-score value larger than 3, the red point for absolute Z-score value ranging from two to three, and the green point for absolute Z-score value ranging from zero to two. The thick bar denotes two standard errors, and the thin bar for three standard errors. Figures were grouped via the second population in the $f_4$-statistics of Inland/Coastal Neolithic/Bronze Age northern East Asian/Tibet Plateau/Siberia/Japan, which was labeled as the red colour with some significant negative values. Significant negative values denoted the included second populations harboured more central/southern Sinitic or TK related ancestry. Significant positive $f_4$ values indicated the first population had more central/southern Sinitic or TK related ancestry.

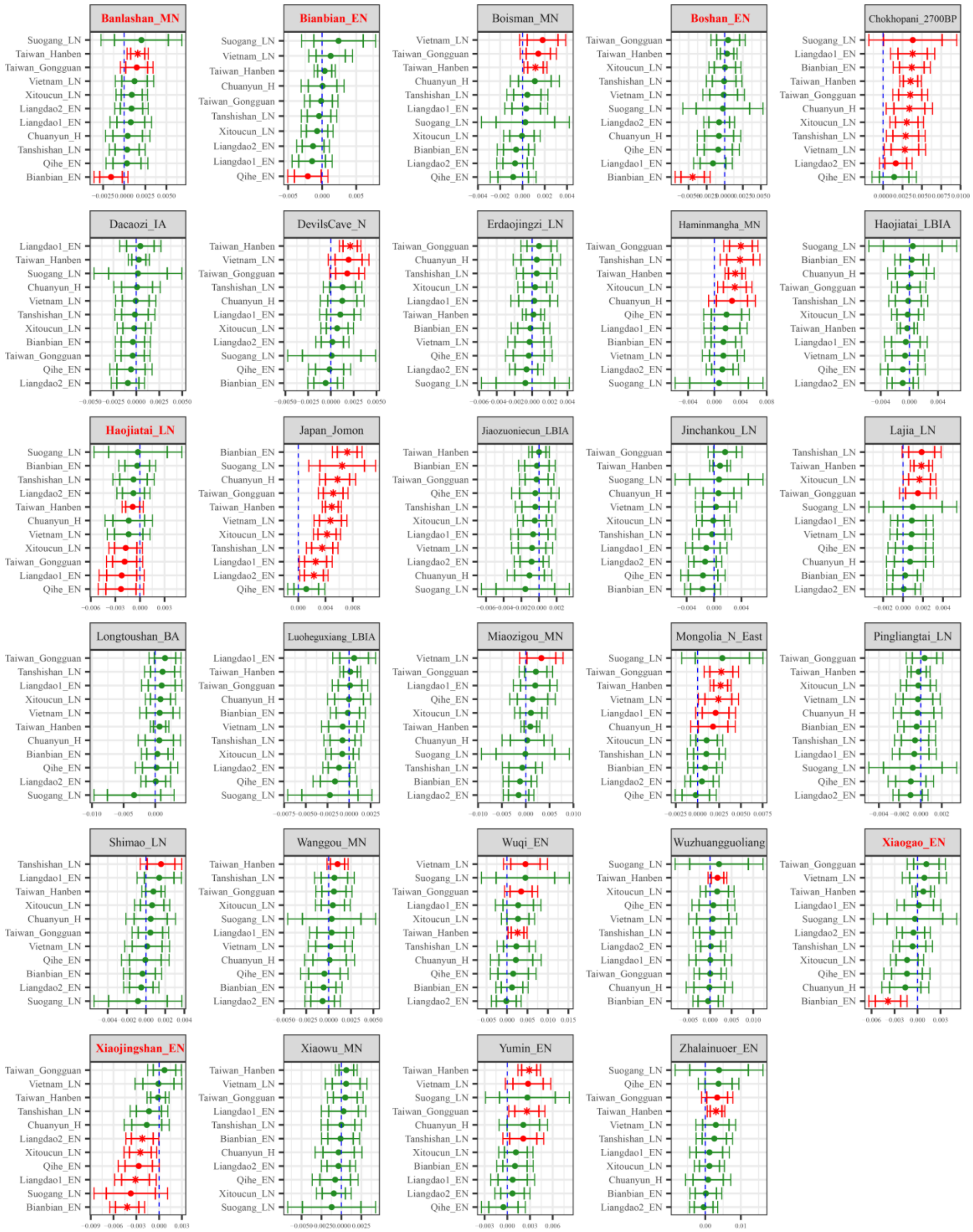

$f_4$(Wadian_LN, Northern_EA_Tibetan_Siberian; Southern EA, Mbuti)

**Supplementary Fig. 39B. Temporal changes of shared genetic drift of ancient populations from Yellow River Basin and other region of northern East Asia assessed via *$f_4$(Wadian_LN, Inland/Coastal Neolithic/Bronze Age northern East Asian/Tibet Plateau/Siberia/Japan; Inland/Coastal Neolithic/Bronze Age southern East Asian, Mbuti)*.**

The dashed blue line indicates the zero $f_4$ value. The red asterisk indicates the absolute Z-score value larger than 3, the red point for absolute Z-score value ranging from two to three, and the green point for absolute Z-score value ranging from zero to two. The thick bar denotes two standard errors, and the thin bar for three standard errors. Figures were grouped via the second population in the $f_4$-statistics of Inland/Coastal Neolithic/Bronze Age northern East Asian/Tibet Plateau/Siberia/Japan, which was labeled as the red colour with some significant negative values. Significant negative values denoted the included second populations harboured more ancient southern East Asian related ancestry. Significant positive $f_4$ values indicated the first population had more ancient southern East Asian related ancestry.

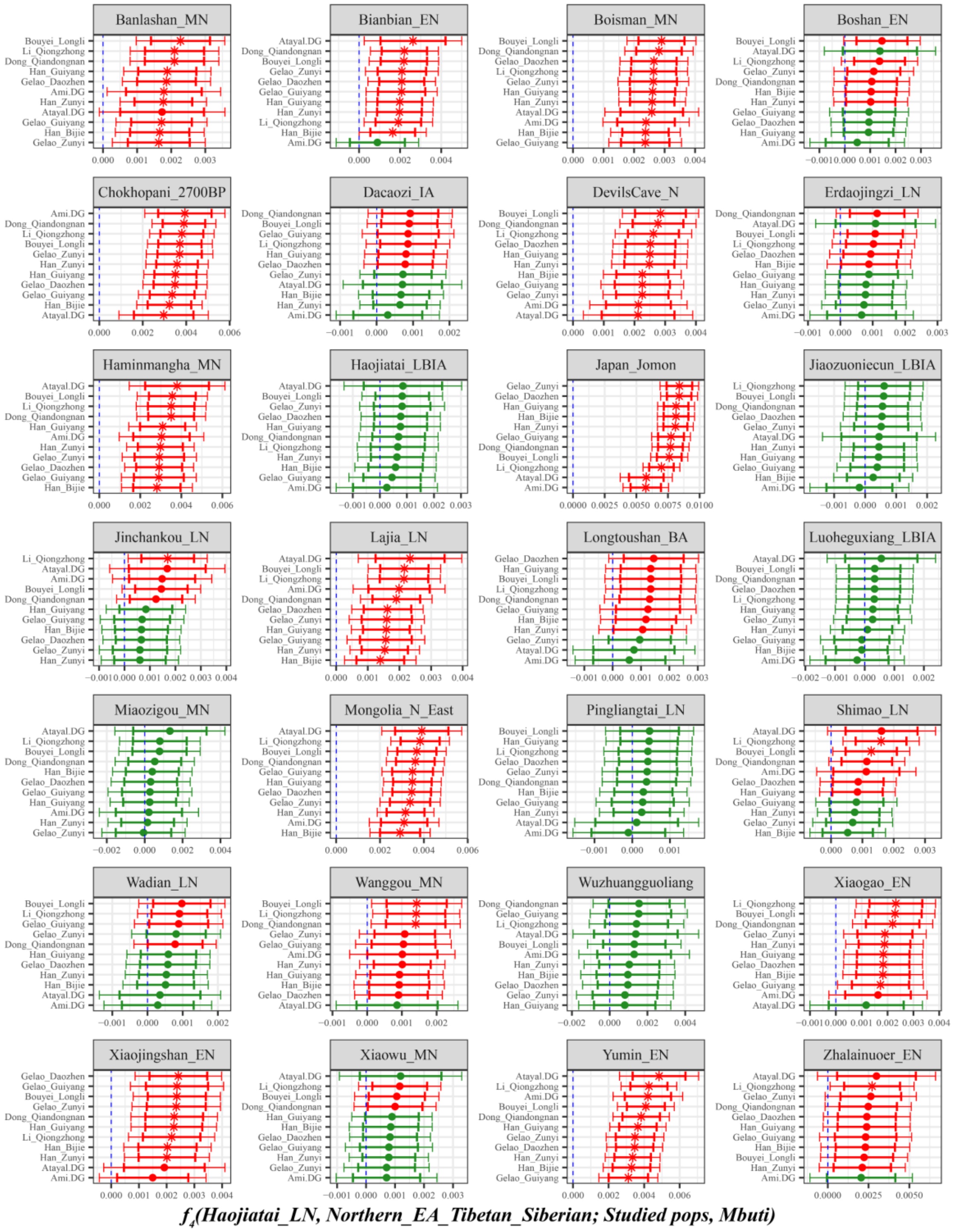

**Supplementary Fig. 40A. Temporal changes of shared genetic drift of ancient populations from Yellow River Basin and other region of northern East Asia assessed via *$f_4$(Haojiatai_LN, Inland/Coastal Neolithic/Bronze Age northern East Asian/Tibet Plateau/Siberia/Japan; Studied inland TK/Sinitic, Mbuti)*.**

The dashed blue line indicates the zero $f_4$ value. The red asterisk indicates the absolute Z-score value larger than 3, the red point for absolute Z-score value ranging from two to three, and the green point for absolute Z-score value ranging from zero to two. The thick bar denotes two standard errors, and the thin bar for three standard errors. Figures were grouped via the second population in the $f_4$-statistics of Inland/Coastal Neolithic/Bronze Age northern East Asian/Tibet Plateau/Siberia/Japan, which was labeled as the red colour with some significant negative values. Significant negative values denoted the included second populations harboured more central/southern Sinitic or TK related ancestry. Significant positive $f_4$ values indicated the first population had more central/southern Sinitic or TK related ancestry.

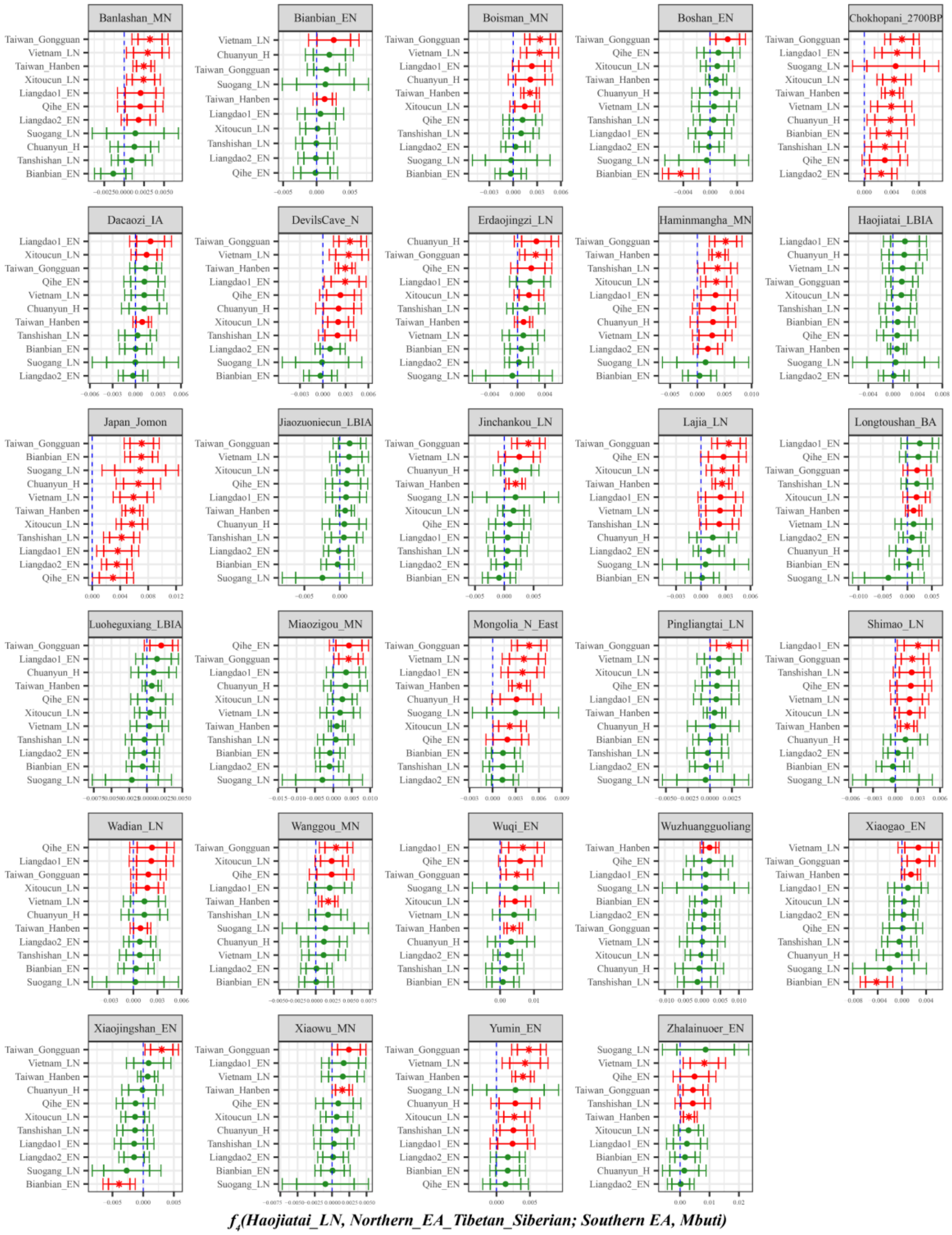

**Supplementary Fig. 40B. Temporal changes of shared genetic drift of ancient populations from Yellow River Basin and other region of northern East Asia assessed via $f_4$*(Haojiatai_LN, Inland/Coastal Neolithic/Bronze Age northern East Asian/Tibet Plateau/Siberia/Japan; Inland/Coastal Neolithic/Bronze Age southern East Asian, Mbuti)*.**

The dashed blue line indicates the zero $f_4$ value. The red asterisk indicates the absolute Z-score value larger than 3, the red point for absolute Z-score value ranging from two to three, and the green point for absolute Z-score value ranging from zero to two. The thick bar denotes two standard errors, and the thin bar for three standard errors. Figures were grouped via the second population in the $f_4$-statistics of Inland/Coastal Neolithic/Bronze Age northern East Asian/Tibet Plateau/Siberia/Japan, which was labeled as the red colour with some significant negative values. Significant negative values denoted the included second populations harboured more ancient southern East Asian related ancestry. Significant positive $f_4$ values indicated the first population had more ancient southern East Asian related ancestry.
