## Supplementary Figures S1-20 for "New insights from the combined discrimination of modern/ancient genome-wide shared alleles and haplotypes: Differentiated demographic history reconstruction of Tai-Kadai and Sinitic people in South China"

[1]State Key Laboratory of Cellular Stress Biology, School of Life Sciences, Department of Anthropology and Ethnology, Institute of Anthropology, State Key Laboratory of Marine Environmental Science, Xiamen University, Xiamen 361005, PR China
[2]Institute of Forensic Medicine, West China School of Basic Science and Forensic Medicine, Sichuan University, Chengdu 610000, PR China
[3]School of Forensic Medicine, Zunyi Medical University, Zunyi 563000, Guizhou, PR China
[4]Department of Forensic Medicine, College of Basic Medicine, Chongqing Medical University, Chongqin 400016, PR China
[5]Department of Forensic Medicine, Guizhou Medical University, Guiyang 550000, PR China
[6]Hunan Key Lab of Bioinformatics, School of Computer Science and Engineering, Central South University, Changsha, 410075, PR China
[7]School of Humanities, Nanyang Technological University, Nanyang, 639798, Singapore
[8]School of Basic Medical Sciences, Zhejiang University School of Medicine, Hangzhou 310000, PR China
[9]Institute of Asian Civilizations, Zhejiang University, Hangzhou 310000, PR China
*These authors contributed equally to this work and should be considered co-first authors.

**[#]Corresponding author**
**Guanglin He**
**Affiliation**: State Key Laboratory of Cellular Stress Biology, School of Life Sciences, Department of Anthropology and Ethnology, Institute of Anthropology, State Key Laboratory of Marine Environmental Science, Xiamen University, Xiamen 361005, PR China; School of Humanities, Nanyang Technological University
**Hui-Yuan Yeh**
**Affiliation**: School of Humanities, Nanyang Technological University, Nanyang, 639798, Singapore
**Chuan-Chao Wang**
**Affiliation**: State Key Laboratory of Cellular Stress Biology, School of Life Sciences, Department of Anthropology and Ethnology, Institute of Anthropology, State Key Laboratory of Marine Environmental Science, Xiamen University, Xiamen 361005, PR China

### Contents of Supplementary Figures

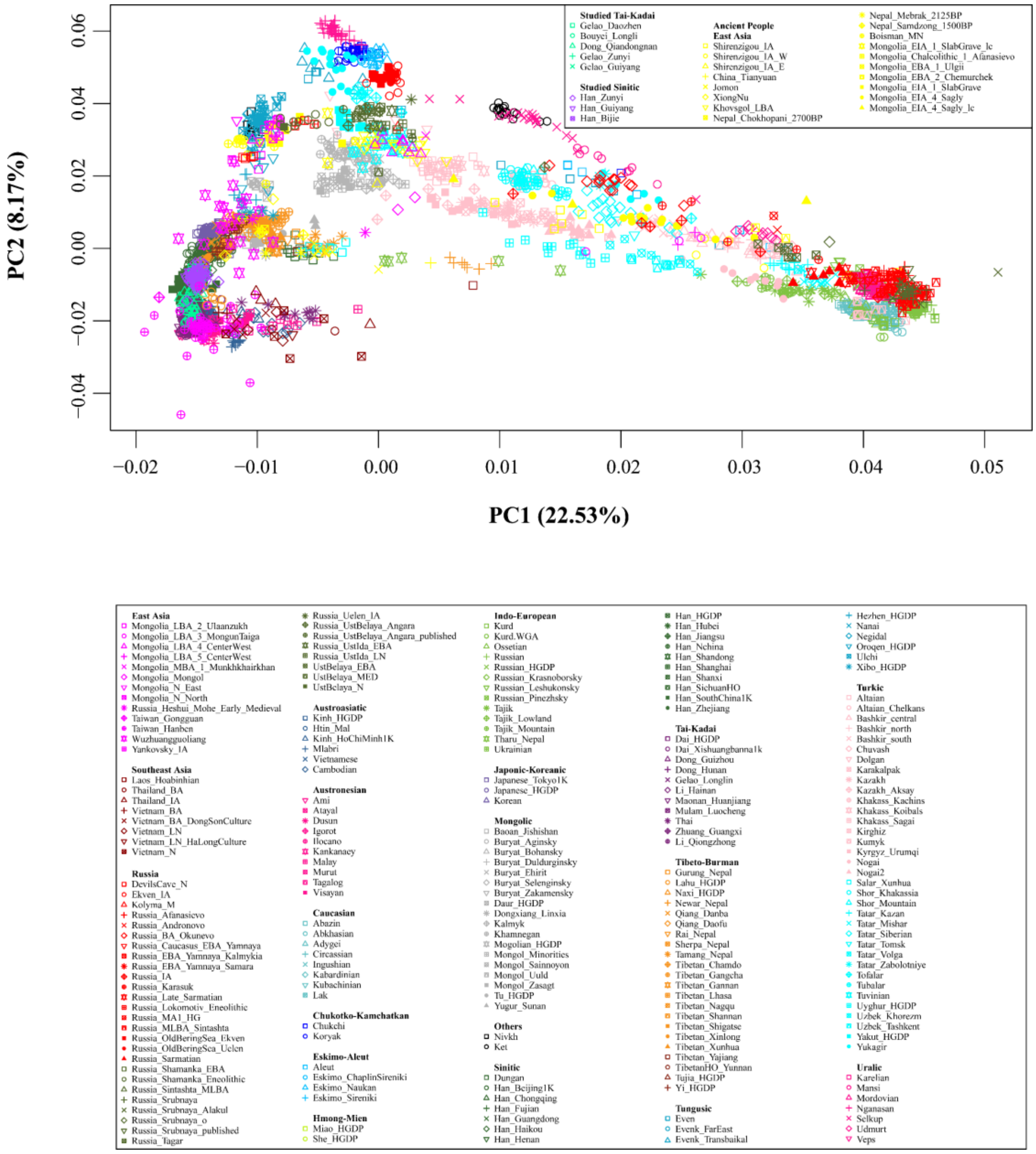

**Supplementary Fig. 1. Eurasian-based principal component analysis.**
Eurasian-based principal component analysis was conducted based on the genetic variations of modern Eurasian populations and Eurasian ancient people were projected onto it.

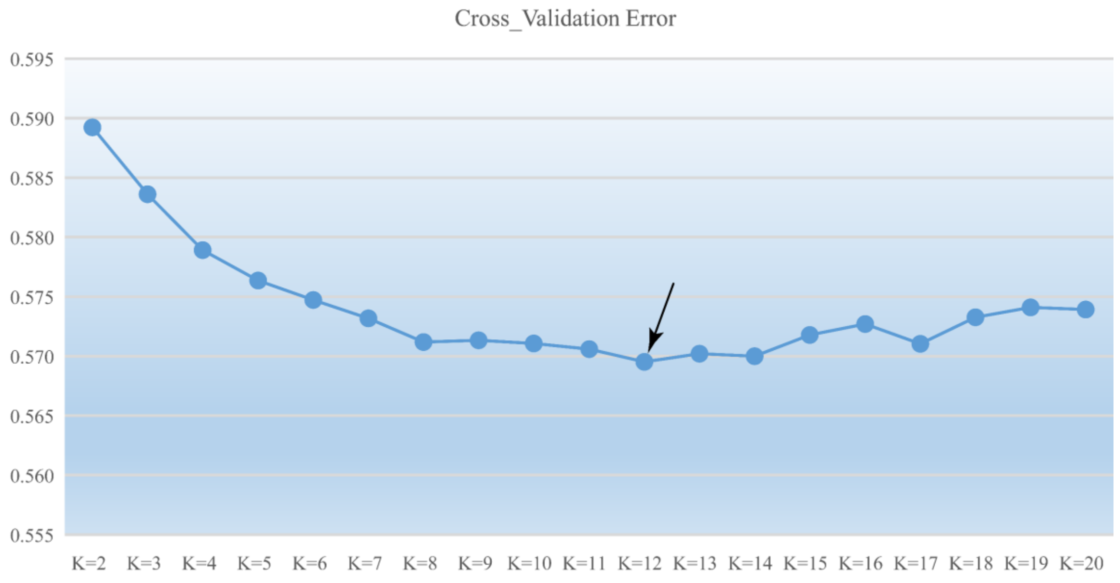

**Supplementary Fig. 2. The cross-validation error of ADMIXTURE analysis.**
The optimal K value of 12 with the lowest cross validation error was identified.

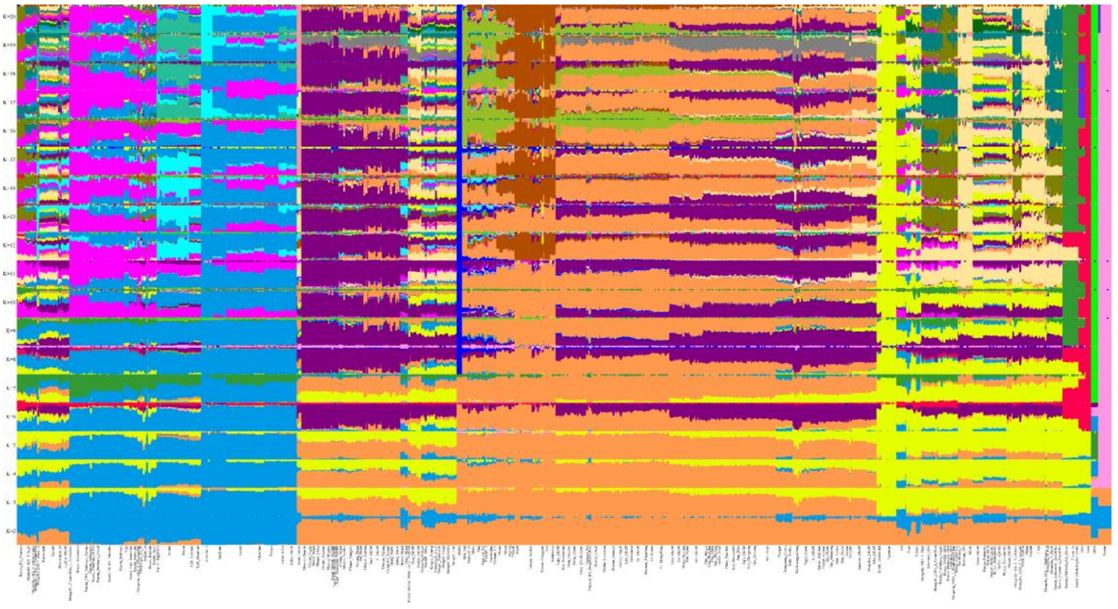

**Supplementary Fig. 3. ADMIXTURE results with K values ranging from 2 to 20 showed the ancestral composition of studied and reference individuals.**

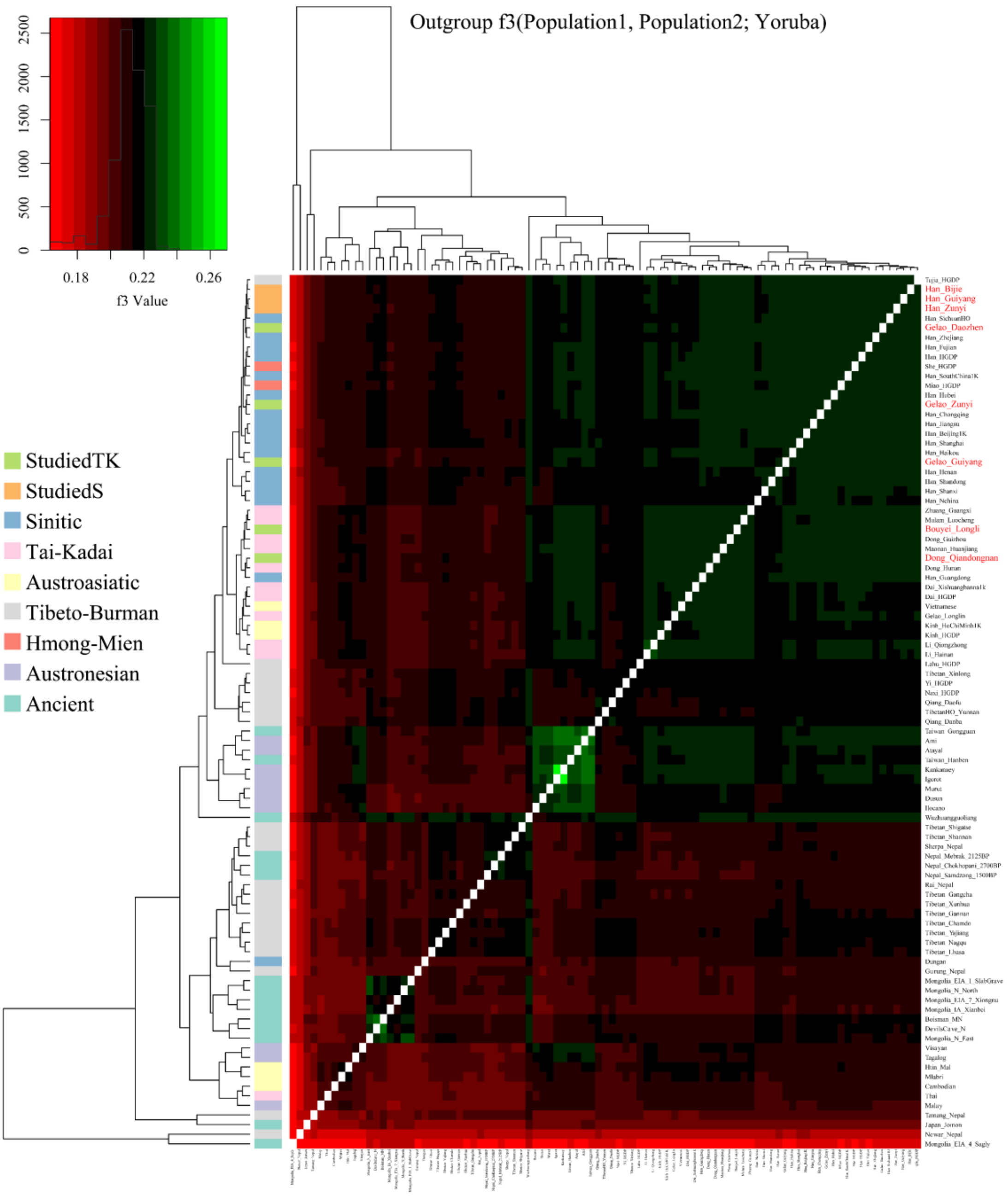

**Supplementary Fig. 4. The shared drift between newly studied populations and reference populations measured by outgroup *f*$_3$*-statistics* of the form *f*$_3$*(X, Y; Yoruba)*.**

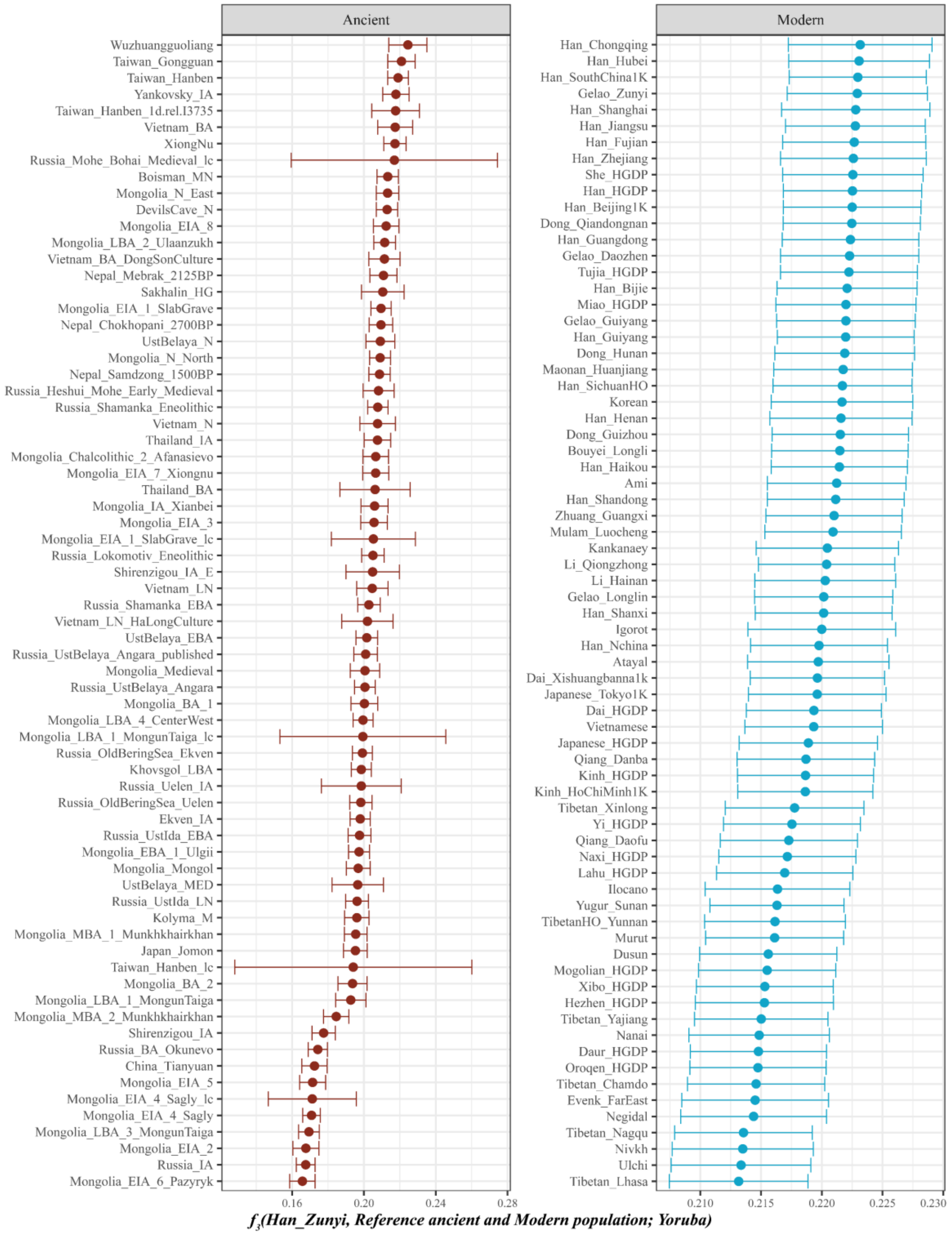

**Supplementary Fig. 5. The shared drift between Zunyi Han and reference ancient and modern populations.**

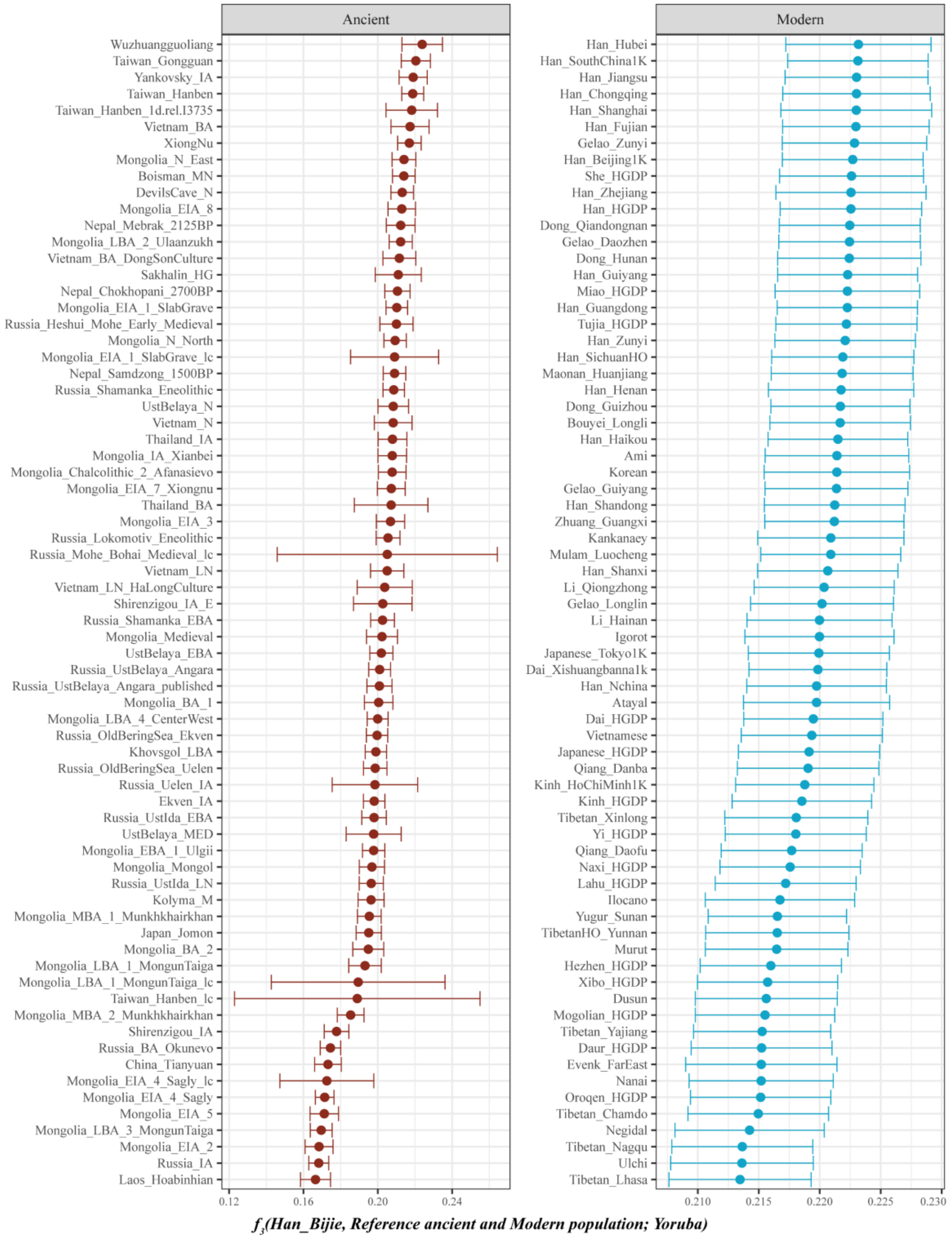

**Supplementary Fig. 6. The shared drift between Bijie Han and reference ancient and modern populations.**

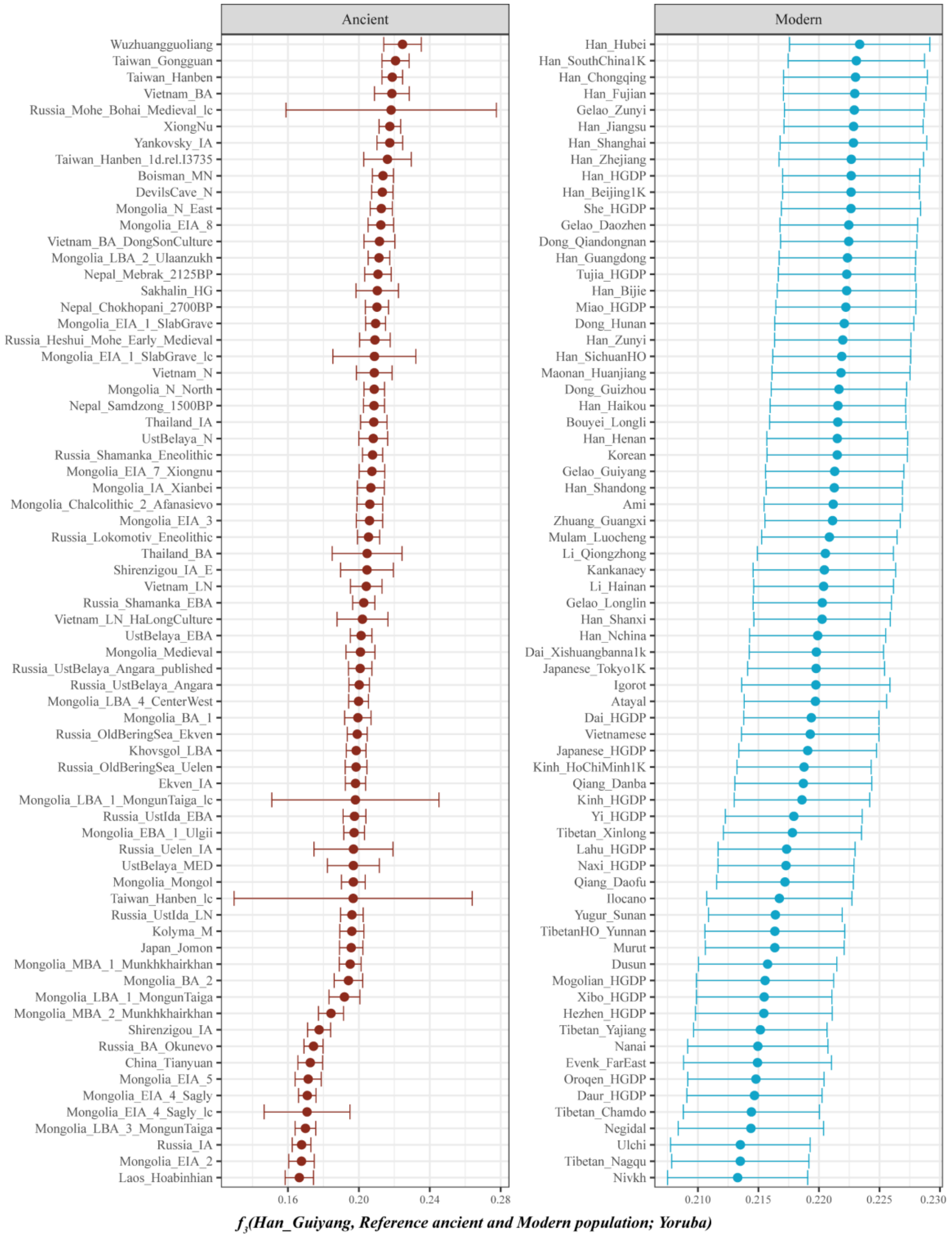

**Supplementary Fig. 7. The shared drift between Guiyang Han and reference ancient and modern populations.**

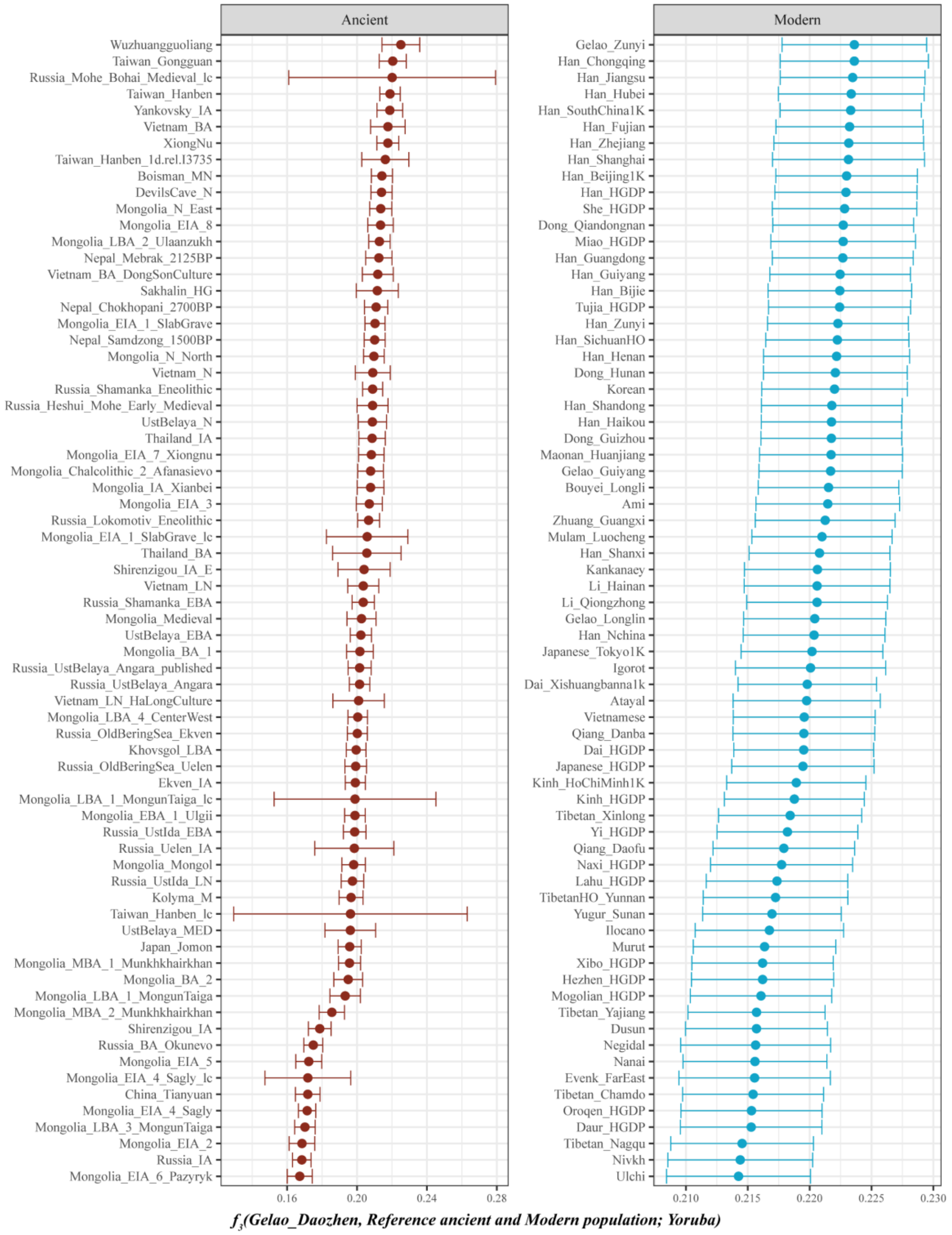

**Supplementary Fig. 8. The shared drift between Daozhen Gelao and reference ancient and modern populations.**

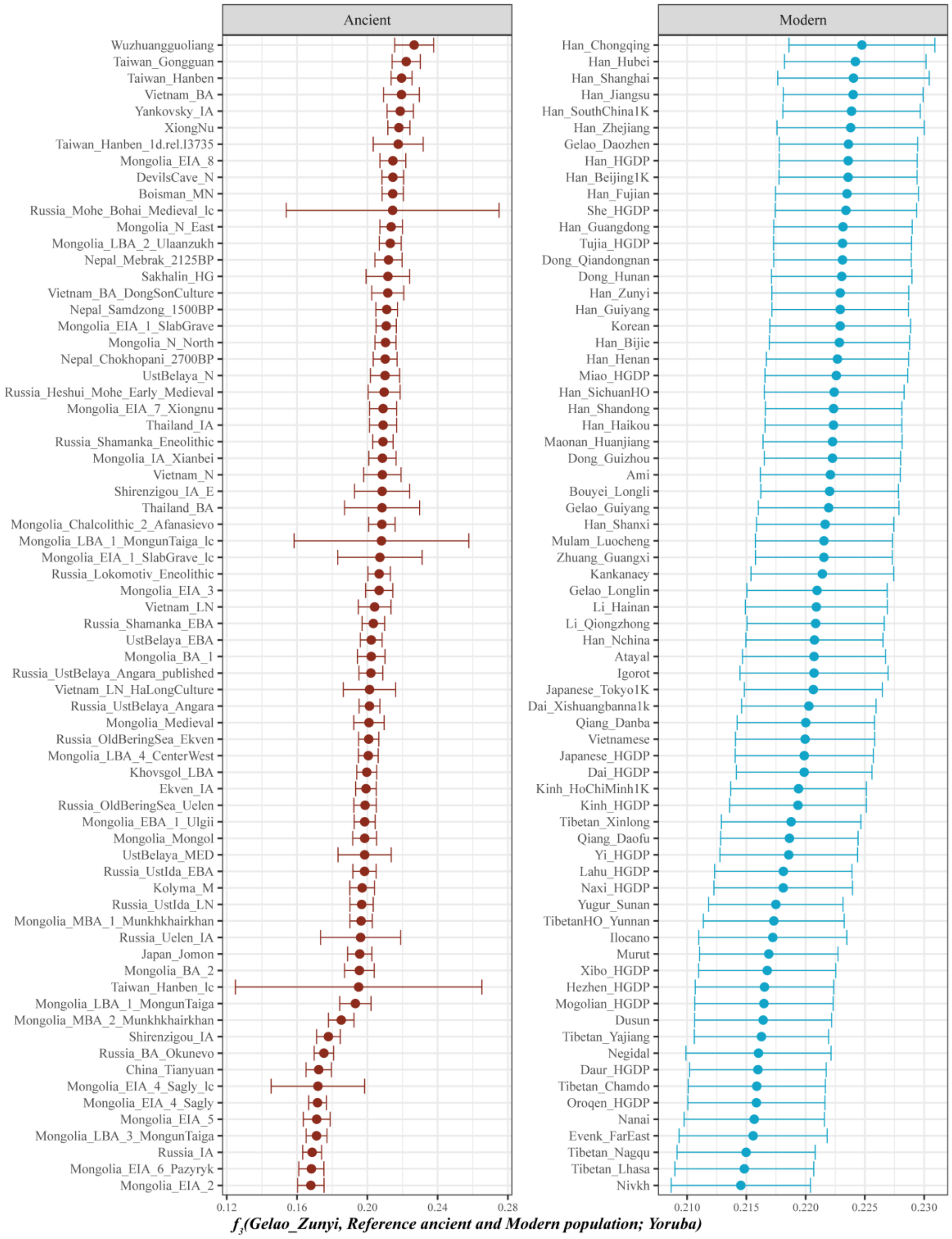

**Supplementary Fig. 9. The shared drift between Zunyi Gelao and reference ancient and modern populations.**

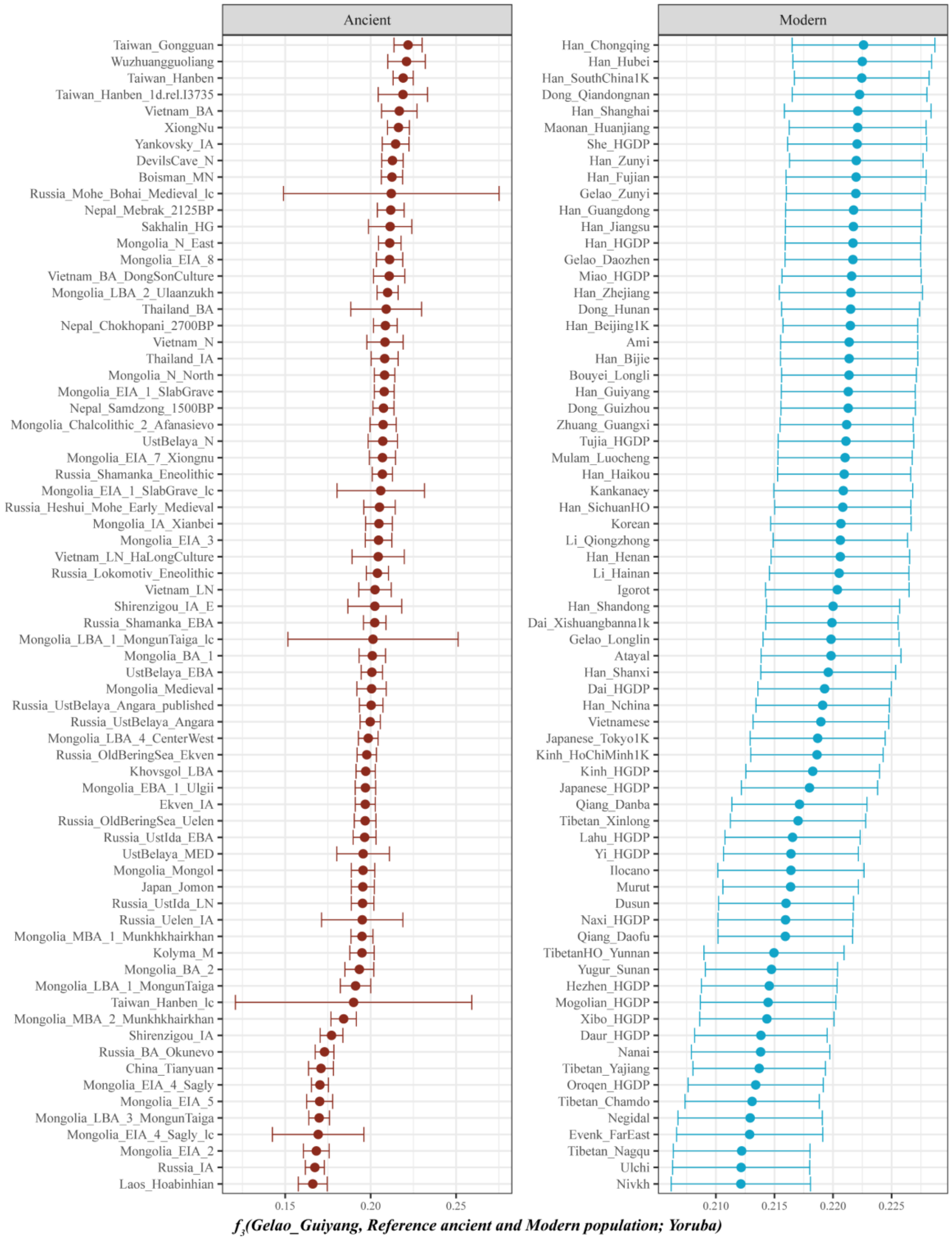

**Supplementary Fig. 10. The shared drift between Guiyang Gelao and reference ancient and modern populations.**

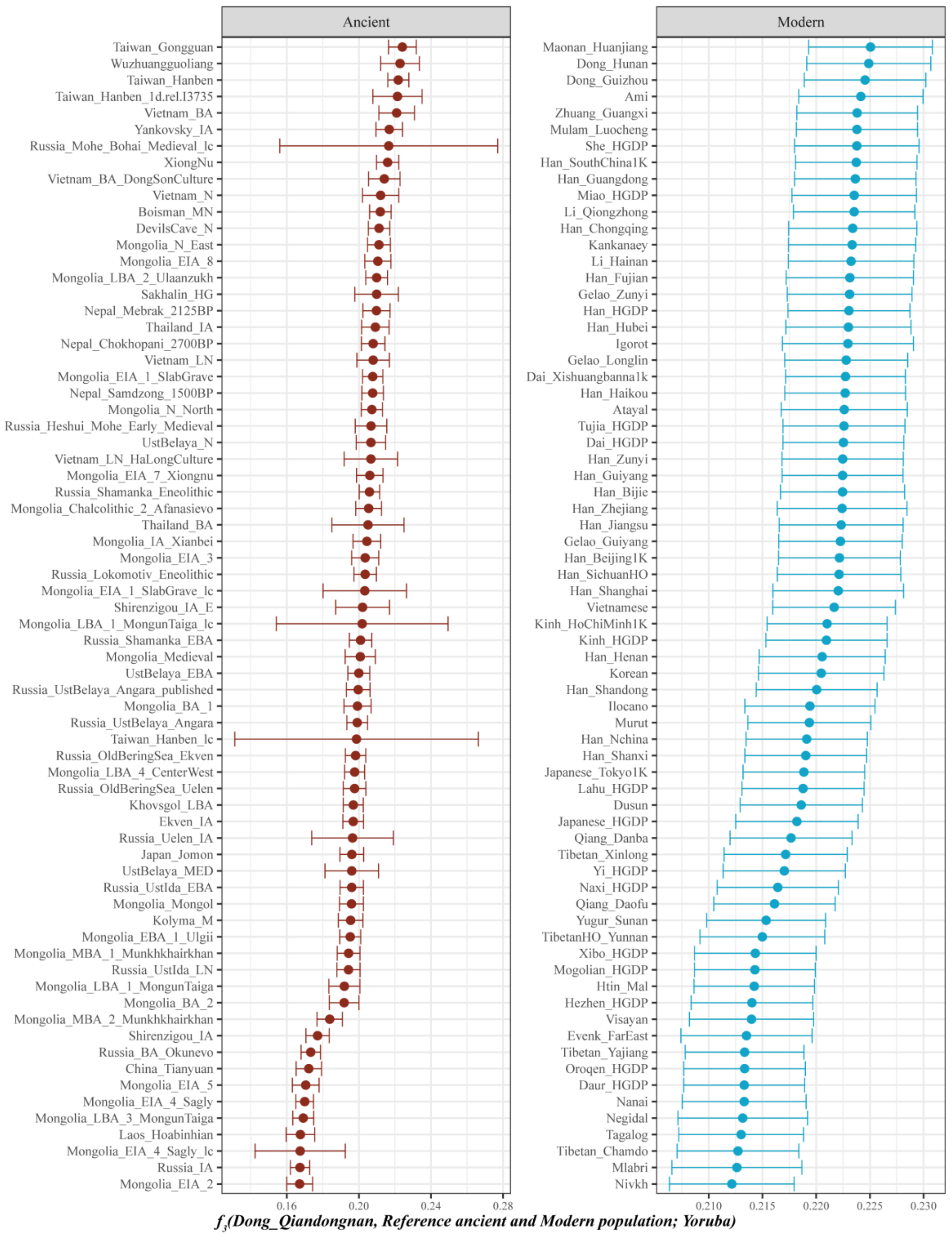

**Supplementary Fig. 11. The shared drift between Qiandongnan Dong and reference ancient and modern populations.**

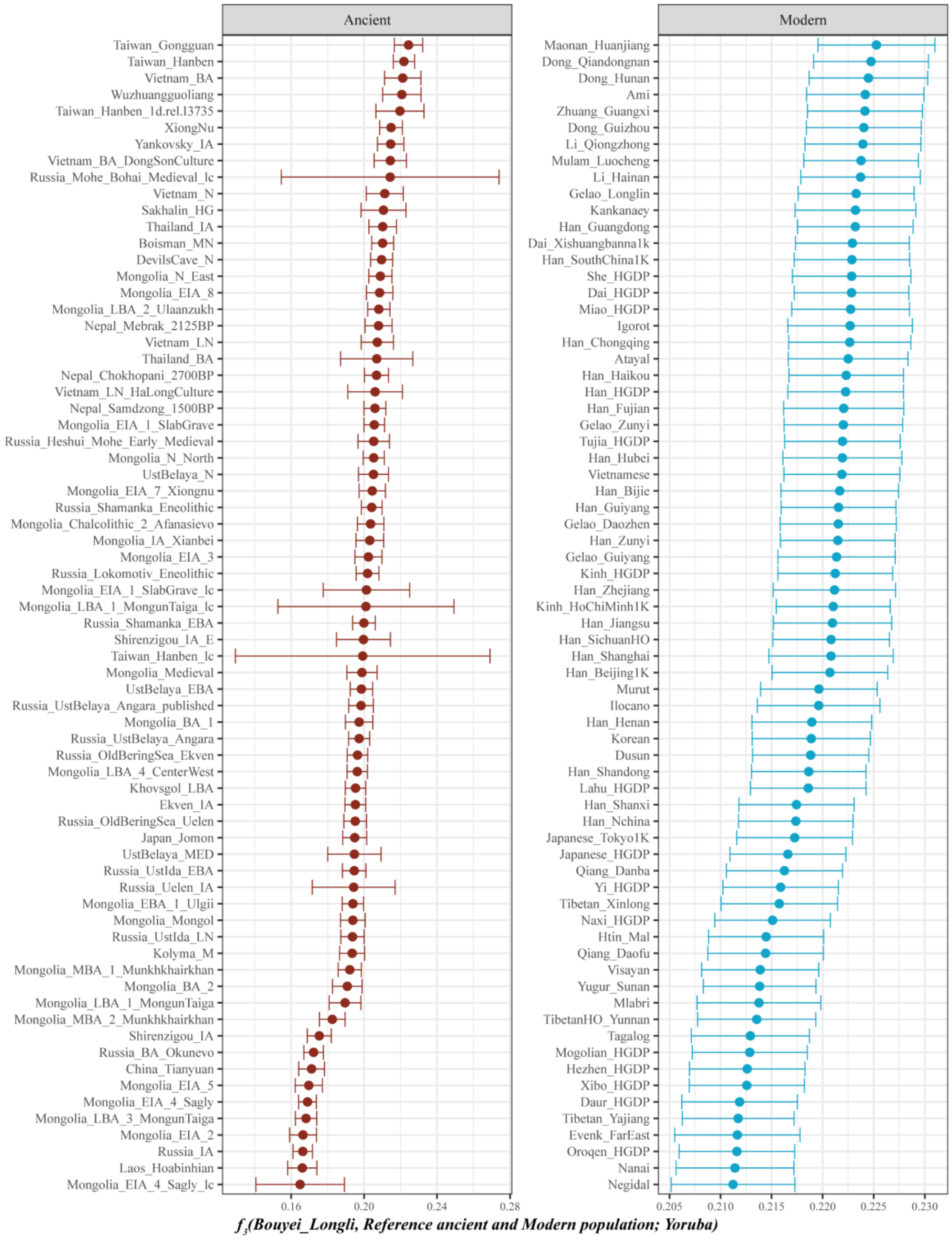

**Supplementary Fig. 12. The shared drift between Longli Bouyei and reference ancient and modern populations.**

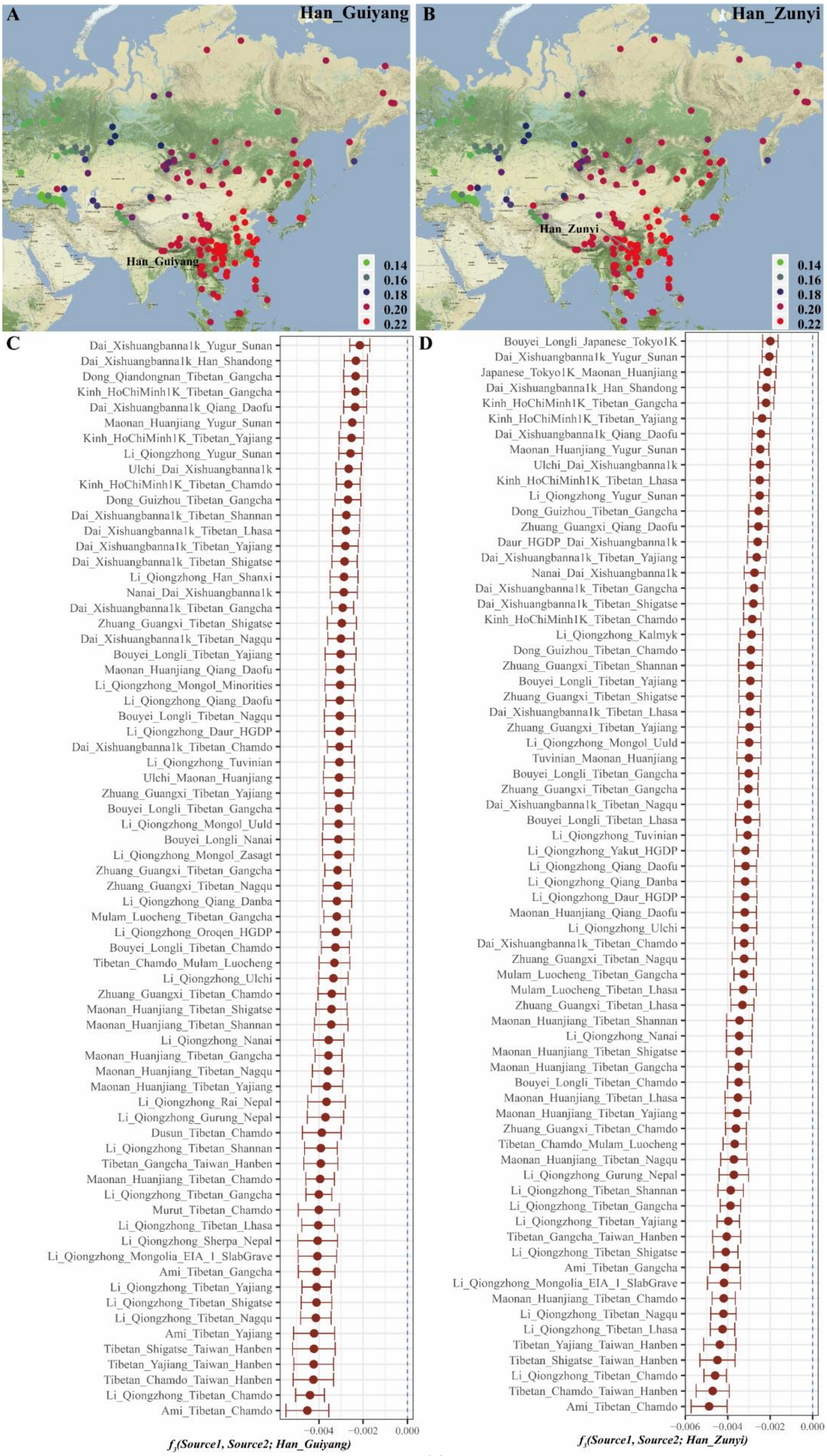
A
Han_Guiyang
Han_Guiyang
0.14
0.16
0.18
0.20
0.22
B
Han_Zunyi
Han_Zunyi
0.14
0.16
0.18
0.20
0.22
C
Dai_Xishuangbanna1k_Yugur_Sunan
Dai_Xishuangbanna1k_Han_Shandong
Dong_Qiandongnan_Tibetan_Gangcha
Kinh_HoChiMinh1K_Tibetan_Gangcha
Dai_Xishuangbanna1k_Qiang_Daofu
Maonan_Huanjiang_Yugur_Sunan
Kinh_HoChiMinh1K_Tibetan_Yajiang
Li_Qiongzhong_Yugur_Sunan
Ulchi_Dai_Xishuangbanna1k
Kinh_HoChiMinh1K_Tibetan_Chamdo
Dong_Guizhou_Tibetan_Gangcha
Dai_Xishuangbanna1k_Tibetan_Shannan
Dai_Xishuangbanna1k_Tibetan_Lhasa
Dai_Xishuangbanna1k_Tibetan_Yajiang
Dai_Xishuangbanna1k_Tibetan_Shigatse
Li_Qiongzhong_Han_Shanxi
Nanai_Dai_Xishuangbanna1k
Dai_Xishuangbanna1k_Tibetan_Gangcha
Zhuang_Guangxi_Tibetan_Shigatse
Dai_Xishuangbanna1k_Tibetan_Nagqu
Bouyei_Longli_Tibetan_Yajiang
Maonan_Huanjiang_Qiang_Daofu
Li_Qiongzhong_Mongol_Minorities
Li_Qiongzhong_Qiang_Daofu
Bouyei_Longli_Tibetan_Nagqu
Li_Qiongzhong_Daur_HGDP
Dai_Xishuangbanna1k_Tibetan_Chamdo
Li_Qiongzhong_Tuvinian
Ulchi_Maonan_Huanjiang
Zhuang_Guangxi_Tibetan_Yajiang
Bouyei_Longli_Tibetan_Gangcha
Li_Qiongzhong_Mongol_Uuld
Bouyei_Longli_Nanai
Li_Qiongzhong_Mongol_Zasagt
Zhuang_Guangxi_Tibetan_Gangcha
Zhuang_Guangxi_Tibetan_Nagqu
Li_Qiongzhong_Qiang_Danba
Mulam_Luocheng_Tibetan_Gangcha
Li_Qiongzhong_Oroqen_HGDP
Bouyei_Longli_Tibetan_Chamdo
Tibetan_Chamdo_Mulam_Luocheng
Li_Qiongzhong_Ulchi
Zhuang_Guangxi_Tibetan_Chamdo
Maonan_Huanjiang_Tibetan_Shigatse
Maonan_Huanjiang_Tibetan_Shannan
Li_Qiongzhong_Nanai
Maonan_Huanjiang_Tibetan_Gangcha
Maonan_Huanjiang_Tibetan_Nagqu
Maonan_Huanjiang_Tibetan_Yajiang
Li_Qiongzhong_Rai_Nepal
Li_Qiongzhong_Gurung_Nepal
Dusun_Tibetan_Chamdo
Li_Qiongzhong_Tibetan_Shannan
Tibetan_Gangcha_Taiwan_Hanben
Maonan_Huanjiang_Tibetan_Chamdo
Li_Qiongzhong_Tibetan_Gangcha
Murut_Tibetan_Chamdo
Li_Qiongzhong_Tibetan_Lhasa
Li_Qiongzhong_Sherpa_Nepal
Li_Qiongzhong_Mongolia_EIA_1_SlabGrave
Ami_Tibetan_Gangcha
Li_Qiongzhong_Tibetan_Yajiang
Li_Qiongzhong_Tibetan_Shigatse
Li_Qiongzhong_Tibetan_Nagqu
Ami_Tibetan_Yajiang
Tibetan_Shigatse_Taiwan_Hanben
Tibetan_Yajiang_Taiwan_Hanben
Tibetan_Chamdo_Taiwan_Hanben
Li_Qiongzhong_Tibetan_Chamdo
Ami_Tibetan_Chamdo
-0.004
-0.002
0.000
f3(Source1, Source2; Han_Guiyang)
D
Bouyei_Longli_Japanese_Tokyo1K
Dai_Xishuangbanna1k_Yugur_Sunan
Japanese_Tokyo1K_Maonan_Huanjiang
Dai_Xishuangbanna1k_Han_Shandong
Kinh_HoChiMinh1K_Tibetan_Gangcha
Kinh_HoChiMinh1K_Tibetan_Yajiang
Dai_Xishuangbanna1k_Qiang_Daofu
Maonan_Huanjiang_Yugur_Sunan
Ulchi_Dai_Xishuangbanna1k
Kinh_HoChiMinh1K_Tibetan_Lhasa
Li_Qiongzhong_Yugur_Sunan
Dong_Guizhou_Tibetan_Gangcha
Zhuang_Guangxi_Qiang_Daofu
Daur_HGDP_Dai_Xishuangbanna1k
Dai_Xishuangbanna1k_Tibetan_Yajiang
Nanai_Dai_Xishuangbanna1k
Dai_Xishuangbanna1k_Tibetan_Gangcha
Dai_Xishuangbanna1k_Tibetan_Shigatse
Kinh_HoChiMinh1K_Tibetan_Chamdo
Li_Qiongzhong_Kalmyk
Dong_Guizhou_Tibetan_Chamdo
Zhuang_Guangxi_Tibetan_Shannan
Bouyei_Longli_Tibetan_Yajiang
Zhuang_Guangxi_Tibetan_Shigatse
Dai_Xishuangbanna1k_Tibetan_Lhasa
Zhuang_Guangxi_Tibetan_Yajiang
Li_Qiongzhong_Mongol_Uuld
Tuvinian_Maonan_Huanjiang
Bouyei_Longli_Tibetan_Gangcha
Zhuang_Guangxi_Tibetan_Gangcha
Dai_Xishuangbanna1k_Tibetan_Nagqu
Bouyei_Longli_Tibetan_Lhasa
Li_Qiongzhong_Tuvinian
Li_Qiongzhong_Yakut_HGDP
Li_Qiongzhong_Qiang_Daofu
Li_Qiongzhong_Qiang_Danba
Li_Qiongzhong_Daur_HGDP
Maonan_Huanjiang_Qiang_Daofu
Li_Qiongzhong_Ulchi
Dai_Xishuangbanna1k_Tibetan_Chamdo
Zhuang_Guangxi_Tibetan_Nagqu
Mulam_Luocheng_Tibetan_Gangcha
Mulam_Luocheng_Tibetan_Lhasa
Zhuang_Guangxi_Tibetan_Lhasa
Maonan_Huanjiang_Tibetan_Shannan
Li_Qiongzhong_Nanai
Maonan_Huanjiang_Tibetan_Shigatse
Maonan_Huanjiang_Tibetan_Gangcha
Bouyei_Longli_Tibetan_Chamdo
Maonan_Huanjiang_Tibetan_Lhasa
Maonan_Huanjiang_Tibetan_Yajiang
Zhuang_Guangxi_Tibetan_Chamdo
Tibetan_Chamdo_Mulam_Luocheng
Maonan_Huanjiang_Tibetan_Nagqu
Li_Qiongzhong_Gurung_Nepal
Li_Qiongzhong_Tibetan_Shannan
Li_Qiongzhong_Tibetan_Gangcha
Li_Qiongzhong_Tibetan_Yajiang
Tibetan_Gangcha_Taiwan_Hanben
Li_Qiongzhong_Tibetan_Shigatse
Ami_Tibetan_Gangcha
Li_Qiongzhong_Mongolia_EIA_1_SlabGrave
Maonan_Huanjiang_Tibetan_Chamdo
Li_Qiongzhong_Tibetan_Nagqu
Li_Qiongzhong_Tibetan_Lhasa
Tibetan_Yajiang_Taiwan_Hanben
Tibetan_Shigatse_Taiwan_Hanben
Li_Qiongzhong_Tibetan_Chamdo
Tibetan_Chamdo_Taiwan_Hanben
Ami_Tibetan_Chamdo
-0.006
-0.004
-0.002
0.000
f3(Source1, Source2; Han_Zunyi)

**Supplementary Fig. 13. Results of $f_3$-*statistics* for Guiyang and Zunyi Han.**
(A) The shared alleles between Guiyang Han and reference modern populations measured by *$f_3$(Han_Guiyang, Reference modern population; Yoruba);* (B) The shared alleles between Zunyi Han and reference modern populations measured by *$f_3$(Han_Zunyi, Reference modern population; Yoruba)*; (C)The potential source populations of Guiyang Han measured by *$f_3$(Source1, Source2; Han_Guiyang)*; (D)The potential source populations of Zunyi Han measured by *$f_3$(Source1, Source2; Han_Zunyi)*.

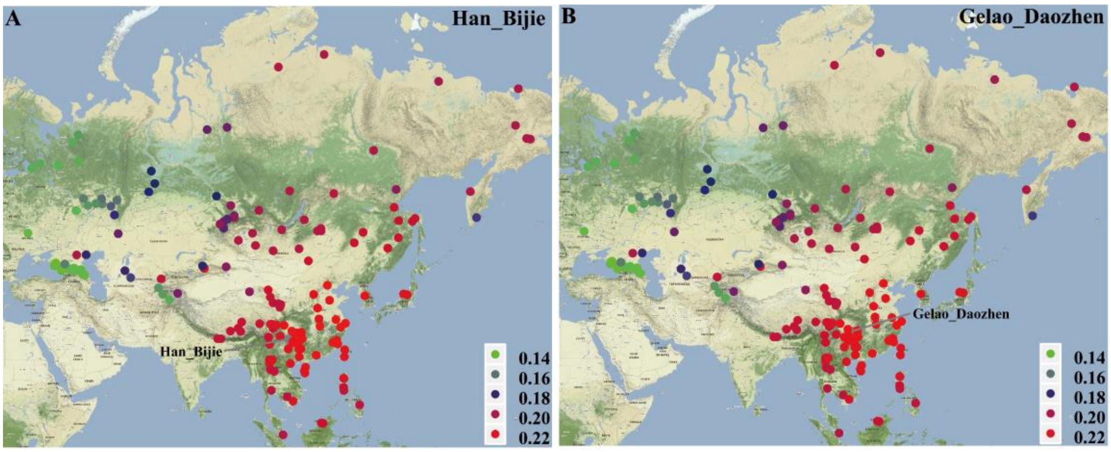

**Supplementary Fig. 14. Results of outgroup $f_3$-*statistics* for Bijie Han and Daozhen Gelao.**
(A) The shared alleles between Bijie Han and reference modern populations measured by *$f_3$(Han_Bijie, Reference modern population; Yoruba)*; (B) The shared alleles between Daozhen Gelao and reference modern populations measured by *$f_3$(Gelao_Daozhen, Reference modern population; Yoruba)*.

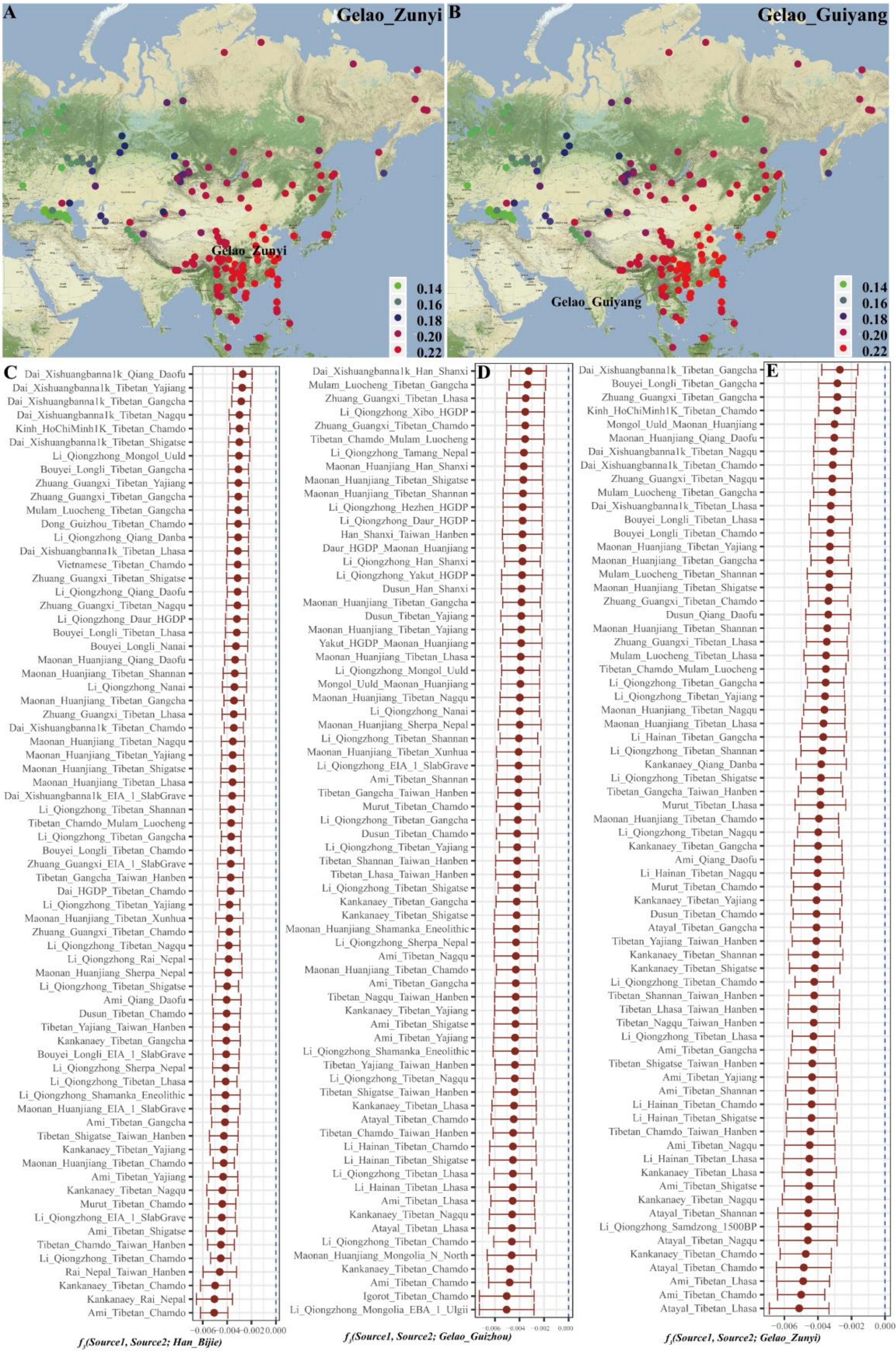

**Supplementary Fig. 15. Results of $f_3$-*statistics* for Bijie Han, Guiyang, Zunyi and Guiyang Gelaos.** (A) The shared alleles between Zunyi Gelao and reference modern populations measured by $f_3$*(Gelao_Zunyi, Reference modern population; Yoruba)*; (B) The shared alleles between Guiyang Gelao and reference modern populations measured by $f_3$*(Gelao_Guiyang, Reference modern population; Yoruba)*; (C)The potential source populations of Bijie Han measured by $f_3$*(Source1, Source2; Han_Bijie)*;

(D) The potential source populations of Guizhou Gelao measured by $f_3$*(Source1, Source2; Gelao_Guizhou)*; (E) The potential source populations of Zunyi Gelao measured by $f_3$*(Source1, Source2; Gelao_Zunyi)*.

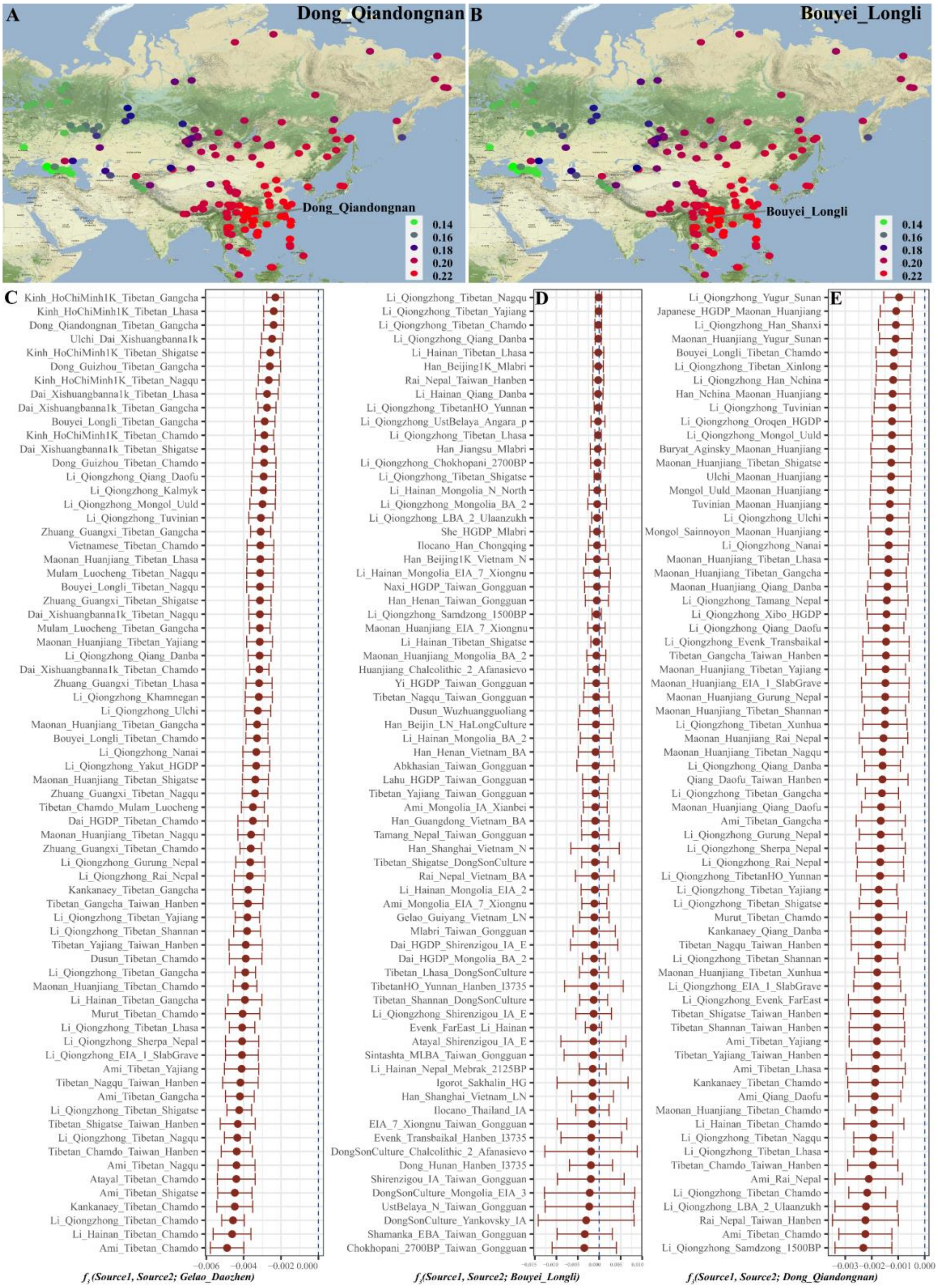

**Supplementary Fig. 16. Results of $f_3$*-statistics* for Dong, Bouyei and Daozhen Gelao.**

(A) The shared alleles between Qiandongnan Dong and reference modern populations measured by $f_3$*(Dong_Qiandongnan, Reference modern population; Yoruba)*; (B) The shared alleles between Longli Bouyei and reference modern populations measured by $f_3$*(Bouyei_Longli, Reference modern population; Yoruba)*; (C) The potential source populations of Daozhen Gelao measured by $f_3$*(Source1, Source2;*

*Gelao_Daozhen)*; (D) The potential source populations of Longli Bouyei measured by *f₃(Source1, Source2; Bouyei_Longli)*; (E) The potential source populations of Qiandongnan Dong measured by *f₃(Source1, Source2; Dong_Qiandongnan)*.

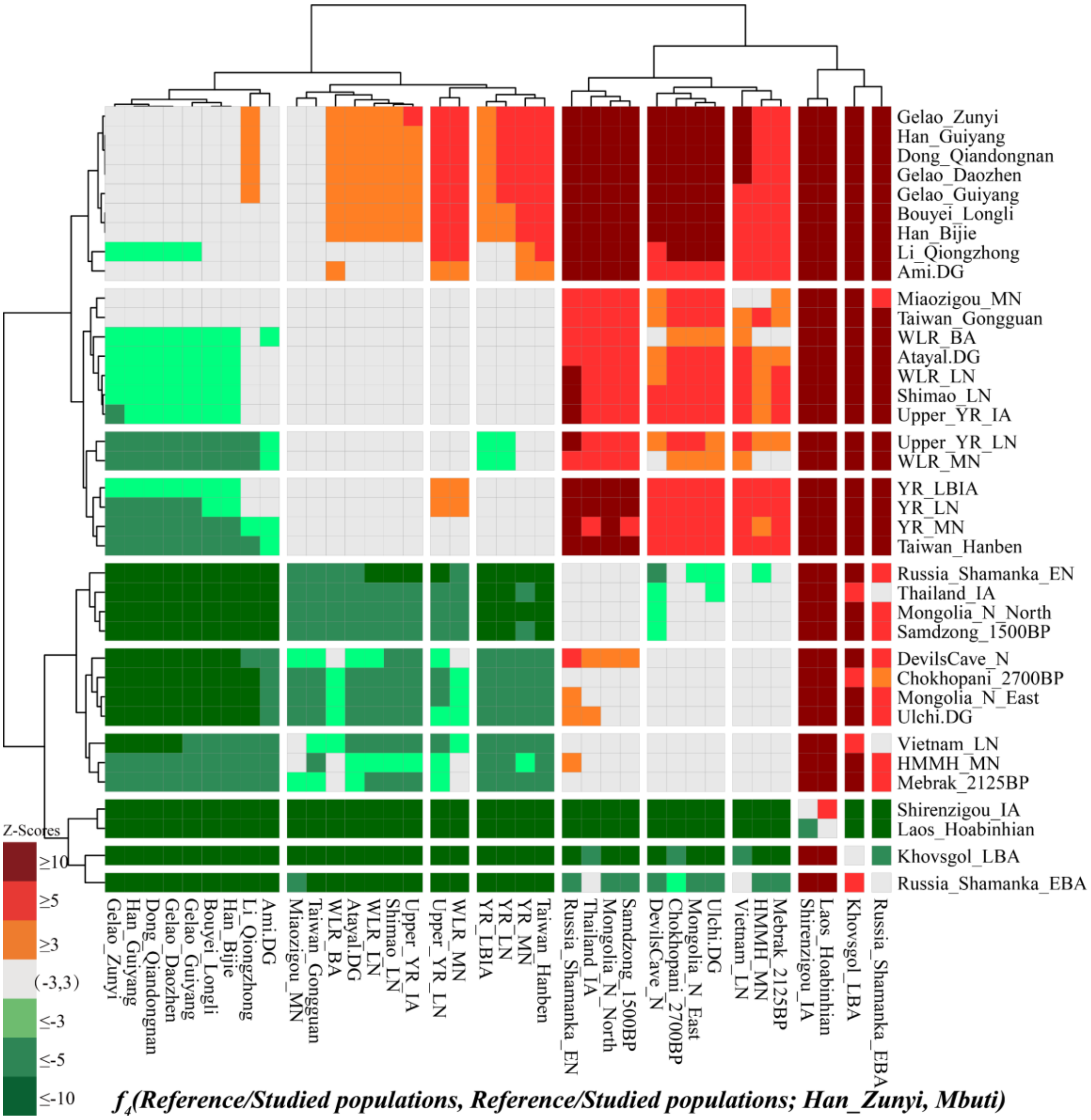

**Supplementary Fig. 17. The Z-scores of *f₄(Reference population1, Reference population2; Han_Zunyi, Mbuti)* revealed the allele sharing between Zunyi Han and reference populations.**

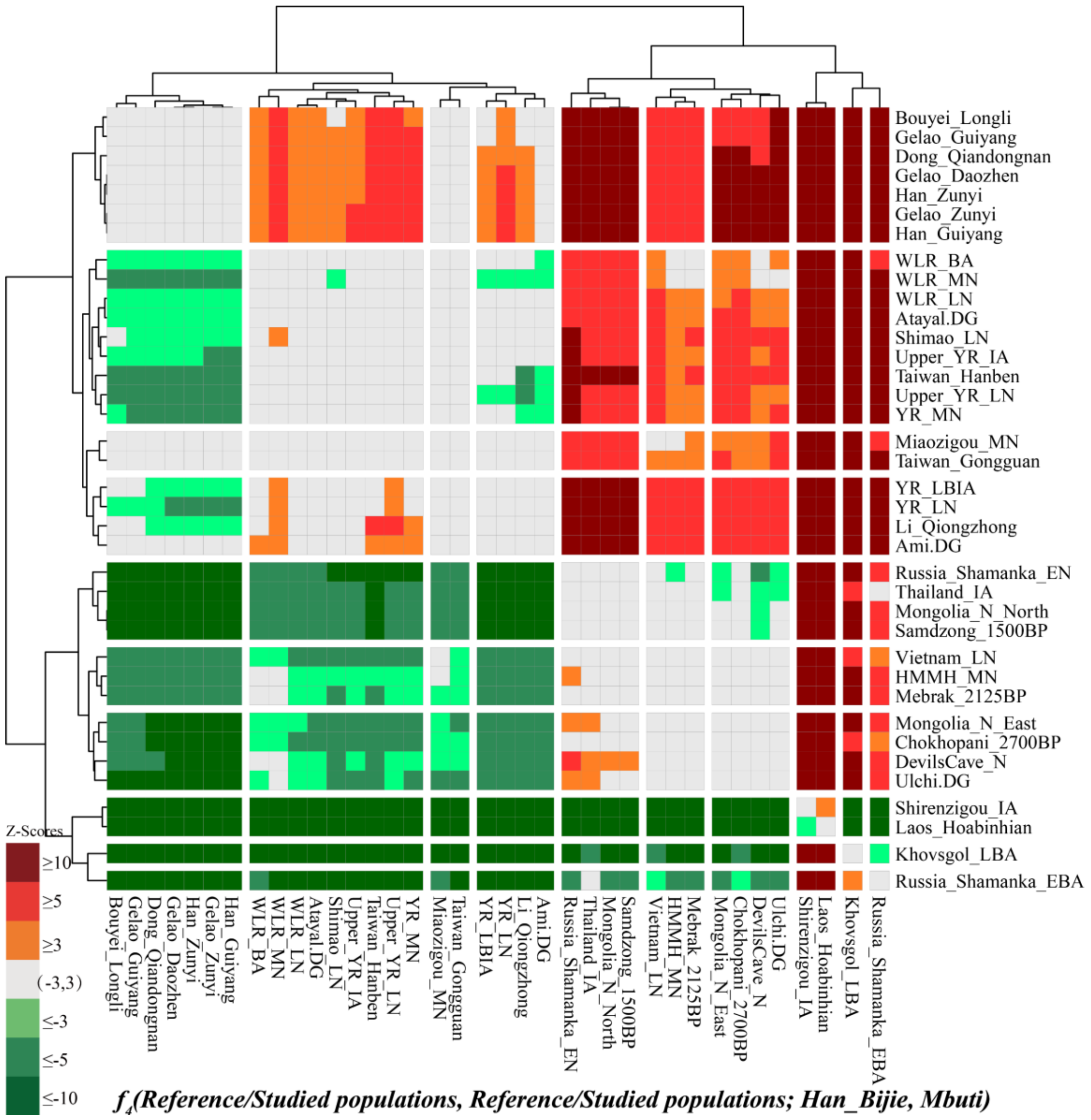

**Supplementary Fig. 18. The Z-scores of *$f_4$(Reference population1, Reference population2; Han_Bijie, Mbuti)* revealed the allele sharing between Bijie Han and reference populations.**

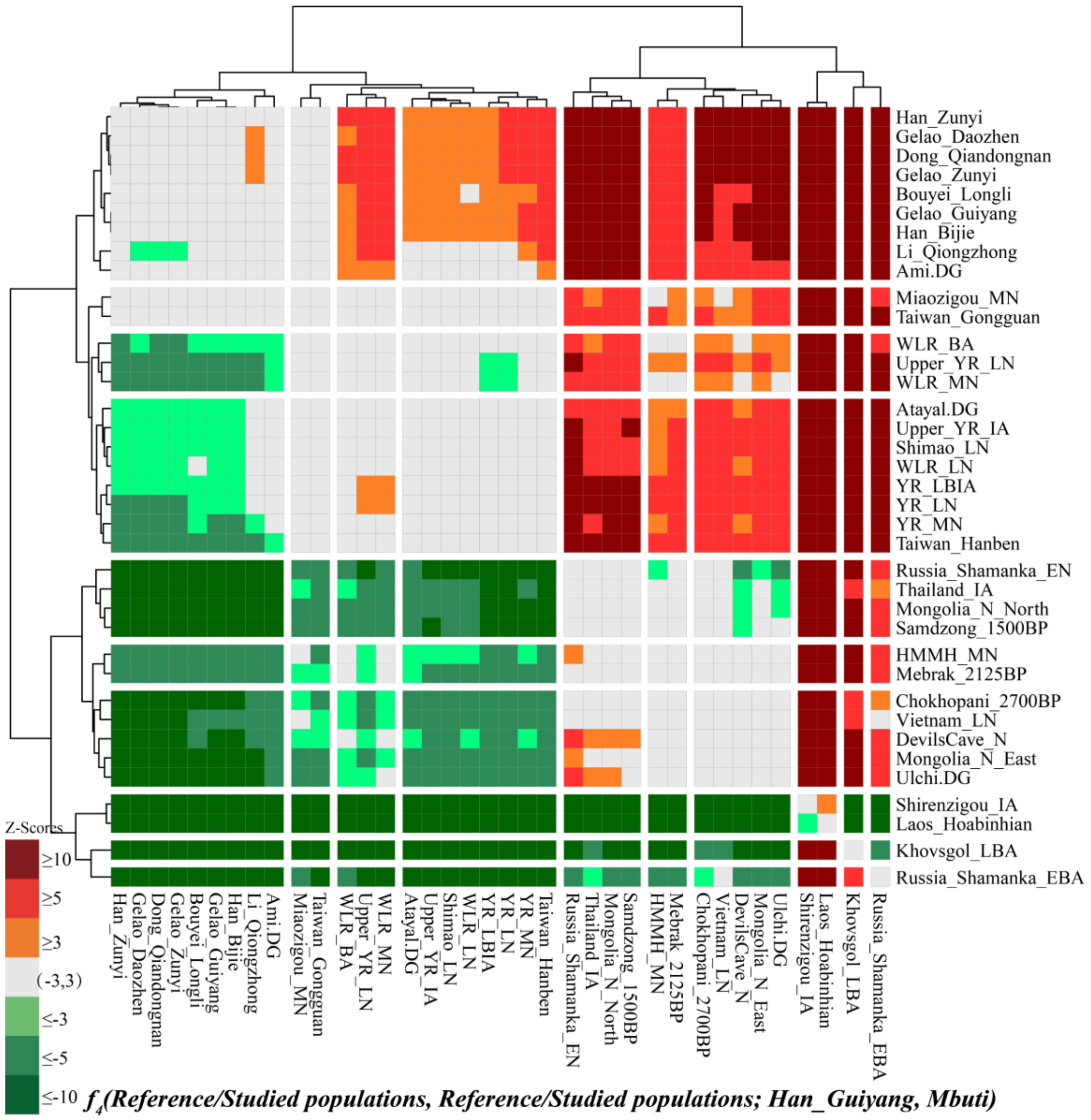

**Supplementary Fig. 19. The Z-scores of *$f_4$(Reference population1, Reference population2; Han_Guiyang, Mbuti)* revealed the allele sharing between Guiyang Han and reference populations.**

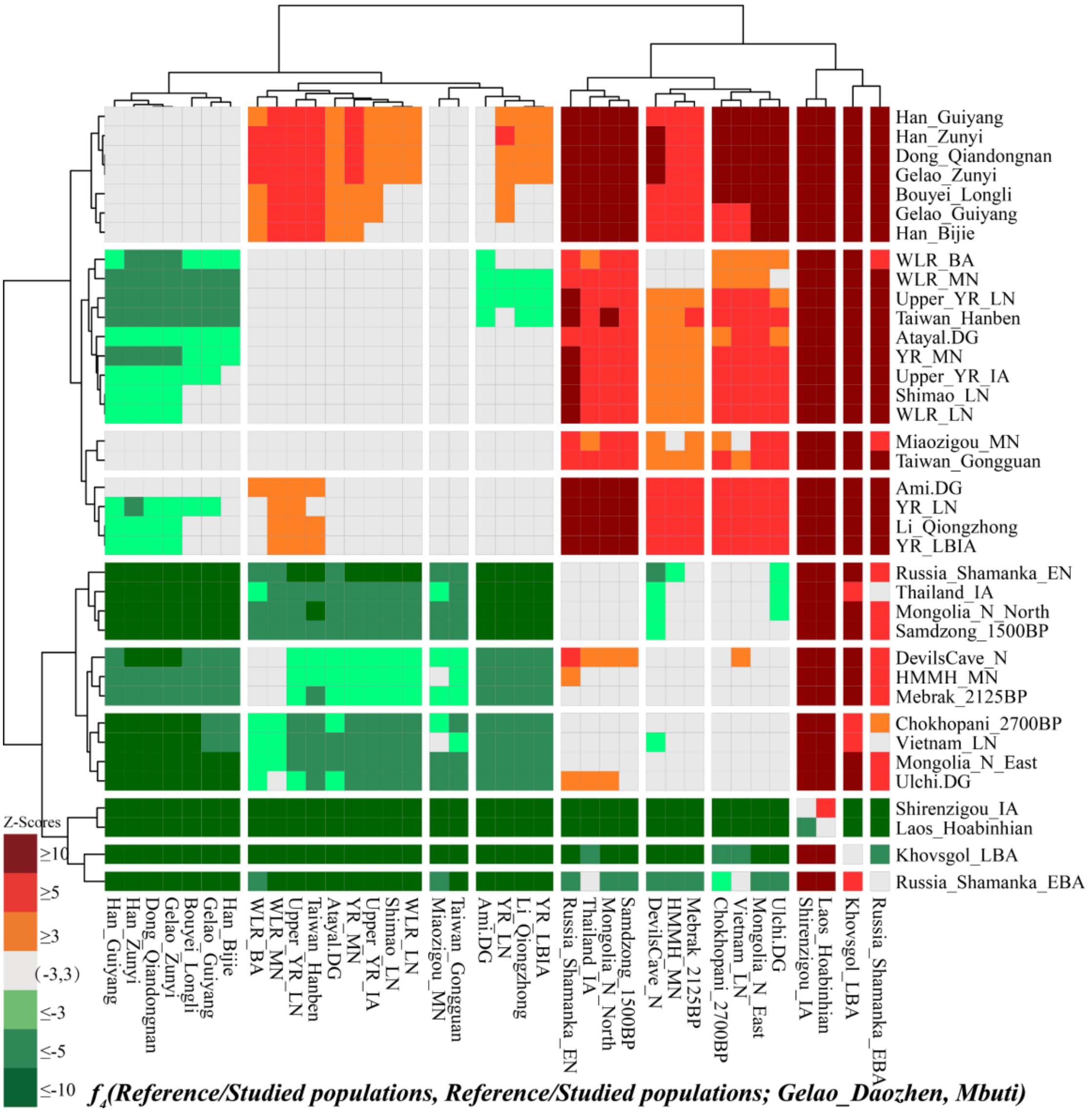

**Supplementary Fig. 20. The Z-scores of $f_4$*(Reference population1, Reference population2; Gelao_Daozhen, Mbuti)* revealed the allele sharing between Daozhen Gelao and reference populations.**
